# DISTINCT METABOLIC STATES DIRECT RETINAL PIGMENT EPITHELIUM CELL FATE DECISIONS

**DOI:** 10.1101/2023.09.26.559631

**Authors:** J. Raúl Perez-Estrada, Jared A. Tangeman, Maeve Proto-Newton, Harshavardhan Sanaka, Byran Smucker, Katia Del Rio-Tsonis

## Abstract

During tissue regeneration, proliferation, dedifferentiation, and reprogramming are necessary to restore lost structures. However, it is not fully understood how metabolism intersects with these processes. Chicken embryos can regenerate their retina through retinal pigment epithelium (RPE) reprogramming when treated with fibroblast factor 2 (FGF2). Using transcriptome profiling, we uncovered extensive regulation of gene sets pertaining to proliferation, neurogenesis, and glycolysis throughout RPE-to-neural retina reprogramming. By manipulating cell media composition, we determined that glucose, glutamine, or pyruvate are sufficient to support RPE reprogramming identifying glycolysis as a requisite. Conversely, the induction of oxidative metabolism by activation of pyruvate dehydrogenase induces Epithelial-to-mesenchymal transition (EMT), while simultaneously blocking the activation of neural retina fate. We also identify that EMT is partially driven by an oxidative environment. Our findings provide evidence that metabolism controls RPE cell fate decisions and provide insights into the metabolic state of RPE cells, which are prone to fate changes in regeneration and pathologies, such as proliferative vitreoretinopathy.

## INTRODUCTION

Metabolic activity has shown to be a crucial cellular attribute that can be altered throughout many cellular and molecular functions, such as cell signaling, cell proliferation, cell differentiation, and cell reprogramming (Agathocleous and Harris, 2013; Wu et al., 2016). Reprogramming somatic cells into induced pluripotent stem cells (iPSC) using Yamanaka factors (Oct3/4, Sox2, Klf4, c-Myc) has upended modern biology and opened new avenues for the development of cell therapies, including tissue replacement therapies (Shi et al., 2017; Takahashi and Yamanaka, 2006). Considerable molecular alterations, such as the activation/repression of signaling pathways, epigenetic arrangements, and metabolic remodeling, occur during the process of a somatic cell becoming an iPSC (Polo et al., 2012; Takahashi and Yamanaka, 2006). Changes in metabolic activity are one of the major molecular alterations that occur in response to the reprogramming factors, resulting in a switch from oxidative phosphorylation (OXPHOS) to glycolytic metabolism as cells transition to the pluripotent state (Teslaa and Teitell, 2014; Wu et al., 2016; Panopoulos et al., 2011). This metabolic switch is required for cell reprogramming, as glycolysis inhibition and OXPHOS activation cause a decrease in reprogramming efficiency, whereas inversely, glycolysis activation and OXPHOS inhibition promote pluripotency (Sun et al., 2020; Panopoulos et al., 2011). These observations suggest that any cell that undergoes reprogramming or a fate change might experience a deterministic metabolic rearrangement. In this regard, during regeneration, the reprogramming of resident cells is critical for recovering tissues or body parts that have been lost (Min and Whited, 2023; Barbosa-Sabanero et al., 2012; Jopling et al., 2011; Chiba, 2014; Wan and Goldman, 2016). However, how does metabolism affect cell reprogramming in the context of regeneration? This question remains poorly studied.

One of the best examples of cell reprogramming during regeneration is blastema formation. In many organisms, following the loss of body parts (e.g., limbs, tails, fins, etc.), a mass of cells, called *the blastema,* is formed around the injured area (Min and Whited, 2023; Jopling et al., 2011). The blastema is necessary for many forms of epimorphic regeneration and is formed by the reprogramming of cells surrounding the injured area, which transit to a less differentiated state and then proliferate and differentiate to form a new regenerated structure. Likewise, retina regeneration in several vertebrates also requires cell reprogramming. For example, in zebrafish, the neural retina regenerates through the reprogramming of the Müller glia, while in newts, retina regeneration proceeds through retinal pigment epithelium (RPE) reprogramming (Chiba, 2014; Wan and Goldman, 2016; Barbosa-Sabanero et al., 2012). Interestingly, embryonic amniotes can also regenerate the neural retina through RPE reprogramming, if stimulated with fibroblast growth factor 2 (FGF2) at the time of injury (Spence et al., 2007, 2004). In the chicken, RPE reprogramming can take place at day 4 of development (Hamburger-Hamilton [HH] Stage 24, (Hamburger and Hamilton, 1951)) but by day 5 (HH Stage 27), then RPE loses this ability (Tangeman et al., 2022; Sakami et al., 2008). Cell proliferation, epigenetic remodeling, and the activation of neural genes have been identified as cellular processes that are required to facilitate RPE reprogramming in embryonic chicken (Tangeman et al., 2022; Luz-Madrigal et al., 2020; Tangeman et al., 2021; Luz-Madrigal et al., 2014).

Despite these findings, the metabolic foundations of embryonic RPE reprogramming remain relatively unexplored. Our recent study highlighted the activation of retinol metabolism concurrent with RPE fate restriction, suggesting that the metabolic pathways associated with RPE maturation may limit reprogramming competence (Tangeman et al., 2022). However, the metabolic requirements of RPE cells as they dedifferentiate and change identity during reprogramming are less clear. In the adult eye, there is an intimate coupling of the metabolic states of the RPE and neural retina, and disruption of this balance can drive pathologies such as retinitis pigmentosa and RPE degeneration (Nolan et al., 2022). Thus, it is likely that metabolism plays an instrumental role in establishing and maintaining the RPE lineage during development and throughout life. Given the centrality of metabolism in RPE fate and function, there remains a notable deficit in our understanding of how RPE metabolism determines cell identity, physiology, dysregulation/degeneration, and critically, regenerative competence.

To address these questions, we used the embryonic chicken to dissect the role of metabolism during RPE reprogramming. Using an RPE explant system that facilitates a precise modulation of the metabolic environment, we identified glycolysis as a pathway that is essential for cell reprogramming. In addition, we found that the metabolic reprogramming of RPE from a glycolytic to oxidative metabolism provokes the activation of an epithelial-mesenchymal transition (EMT) program that redirects the cells toward a mesenchymal fate. Importantly, we demonstrate that driving OXPHOS during RPE reprogramming blocks the activation of genes essential for neural retina development and prevents proliferation. Our results provide newfound knowledge regarding the role of metabolism in determining the competence of RPE cells to reprogram into neural retina. We hope these results may provide a foundation for understanding metabolic dysregulation in the pathogenesis of eye diseases, such as PVR, RP, and age-related macular degeneration (AMD).

## RESULTS

### RPE explants reprogram into neural retina

The cellular reprogramming of RPE into neural retina is induced by FGF2 in chicken embryos, (Tangeman et al., 2022; Luz-Madrigal et al., 2014; Spence et al., 2004). However, the limited period of time and accessibility to the *in vivo* system has complicated the elucidation of the metabolic determinants of RPE reprogramming. RPE can be cultured as an explant and reprogrammed into neural retina in the presence of FGF2 (Sakami et al., 2008; Tangeman et al., 2022), but the reprogramming process has not been studied in detail ex vivo. To better understand the RPE explant model, we first performed a phenotypic characterization. RPE sheets were collected from E4 (HH stage 24) embryos and cultured in the presence or absence of FGF2 (Fig. 1 A). By 24h of cell culture, the explants started to show signs of neuroepithelium formation, which became more evident at 48h and well-defined by 96h of culture (Fig. 1 B). *In vivo*, we know that FGF2 is essential for inducing RPE reprogramming, however, it has not been possible to determine whether sustained FGF2 treatment is necessary for reprogramming and neuroepithelial growth. To test this, we removed FGF2 at 1h or 6h after the initiation of the culture. Surprisingly, an hour of FGF2 treatment was enough to induce RPE reprogramming (Fig. S1 A), suggesting that FGF2 is a trigger of cell reprogramming, but that its sustained presence is not necessary for neuroepithelium formation. Similar to in vivo, RPE explants can reprogram when collected at E4, but lose neurocompetence at embryonic day 5 (E5) (Tangeman et al., 2022; Sakami et al., 2008). In addition, E4 RPE explants lose neurocompetence by 48h of culture, as RPE retains an ability to continue differentiating ex vivo (Sakami et al., 2008). However, the exact timepoint that reprogramming competence is lost in culture has not been determined. Therefore, we added FGF2 to E4 explants at different time points in the culture (Fig. S1 B). We found that after 24h, RPE was still able to reprogram, however, after 48h of culture, RPE did not reprogram, demonstrating that RPE progressively loses reprogramming competence between 24h and 48h of culture and is extended at least 24h relative to in vivo.

**Figure 1.**
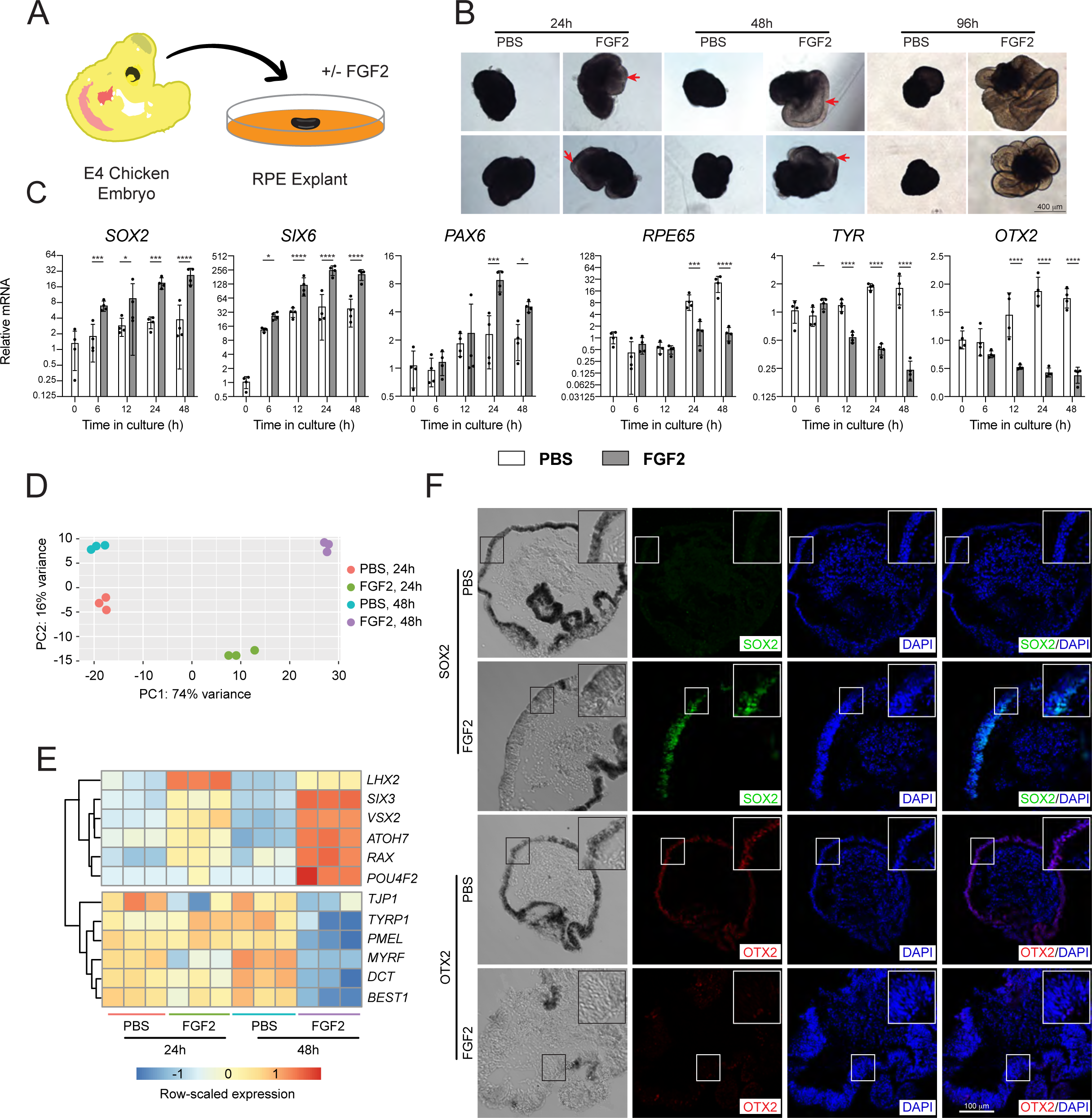
RPE Explants reprogram into neural retina. (A) RPE sheets were collected from chicken embryos at E4 and cultured for up to 4 days. (B) Representative RPE explants at 24h, 48h, and 96h of cell culture. (C) The expression of neural genes *SOX2*, *SIX6*, and *PAX6*, as well as RPE genes *RPE65*, *TYR*, and *OTX2* were measured by RT-qPCR from explants collected at indicated time points. (D) Variance stabilizing transformation was applied to RNAseq normalized counts and principal component analysis was performed, resulting in a clear spatial separation of samples by condition. (E) Row-normalized heatmap displays expression patterns of RPE identity factors (top) and neural retina transcription factors (bottom) in response to FGF2. (F) Immunofluorescence detection of SOX2 and OTX2 in explants collected at 48h of cell culture. Plots show group mean ±SD; p-values from modeling as described in Statistical analysis section. *p<0.05, **p<0.01, *** p<0.001 and **** p<0.0001.

To characterize RPE reprogramming ex vivo at the molecular level, we collected explants at different time points during cell culture (Fig. S1 C) for gene expression analysis of the neural retina markers *SOX2*, *PAX6*, and *SIX6*, and the RPE markers *RPE65*, *TYR*, and *OTX2*. We found that FGF2 treatment induced upregulation of *SOX2* and *SIX6* by 6h, while *PAX6* was upregulated by 24h (Fig. 1 C). In contrast, RPE markers were acutely downregulated in the presence of FGF2 (Fig. 1 C). Downregulation of *TYR* and *OTX2* was observed after 12h of culture, while *RPE65* was significantly downregulated after 24h (Fig. 1 C). Interestingly, in PBS-treated controls, *RPE65* and *OTX2* abundance increased through 48h (Fig. 1 C), indicating RPE maturation. Based on these results, we collected another set of explants at 24h and 48h for RNAseq (Fig. S2). Principal component analysis (PCA) showed that the sequenced samples clustered by condition, underscoring a high concordance between biological replicates and treatment effect (Fig. 1 D). Normalized count plots for retinal and RPE markers were plotted from the RNAseq data, showing the same regulatory patterns for genes previously assessed via RT-qPCR (Fig. 1 C and Fig. S3 A). As expected, FGF2 treatment led to an increased abundance of genes encoding eye field transcription factors (*RAX*, *SIX3*, *LHX2*), which is consistent with previous reports that FGF2 pushes RPE to dedifferentiate toward an early eye field identity (Tangeman et al., 2021, 2022; Luz-Madrigal et al., 2014). Similarly, neural retina specification marker (*VSX2*) was elevated, and retinal ganglion cell determinants (*POU4F2*, *ATOH7*) were elevated by 48h, suggesting the onset of fate determination within the reprogrammed neuroepithelium. In contrast, RPE genes were downregulated in the presence of FGF2 by 48h (Fig. 1 E). In addition, genes associated with the term Generation of neurons (GO:0022008) were also upregulated in reprogramed explants (Fig. S3 B). To confirm the gene expression results, immunohistochemistry was performed for SOX2 and OTX2 in explants collected at 48h of culture (Fig. 1 D). Explants treated with FGF2 showed a positive signal for SOX2, which was absent from untreated explants (Fig. 1 F). In contrast, the explants that were not exposed to FGF2 were OTX2-positive, whereas in FGF2-treated explants OTX2 abundance was low and mostly restricted to pigmented areas (Fig 1 F). Altogether, these data show that the molecular reprogramming of RPE into neural retina is observable at 24h and is robustly detectable by 48h.

### Cell proliferation is a tightly regulated process during RPE reprogramming

From the RNAseq experiment, we detected 19194 genes, of which 3682 were differentially expressed genes (DEGs), fitting the criteria log fold change ≥ 1 & adjusted p-value ≤ 0.05 (Fig. 2 A). The top 10 DEGs that were repressed or activated by FGF2 at 24h and 48h are shown in Figure 2B. We found that all the FGF2-repressed genes at 24 and 48h were different, whereas, amongst the top FGF2-activated genes, *FEZF2*, *KK34*, and *NOS1* were commonly observed at 24h and 48h. *FEZF2* encodes a transcription factor required for neuronal differentiation and specification (Tsyporin et al., 2021; Zhang et al., 2014), while *KK34* encodes an interleukin-like protein (Koskela et al., 2004) and *NOS1* for nitric oxide 1, which is mainly expressed in the brain (Förstermann and Kleinert, 1995). Interestingly, many FGF2-activated genes encode transcription factors such as FEZF2, HMX3, POU3F1, PTF1A, and NHLH2. In contrast, FGF2-repressed genes encode a wide variety of proteins, some of which are involved in cell signaling and transcriptional activity. To identify the possible pathways and cellular processes that act as determinants for RPE reprogramming, we performed pathway enrichment analysis using the differentially expressed genes classified as FGF2-activated or FGF2-repressed genes, at both 24h and 48h (Fig. 2 B). The regulated processes included MAPK cascade, mesenchyme differentiation, ROS metabolic process, DNA metabolic process, and cell cycle (Fig. 2 B). Previously, it was reported that during RPE reprogramming, cells undergo proliferation to form a neuroepithelium (Luz-Madrigal et al., 2020), and that RPE loses proliferative ability concomitant with the loss of neural competence (Tangeman et al., 2022). Accordingly, in RPE explants, genes associated with the cell cycle (KEGG:04110) were robustly upregulated by FGF2 at 24h and 48h (Fig. 2 C). Using EdU incorporation and phospho-histone H3 (pHH3) staining to label the cells that underwent phases S and M of the cell cycle, we confirmed that in FGF2-treated explants cell proliferation was highly increased during cell reprogramming (Fig. 2, D and E). Taken together, these data confirm that cell proliferation is one of the major cellular processes associated with RPE reprogramming ex vivo.

**Figure 2.**
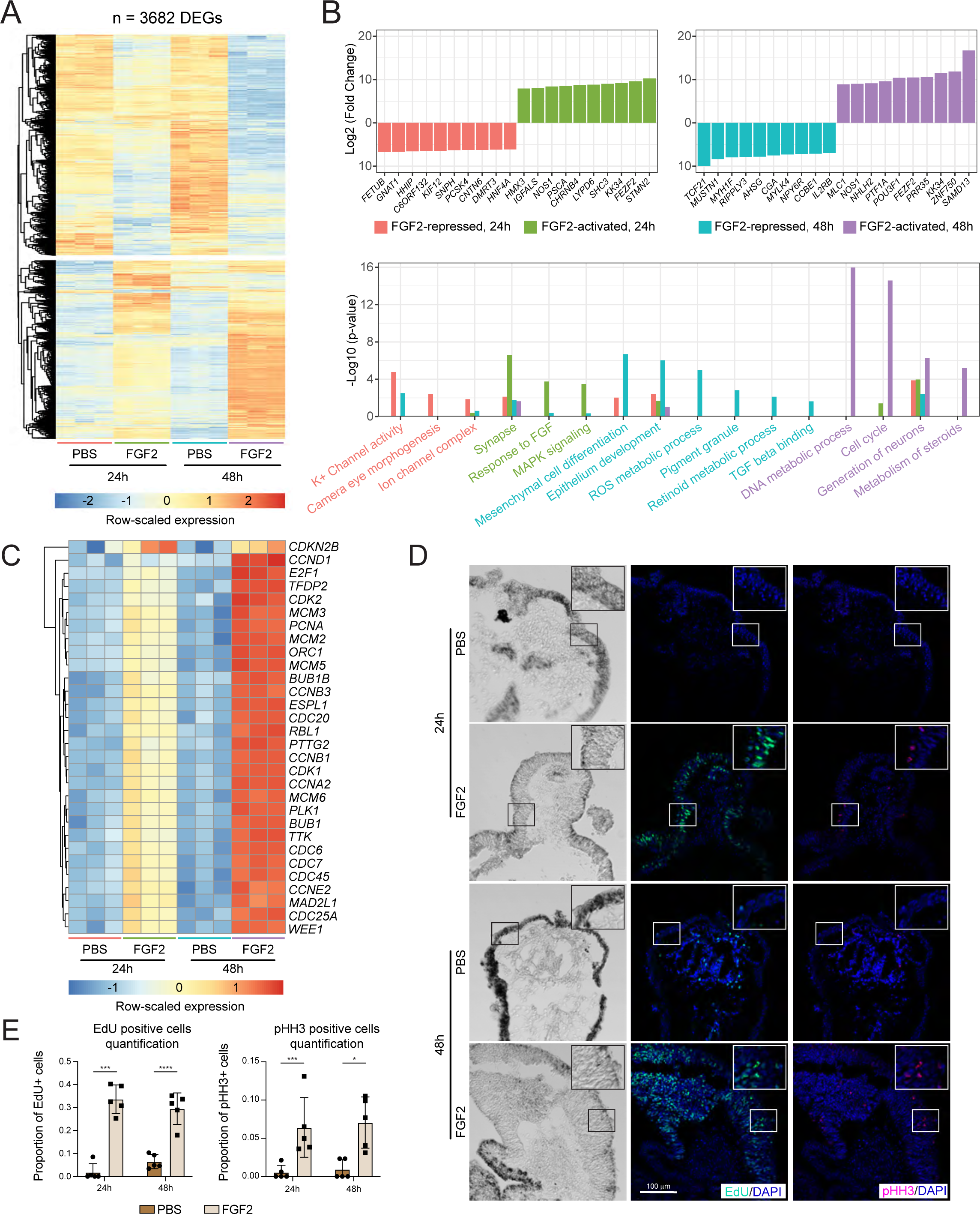
Cell proliferation is highly activated during RPE reprogramming. (A) Heatmap displays row-normalized expression patterns of 3682 identified DEGs, defined by the criteria |log_2_(fold change)| ≥ 1 and adjusted p-value ≤ 0.05. (B) Upper panel: The top 10 up- and down-regulated DEGs, as ranked by log-fold change, are displayed. Log fold change values represent expression change in response to FGF2 relative to PBS, after 24h or 48h in culture. Lower panel: Pathway enrichment analysis was used to assign biological functions to up- and downregulated gene sets in response to FGF2. The adjusted p-value for select terms are displayed in bar charts. (C) The top 30 upregulated genes associated with the term cell cycle (KEGG:04110) are displayed in a row-normalized heatmap. (D) EdU and pHH3 staining were performed in explants at 24h and 48h of cell culture. (E) quantification of EdU or pHH3 positive cells from (D). Plots show group mean ±SD; p-values from modeling as described in Statistical analysis section. *p<0.05, **p<0.01, *** p<0.001 and **** p<0.0001.

### Glycolysis is necessary for RPE reprogramming

Cells with high proliferative activity, such as a wide variety of stem cells, and progenitors, including neural progenitors, and cancer cells, demand large amounts of energy that are provided by metabolic activity, especially glycolysis (Abdel-Haleem et al., 2017; Lunt and Heiden, 2011; DeBerardinis et al., 2008). Thus, we hypothesized that RPE requires glycolysis due to the high cell proliferation observed during its reprogramming. Glycolysis is a pathway of sequential biochemical reactions that produce ATP, pyruvate, and metabolites for amino acid synthesis (Lunt and Heiden, 2011). In contrast, gluconeogenesis results in the production of glucose (Hers and Hue, 1983). Glycolysis and gluconeogenesis share enzymes that catalyze reversible reactions, whereas rate-limiting enzymes are specific to each pathway. In glycolysis, hexokinases I and II (HKI, II), phosphofructokinase (PFK P/L), and pyruvate kinase (PKLR) are glycolytic-specific enzymes, whereas pyruvate carboxylase (PC), phosphoenolpyruvate carboxylase 1 (PCK1), fructose 1,6 biphosphatase 1 (FBP1), and glucose 6-phosphatase (G6P) are specific enzymes of gluconeogenesis (Fig. 3 A). RNAseq analysis revealed that the genes encoding enzymes essential for glycolysis and gluconeogenesis were differentially expressed in the RPE explants (Fig 3, A and B). Interestingly, genes encoding most glycolytic enzymes, including rate-limiting enzymes, apart from *ALDOB*, were upregulated in the presence of FGF2 (Fig. 3 B). *LDH A/B*, which encodes lactate dehydrogenases A and B, respectively, were also expressed and regulated during cell reprogramming. *LDHA* was upregulated at 48h, while *LDHB* was transiently upregulated at 24h (Fig. 3 B). To confirm these observations, the expression levels of selected metabolic genes were determined by RT-qPCR at different time points of RPE reprogramming (Fig. S4). Similar to the results in the RNAseq dataset, *HK1* was upregulated by FGF2 at 24h, while *HK2* was significantly upregulated at 12h (Fig. S4). *GADPH* was also upregulated in the presence of FGF2 (Fig. S4). RT-qPCR analysis confirmed that *LDHA* was positively regulated by FGF2, while *LDHB* was unaffected (Fig S4). LDHA and LDHB are mainly differentiated by their affinity for pyruvate and lactate; LDHA has higher affinity for pyruvate and LDHB for lactate, therefore, their differential expression might affect the glycolytic rate (Mishra and Banerjee, 2019; Read et al., 2001). Contrary to what has been observed in glycolytic genes, levels of *FBP1*, a gluconeogenic gene, were repressed by FGF2 at 48h (Fig. S4). Together these data suggest that glycolysis could be favored and required for RPE reprogramming but not gluconeogenesis.

**Figure 3.**
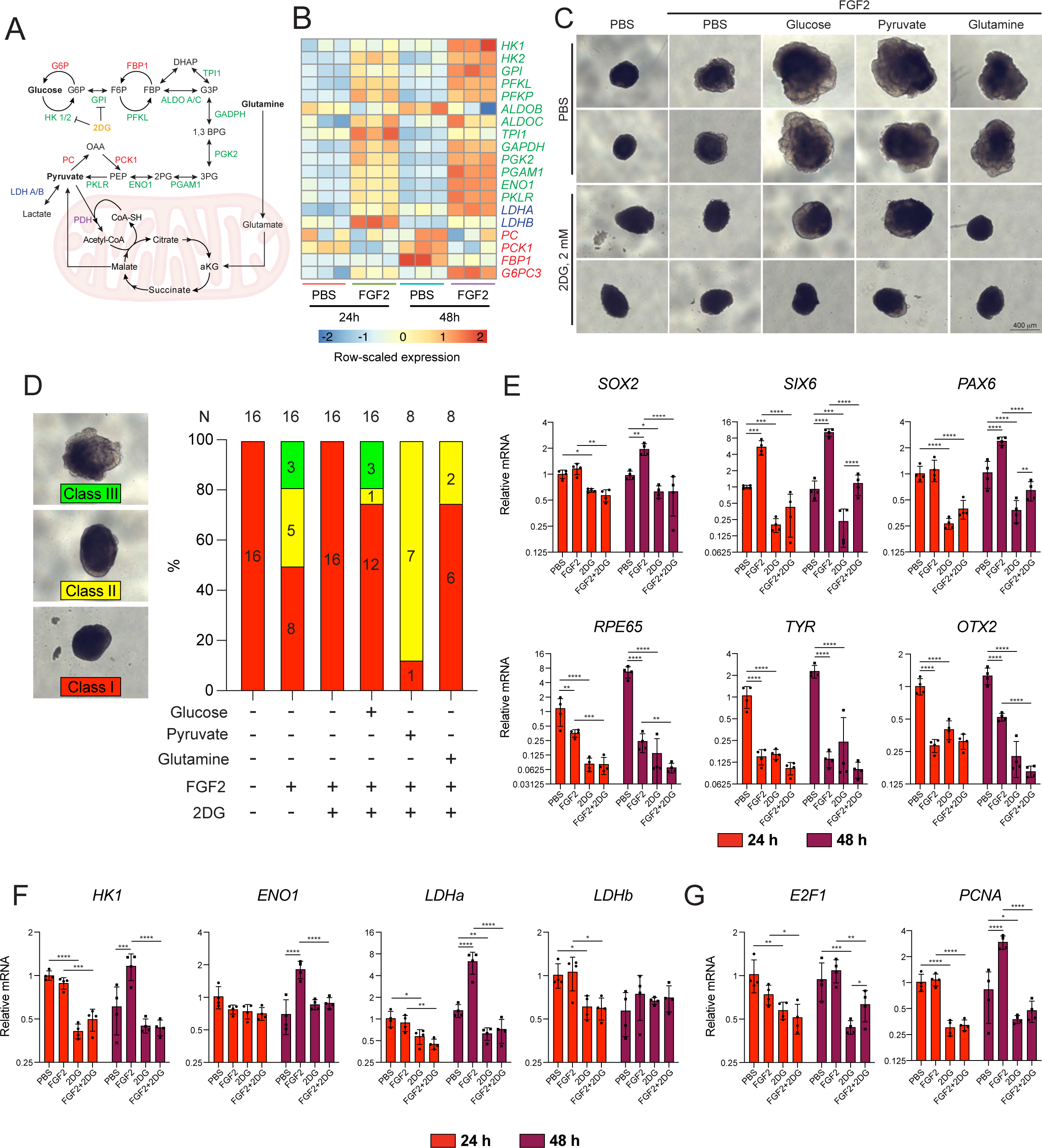
The glycolysis pathway is activated during RPE reprogramming and its inhibition affects cell reprogramming. (A) The glycolysis and gluconeogenesis pathway, including a pathway that glutamine can follow to enter glycolysis/gluconeogenesis. Green font indicates glycolytic enzymes, red font indicates gluconeogenesis-specific enzymes and blue font indicates lactic enzymes. (B) Heatmap displays row-normalized expression values of select genes with functions pertaining to glycolysis or gluconeogenesis, as well as lactic enzymes. (C) Representative RPE explants at 96h cultured in DMEM without glucose, glutamine, and pyruvate and supplemented with glucose, pyruvate, or glutamine at 2 mM, +/-FGF2, and +/- 2DG. (D) quantification of explant phenotype shown in (C). Class I: no reprogrammed RPE; Class II: signs of RPE reprogramming; Class III: reprogrammed RPE. Gene expression analysis of retina and RPE markers genes (E), and metabolic genes (F) of RPE explants cultured in DMEM without glucose, glutamine, and pyruvate and supplemented +/-glucose at 2 mM and/or +/-2DG at 2 mM and collected at indicated time points. Plots show group mean ±SD; p-values from modeling as described in Statistical analysis section. *p<0.05, **p<0.01, *** p<0.001 and **** p<0.0001.

To test whether glycolysis is required for RPE reprogramming, we used 2-Deoxy-D-Glucose (2DG), a glucose analog that inhibits glycolysis by inhibition of hexokinase and glucose-6-phosphate isomerase. 2DG up to 20 mM concentration did not inhibit RPE reprogramming, a phenotype confirmed by morphological examination, as well as gene expression determination of neural retina and RPE markers (Fig. S5, A and B). Since 2DG inhibits glycolysis in a competitive and non-competitive manner (Pajak et al., 2019; Ralser et al., 2008; Aft et al., 2002; Shiraishi et al., 2014), and the final concentration of glucose in the media was 68 mM, we hypothesized that 20 mM of 2DG might be insufficient to inhibit glycolysis. Therefore, we cultured RPE explants in DMEM modified without glucose or pyruvate, and glucose was added separately. At these conditions the RPE was still able to reprogram (Fig. S5 C), suggesting that the cells used a different metabolite as an energy source, such as glutamine. Therefore, RPE explants were cultured in media depleted of glucose, pyruvate, and glutamine. Under these conditions, RPE reprogramming was limited but was rescued when glucose, pyruvate, or glutamine was added at a concentration of 2 mM. Interestingly, the rescue of the reprogramming phenotype was inhibited by 2DG at equimolar concentrations (Fig. 3 C). To quantify the effect of 2DG, we classified RPE reprogramming based on phenotypic observations on day 4 of culture into three classes: I) not reprogrammed RPE, II) rudimentary signs of RPE reprogramming, and III) extensively reprogrammed RPE. In the absence of FGF2, control explants did not reprogram (16/16). In the presence of FGF2, explants class II (5/16) and class III (3/16) were detected even in the absence of glucose, glutamine, and pyruvate, suggesting that amino acids present in the medium might be used as energy and carbon sources. However, none of the explants (16/16) were reprogrammed when glycolysis was inhibited with 2DG (Fig. 3 D). In the presence of FGF2, glucose, and 2DG, most explants were class I (12/16), one was class II, and three were class III. Interestingly, most of the explants cultured in the presence of FGF2, pyruvate, and 2DG showed signs of reprogramming (class II, 7/8), one did not reprogram (class I), and none were classified as class III. On the other hand, in the presence of glutamine and 2DG, most of the explants were class I (6/8) and a few class II (2/8) (Fig. 3 D). When the concentration of glucose was 4 mM and 2DG 2 mM, no class I explants were detected; instead, most of the explants were class II (6/8) and a few class III (2/8), reflecting the competitive nature of 2DG (Fig S5, D and E). Similarly, when glutamine or pyruvate was added at 4 mM concentration and 2DG at 2 mM, only class II explants were detected (Fig. S5, D and E). The observation that 2DG inhibited RPE reprogramming in most of the cases at equimolar concentration, suggests that even when metabolites such as pyruvate, glutamine, and other amino acids can be used as carbon sources and energy production, probably through anaplerotic reactions, the metabolism of those metabolites still requires glycolysis, suggesting pyruvate or glutamine might be used to produce glucose through gluconeogenesis and then metabolized by glycolysis, establishing glucose as an essential metabolite for RPE reprogramming.

To corroborate the effects of 2DG on RPE reprogramming, we determined the gene expression levels of neural retina and RPE markers using RT-qPCR. The results confirmed that 2DG stymied the upregulation of *SOX2*, *SIX6*, and *PAX6* induced by FGF2 (Fig. 3 E). However, the expression of RPE markers was downregulated in explants treated with either FGF2 or 2DG alone, suggesting that glycolysis inhibition might also disrupt RPE identity (Fig. 3 E). Gene expression analysis of metabolic genes also showed that 2DG prevents the upregulation of glycolytic genes *HK1*, *HK2*, *TPI1*, *PGAM1*, *ENO1*, *PKLR*, and *LDHA* observed during cell reprogramming (Fig 3 F and Fig. S5 F). In addition, treatment with 2DG and FGF2 led to downregulation of the cell cycle genes *E2F1* and *PCNA* relative to FGF2-only explants, suggesting that 2DG may limit cell proliferation during reprogramming (Fig. 3 G). However, it is important to highlight that *E2F1* response to FGF2 was lower in comparison to RPE explants cultured in glucose at 68 mM concentration (Fig. 2 C). *E2F1* has been identified as a global regulator of metabolism including glucose metabolism (Denechaud et al., 2017), indicating glucose concentration (2 mM versus 68 mM) was responsible for those differences. Altogether, the data show that glycolysis is necessary for RPE to neural retina reprogramming and points toward glucose as a required metabolite.

### Activation of OXPHOS metabolism inhibits RPE reprogramming

Glycolysis inhibition demonstrated the importance of proper metabolism for RPE reprogramming. Therefore, we next investigated the effect of OXPHOS activation by promoting pyruvate dehydrogenase (PDH) activity using dichloroacetate (DCA). DCA is an inhibitor of pyruvate dehydrogenase kinase (PDK), which inactivates PDH by phosphorylation, controlling the catabolism of pyruvate through the tricarboxylic acid cycle (TCA) (Wang et al., 2021; Stacpoole and Greene, 1992; Michelakis et al., 2008; Rodrigues, 2015; Stacpoole, 1989; Takubo et al., 2013) (Fig. 4 A). We hypothesized that OXPHOS activation would inhibit RPE reprogramming, similar to what is observed during iPSC formation (Sun et al., 2020). DCA alone did not clearly affect the morphology of RPE explants at the concentrations tested, except for an apparent increase in the size of the explants when they were treated at 20 mM (Fig. 4 B). Interestingly, DCA inhibited RPE reprogramming in a concentration-dependent manner, with 100% inhibition at 20 mM (Fig. 4 B). However, the phenotypic characteristics of the explants differed from those treated with 2DG at an equimolar concentration (Fig. 3 C), as RPE explants treated with DCA at 20 mM were apparently larger than the control explants (Fig. 4 B). The inhibition of RPE reprogramming by DCA was also observed in RPE explants cultured in DMEM modified without glucose and pyruvate, a medium that only contains glutamine as the carbon source (Fig. S6 A), indicating that the effect depends only on OXPHOS activation, but not of the carbon source. H&E staining revealed a lack of a neuroepithelium formation in FGF+DCA treated explants; instead, an apparent disorganization of the RPE was noted even in the absence of FGF2 (Fig. S6 B, red arrows). The phenotypic observations were confirmed by RT-qPCR analyses. We observed overexpression of *SOX2* and *SIX6* induced by FGF2 in the presence of DCA, but their expression levels were significantly lower than those observed in FGF2 alone, except for *SOX2* at 24h. However, *PAX6* levels were mostly unaffected, and DCA alone upregulated its gene expression at 24h and 48h (Fig. 4 C). Interestingly, DCA treatment alone and in combination with FGF2, led to repression of *RPE65* and *TYR* whereas *OTX2* downregulation by FGF2 was not observed in presence of DCA (Fig. 4 C). These observations suggest that OXPHOS activation not only affected RPE reprogramming but also perturbed RPE identity.

**Figure 4.**
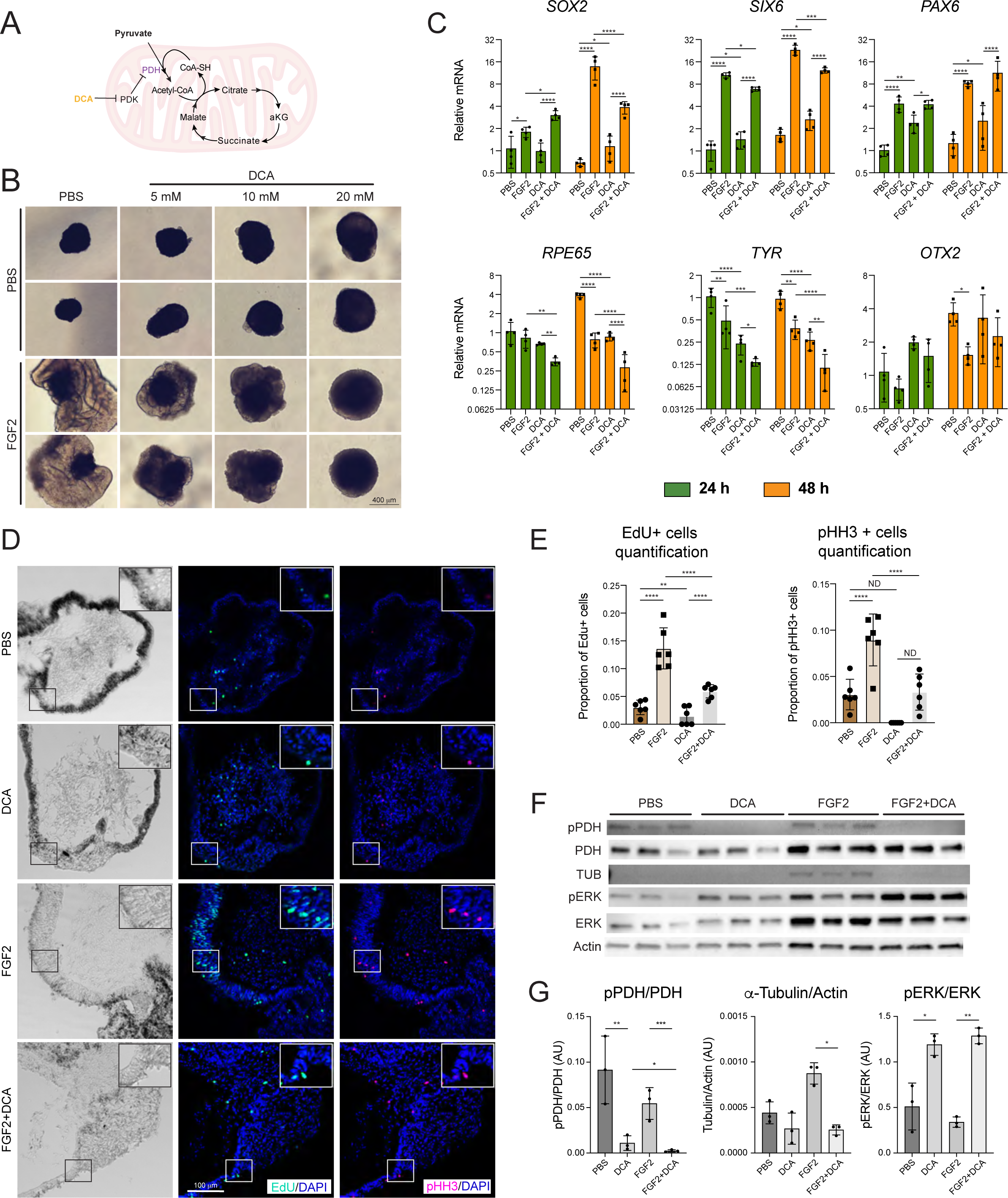
Activation of OXPHOS metabolism blocks RPE reprogramming, inhibits cell proliferation, and activates ERK. (A) The OXPHOS pathway and its activation by inhibition of PDK via DCA. (B) Representative RPE explants treated with FGF2 and DCA at varying concentrations after 96h of cell culture. (C) Levels of relative gene expression of neural genes *SOX2*, *SIX6*, and *PAX6*, as well as RPE genes *RPE65*, *TYR*, and *OTX2* were measured by RT-qPCR using explants collected at indicated conditions and time points. (D) EdU and pHH3 staining were performed in explants after 48h of cell culture. (E) Quantification of EdU or pHH3 positive cells from (D). (F) Representative western blot (WB) of RPE explants at 48h of cell culture at indicated conditions. (G) Protein bands quantification by densitometry of WB showed in (F). Plots show group mean ±SD; p-values from modeling as described in Statistical analysis section. *p<0.05, **p<0.01, *** p<0.001 and **** p<0.0001.

As we determined above, cell proliferation is a key cellular process for RPE reprogramming; therefore, we investigated whether cell proliferation was also affected by OXPHOS activation. We observed a comparable number of EdU and pHH3 positive cells in PBS- and DCA-treated explants at 24h and 48h (Fig. 4 D and E; Fig. S6 C and D). As was expected, FGF2 promoted cell proliferation, an effect that was inhibited by DCA (Fig. 4 D and E; Fig. S6 C and D). In addition, the expression of *PCNA* and *E2F1* was downregulated in explants treated with FGF2+DCA (Fig. S6 E). To ensure that DCA is exerting the intended effect, we performed western blot assays to detect the phosphorylated form of PDH (pPDH). WB analysis revealed a decrease of pPDH induced by DCA, at 24 or 48h of cell culture; however, the effect was more pronounced and became statistically significant at 48h (Fig. 4 F and G; Fig. S6 F and G). This result confirmed that DCA inhibited PDK, and in consequence, PDH was activated. Since FGF2 signals through MAPK kinases pathway, a pathway that is required for RPE reprogramming (Spence et al., 2007, 2004), we asked whether DCA might affect ERK phosphorylation. Surprisingly, DCA alone promoted ERK phosphorylation and become more prominent in the presence of FGF2 (Fig. 4 F and G; Fig. S6 F and G). To confirm that DCA affected RPE programming into neural retina at the protein level, a WB against α-tubulin, a protein that is highly present in neural tissues (Gloster et al., 1994; Lewis et al., 1985), was included. We observed that at 48h of cell culture, α-tubulin was present in FGF2-treated explants, but not in explants treated with PBS, DCA, or FGF2+DCA, confirming DCA inhibited neural fate induced by FGF2. Altogether, the data showed that OXPHOS activation, by promoting PDH activity, inhibits RPE reprogramming.

### OXPHOS activation redirects RPE reprogramming toward an EMT program

The phenotypic and molecular characteristics observed following OXPHOS activation suggested initiation of the cell reprogramming process but without resulting in retina formation as a final cellular fate. In order to elucidate this, we collected explants treated with DCA for RNAseq at 24h and 48h and compared them with the samples treated with FGF2 alone (Fig. S2). PCA showed a clear clustering of the samples by treatment (Fig. 5 A and Fig. S7 A-D), indicating a robust effect of OXPHOS activation on the gene expression. Although we observed a clear effect of DCA at both 24h and 48h, measured gene expression responses were generally much stronger at 48h; therefore, we decided to focus our study on potential fate changes in the samples collected at 48h. When gene expression patterns were analyzed, we noted that the expression patterns of some genes normally affected by FGF2, such as *FGFR1*, were not affected by DCA (Fig. 5 B).

**Figure 5.**
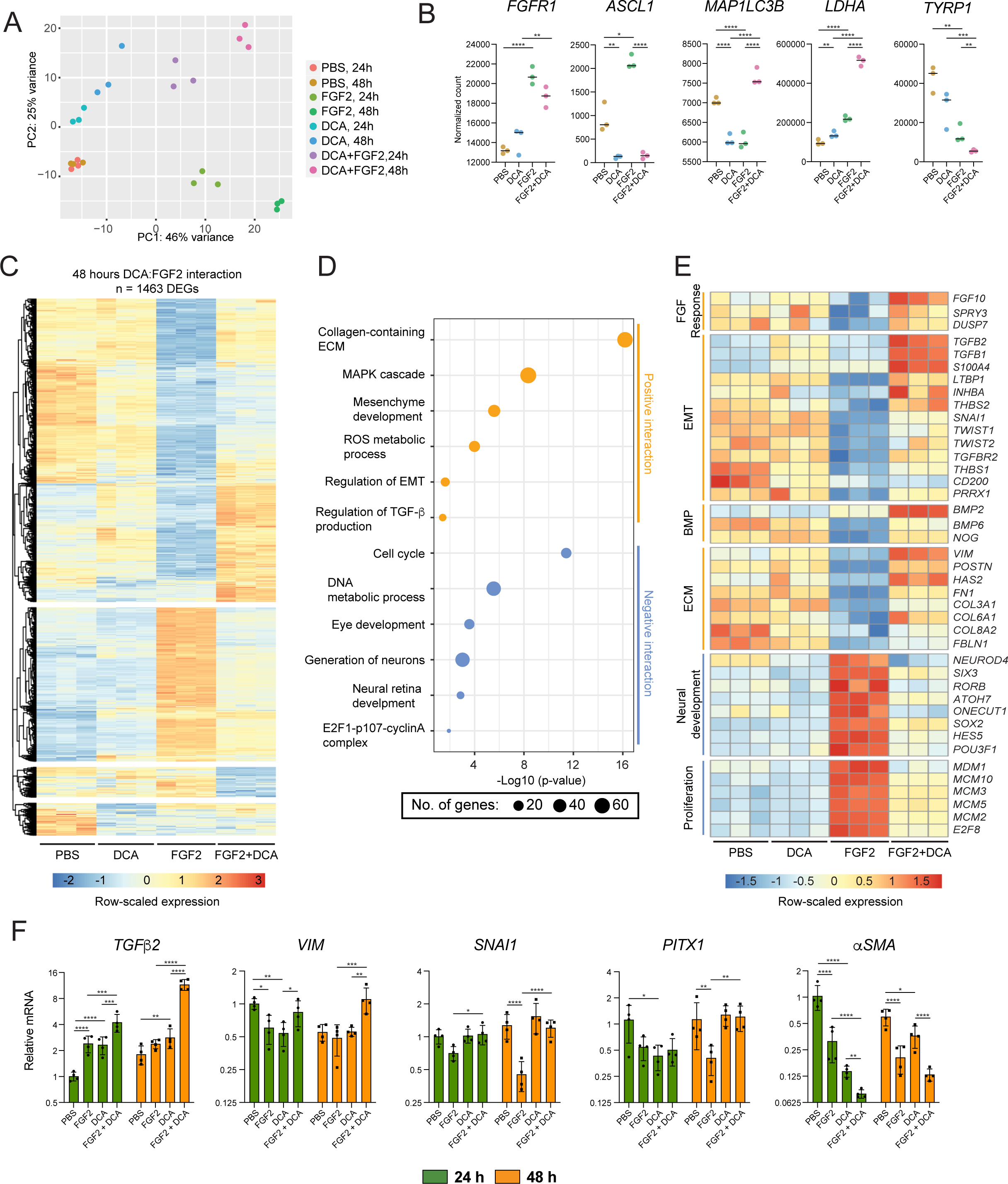
The activation of OXPHOS metabolism during RPE reprogramming turns on an EMT program. (A) Two-dimensional principal component analysis (PCA) displays RNAseq samples colored by condition. Samples show a clear spatial separation along PCA axes. (B) Representative RNAseq count plots display normalized expression values for genes of interest from explants samples after 48h of cell culture. (C) An interaction analysis was performed to identify genes that exhibit altered responses to FGF2 in the presence of DCA at 48 hours. Genes with adjusted p-value ≤ 0.05 and |log_2_(fold change)| ≥ 1 are displayed in a row-normalized heatmap. (D) Pathway enrichment analysis was performed on gene sets with positive or negative log_2_(fold change) values, as identified in C. (E) The expression patterns for genes of interest that display a significant DCA:FGF2 interaction are displayed in a row-normalized heatmap. (F) Levels of relative gene expression for EMT and mesenchyme associated genes were determined by RT-qPCR using explants collected at indicated conditions and time points. Plots show group mean ±SD; p-values from modeling as described in Statistical analysis section. *p<0.05, **p<0.01, *** p<0.001 and **** p<0.0001.

However, other classes of genes that were upregulated by FGF2, such as the neural retina factor *ASCL1*, were instead downregulated in the presence of DCA (Fig. 5 B). Genes with an opposite behavior were also found; this was the case for *MAP1LC3B*, a gene important for autophagy a process that can be regulated by FGF2/FGFR1 (Yuan et al., 2017), was downregulated by FGF2, but for which FGF+DCA treatment promoted expression (Fig. 5 B). In addition, for another set of genes, DCA potentiated the effect of FGF2; for example, *LDHA* was upregulated by FGF2, which was exacerbated by FGF2+DCA. Moreover, *TYRP1* was downregulated by FGF2, and further downregulated when both FGF2 and DCA were present (Fig. 5 B).

These observations indicated that the gene expression of many genes was altered by an interaction between FGF2 and DCA. Therefore, we performed an interaction analysis to answer the question “*How is the effect of FGF2 modified based on the addition of DCA?*” From this analysis, we identified 1463 genes with a significant interaction (Fig. 5 C). Pathway enrichment analysis of the identified genes set with positive log_2_(fold change) values revealed that the addition of DCA enhances the treatment effects of FGF2 for genes associated with extracellular matrix (ECM), MAPK cascade, mesenchyme development, ROS metabolism, the production of TGF-β, and EMT (Fig. 5 D). On the other hand, we observed a negative interaction for genes associated with cell cycle, DNA metabolic process, eye development, the generation of neurons as well as neural retina development (Fig. 6 D). Thus, these negatively regulated genes, and their associated processes, represent a class of biological effects elicited by FGF2 that are muted by the addition of DCA. Accordingly, these findings can be reconciled with the observed inhibition of cell proliferation observed with FGF2+DCA treatment (Fig. 4 D and E; Fig. S6 C, D and F), as well as the accumulation of pERK in the presence of DCA (Fig. 4 F and G; Fig. S6 F and G). Select genes were plotted in a heatmap, revealing the nature of the interactions for genes associated with FGF2 response, EMT, BMP signaling, as well as ECM. These gene sets were downregulated by FGF2 alone, but upon addition of both FGF2 and DCA, failed to be downregulated, or were even increased in expression (Fig. 5 E). In contrast, genes essential for neural development and proliferation were robustly activated by FGF2 alone, an effect that was largely abrogated by combinatorial treatment of FGF2 and DCA, demonstrating that DCA impairs the activation of essential neural reprogramming factors (Fig 5 E).

**Figure 6.**
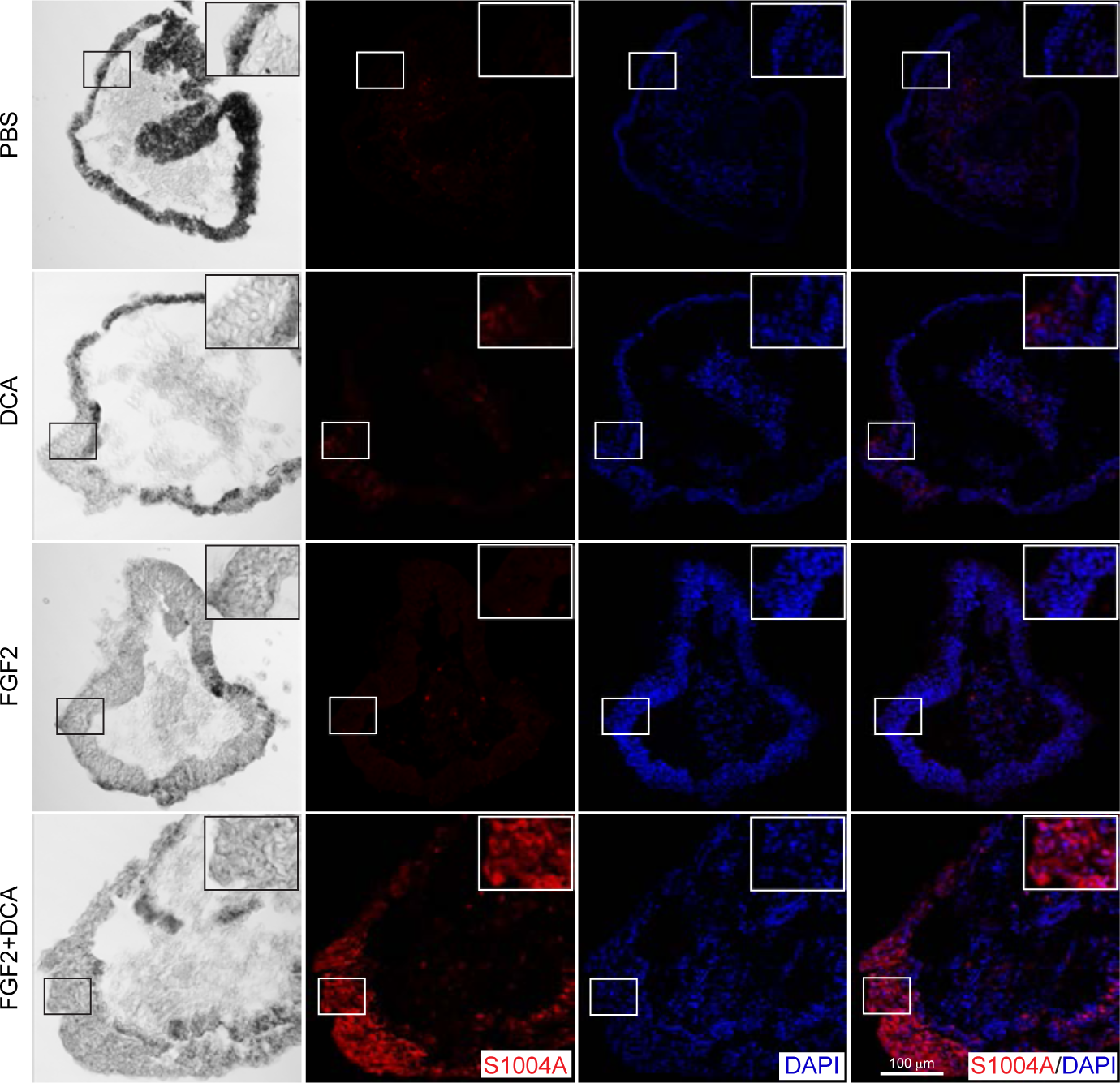
The activation of OXPHOS metabolism induces the presence of EMT protein S1004A in the RPE. Representative immunofluorescence images to detect EMT protein S1004A using explant cryosections collected after 48h of cell culture.

These results suggest that one of the main processes induced by the activation of OXPHOS metabolism during RPE reprogramming is an EMT program, and consequently, the redirection of RPE reprogramming toward mesenchymal fate. The determination of expression of select gene markers of EMT and mesenchyme fate revelated that *TGFβ2* and *vimentin* (*VIM*) were robustly upregulated in the FGF2+DCA condition by 48h, while *SNAI1* and *PITX1*, which were downregulated by FGF2 alone, were unchanged upon the addition of FGF2+DCA (Fig. 5 F). Notably, the addition of DCA did not modify the expression of *αSMA* relative to FGF2 alone; similar gene expression patterns were observed from RNAseq data (Fig. S7 E), collectively indicating that RPE was reprogramming into mesenchyme. However, during the RPE collection for the explants, a small amount of periocular mesenchyme tissue remains associated with the RPE, leaving the possibility that the observed gene expression alterations could be attributed to the mesenchyme cells associated with the explant. Thus, immunofluorescence against S100A4, a marker of EMT that also is present when RPE acquires mesenchyme characteristics (Chen et al., 2012; Lo et al., 2011), was performed on explants collected at 48h. S100A4 was not detected in the RPE or the reprogramed RPE but was detected at low levels in the RPE of DCA-treated explants (Fig. 6). Interestingly, S100A4 was robustly present throughout the corresponding RPE regions of explants treated with FGF2+DCA, which are discernable by the presence of pigment (Fig. 6). Altogether, the observations show that the activation of OXPHOS during FGF2 treatment turns on an EMT program that redirects RPE away from neural lineages and toward a mesenchymal fate.

### OXPHOS-induced EMT is partially driven by ROS

Activation of OXPHOS metabolism can promote reactive oxygen species formation (Mailloux, 2020; Ruggieri et al., 2014). In addition, it has been shown that ROS can induce EMT (Radisky et al., 2005; Jiang et al., 2017), therefore, the formation of ROS following the induction of OXPHOS might have an active role in the induction of EMT. To test this hypothesis, we supplemented the media with the antioxidant N-acetyl cysteine (NAC). NAC at 1 or 10 mM did not show any obvious phenotypic effects to the RPE explants (Fig. S8 A) and did not affect RPE reprogramming induced by FGF2 at 10 mM (Fig. 7 A). Surprisingly, NAC at 10 mM, but not at 1 mM, partially rescued the RPE reprogramming inhibited by DCA (Fig. 7 A and Fig. S8 B). The classification of the phenotype of the explants on day 4 of culture in class I (no reprogramming), class II (partially reprogrammed), and class III (reprogrammed), showed that DCA inhibited 100% of the reprogramming of the RPE, however, the addition of both DCA and NAC partially rescued reprogramming competence, with 30% of the explants showing signs of reprogramming, and 70% fully reprogrammed, although the total amount of neuroepithelium produced was lower compared to the explants treated with FGF2 alone or FGF2+NAC (Fig 8 A and B). Using RT-qPCR, we evaluated the effect of NAC at 10 mM concentration on RPE reprogramming using explants collected at 48h. We observed that the expression of neural retina markers *SOX2*, *SIX6*, and *PAX6* were not affected by either NAC or DCA in the absence of FGF2 (Fig. 7 C). In FGF2-treated explants, NAC also did not affect the gene expression of neural retina markers *SIX6* or *PAX6* (Fig. 7 C). However, FGF2+DCA induced the downregulation of *SOX2*, consistent with our RT-qPCR and RNAseq results (Fig. 5 E), but this downregulation was blocked by the addition of NAC (Fig. 7 C). On the other hand, media supplementation with NAC did not significantly affect the expression of the RPE markers previously observed to be regulated by DCA treatment (Fig. 4 C), in either FGF2-treated or no-treated explants (Fig. 7 C). Gene expression analysis revealed that NAC did not affect the expression of EMT markers in the absence of FGF2. Furthermore, within FGF2-treated explants, the induction of *TGFβ2* by OXPHOS was reduced by NAC, and a similar effect was observed for *VIM* gene expression levels, but no effect was detected for *SNAI*1 or *aSMA* (Fig. 7 D). Thus, limiting ROS production via the addition of NAC may partially inhibit the activation of EMT machinery that is caused by the DCA:FGF2 interaction. Regardless of the effect of NAC on EMT genes, no rescue effect was observed on cell cycle genes *E2F1* and *PCNA*, which were downregulated by DCA even when NAC was present (Fig. 7 D). These data suggest that ROS production is one component of OXPHOS metabolism during RPE reprogramming and plays an active role in the regulation of some neural genes and certain attributes of the EMT program. However, ROS production is not the only process involved and cannot explain the totality of the DCA:FGF2 interaction, such as effects related to proliferation and the full scope of the changes in the expression of EMT machinery.

**Figure 7.**
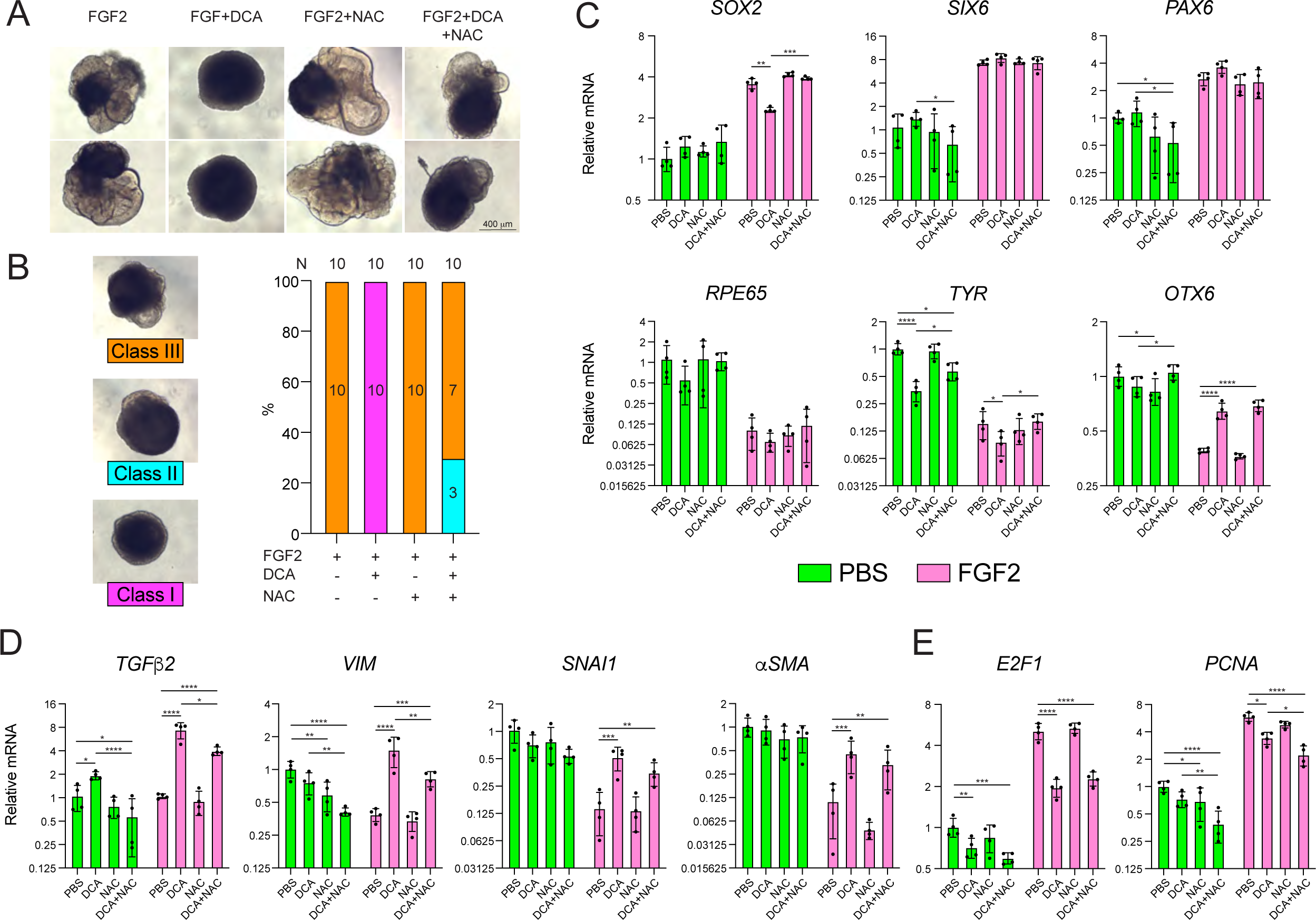
EMT activation by OXPHOS is partially reverted by the antioxidant NAC. (A) Representative RPE explants after 96h of culture treated with the indicated molecules. (B) Quantification of the occurrence of explant phenotypes shown in (A). Class I: no reprogrammed RPE; Class II: signs of RPE reprogramming; Class III: reprogrammed RPE. Gene expression analysis of retina and RPE markers genes (C), and EMT and mesenchyme genes (D) using RPE explants collected from the indicated conditions and time points. Plots show group mean ±SD; p-values from modeling as described in Statistical analysis section. *p<0.05, **p<0.01, *** p<0.001 and **** p<0.0001.

**Figure 8.**
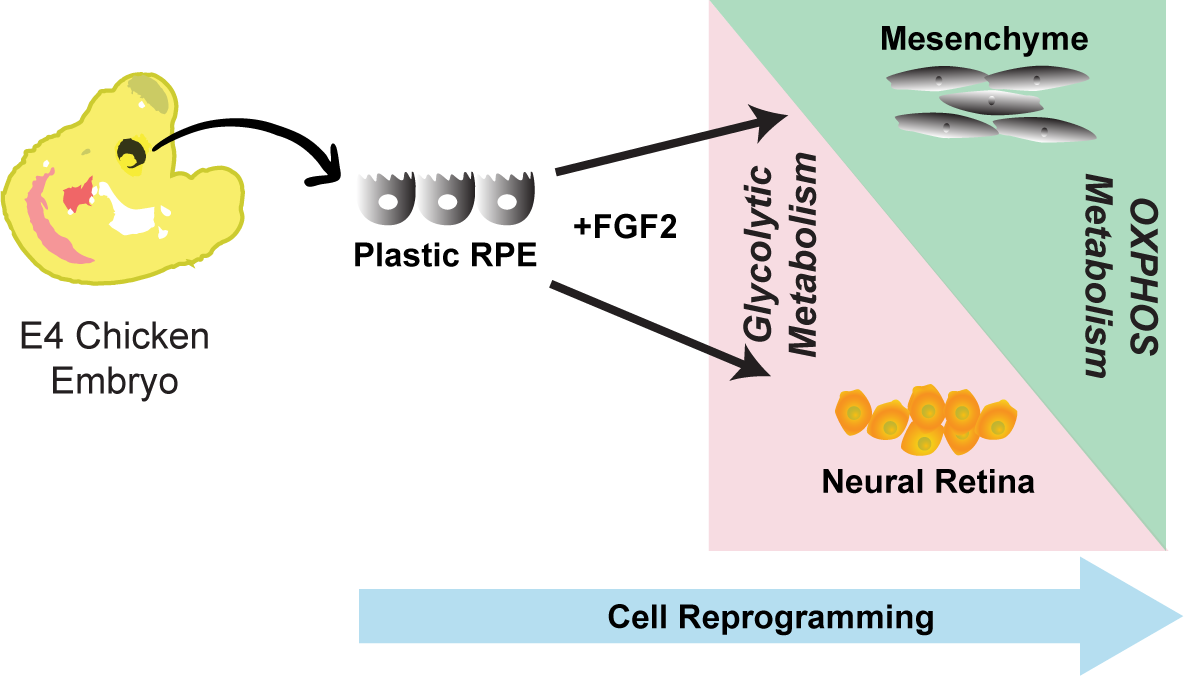
Proposed mechanisms for the role of glycolysis and OXPHOS in directing cell fate decisions during RPE reprogramming. Plastic RPE requires glycolytic metabolism to reprogram into neural retina while turning on OXPHOS metabolism induces an EMT program that favors mesenchyme formation.

## DISCUSSION

In this study, we comprehensively characterize RPE explants collected from chicken embryos as a model to study RPE reprogramming into neural retina ex vivo. Using this model, we elucidate the role of glycolysis and OXPHOS as RPE cells reprogram into neural retina. Our Gene expression analysis confirmed that RPE reprogramming ex vivo initiates with the presence of FGF2, the inducer of chicken RPE reprogramming into retina, by inductions of a loss of RPE identity, characterized by downregulation of RPE-associated genes and upregulation of neural and proliferation genes, similar to what is observed *in vivo* (Tangeman et al., 2022, 2021; Luz-Madrigal et al., 2014). This process is initiated by 6h *in vivo* (Tangeman et al., 2022, 2021; Luz-Madrigal et al., 2014) but *in vitro* is robustly observed by 24h after FGF2 addition (Fig. 1 C, D, and E). Interestingly, our results point out FGF2 as a necessary trigger of reprogramming but not required for reprogramed RPE growing, since FGF2 removal after an hour of exposition did not stop RPE reprogramming (Fig S1 A), a phenomenon that has not been possible to determine before *in vivo*.

The RPE explants model revealed remodeling of metabolic activity during RPE reprogramming. In this regard, we observed an upregulation of glycolytic genes following FGF2 addition (Fig. 3 A and B). Using different cell media conditions, we showed that RPE cells can use glucose, pyruvate, or glutamine as carbon sources to reprogram. Inhibition of glycolysis by 2DG, an inhibitor of the first two glycolytic enzymes, HK and GPI (Fig. 3 A), blocked RPE reprogramming, even in cases where pyruvate or glutamine served as the sole carbon source (Fig. 3 C). Pyruvate is the final product of glycolysis; therefore, its catabolism produces energy through the TCA cycle and does not require glycolysis (Fig. 3 A). Similarly, the use of glutamine for ATP production does not require glycolysis (Stumvoll et al., 1999) (Fig. 3 A). Hence, the hindrance of RPE reprogramming caused by 2DG in the presence of pyruvate or glutamine as the primary carbon sources suggests that, even in the absence of exogenous glucose, glycolysis is still active; sustaining glycolysis at these conditions necessitates the production of glucose. Glucose is synthesized by gluconeogenesis using metabolites such as pyruvate and glutamine, although gluconeogenesis is primarily regarded as limited to the liver, muscle, kidney, and intestine (Scrutton and Utter, 1968; Mithieux and Gautier-Stein, 2014). Our RNAseq analysis reveals expression of gluconeogenic genes in the RPE (Fig. 3 A and B), providing evidence that upon glucose depletion, glucose might be produced endogenously identifying RPE as a gluconeogenic tissue. However, to date, it has not been shown that RPE is able to perform gluconeogenesis, although RPE and retina have been described as metabolically active tissues (MacGregor et al., 1986; Hurley et al., 2015; Xu et al., 2020). RPE has the capability to produce pyruvate, malate, and citrate from both glucose and lactate. However, the production of citrate and malate was more extensive when lactate was utilized as a precursor (Kanow et al., 2017). Although RPE also metabolizes glucose, lactate can inhibit glucose metabolism through glycolysis, forming a metabolic flux that stimulates glucose metabolism in the retina (Kanow et al., 2017). The presence of various metabolites utilized by RPE, such as lactate, citrate, pyruvate, etcetera, which can potentially contribute to the gluconeogenesis pathway, provides compelling evidence supporting gluconeogenesis occurring in the RPE.

Interestingly, gluconeogenesis in the eye has been observed in the retina in both amphibians and mammals although it is limited to Müller glia, cells that also store glycogen (Kanow et al., 2017; Viegas and Neuhauss, 2021; Scrutton and Utter, 1968). However, it is unclear whether it is also present in photoreceptors (Hurley et al., 2015; Goldman, 1990; Mamczur et al., 2010). Based on this, in the context of RPE reprogramming, the RPE and reprogrammed RPE might not have similar gluconeogenic capabilities. In this regard, some insights can be obtained from our RNAseq data, which shows similar levels of gluconeogenesis-related genes, including *PC*, *PCK1*, and *FBP1*, in PBS- and FGF2-treated samples at 24h (Fig. 3 B). However, in PBS-treated samples, expression levels of these genes were higher at 48h than at 24h and no alterations were noticed in the reprogrammed RPE explants, although higher expression of *G6PC3*, the catalytic subunit of the gluconeogenic enzyme G6P, was observed in reprogramed RPE at 48h (Fig. 3 B). Nevertheless, *FBP1* was more highly expressed at 48h in PBS-treated explants compared to FGF2-treated explants (Fig. 3 A, Fig. 4S A). Altogether, these observations indicate that gluconeogenesis might be a specific metabolic characteristic of the RPE rather than the retina; hence, it might be part of RPE identity or maturation. Therefore, as RPE transitions into neural retina is likely to lose its ability to synthesize glucose, becoming a glycolytic tissue, as RNAseq indicates (Fig. 3 B). Thus, we propose that RPE cell reprogramming includes metabolic reprogramming to proceed.

The inhibition of RPE reprograming by disturbing glycolysis even in the presence of pyruvate or glutamine, not only emphasizes the significance of glucose as an energy source but also reveals that glucose serves functions beyond energy production. Interestingly, shifting glycolytic metabolism toward OXPHOS via the activation of PDH, a metabolic state where ATP production will be highly favored, blocks RPE reprogramming into neural retina, reinforcing the importance of glucose source of metabolites for appropriate cell reprogramming of the RPE. Unexpectedly, interaction analysis of RNAseq data from FGF2 and DCA-treated explants, revealed that OXPHOS enhances the effect of FGF2 on gene expression related to mesenchyme development, ROS metabolism, and EMT, but negatively impacts the effect of FGF2 on genes related to cell cycle, the generation of neurons, and neural retina development (Fig. 5 D), which are prerequisites for RPE reprogramming (Tangeman et al., 2022). On the other hand, genes normally downregulated by FGF2, such as those involved in ECM, BMP signaling, FGF response, and EMT, failed to be downregulated, or in some cases were even upregulated, when OXPHOS was activated (Fig. 5 E). Amongst the upregulated genes are *INHBA (Activin)*, *BMP2/6*, and *TGFβ1/2*, which belong to the TGFβ superfamily of structurally related cytokines. BMP factors are critical effectors of RPE and neural retina fate (Steinfeld et al., 2017; Müller et al., 2007), and similarly, activin signaling has been shown to restrict RPE neuro-competence (Sakami et al., 2008; Fuhrmann et al., 2000). Therefore, OXPHOS activation triggers the repression of the neural fate program and simultaneously activates signals that restrict RPE neuro-competence.

Several members of the TGFβ superfamily, such as TGFβ1 and Activin are positive regulators of EMT (Kahata et al., 2018), suggesting that OXPHOS activation during RPE reprogramming might elicit an EMT program that redirects RPE cells toward a mesenchymal fate. This observation is supported by gene expression patterns of EMT effectors and mesenchymal markers, as well as the observed presence of S1004A in the RPE, a calcium-binding protein present in EMT progression across several cellular contexts, including RPE (Lo et al., 2011; Chen et al., 2012). Recently, Zu et al., 2023, showed acetate controls endothelial-to-mesenchymal transition (EndMT), and identified inhibition of PDK4 as one of the main components (Zhu et al., 2023). At a molecular level, EndMT requires an increase in TFGβ signaling (Chen et al., 2019). Interestingly, Zu et al, found that TFGβ suppresses the expression of PDK4, promoting the synthesis of acetate, which is used by ACSS2 to produce Ac-CoA. Ac-CoA overproduction causes acetylation of TGFβ receptor ALK5 as well as the mediators SMAD2 and SMAD4, allowing long-term TGF-β signaling and EndMT. Surprisingly, we found that PDK4 inhibition by DCA results in overexpression of *TGFβ1* and *TGFβ2* (Fig. 5 E and F), thus, the overproduction of acetate by an increase of PDH activity could potentially activate TGFβ signaling and be responsible for EMT program in the RPE in an FGF2-dependent manner.

In the human cell line ARPE-19, DCA prevented EMT induced by TGFβ (Shukal et al., 2020). The authors found that TGFβ induced the expression of EMT markers, wound healing response, and phosphorylation of ERK, effects that were reduced by DCA (Shukal et al., 2020). The clearly contradictory observations between Shukal et al. and our study may be due to differences in RPE origin. ARPE-19 is a spontaneously arising human RPE cell line that retains some phenotypic and expression characteristics of native RPE cells, but also retains substantially different molecular signatures than primary human or iPSC-derived RPE (Samuel et al., 2016; Markert et al., 2022; Dunn et al., 1996). On the other hand, the RPE explants used in our study were collected from chicken embryos and studied in the context of cell reprogramming. A recent study showed that RPE derived from iPSC cells has a transcriptomic signature that suggests it is in a more immature state than primary-derived RPE, a state that makes it more susceptible to EMT transformation induced by TGFβ or TNFα (Markert et al., 2022). Thus, differences in RPE origins could explain the contradictory findings, but also reveal that the response to the same metabolic disturbance can produce an opposite outcome depending on the cellular and environmental context.

In addition to energy production, OXPHOS activation can also cause ROS formation by TCA and electron transport chain (ETC) (Hernansanz-Agustín and Enríquez, 2021; Hamanaka and Chandel, 2010), a phenom also observed upon OXPHOS activation by DCA (Ruggieri et al., 2014). It has been described that mitochondrial ROS can affect and regulate cellular processes such as proliferation, differentiation, and apoptosis through the regulation of signaling pathways, including PI3K, MAPK, or activating transcription factors, such as HIF1a and FOXO (Hamanaka and Chandel, 2010; Schieber and Chandel, 2014). Furthermore, ROS can also induce EMT, a process principally studied in cancer cells (Jiang et al., 2017; Radisky et al., 2005). All this evidence suggests ROS participation in promoting EMT. Interestingly, we observed that the ROS scavenger NAC, partially inhibited EMT progression of the RPE and overexpression of *TGFβ2* when OXPHOS was activated and concomitantly rescued retinal formation (Fig. 7 A and D). Therefore, ROS hereby contributes to the EMT program observed during RPE reprogramming, which could represent a linear sequence of events supported by our data.

Eye disorders such as diabetic retinopathy, proliferative vitreous retinopathy, and AMD, are multifactorial diseases characterized by RPE dysfunction and degeneration (Yang et al., 2021). The accumulation of connective tissue, a process known as fibrosis, is commonly observed in the context of vitreoretinal diseases, and is caused by EMT of RPE cells (Hiscott et al., 1999). Several studies have shown that ROS, TGFβ, and TNFα are some of the drivers of EMT in RPE cells (Boles et al., 2020; Yang et al., 2020). Furthermore, glucose imbalance is strongly correlated with the development of diabetic retinopathy and EMT of RPE cells (Cheung and Wong, 2007; Che et al., 2016), indicating that metabolic state might play an active role in EMT and vitreoretinal diseases. In this study, we show evidence that glycolytic metabolism is essential for appropriate RPE reprogramming into neural retina while metabolic reprogramming of RPE toward OXPHOS induces EMT redirecting the final cellular fate into mesenchyme (Fig. 8). Therefore, our data provide new insights into the importance of metabolism in RPE cell decisions and could point toward a potential role for metabolism dysfunction during vitreoretinal disease progression. It remains to be investigated whether the metabolic state of mature RPE cells may be inhibitory to neurogenesis or bias them toward fibrotic lineages following injury. Similarly, important insights could be garnered by comparing the metabolic state of regeneration-competent RPE, such as is found in the embryonic chicken, to the metabolism observed in the RPE of adult mammals. As suggested in our study, it is possible that the attenuation of ROS and a switch toward glycolytic metabolism could represent two essential steps toward unlocking latent neurogenic competence from RPE cells, or at least for preventing fibrotic outcomes. Future studies on this frontier could be oriented toward leveraging metabolism to expand the capacity of the RPE to reprogram into neural retina across these contexts. As shown in our study, intrinsic metabolic alterations and the composition of available metabolites can drastically affect the outcome of RPE reprogramming, and consequently, the formation of neural retina. Thus, future paradigms seeking to use RPE therapeutically, whether for RPE transplantation or neural retina regeneration, are likely to benefit from a more comprehensive understanding of the interplay between metabolism and RPE cell fate maintenance.

## CONCLUSION

In this study, we provide new insights into the metabolic characteristics of the RPE, as well as the role of metabolism in determining RPE reprogramming potential. Our data indicate that RPE is an actively glycolytic tissue that also has the capacity to synthesize glucose. We identify glycolysis as necessary for appropriate RPE reprogramming into neural retina, and revealed that the activation of OXPHOS metabolism during cell reprogramming elicits an EMT program, causing RPE cells to acquire mesenchyme features. Importantly, this EMT program can be partially attenuated by antioxidant treatment. These findings provide new knowledge to better understand common RPE diseases such as diabetic retinopathy or proliferative vitreoretinopathy, which, during their pathogenesis, show EMT characteristics and are commonly present in people who suffer previous metabolic alteration caused by disorders, such as diabetes or acute injury.

## MATERIALS AND METHODS

### *Chicken* embryos, RPE explants culture, and small molecule treatments

Fertilized white leghorn chicken eggs were obtained from Michigan State University. Eggs were incubated in a humidified rotating incubator at 37°C. At day 4 of development (E4, Hamburger & Hamilton Stage 24) RPE sheets were collected and cultured as previously described, with slight modifications (Sakami et al., 2008). Briefly, RPE tissue was dissected from chicken embryos at E4 (Stage 24, (Hamburger and Hamilton, 1951)), with a small amount of associated mesenchyme, in modified HBSS buffer (without calcium chloride, magnesium sulfate, and sodium bicarbonate, supplemented with 5mM HEPES and 0.6% D-glucose). RPE sheets were washed in HBSS solution and cultured in 500 µl of explant medium in 24-well plates. The plates were incubated in an orbital shaker at 50 rpm (3-D, Fixed Tilt Platform Rotator, Grant Instruments) at 37 °C with 5% CO2. An explant medium was prepared as described previously (Sakami et al., 2008). DMEM/F12 (HyClone, #SH30026), was supplemented with 0.9% D-glucose (Sigma, #G-7021), 0.1125% NaHCO3 (Sigma-Aldrich, #S5761), 20 mM HEPES (Boston BioProducts Inc, #BB-2076-K), 5% FBS (Fisher brand, #FB12999102), and antibiotics (Gibco, #15-240-062). In addition, DMEM, with no glucose, no phenol red, and no glutamine (Thermo Fisher, #A14430) or DMEM with no glucose (Thermo Fisher, #11966025) were supplemented with 20 mM HEPES, 5% FBS and antibiotics, and used in the indicated experiments. FGF2 (R&D Systems, #3718-FB-025) was added at a concentration of 100 ng/ml as specified. Sodium dichloroacetate (Sigma-Aldrich, #347795-10G) 1M, 2-Deoxy-D-glucose (2DG, Sigma-Aldrich, #D-8375-1G) 1M, and N-acetyl cysteine (Sigma-Aldrich, #A9165-5G) 0.5M stock solutions were prepared in water and added directly to the cell media, as specified. NAC stock was adjusted at pH 7 with NaOH. A stock of D-glucose (Sigma-Aldrich, #G-7021) was prepared in water at a final concentration of 2 M and used for indicated experiments. Pyruvate (HyClone, #SH30239.01) and GlutaMAX (Gibco, #35050079), a substitute for L-Glutamine, were used in the indicated experiments.

### Immunoblotting

Three RPE explants were used per biological sample. RPE explants were collected in 60 μl of lysis buffer (Tris-HCl 62.5 mM, SDS 2%, pH 6.8) with protease (Roche, #1183617001) and phosphatase inhibitors (ThermoFisher, #A32957). The samples were sonicated, and protein quantification was performed using the Bradford assay (ThermoFisher, #1863028). 10 μg of protein was separated by electrophoresis on 4-20% Mini-PROTEAN TGX Precast Protein (Biorad, #4561096) and transferred to PVDF membranes (Biorad, #1704274). The membranes were blocked with EveryBlot Blocking Buffer (Biorad, #12010020) for 10 minutes and incubated overnight at room temperature with primary antibodies diluted in EveryBlot Blocking Buffer at the indicated concentrations (Table S1). After overnight incubation, the membranes were washed three times, 5 minutes each, with TBST buffer and incubated with secondary antibodies diluted in TBST at the indicated concentrations (Table S1). When the secondary antibody was HRP-linked, SuperSignal chemiluminescent substrate was used (ThermoFisher, #43580) following the manufactures’ instructions. The membranes were scanned in a ChemiDocMP scan (Biorad Laboratories, Inc). Images obtained from ChemiDocMP were analyzed and quantified using Image Lab version 6.1.0 (Biorad Laboratories, Inc). Actin, selected as a housekeeping protein, was detected using a monoclonal primary antibody and a fluorescent conjugated secondary antibody (Table S1). It was detected together with other target proteins only when the other primary antibody corresponded to a different host species (Table S1). Because the primary antibodies to detect the total and phosphorylated isoforms of ERK and PDH were developed in the same host species, it was no possible to precisely detect both isoform proteins from the same blot. Therefore, two WB were run at the same time using the same loading mix of each sample. One of the blots were used to detect the total protein and the other the phosphorylated isoform. Following this procedure, we could accurately determine the phosphorylated status of ERK and PDH.

### Histology, Immunostaining, and EdU detection

Explants used for EdU detection were exposed to EdU (Invitrogen, #A10044) for an hour prior to collection by adding 5 μl of EdU 2 mM to 500 μl of cell media. Explants just used for immunodetection were not exposed to EdU. Then, RPE explants were washed thrice with PBS for 5 minutes and fixed with 4% PFA for 15 min. After fixation, explants were washed three times with PBS and embedded in O.C.T. Compound (Fisher, #4585) and cryosections of 10 µm were obtained using a cryotome (Cryostar NX50, ThermoFisher) and collected on super frost plus microscope slides (Fisher Scientific, #22-037-246). For hematoxylin and eosin staining (H&E) the slides were rinsed with PBS and histology staining following a previously described (Luz-Madrigal et al., 2014). For immune fluoresces and EdU detections, the slides with tissue sections were rinsed three times with PBS and incubated for 5 minutes in 5% saponin (Sigma-Aldrich, #S7900) solutions following three washes of PBS. For EdU, click-it solutions were prepared according to the kit protocol (Invitrogen, #C10337), and each slide was covered with 150 μl of click-it solution for 30 min. Then, the click-it solution was removed, and the explants were washed three times with PBS. Afterward, slides were blocked with 200 μl donkey serum (Sigma-Aldrich, #D9663) prepared at 10% in PBS for an hour. Then, the blocking solution was removed and 150 µl primary antibody (Table S1) was prepared in blocking solutions and added to incubate overnight at 4°C. The primary antibody was removed, and three washes of PBS were performed, followed by 3 washes of PBST. Then, the slides were incubated in 100 µl of secondary antibody prepared in a blocking solution for 2h. Afterward, three washes with PBS, for 5 minutes each, were performed, and the slides were incubated for 5 minutes in 100 µl of DAPI solution 1 μg/μl (Sigma-Aldrich, #10236276001) before finishing with three more washes of PBS. If the samples were just processed for immunofluorescence, EdU detection steps were skipped. The slides were mounted with Fluormount (Sigma-Aldrich, #F4680) and cover slipped. For imaging, confocal images were obtained using 710 Laser Scanning Confocal System (Jena, Germany) using a × 20/0.80 Numeric Aperture (NA) = 0.55 WD objective lens or EC Plan-Neofluar.

### RNA isolation and RT-qPCR gene expression

RPE explants were collected in 200 μl DNA/RNA Shield buffer (Zymo Research, #1220-25) and stored at −20 °C. Each biological sample was a pool consisting of three explants. Total RNA was isolated using the Quick-RNA Microprep Plus Kit Microprep (Zymo Research, #R1051) following the manufacturer’s instructions. Total RNA samples were analyzed for quantity and quality using Nanodrop ND-2000 Spectrophotometer (Thermo Scientific) and Agilent 2100 Bioanalyzer (Agilent Technologies), respectively. 200 ng of RNA was used as a template to synthesize cDNA using QuantiTect Reverse Transcription kit (Qiagene, #205313) according to manufacturer’s instructions. The synthesized cDNA was diluted at 1:10 ratio with pure water. 2 μl of the cDNA dilution were used for quantitative PCR (qPCR) reaction. The final qPCR reaction contained 2μl of diluted cDNA, 10 μl of TB Green® Advantage® qPCR Premix (Takara, #639676), and 50 nM of each primer adjusted to 20 µl with water. qPCR reactions were set up in duplicate in the Rotor-Gene Q thermocycler 5 plex (Qiagen, Germantown, MD, USA). Primers reported here were designed using Primer BLAST (https://www.ncbi.nlm.nih.gov/tools/primer-blast/, Table S2) and obtained from IDT Technologies. The comparative ΔΔCt method was used to determine the relative gene expression levels compared with the housekeeping gene (*RPLP0*). Four biological samples were used under each condition.

### RNA-seq Library Preparation and Sequencing

Purified RNA was collected in triplicate, as described above, and quantified using the Qubit 4 and Qubit RNA HS Kit (ThermoFisher, #Q32852). RNA integrity was validated using the Agilent 6000 Pico Kit (Agilent, #5067-1513). For each sample, 100 ng of total intact RNA was used with the NEBNext® Poly(A) mRNA Magnetic Isolation Module (NEB, #E7490S), and the enriched mRNA was prepped for sequencing with the NEBNext® Ultra™ II Directional RNA Library Prep with Sample Purification Beads (NEB, #E7765S) according to the manufacturer’s instructions. Thirteen PCR amplification cycles were used, and indexing was performed using single index oligos (NEB, 7710/7730). The final amplified libraries were validated using the Agilent High Sensitivity DNA Kit (Agilent, 5067-4626) and quantified using the Qubit dsDNA HS Kit (Thermo Fisher, Q32851). Samples were pooled in equimolar ratios before sequencing across two lanes of Illumina HiSeq X Ten at the Novogene sequencing core using 150 base pair paired end reads.

### RNA-seq data analysis

Raw reads were analyzed using FastQC v0.11.9 (Babraham Bioinformatics - FastQC A Quality Control tool for High Throughput Sequence Data) and MultiQC v1.9 (Ewels et al., 2016) for quality assessments. Adapters and low-quality bases were removed with Trim Galore v 0.6.4_dev (Krueger, 2016) with the parameters *–stringency 3 –paired –length 36*. Trimmed reads were aligned to the chicken genome GRCg6a using the STAR alignment tool v2.7.5b (Dobin et al., 2013) and parameters *–quantMode GeneCounts TranscriptomeSAM –runThreadN 30 –limitBAMsortRAM 60000000000 – readFilesCommand zcat –genomeLoad LoadAndKeep –outSAMstrandField intronMotif – outSAMtype BAM SortedByCoordinate*. The genome index for star was generated with the parameters *–runMode genomeGenerate –sjdbOverhang 149 – genomeSAindexNbases 13*, and splice sites were incorporated from Ensembl release 106 (Cunningham et al., 2022). Gene counts were generated with Stringtie v2.1.4 (Pertea et al., 2015) using the parameters *–rf -e*, and genes with less than 10 raw counts were filtered from the analysis. Differential expression testing was performed using DESeq2 v1.34.0, and genes with p-adjusted value ≤ 0.05 and |log_2_(fold change)| ≥ 1 were considered differentially expressed throughout the manuscript (Love et al., 2014). For the interaction analysis, the formula design used employed a 7-term model, *∼ FGF2 + DCA + time + FGF2:time + DCA:time + FGF2:DCA:time + DCA:FGF2*. To test whether the effect of FGF2 was different across DCA levels at 48 hours, the DESeq2 *results ()* function was used to add the parameter estimates *FGF2Yes.DCAYes* and *FGF2Yes.DCAYes.time48h*. Pathway enrichment analysis was performed with g:Profiler with chicken genes converted to human orthologs (Kolberg et al., 2020; Raudvere et al., 2019). Multiple hypothesis testing for enriched pathways was performed using the g: SCS algorithm within g: Profiler.

### Statistical analysis

Experiments measuring relative gene expression by RT-qPCR or RNAse (see Fig. 1 C, 3 E, 3 F, 3 G, 4 C, 5 B, 5 F, 7 C, 7 D, 7 E, S3 A, S 4, S5 B, S5 F and S6E) were log(2)-transformed and then analyzed using ANOVA. Specific comparisons were made, and these p-values were adjusted to control for multiple comparisons by the false discovery rate (FDR) procedure (Benjamini and Hochberg, 1995), with the exception of analyses comparing treatments to control (Fig. 1 C and Fig. S4) in which case we used Dunnett’s multiple comparisons procedure. EdU and pHH3 quantification (count responses; see Fig. 2 E, 4E, and S6D) were analyzed using negative binomial regression (Lindén and Mäntyniemi, 2011; White and Bennetts, 1996; Hoef and Boveng, 2007), and multiple comparisons were also corrected using FDR. Immunoblotting experiments, which involved ratios related to PDH/pPDH, pERK/ERK, and Tubulin/Actin (see Fig. 4 G and S6 G), were also log(2)-transformed to stabilize unit-to-unit variance of these ratios, before analyzing the responses via ANOVA and FDR-adjusted p-values. One of the key assumptions for the use of the linear model is that each observation has the same variance. In this paper, there are a large number of statistical tests performed based on ANOVA models, and unsurprisingly there are some instances of what seems to be visual evidence of nonconstant variance across experimental groups. We acknowledge the fact that because of this, some of the resulting p-values may only be approximately valid. However, we typically do not see systematic differences in variation, as a function of the fitted values; furthermore, our conclusions are not based upon any particular p-value, but on the weight of the results as a whole. Thus, even if some of the experimental data does not perfectly meet the model assumptions, our analysis given here still provides evidence in support of our basic contentions. For the sake of transparency, we have included the data along with a supplementary document detailing our statistical analyses (SFile_Statistics). All statistics were performed in the R environment. For data plotting R environment or Prism 10.0.1 for MAC, were used.

## Supporting information

Stables

Statistical analysis

## AUTHOR CONTRIBUTIONS

JRPE and KDRT were responsible for the conception, design, and writing of this study. JRPE performed explant collection and culture, total RNA isolation, RT-qPCR, WB, immunostaining, and data analysis. JAT prepared the RNAseq libraries and analyzed RNAseq data. MPN and HS assisted with explants collection and immunostaining. BS assisted with statistical analysis. All authors reviewed and approved the final manuscript.

## FUNDING

This work was supported by National Eye Institute grant R01 EY026816, the Miami University Rapid Grant, and the John W. Steube endowed Professorship to KDRT. Further support was provided by National Institute of Neurological Disorders and Stroke grant F99 NS129167 to JAT. Support was also derived from the Undergraduate Research Award Miami University program to MPN and HS, as well as the Miami University Honors Program and Undergraduate Summer Scholars Miami University program to MPN and HS, respectively.

## ACKNOWLEDGMENTS

We acknowledge and thank the staff (Dr. Andor Kiss & Ms. Xiaoyun Deng) of the Center for Bioinformatics & Functional Genomics (CBFG) at Miami University for instrumentation and computational support. The authors further thank Zachery Oestreicher of the Center for Advanced Microscopy and Imaging (CAMI) at Miami University for providing instrumentation support. We also thank Erika Grajales-Esquivel for her managerial assistance and critical reading of the manuscript.

---

**Figure S1. Transient FGF2 treatment is sufficient to trigger RPE reprogramming, and the E4 RPE loses reprogramming competence between 24h and 48h in culture.** (A) Timeline of FGF2 removal at 1h or 6h of cell culture and representative images of explants. (B) Timeline of FGF2 additions at indicated time points of cell culture and representative images of explants. (C) Timeline RPE explants collection at different time points.

**Figure S2. RNAseq quality metrics.** (**A)** The table summarizes the quality metrics for each RNA-seq sample used throughout the study, including raw read duplication rates, nucleotide composition, and sequencing depth. Quality/adapter trimming statistics and alignment rates are also provided**. (B**) Alignment rates of reads to the chicken genome using STAR.

**Figure S3. Gene associated with neural fate are upregulated during RPE reprogramming. (A)** RNA-seq count plots display normalized expression values for neural retina and RPE genes in the presence or absence of FGF2. Y-axis is log scaled. (*) denotes adjusted p-value ≤ 0.05, n.s. = not significant**. (B)** The top 60 up-regulated genes associated with the term Generation of neurons (GO:0022008) are displayed in a row-normalized heatmap.

**Figure S4. Metabolic genes are upregulated by FGF2.** Expression patterns for glycolytic genes *HK1*, *HK2*, *GADPH*, *LDHA*, and *LDHB*, as well as the glucogenic gene *FBP1*, were determined by RT-qPCR in explants collected at different time points in presence or absence of FGF2. Plots show group mean ±SD; p-values from modeling as described in Statistical analysis section. *p<0.05, **p<0.01, *** p<0.001 and **** p<0.0001.

**Figure S5. Glucose, pyruvate, and glutamine affect RPE reprogramming.** (A) Representative RPE explants cultured in explant medium (see materials and methods) with or without FGF2 or 2DG at indicated concentrations were photographed after 96h of culture. (B) Gene expression determination of retina and RPE markers using explants cultured in explant medium with or without FGF2 or 2DG at 20 mM and collected after 24h or 48h of culture. (C) Representative RPE explants cultured in DMEM without glucose (see materials and methods) and in the presence or absence of FGF2 and glucose were photographed after 96h of culture. (D) Representative RPE explants at 96h of culture in DMEM without glucose, glutamine, and pyruvate and supplemented with glucose, pyruvate, or glutamine individually at 4 mM with FGF2, and 2DG 2 mM addition. (D) quantification of explants according to the phenotypic classification shown in Figure 3D. (F) Gene expression analysis of glycolysis-associated genes using RPE explants cultured in DMEM without glucose, glutamine, or pyruvate and supplemented +/-glucose at 2 mM and/or +/-2DG at 2 mM and collected at indicated time points. Plots show group mean ±SD; p-values from modeling as described in Statistical analysis section. *p<0.05, **p<0.01, *** p<0.001 and **** p<0.0001.

**Figure S6. OXPHOS activation inhibits RPE reprogramming independently of the available carbon source.** (A) Representative images of RPE explants cultured in DMEM, no glucose-supplemented medium (see materials and methods) with or without FGF2 addition and DCA at 20 mM after 96h of culture. H&E staining of RPE explants cultured for 96h in explant medium and treated with +/-FGF2 and/or +/-DCA 20 mM. (C) EdU and pHH3 staining were performed in explants after 24h of cell culture. (D) quantification of EdU or pHH3 positive cells from (C). Plots show mean ±SD. (E) Levels of relative gene expression of cell cycle genes *PCNA* and *E2F1* measured by RT-qPCR from explants collected at indicated conditions and time points. (F) Representative WB of RPE explants at 24h of cell culture and indicated conditions. (G) Protein band quantification by densitometry of WB shown in (F). Plots show group mean ±SD; p-values from modeling as described in Statistical analysis section. *p<0.05, **p<0.01, and *** p<0.001.

**Figure S7 Principal component analysis (PCA) of RNAseq data.** (A) 3-dimensional PCA displays RNA-seq samples colored by condition. Samples show a clear spatial separation along PCA axis 1 when colored by FGF2 treatment (B) and a clear separation along PCA axis 2 when colored by DCA treatment (C). PCA axis 3 captures variation attributable to time in culture (D). (E) Normalized counts for genes encoding EMT markers are displayed.

**Figure S8. NAC does not affect RPE explant morphology in the absence of FGF2.** (A) Representative RPE explants cultured in an explant medium with or without NAC at 1 mM or 10 mM and photographed after 96h of culture. (B) Representative RPE explant following culture in explant medium containing FGF2 and DCA at 20 mM and +/-NAC 1 nM and photographed after 96h of culture.

**SFile_STables.docx.** File contains Table 1S and 2S. **SFile_Statistics.pdf.** File contains full statistical analysis. **SFile_RawData.xlsx.** File contains raw data.

**SFile_DE_Testing.xlsx**. File contains differential expression statistics for RNA-seq based comparisons used in the study, calculated with DESeq2.

**SFile_Gene_Ontology.xlsx**. File contains gene ontology results for FGF-responsive genes following 24 and 28 hours of culture, as well as interaction analysis performed at 48 hours.

**Data availability**. Upon request

## Notes

### Competing Interest Statement

The authors have declared no competing interest.

