## Supplementary material for "DISTINCT METABOLIC STATES DIRECT RETINAL PIGMENT EPITHELIUM CELL FATE DECISIONS": Figures

### Figure S1

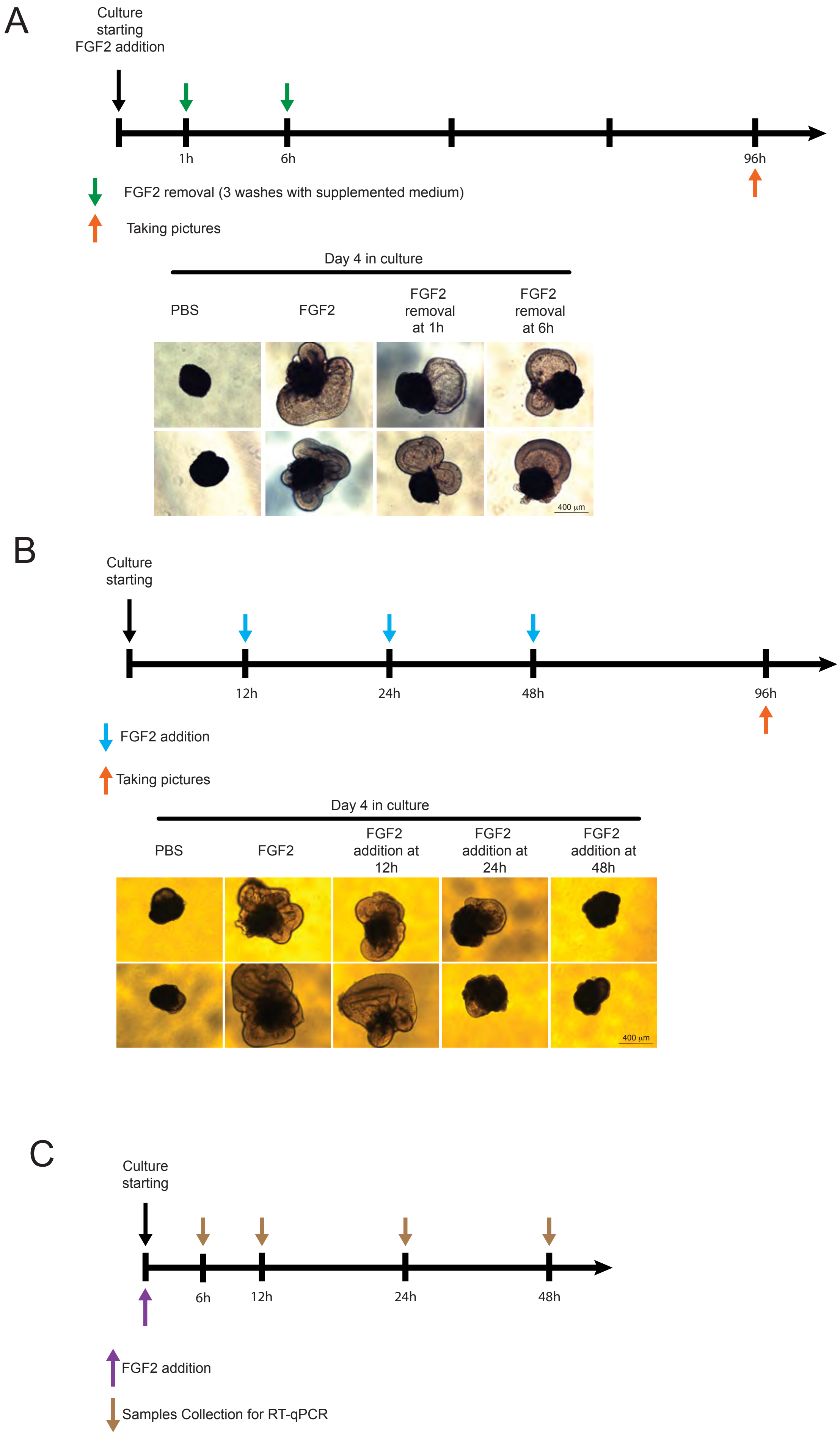

A
Culture starting FGF2 addition
1h
6h
96h
FGF2 removal (3 washes with supplemented medium)
Taking pictures
Day 4 in culture
PBS
FGF2
FGF2 removal at 1h
FGF2 removal at 6h
400 µm
B
Culture starting
12h
24h
48h
96h
FGF2 addition
Taking pictures
Day 4 in culture
PBS
FGF2
FGF2 addition at 12h
FGF2 addition at 24h
FGF2 addition at 48h
400 µm
C
Culture starting
6h
12h
24h
48h
FGF2 addition
Samples Collection for RT-qPCR

### Figure S2

A

| Sample | %Aligned | M Aligned | % BP Trimmed | % Dups | % GC | % M Seqs |
|---|---|---|---|---|---|---|
| DCA+FGF2 24h, S1 | 93.6% | 13.6 | 3.9% | 50.3% | 49% | 14.6 |
| DCA+FGF2 24h, S2 | 93.5% | 15.6 | 3.9% | 50.7% | 49% | 16.8 |
| DCA+FGF2 24h, S3 | 93.3% | 15.6 | 3.8% | 51.5% | 49% | 16.8 |
| PBS 48h, S1 | 93.9% | 16.3 | 3.2% | 49.7% | 48% | 17.4 |
| PBS 48h, S2 | 93.8% | 20.0 | 3.2% | 53.0% | 48% | 21.4 |
| PBS 48h, S3 | 93.7% | 16.2 | 3.4% | 49.6% | 48% | 17.4 |
| DCA 48h, S1 | 93.5% | 15.1 | 2.8% | 49.1% | 48% | 16.2 |
| DCA 48h, S2 | 93.5% | 15.1 | 4.1% | 50.1% | 49% | 16.2 |
| DCA 48h, S3 | 93.5% | 16.5 | 3.8% | 51.1% | 49% | 17.7 |
| FGF2 48h, S1 | 93.0% | 15.2 | 3.3% | 50.7% | 49% | 16.4 |
| PBS 24h, S1 | 92.7% | 16.2 | 6.2% | 53.0% | 49% | 17.5 |
| FGF2 48h, S2 | 93.1% | 17.6 | 3.1% | 52.9% | 49% | 19.0 |
| FGF2 48h, S2 | 93.1% | 30.1 | 3.4% | 58.3% | 49% | 32.5 |
| DCA+FGF2 48h, S1 | 93.1% | 16.9 | 3.3% | 52.1% | 49% | 18.2 |
| DCA+FGF2 48h, S2 | 92.7% | 14.7 | 4.9% | 51.0% | 50% | 15.9 |
| DCA+FGF2 48h, S3 | 93.1% | 15.5 | 3.3% | 50.9% | 49% | 16.7 |
| PBS 24h, S2 | 93.5% | 15.6 | 3.2% | 49.6% | 49% | 16.8 |
| PBS 24h, S3 | 93.6% | 15.3 | 3.4% | 48.5% | 48% | 16.4 |
| DCA 24h, S1 | 93.8% | 15.3 | 3.8% | 49.6% | 48% | 16.3 |
| DCA 24h, S2 | 93.7% | 16.7 | 4.5% | 51.4% | 48% | 17.9 |
| DCA 24h, S3 | 93.5% | 19.0 | 4.4% | 53.2% | 49% | 20.4 |
| FGF2 24h, S1 | 93.6% | 15.6 | 3.3% | 50.9% | 49% | 16.7 |
| FGF2 24h, S2 | 93.5% | 14.1 | 3.6% | 49.1% | 49% | 15.1 |
| FGF2 24h, S3 | 93.5% | 28.9 | 4.5% | 58.0% | 49% | 31.0 |

B

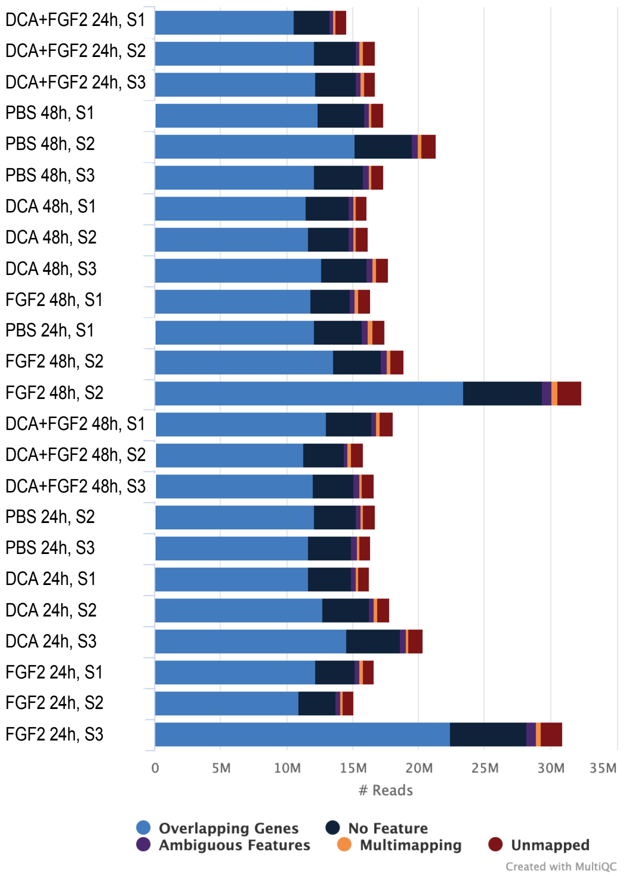

### Figure S3

A

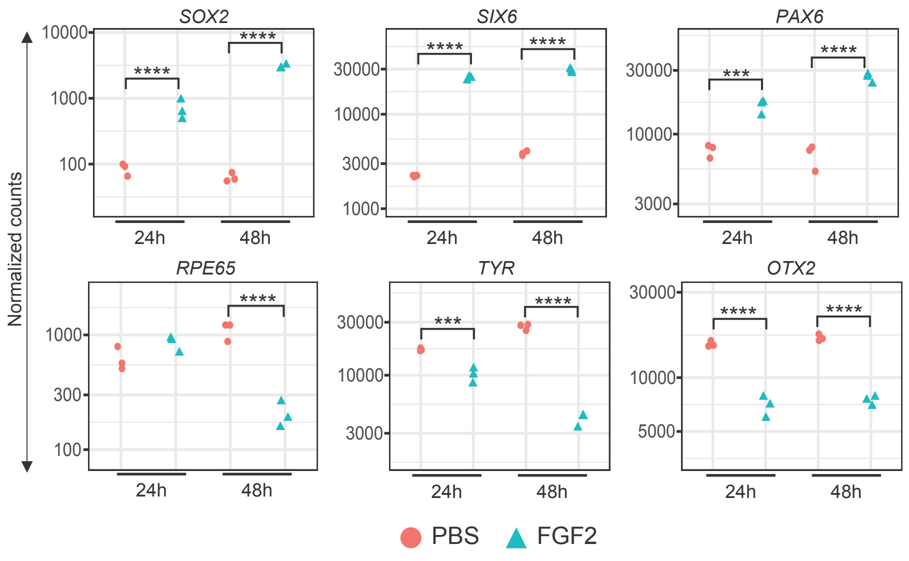
Normalized counts
SOX2
SIX6
PAX6
RPE65
TYR
OTX2
24h
48h
PBS
FGF2

B

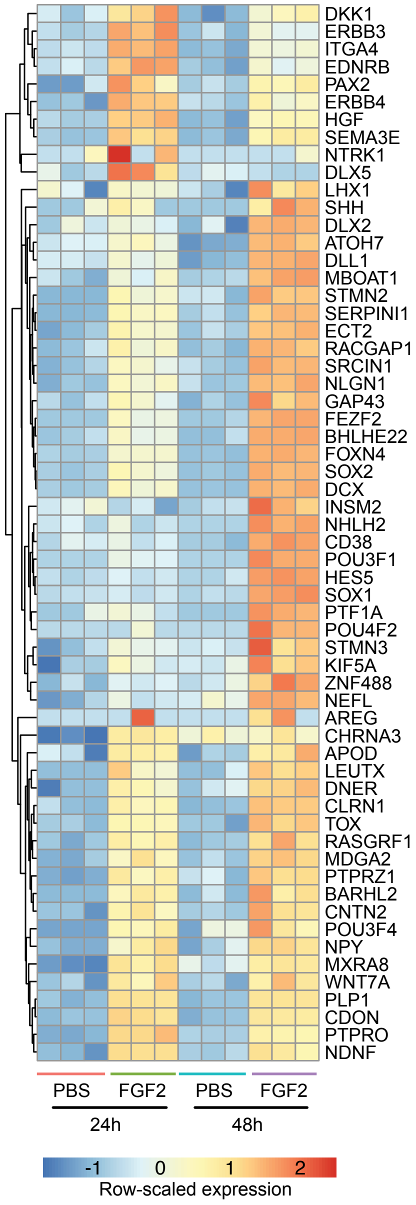
DKK1
ERBB3
ITGA4
EDNRB
PAX2
ERBB4
HGF
SEMA3E
NTRK1
DLX5
LHX1
SHH
DLX2
ATOH7
DLL1
MBOAT1
STMN2
SERPINI1
ECT2
RACGAP1
SRCIN1
NLGN1
GAP43
FEZF2
BHLHE22
FOXN4
SOX2
DCX
INSM2
NHLH2
CD38
POU3F1
HES5
SOX1
PTF1A
POU4F2
STMN3
KIF5A
ZNF488
NEFL
AREG
CHRNA3
APOD
LEUTX
DNER
CLRN1
TOX
RASGRF1
MDGA2
PTPRZ1
BARHL2
CNTN2
POU3F4
NPY
MXRA8
WNT7A
PLP1
CDON
PTPRO
NDNF
PBS
FGF2
PBS
FGF2
24h
48h
-1
0
1
2
Row-scaled expression

### Figure S4

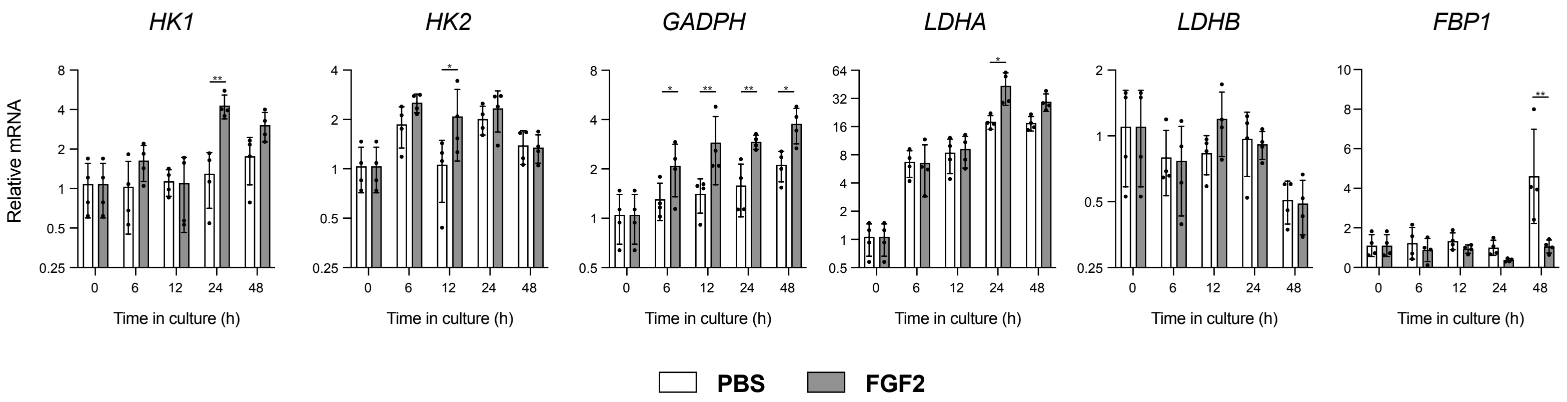

HK1
HK2
GADPH
LDHA
LDHB
FBP1
Relative mRNA
Time in culture (h)
0
6
12
24
48
PBS
FGF2

### Figure S5

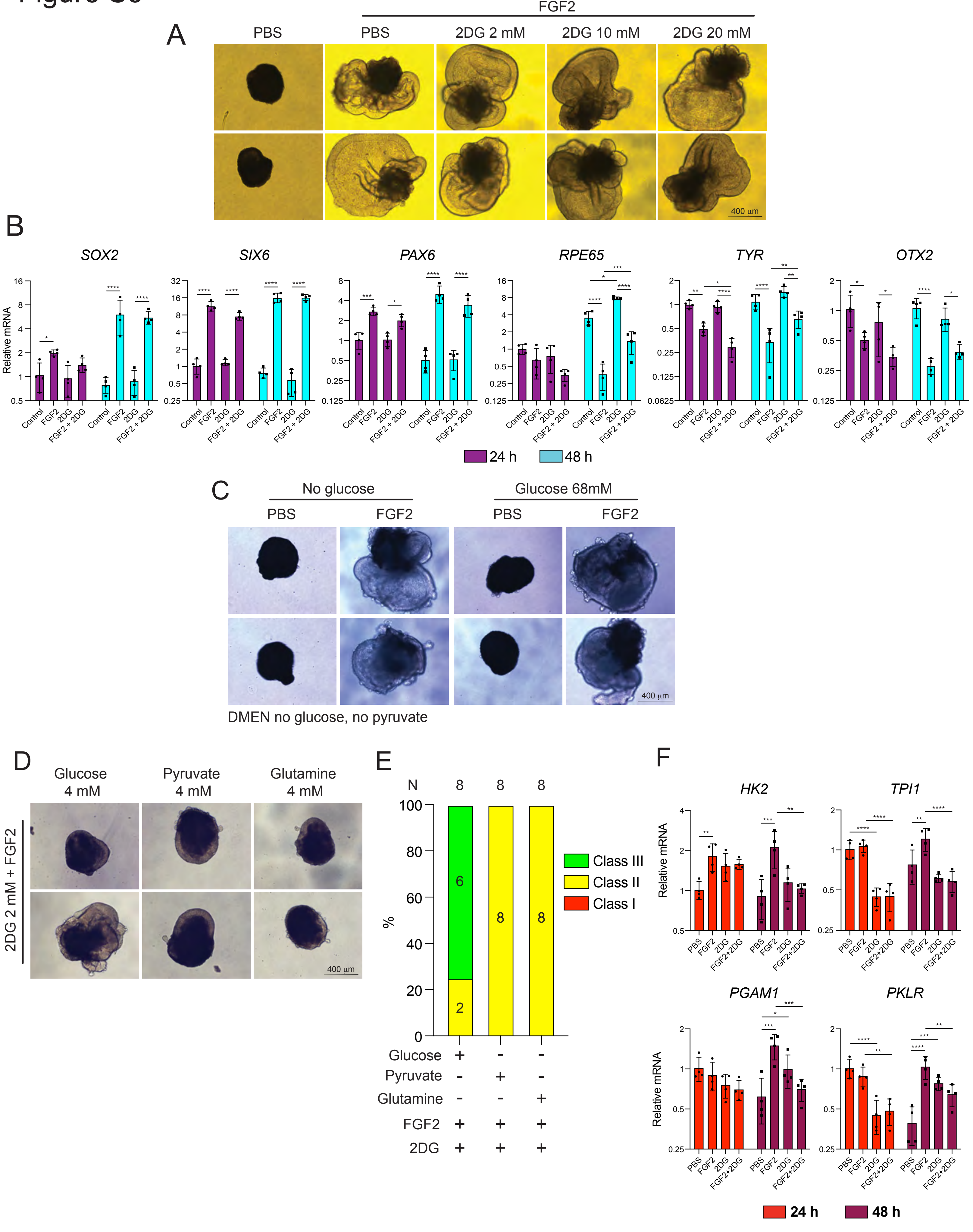

A
FGF2
PBS
PBS
2DG 2 mM
2DG 10 mM
2DG 20 mM
400 µm
B
SOX2
SIX6
PAX6
RPE65
TYR
OTX2
Relative mRNA
Control
FGF2
2DG
FGF2 + 2DG
24 h
48 h
C
No glucose
Glucose 68mM
PBS
FGF2
PBS
FGF2
400 µm
DMEN no glucose, no pyruvate
D
Glucose 4 mM
Pyruvate 4 mM
Glutamine 4 mM
2DG 2 mM + FGF2
400 µm
E
N
8
8
8
%
6
2
8
8
Class III
Class II
Class I
Glucose + - -
Pyruvate - + -
Glutamine - - +
FGF2 + + +
2DG + + +
F
HK2
TPI1
PGAM1
PKLR
Relative mRNA
PBS
FGF2
2DG
FGF2+2DG
24 h
48 h

### Figure S6

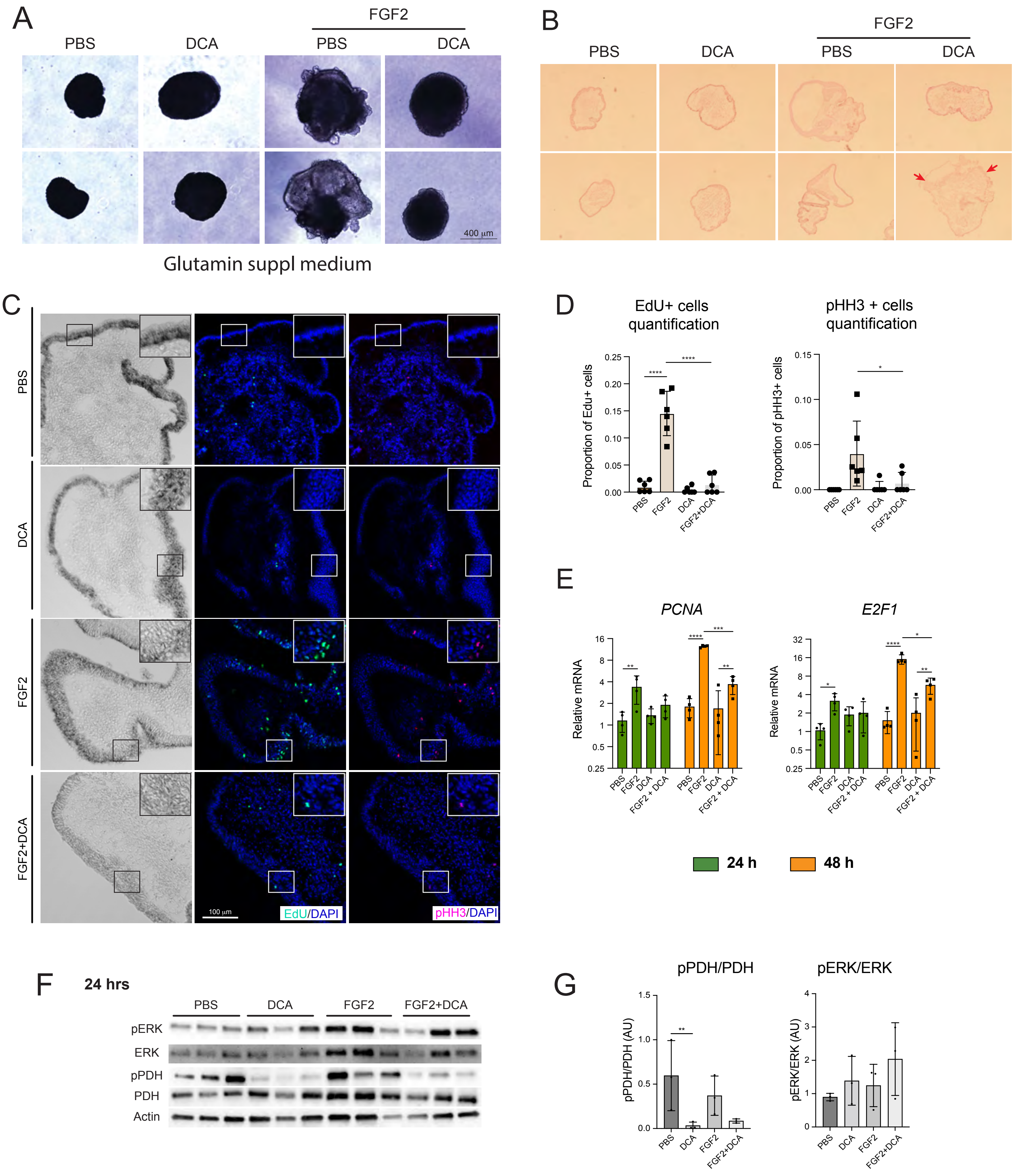

A
PBS
DCA
FGF2
PBS
DCA
400 μm
Glutamin suppl medium
B
PBS
DCA
FGF2
PBS
DCA
C
PBS
DCA
FGF2
FGF2+DCA
100 μm
EdU/DAPI
pHH3/DAPI
D
EdU+ cells quantification
Proportion of Edu+ cells
pHH3 + cells quantification
Proportion of pHH3+ cells
PBS
FGF2
DCA
FGF2+DCA
E
PCNA
E2F1
Relative mRNA
PBS
FGF2
DCA
FGF2 + DCA
24 h
48 h
F
24 hrs
PBS
DCA
FGF2
FGF2+DCA
pERK
ERK
pPDH
PDH
Actin
G
pPDH/PDH
pPDH/PDH (AU)
pERK/ERK
pERK/ERK (AU)

### Figure S7

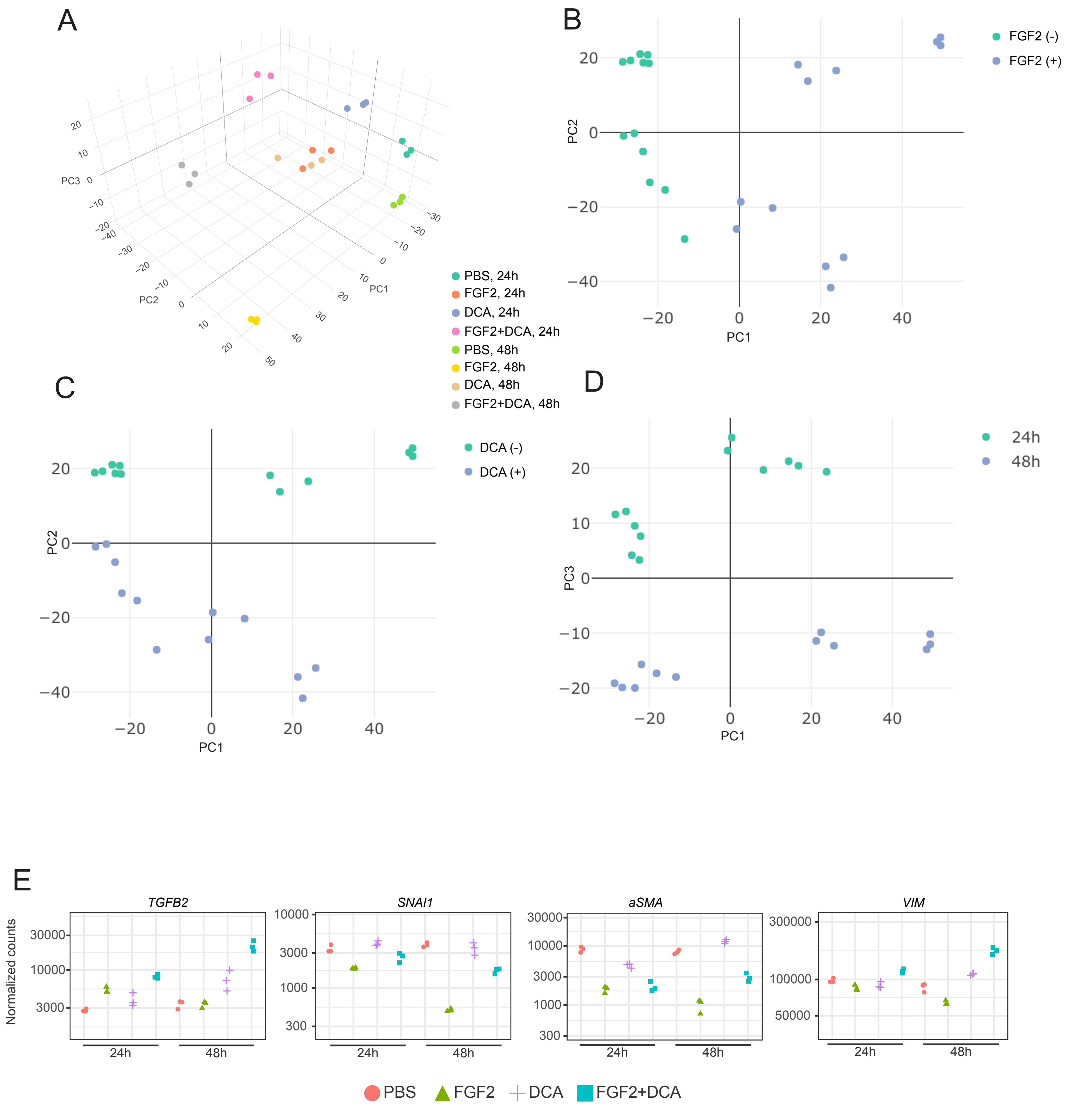

A
B
C
D
E
PC1
PC2
PC3
PBS, 24h
FGF2, 24h
DCA, 24h
FGF2+DCA, 24h
PBS, 48h
FGF2, 48h
DCA, 48h
FGF2+DCA, 48h
FGF2 (-)
FGF2 (+)
DCA (-)
DCA (+)
24h
48h
Normalized counts
TGFB2
SNAI1
aSMA
VIM
PBS
FGF2
DCA
FGF2+DCA

### Figure S8

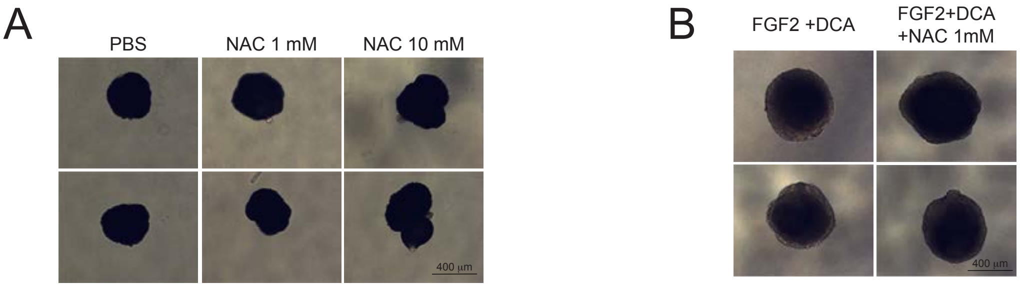
