## Supplementary material for "DISTINCT METABOLIC STATES DIRECT RETINAL PIGMENT EPITHELIUM CELL FATE DECISIONS": Stables

**Table 1S.** Antibodies used for WB and immunofluorescence

| Antibody | Ventor | Catalog | Concentration | Application |
|---|---|---|---|---|
| Anti-PDHA1 antibody [8D10E6] ab110334 | Abcam | ab110334 | 1 μg/mL | WB |
| Anti-phospho-Histone H3 (Ser10) Antibody | Millipore Sigma | 06-570 | 2.5 μg/mL | IF |
| Anti-rabbit IgG, HRP-linked Antibody | Cell Signaling | #7074 | Dil: 1:2500 | WB |
| Anti-β-Actin antibody, Mouse monoclonal | Millipore Sigma | A1978 | 0.01 μg/mL | WB |
| Goat anti-Mouse IgG (H+L) Highly Cross-Adsorbed Secondary Antibody, Alexa Fluor™ 546 | ThermoFisher | A-11030 | Dil: 1:300 | IF |
| Goat anti-Rabbit IgG (H+L) Highly Cross-Adsorbed Secondary Antibody, Alexa Fluor™ 568 | ThermoFisher | A-11036 | Dil: 1:300 | IF |
| p44/42 MAPK (Erk1/2) Antibody | Cell Signaling | #9102 | Dil: 1:750 | WB |
| Phospho-p44/42 MAPK (Erk1/2) | Cell Signaling | #9101 | Dil: 1:750 | WB |
| PhosphoDetect™ Anti-PDH-E1α ($pSer^{300}$) | Millipore Sigma | AP1064 | 1 μg/mL | WB |
| Recombinant Anti-Otx2 antibody | Abcam | ab183951 | Dil: 1:200 | IF |
| S100A4-1 | DSHB | CPTC-S100A4-1 | 5 μg/mL | IF |
| Sox-2 Antibody (E-4) | Santa Crus Biotechnology | sc-365823 | 1 μg/mL | IF |
| StarBright Blue 700 Goat Anti-Mouse IgG | Biorad | 12004158 | Dil: 1:1500 | WB |
| Tubulin, alpha | DSHB | 4A1 | 0.5 μg/mL | WB |

**IF: Immunofluorescence; WB: Western blot**

**Table 2S.** Primers used for RT-qPCR gene expression.

| Gene | GenBank ID | Forward primer | Reverse primer | PCR amplicon size (bp) | Reference |
|---|---|---|---|---|---|
| aSMA (ACTA2) | NM_001031229.1 | GTGCCAGCCATGTATGTAGC | ACACCATCCCCAGAGTCAAG | 92 | N/A |
| E2F1 | NM_205219.1 | GGACGATCTCATCCAGACGTG | ACGTAGGCTGCGTGCT | 78 | N/A |
| ENO1 | NM_205120.2 | ACGCTACTTGGGAAAAGGTGT | TGCTCCACCACGTTGACATT | 96 | N/A |
| FBP1 | NM_001278048.2 | TAATCTTGTGGCAGCGGGTT | TTCTCCGATTGCCGGATCAA | 109 | N/A |
| GADPH | NM_204305.1 | CCATGTTTGTGATGGGTGTC | CTCCACAATGCCAAAGTTGT | 131 | N/A |
| HK1 | NM_204101.1 | CAGTGGGACACGGCTTTTTG | CTTAGATTGTCGGCACGGGA | 124 | N/A |
| HK2 | NM_204212.2 | TAAGATCCGCGAGAACCGTG | CATGATGGCCGAGAAGTGTG | 97 | N/A |
| LDHA | NM_205284.2 | TTGGCCTTTCTGTGGCAGAT | ATTCCGTGCATGCCCTTAACA | 94 | N/A |
| LDHB | NM_204177.2 | ACGTTATGGCGACCCTGAAG | ATCACAAAGACCCTTGCCGA | 143 | N/A |
| OTX2 | NM_204520.2 | GTCGGTTATCCCGCCACC | TTTTCAAGGCCACCTCCTCC | 139 | N/A |
| PAX6 | NM_205066.1 | GGCAGAAGATCGTGGAACTC | TTCGTAATACCTGCCCAAAA | 152 | Luz-Madrigal et al., 2014 |
| PCNA | NM_204170.3 | GCGTCAACCTAAACAGCATGTC | GCCAACGTATCCGCATTGTC | 94 | N/A |
| PGAM1 | NM_001031556.3 | TGTCAAGCATTTGGAAGGCAT | CACGATTGGGATACCGGTGG | 73 | N/A |
| PKLR | NM_205469.1 | CATGCAGCACGCTATTGCTC | GTGGTGTACACTGTGGCGTA | 88 | N/A |
| PTX1 | NM_001167684.2 | TTTGGAGAGACTGCCCGAGA | GCTTCTCTCTTTTTCTGGCACC | 151 | N/A |
| RPE65 | NM_204884.1 | CCTACCACCGGAGGTTTGTT | GGTCTGGGTAGGCGTAGGTA | 99 | N/A |
| RPLP0 | NM_204987.2 | GGAGCTCACAGCTCGTCTTT | TAGTTGGACTTCCACGTCGC | 92 | N/A |
| SIX6 | NM_001389365.1 | AGGTGGGCAACTGGTTCAAA | CTGCTGCTGTAGCCTGTTCT | 74 | N/A |
| SNAI1 | NM_205142.1 | CCCTGTGTCTGCAAGATGTG | CGCAGATTAGAACGGTCAGC | 137 | N/A |
| SOX2 | NM_205188.2 | TGAACGGATCGCCTACCTAC | CTGGATTCCGTCTTGACCAC | 97 | Luz-Madrigal et al., 2014 |
| TGFB2 | NM_001031045.3 | GCCATCCCACCAAGCTATTA | AGTTGGACGCATTTTTCTCC | 82 | N/A |
| TYR | NM_204160.1 | TTTGCTGATCCACACACTGC | GATCATTCGCAGAGCCTTGT | 112 | N/A |
| VIM | NM_001048076.2 | TCGCCATCTTCGTGAGTACC | TTCTCCCTCCAGCAGTTTTC | 91 | N/A |
| TPI | NM_205451.1 | AGGAGAGATCAGCCCAGCAA | ATCCAGCTTCTCCCCAATGC | 172 | NA |
