## Supplementary material for "DISTINCT METABOLIC STATES DIRECT RETINAL PIGMENT EPITHELIUM CELL FATE DECISIONS": Statistical analysis

### Statistical Supplementary Material for “Distinct Metabolic States Direct Retinal Pigment Epithelium Cell Fate Decisions” by Perez-Estrada et al.

We have numbered the experiments and included cross-references with the Figures in the paper and/or supplementary material. For each experiment, we provide a basic description and its statistical analysis.

Note that for the experiments with non-count response variables, we display residual plots throughout in order to assess the congruence of model assumptions (most importantly, that all experimental units have the same variance) with the datasets. There are many such models, and so it is difficult to assess and respond to each plot/model. Most potential departures are the result of non-extreme outliers, and we note that in such cases the outliers actually inflate the error variance estimate, compared to a similar dataset without an outlier, resulting in conservative inference. Overall, violations of model assumptions is not overly serious or pervasive.

### Experiment 1 (Figures 1C, S4)

Response is relative gene expression of a number of genes; two factors: Chemical (levels: PBS, FGF2) and Time (levels: 0, 6, 12, 24, 48). However, the factors are not crossed because there is no response for FGF2 at Time 0. So 9 treatments, 4 replications per treatment, for a total of 36 experimental units. For this and future relative gene expression responses, we transform using log base 2.

Comparisons for Figure 1C (and S4), for each gene:

- PBS 0h vs PBS 6h, 12h, 24h, and 48h
- PBS 0h vs FGF2 6h, 12h, 24h, and 48h
- PBS vs FGF2 at each time point (6h, 12h, 24h, and 48h)

#### Data and Visualization

For each of these datasets (Fig 1C, Fig S4), we perform two different analyses. One for the first two types of comparisons and one for the last one.

##### Plots for Comparison with Control

First, we compare `PBS 0h` to the other eight treatments.

```
d1C <- read.xlsx(xlsxFile = "MetabolismPaperRawData_Byran_working.xlsx",
                 colNames = TRUE, sheet="Fig 1_C")
glimpse(d1C)
```

```
## Rows: 36
## Columns: 9
## $ Chemical   <chr> "PBS", "PBS", "PBS", "PBS", "PBS", "PBS", "PBS", "PBS", "FG~
## $ Sample     <dbl> 1, 2, 3, 4, 1, 2, 3, 4, 1, 2, 3, 4, 1, 2, 3, 4, 1, 2, 3, 4,~
## $ `Time.(h)` <dbl> 0, 0, 0, 0, 6, 6, 6, 6, 6, 6, 6, 6, 12, 12, 12, 12, 12, 12,~
## $ SOX2       <dbl> 1.5574696, 0.2556121, 1.0195235, 2.4637789, 0.9540158, 1.07~
## $ SIX6       <dbl> 1.0291093, 0.6182058, 1.2174786, 1.2910531, 13.7008965, 13.~
## $ PAX6       <dbl> 0.9840290, 0.5874487, 1.0243051, 1.6888565, 0.7258499, 0.70~
## $ RPE65      <dbl> 0.9828628, 0.5734134, 1.2908035, 1.3746088, 0.3257529, 0.20~
## $ TYR        <dbl> 1.1031137, 0.6265110, 1.1446581, 1.2640819, 1.1734015, 0.74~
## $ OTX2       <dbl> 1.0346092, 0.8664992, 0.9119958, 1.2231020, 0.9470746, 0.73~
```

```
dS4 <- read.xlsx(xlsxFile = "MetabolismPaperRawData_Byran_working.xlsx",
                 colNames = TRUE, sheet="Fig 4S")
glimpse(dS4)
```

```
## Rows: 36
## Columns: 9
## $ Chemical   <chr> "PBS", "PBS", "PBS", "PBS", "PBS", "PBS", "PBS", "PBS", "FG~
## $ Sample     <dbl> 1, 2, 3, 4, 1, 2, 3, 4, 1, 2, 3, 4, 1, 2, 3, 4, 1, 2, 3, 4,~
## $ `Time.(h)` <dbl> 0, 0, 0, 0, 6, 6, 6, 6, 6, 6, 6, 6, 12, 12, 12, 12, 12, 12,~
## $ HK1        <dbl> 0.7863192, 0.6185946, 1.6950020, 1.2128993, 0.5300377, 1.08~
## $ HK2        <dbl> 1.0842404, 0.8523087, 1.4653182, 0.7384915, 1.1842694, 2.11~
## $ GADPH      <dbl> 1.4746027, 0.6379957, 0.9396206, 1.1312399, 1.7912290, 1.02~
## $ LDHA       <dbl> 1.2545427, 0.5756463, 1.4962003, 0.9254844, 6.0227023, 4.20~
## $ LDHB       <dbl> 1.4994826, 0.5281534, 1.5729342, 0.8027641, 0.6576684, 0.68~
## $ FBP1       <dbl> 1.2372306, 0.7248728, 1.8258234, 0.6107012, 0.7015782, 0.42~
```

```
# to compare PBS 0h with everything else
d1C_1 <- d1C %>%
  rename(Time=`Time.(h)`) %>%
  mutate(Treatment=paste(Chemical,Time, sep="_"))
glimpse(d1C_1)
```

```
## Rows: 36
## Columns: 10
## $ Chemical  <chr> "PBS", "PBS", "PBS", "PBS", "PBS", "PBS", "PBS", "PBS", "FGF~
## $ Sample    <dbl> 1, 2, 3, 4, 1, 2, 3, 4, 1, 2, 3, 4, 1, 2, 3, 4, 1, 2, 3, 4, ~
## $ Time      <dbl> 0, 0, 0, 0, 6, 6, 6, 6, 6, 6, 6, 6, 12, 12, 12, 12, 12, 12, ~
## $ SOX2      <dbl> 1.5574696, 0.2556121, 1.0195235, 2.4637789, 0.9540158, 1.077~
## $ SIX6      <dbl> 1.0291093, 0.6182058, 1.2174786, 1.2910531, 13.7008965, 13.5~
## $ PAX6      <dbl> 0.9840290, 0.5874487, 1.0243051, 1.6888565, 0.7258499, 0.700~
## $ RPE65     <dbl> 0.9828628, 0.5734134, 1.2908035, 1.3746088, 0.3257529, 0.203~
## $ TYR       <dbl> 1.1031137, 0.6265110, 1.1446581, 1.2640819, 1.1734015, 0.743~
## $ OTX2      <dbl> 1.0346092, 0.8664992, 0.9119958, 1.2231020, 0.9470746, 0.733~
## $ Treatment <chr> "PBS_0", "PBS_0", "PBS_0", "PBS_0", "PBS_6", "PBS_6", "PBS_6~
```

```
dS4_1 <- dS4 %>%
  rename(Time=`Time.(h)`) %>%
  mutate(Treatment=paste(Chemical,Time, sep="_"))
glimpse(dS4_1)
```

```
## Rows: 36
## Columns: 10
## $ Chemical  <chr> "PBS", "PBS", "PBS", "PBS", "PBS", "PBS", "PBS", "PBS", "FGF~
## $ Sample    <dbl> 1, 2, 3, 4, 1, 2, 3, 4, 1, 2, 3, 4, 1, 2, 3, 4, 1, 2, 3, 4, ~
## $ Time      <dbl> 0, 0, 0, 0, 6, 6, 6, 6, 6, 6, 6, 6, 12, 12, 12, 12, 12, 12, ~
## $ HK1       <dbl> 0.7863192, 0.6185946, 1.6950020, 1.2128993, 0.5300377, 1.082~
## $ HK2       <dbl> 1.0842404, 0.8523087, 1.4653182, 0.7384915, 1.1842694, 2.119~
## $ GADPH     <dbl> 1.4746027, 0.6379957, 0.9396206, 1.1312399, 1.7912290, 1.025~
## $ LDHA      <dbl> 1.2545427, 0.5756463, 1.4962003, 0.9254844, 6.0227023, 4.200~
## $ LDHB      <dbl> 1.4994826, 0.5281534, 1.5729342, 0.8027641, 0.6576684, 0.689~
## $ FBP1      <dbl> 1.2372306, 0.7248728, 1.8258234, 0.6107012, 0.7015782, 0.425~
## $ Treatment <chr> "PBS_0", "PBS_0", "PBS_0", "PBS_0", "PBS_6", "PBS_6", "PBS_6~
```

```
d1C_1long <- d1C_1 %>%
  pivot_longer(cols=4:9,names_to="Gene",values_to="RGE")
glimpse(d1C_1long)
```

```
## Rows: 216
## Columns: 6
## $ Chemical  <chr> "PBS", "PBS", "PBS", "PBS", "PBS", "PBS", "PBS", "PBS", "PBS~
## $ Sample    <dbl> 1, 1, 1, 1, 1, 1, 2, 2, 2, 2, 2, 2, 3, 3, 3, 3, 3, 3, 4, 4, ~
## $ Time      <dbl> 0, 0, 0, 0, 0, 0, 0, 0, 0, 0, 0, 0, 0, 0, 0, 0, 0, 0, 0, 0, ~
## $ Treatment <chr> "PBS_0", "PBS_0", "PBS_0", "PBS_0", "PBS_0", "PBS_0", "PBS_0~
## $ Gene      <chr> "SOX2", "SIX6", "PAX6", "RPE65", "TYR", "OTX2", "SOX2", "SIX~
## $ RGE       <dbl> 1.5574696, 1.0291093, 0.9840290, 0.9828628, 1.1031137, 1.034~
```

```
ggplot(d1C_1long, aes(x=Treatment,y=log2(RGE))) +
  geom_jitter(width=.05,height=0)+
  facet_wrap(~Gene, nrow=3)+
  theme(axis.text.x = element_text(angle = 90, vjust = 0.5, hjust=1)) +
  labs(title="For Figure 1C: Comparisons with Control")
```

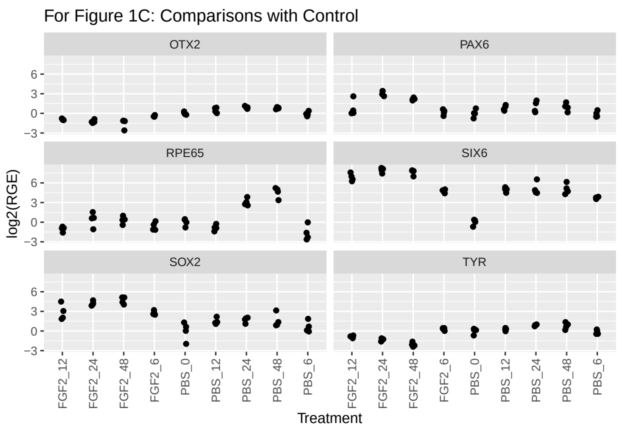

```
dS4_1long <- dS4_1 %>%
  pivot_longer(cols=4:9,names_to="Gene",values_to="RGE")
glimpse(dS4_1long)
```

```
## Rows: 216
```

```
## Columns: 6
## $ Chemical  <chr> "PBS", "PBS", "PBS", "PBS", "PBS", "PBS", "PBS", "PBS", "PBS~
## $ Sample    <dbl> 1, 1, 1, 1, 1, 1, 2, 2, 2, 2, 2, 2, 3, 3, 3, 3, 3, 3, 4, 4, ~
## $ Time      <dbl> 0, 0, 0, 0, 0, 0, 0, 0, 0, 0, 0, 0, 0, 0, 0, 0, 0, 0, 0, 0, ~
## $ Treatment <chr> "PBS_0", "PBS_0", "PBS_0", "PBS_0", "PBS_0", "PBS_0", "PBS_0~
## $ Gene      <chr> "HK1", "HK2", "GADPH", "LDHA", "LDHB", "FBP1", "HK1", "HK2",~
## $ RGE       <dbl> 0.7863192, 1.0842404, 1.4746027, 1.2545427, 1.4994826, 1.237~
```

```
ggplot(dS4_1long, aes(x=Treatment,y=log2(RGE))) +
  geom_jitter(width=.05,height=0)+
  facet_wrap(~Gene, nrow=3)+
  theme(axis.text.x = element_text(angle = 90, vjust = 0.5, hjust=1)) +
  labs(title="For Figure S4: Comparisons with Control")
```

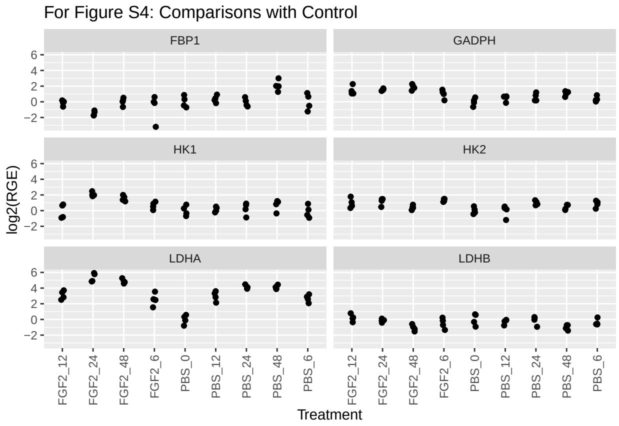

##### Plots to Compare PBS vs. FGF2 at each Time Point

Now, we compare PBS vs. FGF2 at each time point. To facilitate this, we omit PBS at Time 0 (because FGF2 not measured at Time 0).

```
# to compare PBS vs. FGF2 at each time point (omit PBS_0 and use as two factors)
d1C_2long <- d1C %>%
  rename(Time=`Time.(h)`) %>%
  filter(Time != 0) %>%
  pivot_longer(cols=4:9,names_to="Gene",values_to="RGE") %>%
  mutate(Time=as.factor(Time))
glimpse(d1C_2long)
```

```
## Rows: 192
## Columns: 5
## $ Chemical <chr> "PBS", "PBS", "PBS", "PBS", "PBS", "PBS", "PBS", "PBS", "PBS"~
## $ Sample   <dbl> 1, 1, 1, 1, 1, 1, 2, 2, 2, 2, 2, 2, 3, 3, 3, 3, 3, 3, 4, 4, 4~
## $ Time     <fct> 6, 6, 6, 6, 6, 6, 6, 6, 6, 6, 6, 6, 6, 6, 6, 6, 6, 6, 6, 6, 6~
## $ Gene     <chr> "SOX2", "SIX6", "PAX6", "RPE65", "TYR", "OTX2", "SOX2", "SIX6~
## $ RGE      <dbl> 0.9540158, 13.7008965, 0.7258499, 0.3257529, 1.1734015, 0.947~
```

```
dS4_2long <- dS4 %>%
  rename(Time=`Time.(h)`) %>%
```

```r
  filter(Time != 0) %>%
  pivot_longer(cols=4:9,names_to="Gene",values_to="RGE") %>%
  mutate(Time=as.factor(Time))
glimpse(dS4_2long)
```

```
## Rows: 192
## Columns: 5
## $ Chemical <chr> "PBS", "PBS", "PBS", "PBS", "PBS", "PBS", "PBS", "PBS", "PBS"~
## $ Sample   <dbl> 1, 1, 1, 1, 1, 1, 2, 2, 2, 2, 2, 2, 3, 3, 3, 3, 3, 3, 4, 4, 4~
## $ Time     <fct> 6, 6, 6, 6, 6, 6, 6, 6, 6, 6, 6, 6, 6, 6, 6, 6, 6, 6, 6, 6, 6~
## $ Gene     <chr> "HK1", "HK2", "GADPH", "LDHA", "LDHB", "FBP1", "HK1", "HK2", ~
## $ RGE      <dbl> 0.5300377, 1.1842694, 1.7912290, 6.0227023, 0.6576684, 0.7015~
```

```r
ggplot(d1C_2long, aes(x=Time,y=log2(RGE),color=Chemical)) +
  geom_jitter(width=.05,height=0) +
  facet_wrap(~Gene, nrow=3) +
  labs(title="For Figure 1C: Time Comparisons")
```

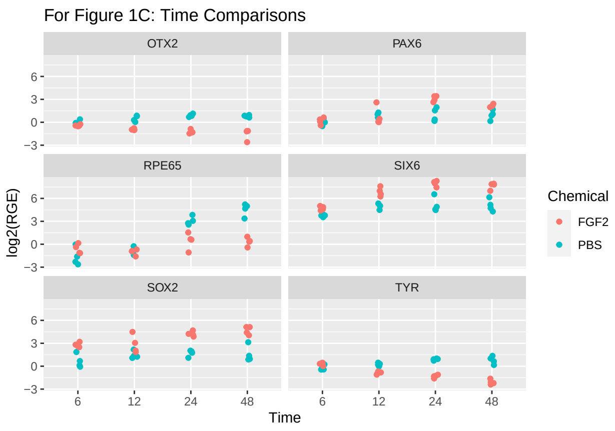

```r
ggplot(dS4_2long, aes(x=Time,y=log2(RGE),color=Chemical)) +
  geom_jitter(width=.05,height=0) +
  facet_wrap(~Gene, nrow=3) +
  labs(title="For Figure S4: Time Comparisons")
```

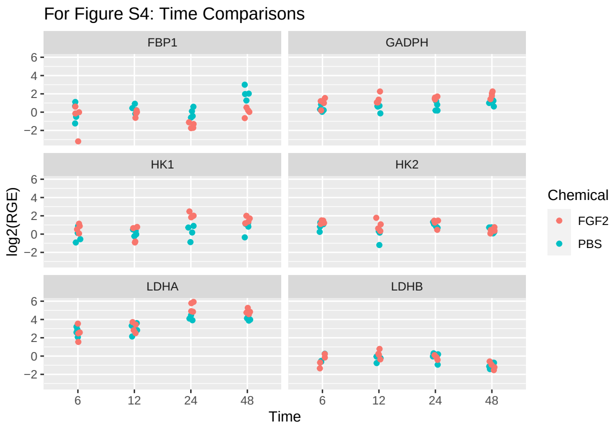

#### Data Analysis

Again, we have two different analyses. The first is comparing the 8 PBS/FGF2 treatments at Time > 0 with the baseline “PBS at Time 0” treatment. The second is to compare the two Chemical treatments at each time point.

##### Comparison with Control

For the first, we’ll fit a one-way ANOVA, where the factor has nine levels.

**Figure 1C** We first fit the models, then examine residual plots.

```
a1C_1SOX2 <- aov(log2(SOX2)~Treatment, data=d1C_1)
a1C_1SIX6 <- aov(log2(SIX6)~Treatment, data=d1C_1)
a1C_1PAX6 <- aov(log2(PAX6)~Treatment, data=d1C_1)
a1C_1RPE65 <- aov(log2(RPE65)~Treatment, data=d1C_1)
a1C_1TYR <- aov(log2(TYR)~Treatment, data=d1C_1)
a1C_1OTX2 <- aov(log2(OTX2)~Treatment, data=d1C_1)
plot(a1C_1SOX2,which=1:2)
```

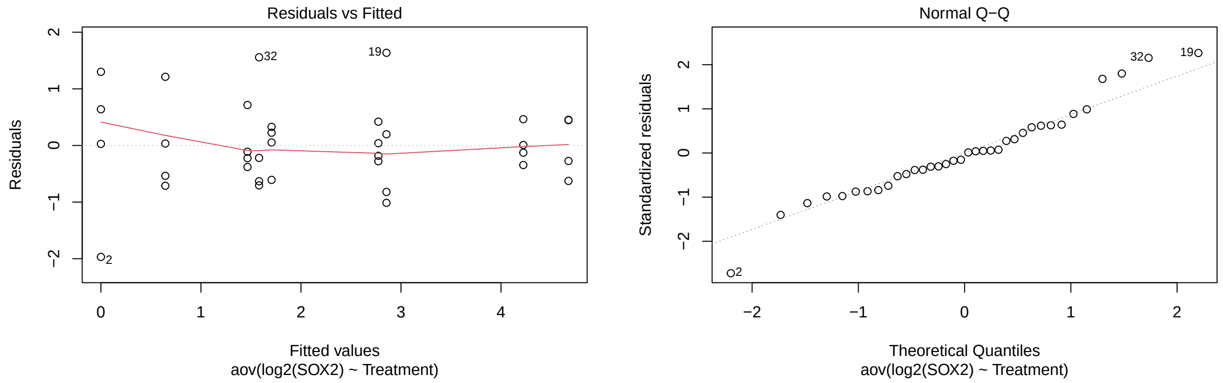

```
plot(a1C_1SIX6,which=1:2)
```

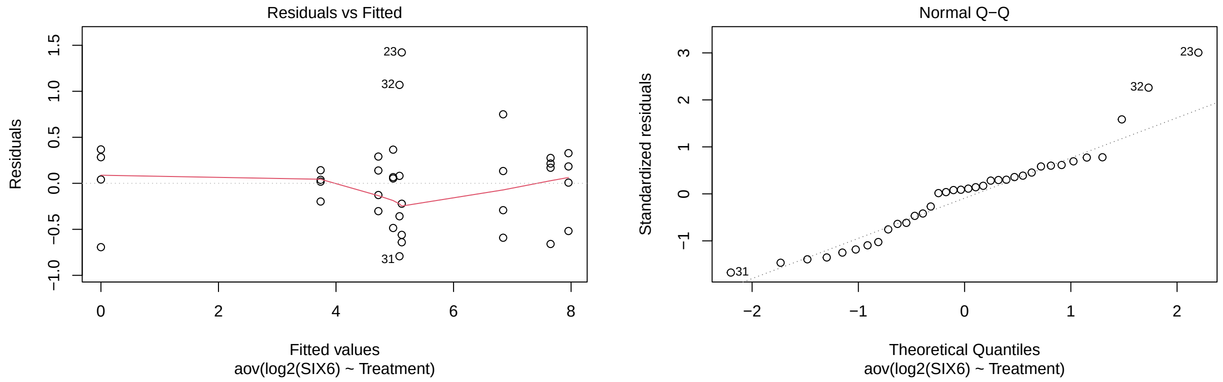

```
plot(a1C_1PAX6,which=1:2)
```

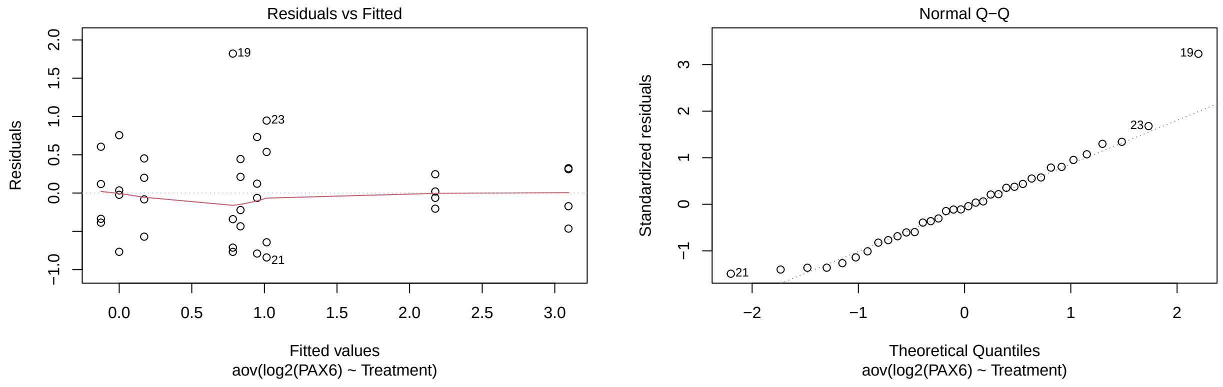

```
plot(a1C_1RPE65,which=1:2)
```

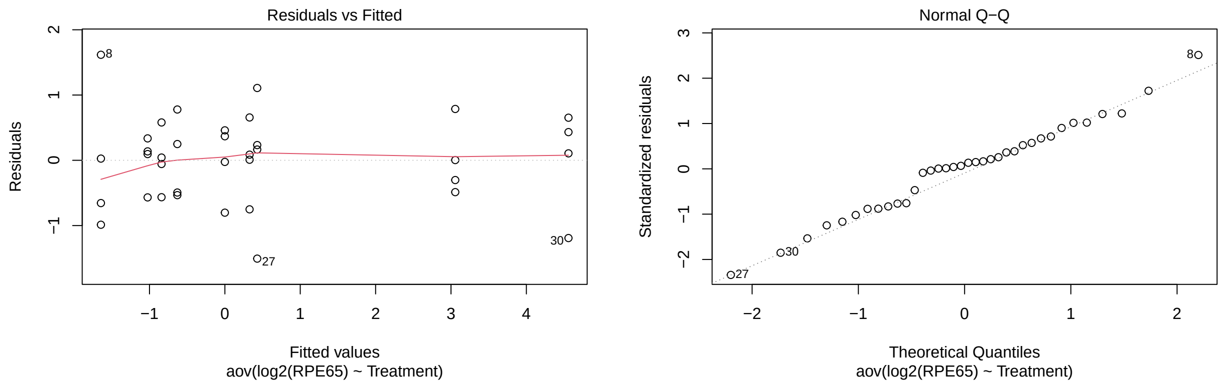

```
plot(a1C_1TYR,which=1:2)
```

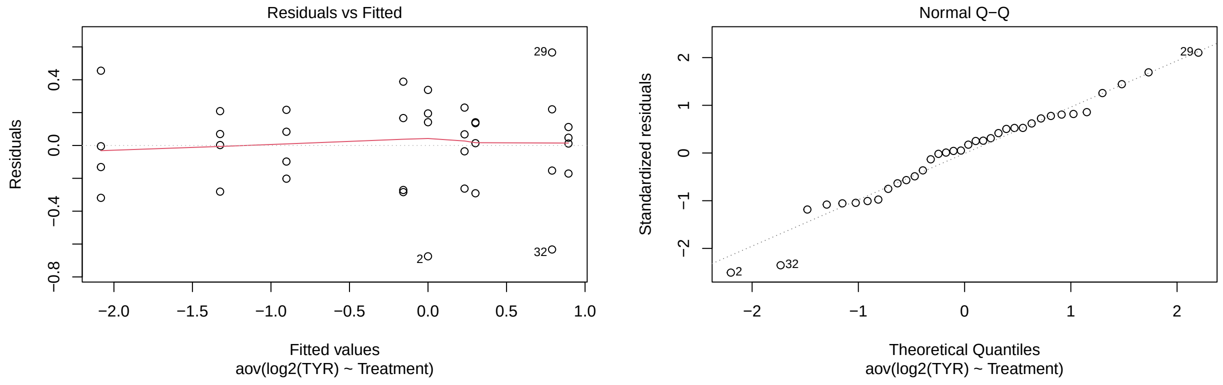

```
plot(a1C_1OTX2,which=1:2)
```

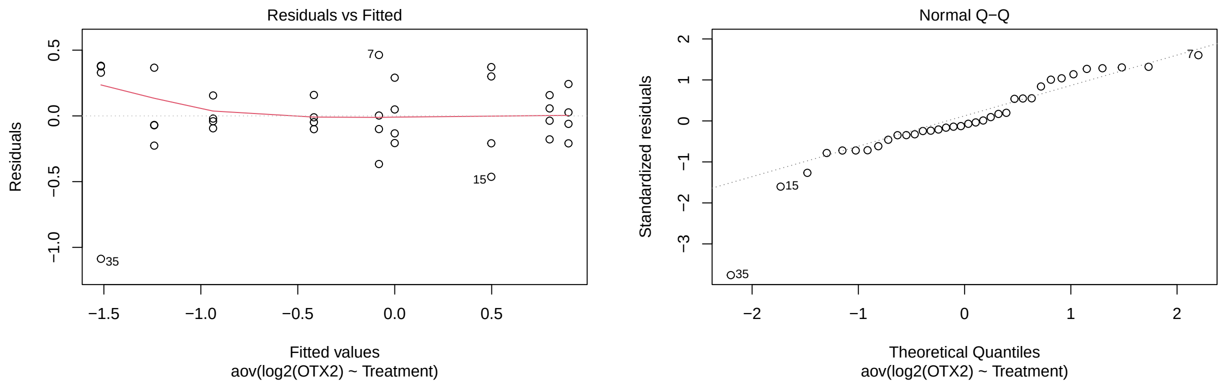

```r
emmeans(a1C_1SOX2,trt.vs.ctrlk ~ Treatment, ref=5)
```

```
## $emmeans
##  Treatment emmean    SE df lower.CL upper.CL
##  FGF2_12    2.855 0.417 27    1.999    3.711
##  FGF2_24    4.223 0.417 27    3.367    5.079
##  FGF2_48    4.675 0.417 27    3.819    5.531
##  FGF2_6     2.774 0.417 27    1.918    3.630
##  PBS_0      0.000 0.417 27   -0.856    0.856
##  PBS_12     1.466 0.417 27    0.610    2.322
##  PBS_24     1.706 0.417 27    0.851    2.562
##  PBS_48     1.581 0.417 27    0.725    2.437
##  PBS_6      0.644 0.417 27   -0.212    1.500
##
## Results are given on the log2 (not the response) scale.
## Confidence level used: 0.95
##
## $contrasts
##  contrast        estimate   SE df t.ratio p.value
##  FGF2_12 - PBS_0    2.855 0.59 27   4.840  0.0003
##  FGF2_24 - PBS_0    4.223 0.59 27   7.159  <.0001
##  FGF2_48 - PBS_0    4.675 0.59 27   7.925  <.0001
##  FGF2_6 - PBS_0     2.774 0.59 27   4.702  0.0005
##  PBS_12 - PBS_0     1.466 0.59 27   2.485  0.1106
##  PBS_24 - PBS_0     1.706 0.59 27   2.893  0.0463
##  PBS_48 - PBS_0     1.581 0.59 27   2.681  0.0736
##  PBS_6 - PBS_0      0.644 0.59 27   1.091  0.7842
##
## Results are given on the log2 (not the response) scale.
## P value adjustment: dunnettx method for 8 tests
```

```r
emmeans(a1C_1SIX6,trt.vs.ctrlk ~ Treatment, ref=5)
```

```
## $emmeans
##  Treatment emmean    SE df lower.CL upper.CL
##  FGF2_12     6.84 0.273 27    6.283    7.404
##  FGF2_24     7.96 0.273 27    7.395    8.516
##  FGF2_48     7.65 0.273 27    7.090    8.211
```

```
##  FGF2_6      4.72 0.273 27    4.160    5.281
##  PBS_0       0.00 0.273 27   -0.561    0.561
##  PBS_12      4.97 0.273 27    4.412    5.533
##  PBS_24      5.12 0.273 27    4.558    5.679
##  PBS_48      5.08 0.273 27    4.520    5.641
##  PBS_6       3.74 0.273 27    3.178    4.299
##
## Results are given on the log2 (not the response) scale.
## Confidence level used: 0.95
##
## $contrasts
##  contrast        estimate    SE df t.ratio p.value
##  FGF2_12 - PBS_0     6.84 0.386 27  17.712  <.0001
##  FGF2_24 - PBS_0     7.96 0.386 27  20.588  <.0001
##  FGF2_48 - PBS_0     7.65 0.386 27  19.800  <.0001
##  FGF2_6 - PBS_0      4.72 0.386 27  12.216  <.0001
##  PBS_12 - PBS_0      4.97 0.386 27  12.868  <.0001
##  PBS_24 - PBS_0      5.12 0.386 27  13.247  <.0001
##  PBS_48 - PBS_0      5.08 0.386 27  13.147  <.0001
##  PBS_6 - PBS_0       3.74 0.386 27   9.674  <.0001
##
## Results are given on the log2 (not the response) scale.
## P value adjustment: dunnettx method for 8 tests
```

```
emmeans(a1C_1PAX6,trt.vs.ctrlk ~ Treatment, ref=5)
```

```
## $emmeans
##  Treatment emmean    SE df lower.CL upper.CL
##  FGF2_12    0.783 0.325 27    0.115    1.450
##  FGF2_24    3.094 0.325 27    2.427    3.762
##  FGF2_48    2.177 0.325 27    1.509    2.844
##  FGF2_6     0.172 0.325 27   -0.495    0.840
##  PBS_0      0.000 0.325 27   -0.668    0.668
##  PBS_12     0.835 0.325 27    0.168    1.503
##  PBS_24     1.015 0.325 27    0.348    1.683
##  PBS_48     0.950 0.325 27    0.282    1.617
##  PBS_6     -0.126 0.325 27   -0.793    0.542
##
## Results are given on the log2 (not the response) scale.
## Confidence level used: 0.95
##
## $contrasts
##  contrast        estimate   SE df t.ratio p.value
##  FGF2_12 - PBS_0    0.783 0.46 27   1.702  0.4225
##  FGF2_24 - PBS_0    3.094 0.46 27   6.725  <.0001
##  FGF2_48 - PBS_0    2.177 0.46 27   4.731  0.0005
##  FGF2_6 - PBS_0     0.172 0.46 27   0.374  0.9925
##  PBS_12 - PBS_0     0.835 0.46 27   1.816  0.3595
##  PBS_24 - PBS_0     1.015 0.46 27   2.207  0.1885
##  PBS_48 - PBS_0     0.950 0.46 27   2.065  0.2422
##  PBS_6 - PBS_0     -0.126 0.46 27  -0.274  0.9974
##
## Results are given on the log2 (not the response) scale.
## P value adjustment: dunnettx method for 8 tests
```

```r
emmeans(a1C_1RPE65,trt.vs.ctrlk ~ Treatment, ref=5)
```

```
## $emmeans
##  Treatment emmean    SE df lower.CL upper.CL
##  FGF2_12   -1.025 0.371 27   -1.787  -0.2625
##  FGF2_24    0.429 0.371 27   -0.333   1.1915
##  FGF2_48    0.328 0.371 27   -0.434   1.0901
##  FGF2_6    -0.631 0.371 27   -1.393   0.1317
##  PBS_0      0.000 0.371 27   -0.762   0.7622
##  PBS_12    -0.840 0.371 27   -1.602  -0.0779
##  PBS_24     3.057 0.371 27    2.294   3.8188
##  PBS_48     4.559 0.371 27    3.796   5.3208
##  PBS_6     -1.644 0.371 27   -2.407  -0.8821
##
## Results are given on the log2 (not the response) scale.
## Confidence level used: 0.95
##
## $contrasts
##  contrast        estimate    SE df t.ratio p.value
##  FGF2_12 - PBS_0   -1.025 0.525 27  -1.950  0.2924
##  FGF2_24 - PBS_0    0.429 0.525 27   0.817  0.9051
##  FGF2_48 - PBS_0    0.328 0.525 27   0.624  0.9589
##  FGF2_6 - PBS_0    -0.631 0.525 27  -1.200  0.7243
##  PBS_12 - PBS_0    -0.840 0.525 27  -1.599  0.4825
##  PBS_24 - PBS_0     3.057 0.525 27   5.818  <.0001
##  PBS_48 - PBS_0     4.559 0.525 27   8.677  <.0001
##  PBS_6 - PBS_0     -1.644 0.525 27  -3.130  0.0269
##
## Results are given on the log2 (not the response) scale.
## P value adjustment: dunnettx method for 8 tests
```

```r
emmeans(a1C_1TYR,trt.vs.ctrlk ~ Treatment, ref=5)
```

```
## $emmeans
##  Treatment emmean    SE df lower.CL upper.CL
##  FGF2_12   -0.902 0.155 27  -1.2202   -0.583
##  FGF2_24   -1.324 0.155 27  -1.6427   -1.005
##  FGF2_48   -2.083 0.155 27  -2.4015   -1.764
##  FGF2_6     0.302 0.155 27  -0.0166    0.621
##  PBS_0      0.000 0.155 27  -0.3187    0.319
##  PBS_12     0.233 0.155 27  -0.0862    0.551
##  PBS_24     0.894 0.155 27   0.5757    1.213
##  PBS_48     0.790 0.155 27   0.4712    1.109
##  PBS_6     -0.157 0.155 27  -0.4761    0.161
##
## Results are given on the log2 (not the response) scale.
## Confidence level used: 0.95
##
## $contrasts
##  contrast        estimate   SE df t.ratio p.value
##  FGF2_12 - PBS_0   -0.902 0.22 27  -4.104  0.0024
##  FGF2_24 - PBS_0   -1.324 0.22 27  -6.027  <.0001
##  FGF2_48 - PBS_0   -2.083 0.22 27  -9.481  <.0001
```

```
##  FGF2_6 - PBS_0     0.302 0.22 27   1.375  0.6198
##  PBS_12 - PBS_0     0.233 0.22 27   1.059  0.8010
##  PBS_24 - PBS_0     0.894 0.22 27   4.072  0.0026
##  PBS_48 - PBS_0     0.790 0.22 27   3.596  0.0087
##  PBS_6 - PBS_0     -0.157 0.22 27  -0.717  0.9364
##
## Results are given on the log2 (not the response) scale.
## P value adjustment: dunnettx method for 8 tests
```

```
emmeans(a1C_1OTX2,trt.vs.ctrlk ~ Treatment, ref=5)
```

```
## $emmeans
##  Treatment  emmean    SE df lower.CL upper.CL
##  FGF2_12   -0.9368 0.167 27   -1.279  -0.5945
##  FGF2_24   -1.2404 0.167 27   -1.583  -0.8981
##  FGF2_48   -1.5152 0.167 27   -1.858  -1.1729
##  FGF2_6    -0.4168 0.167 27   -0.759  -0.0745
##  PBS_0      0.0000 0.167 27   -0.342   0.3423
##  PBS_12     0.4980 0.167 27    0.156   0.8403
##  PBS_24     0.8960 0.167 27    0.554   1.2383
##  PBS_48     0.7988 0.167 27    0.456   1.1411
##  PBS_6     -0.0814 0.167 27   -0.424   0.2610
##
## Results are given on the log2 (not the response) scale.
## Confidence level used: 0.95
##
## $contrasts
##  contrast        estimate    SE df t.ratio p.value
##  FGF2_12 - PBS_0  -0.9368 0.236 27  -3.971  0.0034
##  FGF2_24 - PBS_0  -1.2404 0.236 27  -5.257  0.0001
##  FGF2_48 - PBS_0  -1.5152 0.236 27  -6.422  <.0001
##  FGF2_6 - PBS_0   -0.4168 0.236 27  -1.767  0.3860
##  PBS_12 - PBS_0    0.4980 0.236 27   2.111  0.2238
##  PBS_24 - PBS_0    0.8960 0.236 27   3.798  0.0052
##  PBS_48 - PBS_0    0.7988 0.236 27   3.386  0.0146
##  PBS_6 - PBS_0    -0.0814 0.236 27  -0.345  0.9943
##
## Results are given on the log2 (not the response) scale.
## P value adjustment: dunnettx method for 8 tests
```

**Analysis of Comparisons with Control (Figure S4)** Same as the above for Figure 1C, except using the data for Figure S4.

```
aS4_1FBP1 <- aov(log2(FBP1)~Treatment, data=dS4_1)
aS4_1GADPH <- aov(log2(GADPH)~Treatment, data=dS4_1)
aS4_1HK1 <- aov(log2(HK1)~Treatment, data=dS4_1)
aS4_1HK2 <- aov(log2(HK2)~Treatment, data=dS4_1)
aS4_1LDHA <- aov(log2(LDHA)~Treatment, data=dS4_1)
aS4_1LDHB <- aov(log2(LDHB)~Treatment, data=dS4_1)
plot(aS4_1FBP1,which=1:2)
```

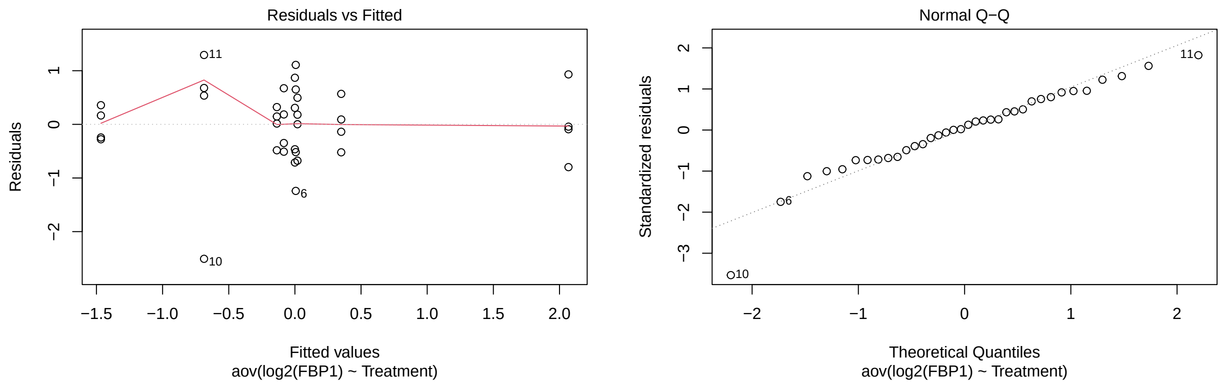

```r
plot(aS4_1GADPH,which=1:2)
```

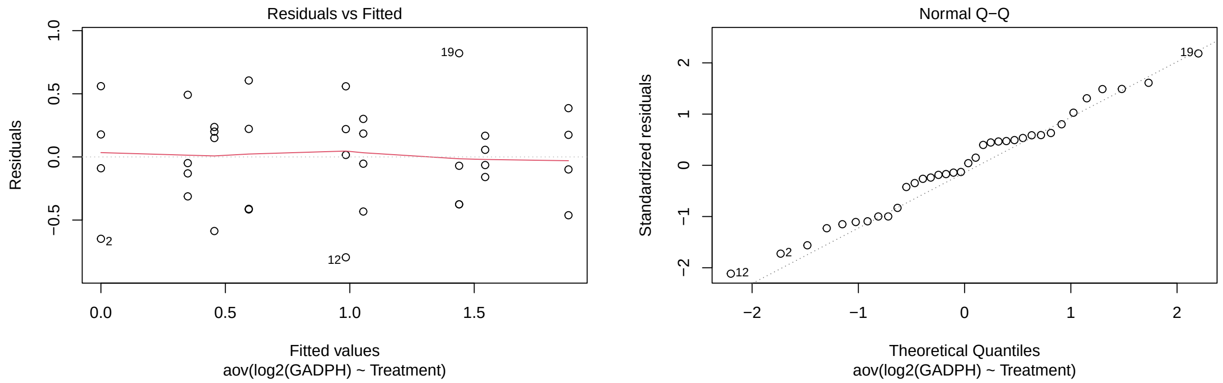

```r
plot(aS4_1HK1,which=1:2)
```

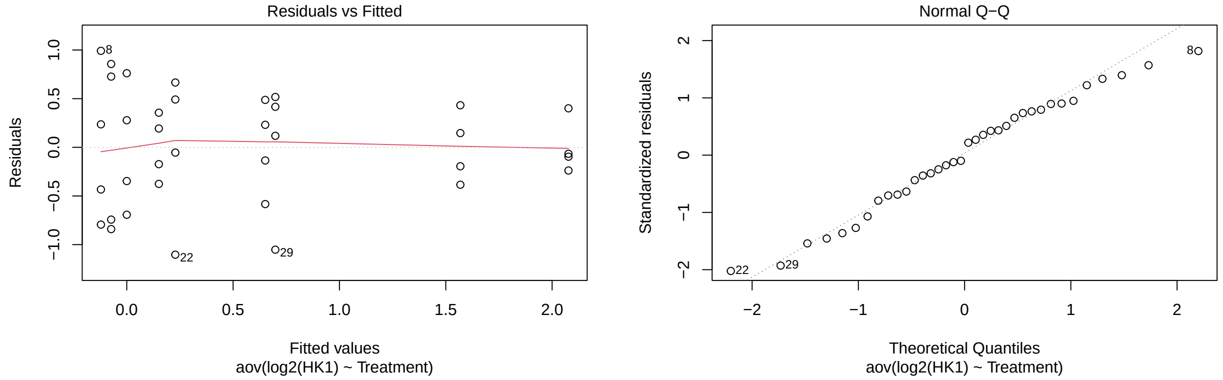

```r
plot(aS4_1HK2,which=1:2)
```

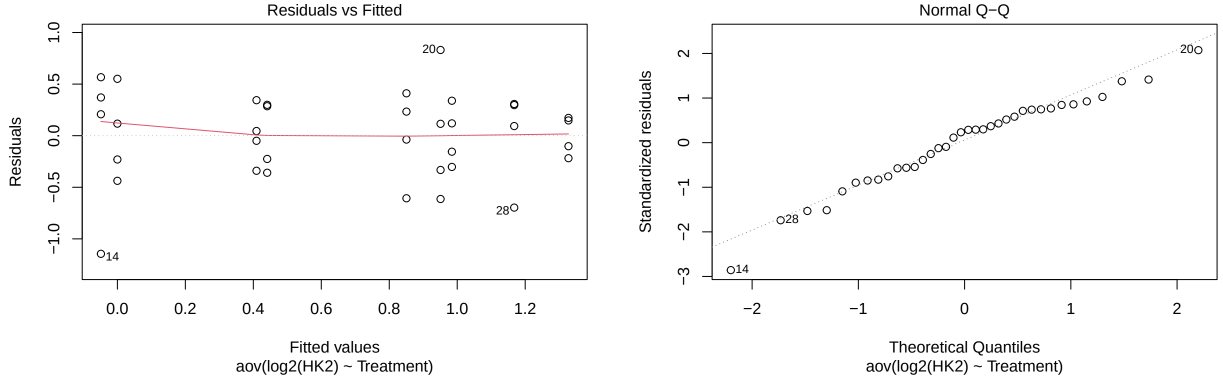

```r
plot(aS4_1LDHA,which=1:2)
```

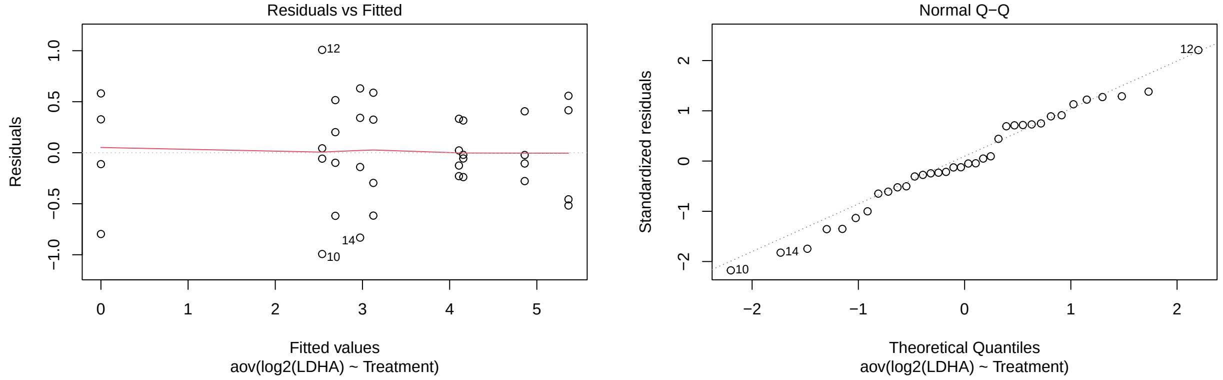

```r
plot(aS4_1LDHB,which=1:2)
```

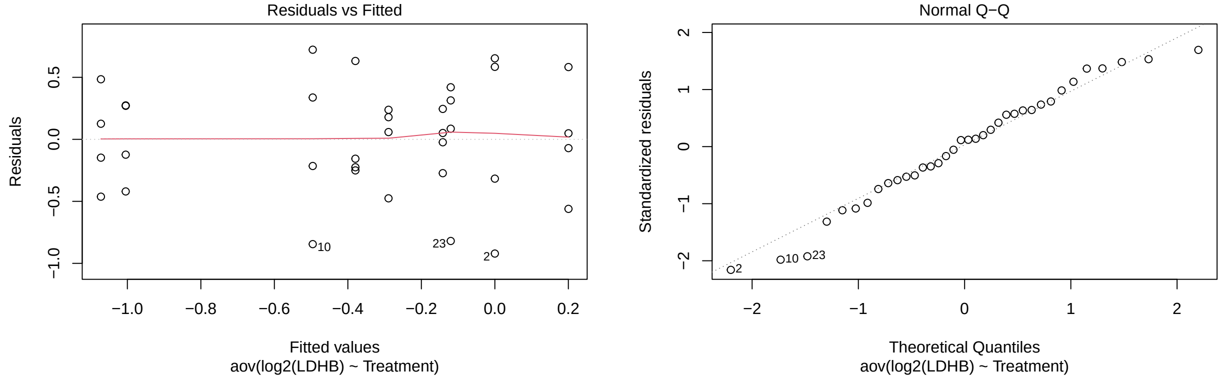

```r
emmeans(aS4_1FBP1,trt.vs.ctrlk ~ Treatment, ref=5)
```

```
## $emmeans
##  Treatment   emmean   SE df lower.CL upper.CL
##  FGF2_12   -0.13646 0.41 27   -0.977    0.704
```

```
##  FGF2_24   -1.46610 0.41 27   -2.307   -0.626
##  FGF2_48    0.01957 0.41 27   -0.821    0.860
##  FGF2_6    -0.68665 0.41 27   -1.527    0.154
##  PBS_0      0.00000 0.41 27   -0.841    0.841
##  PBS_12     0.35009 0.41 27   -0.490    1.191
##  PBS_24    -0.08360 0.41 27   -0.924    0.757
##  PBS_48     2.06771 0.41 27    1.227    2.908
##  PBS_6      0.00656 0.41 27   -0.834    0.847
##
## Results are given on the log2 (not the response) scale.
## Confidence level used: 0.95
##
## $contrasts
##  contrast        estimate    SE df t.ratio p.value
##  FGF2_12 - PBS_0 -0.13646 0.579 27  -0.236  0.9985
##  FGF2_24 - PBS_0 -1.46610 0.579 27  -2.531  0.1007
##  FGF2_48 - PBS_0  0.01957 0.579 27   0.034  1.0000
##  FGF2_6 - PBS_0  -0.68665 0.579 27  -1.185  0.7328
##  PBS_12 - PBS_0   0.35009 0.579 27   0.604  0.9629
##  PBS_24 - PBS_0  -0.08360 0.579 27  -0.144  0.9997
##  PBS_48 - PBS_0   2.06771 0.579 27   3.569  0.0093
##  PBS_6 - PBS_0    0.00656 0.579 27   0.011  1.0000
##
## Results are given on the log2 (not the response) scale.
## P value adjustment: dunnettx method for 8 tests
```

```
emmeans(aS4_1GADPH,trt.vs.ctrlk ~ Treatment, ref=5)
```

```
## $emmeans
##  Treatment emmean    SE df lower.CL upper.CL
##  FGF2_12    1.439 0.217 27   0.9940    1.885
##  FGF2_24    1.544 0.217 27   1.0986    1.989
##  FGF2_48    1.879 0.217 27   1.4333    2.324
##  FGF2_6     0.984 0.217 27   0.5388    1.429
##  PBS_0      0.000 0.217 27  -0.4454    0.445
##  PBS_12     0.456 0.217 27   0.0104    0.901
##  PBS_24     0.594 0.217 27   0.1488    1.040
##  PBS_48     1.054 0.217 27   0.6088    1.500
##  PBS_6      0.349 0.217 27  -0.0963    0.794
##
## Results are given on the log2 (not the response) scale.
## Confidence level used: 0.95
##
## $contrasts
##  contrast        estimate    SE df t.ratio p.value
##  FGF2_12 - PBS_0    1.439 0.307 27   4.689  0.0005
##  FGF2_24 - PBS_0    1.544 0.307 27   5.030  0.0002
##  FGF2_48 - PBS_0    1.879 0.307 27   6.120  <.0001
##  FGF2_6 - PBS_0     0.984 0.307 27   3.206  0.0224
##  PBS_12 - PBS_0     0.456 0.307 27   1.485  0.5522
##  PBS_24 - PBS_0     0.594 0.307 27   1.936  0.2994
##  PBS_48 - PBS_0     1.054 0.307 27   3.434  0.0129
##  PBS_6 - PBS_0      0.349 0.307 27   1.137  0.7597
##
```

```
## Results are given on the log2 (not the response) scale.
## P value adjustment: dunnettx method for 8 tests
```

```r
emmeans(aS4_1HK1,trt.vs.ctrlk ~ Treatment, ref=5)
```

```
## $emmeans
##  Treatment  emmean    SE df lower.CL upper.CL
##  FGF2_12   -0.0734 0.315 27 -0.71956    0.573
##  FGF2_24    2.0769 0.315 27  1.43070    2.723
##  FGF2_48    1.5686 0.315 27  0.92236    2.215
##  FGF2_6     0.6510 0.315 27  0.00484    1.297
##  PBS_0      0.0000 0.315 27 -0.64620    0.646
##  PBS_12     0.1508 0.315 27 -0.49537    0.797
##  PBS_24     0.2283 0.315 27 -0.41795    0.874
##  PBS_48     0.6986 0.315 27  0.05235    1.345
##  PBS_6     -0.1215 0.315 27 -0.76774    0.525
##
## Results are given on the log2 (not the response) scale.
## Confidence level used: 0.95
##
## $contrasts
##  contrast        estimate    SE df t.ratio p.value
##  FGF2_12 - PBS_0  -0.0734 0.445 27  -0.165  0.9996
##  FGF2_24 - PBS_0   2.0769 0.445 27   4.663  0.0005
##  FGF2_48 - PBS_0   1.5686 0.445 27   3.522  0.0104
##  FGF2_6 - PBS_0    0.6510 0.445 27   1.462  0.5664
##  PBS_12 - PBS_0    0.1508 0.445 27   0.339  0.9946
##  PBS_24 - PBS_0    0.2283 0.445 27   0.512  0.9783
##  PBS_48 - PBS_0    0.6986 0.445 27   1.568  0.5010
##  PBS_6 - PBS_0    -0.1215 0.445 27  -0.273  0.9975
##
## Results are given on the log2 (not the response) scale.
## P value adjustment: dunnettx method for 8 tests
```

```r
emmeans(aS4_1HK2,trt.vs.ctrlk ~ Treatment, ref=5)
```

```
## $emmeans
##  Treatment  emmean    SE df lower.CL upper.CL
##  FGF2_12    0.9510 0.231 27   0.4764    1.426
##  FGF2_24    1.1677 0.231 27   0.6932    1.642
##  FGF2_48    0.4093 0.231 27  -0.0652    0.884
##  FGF2_6     1.3274 0.231 27   0.8528    1.802
##  PBS_0      0.0000 0.231 27  -0.4746    0.475
##  PBS_12    -0.0485 0.231 27  -0.5230    0.426
##  PBS_24     0.9843 0.231 27   0.5098    1.459
##  PBS_48     0.4408 0.231 27  -0.0337    0.915
##  PBS_6      0.8506 0.231 27   0.3760    1.325
##
## Results are given on the log2 (not the response) scale.
## Confidence level used: 0.95
##
## $contrasts
##  contrast        estimate    SE df t.ratio p.value
```

```
##  FGF2_12 - PBS_0   0.9510 0.327 27   2.907  0.0448
##  FGF2_24 - PBS_0   1.1677 0.327 27   3.570  0.0093
##  FGF2_48 - PBS_0   0.4093 0.327 27   1.252  0.6945
##  FGF2_6 - PBS_0    1.3274 0.327 27   4.058  0.0027
##  PBS_12 - PBS_0   -0.0485 0.327 27  -0.148  0.9997
##  PBS_24 - PBS_0    0.9843 0.327 27   3.009  0.0355
##  PBS_48 - PBS_0    0.4408 0.327 27   1.348  0.6366
##  PBS_6 - PBS_0     0.8506 0.327 27   2.600  0.0872
##
## Results are given on the log2 (not the response) scale.
## P value adjustment: dunnettx method for 8 tests
```

```
emmeans(aS4_1LDHA,trt.vs.ctrlk ~ Treatment, ref=5)
```

```
## $emmeans
##  Treatment emmean    SE df lower.CL upper.CL
##  FGF2_12     3.12 0.263 27    2.584    3.665
##  FGF2_24     5.36 0.263 27    4.822    5.904
##  FGF2_48     4.86 0.263 27    4.320    5.402
##  FGF2_6      2.54 0.263 27    1.997    3.078
##  PBS_0       0.00 0.263 27   -0.541    0.541
##  PBS_12      2.97 0.263 27    2.432    3.514
##  PBS_24      4.16 0.263 27    3.616    4.697
##  PBS_48      4.11 0.263 27    3.566    4.647
##  PBS_6       2.69 0.263 27    2.148    3.230
##
## Results are given on the log2 (not the response) scale.
## Confidence level used: 0.95
##
## $contrasts
##  contrast        estimate    SE df t.ratio p.value
##  FGF2_12 - PBS_0     3.12 0.373 27   8.386  <.0001
##  FGF2_24 - PBS_0     5.36 0.373 27  14.393  <.0001
##  FGF2_48 - PBS_0     4.86 0.373 27  13.046  <.0001
##  FGF2_6 - PBS_0      2.54 0.373 27   6.811  <.0001
##  PBS_12 - PBS_0      2.97 0.373 27   7.979  <.0001
##  PBS_24 - PBS_0      4.16 0.373 27  11.155  <.0001
##  PBS_48 - PBS_0      4.11 0.373 27  11.022  <.0001
##  PBS_6 - PBS_0       2.69 0.373 27   7.217  <.0001
##
## Results are given on the log2 (not the response) scale.
## P value adjustment: dunnettx method for 8 tests
```

```
emmeans(aS4_1LDHB,trt.vs.ctrlk ~ Treatment, ref=5)
```

```
## $emmeans
##  Treatment emmean    SE df lower.CL upper.CL
##  FGF2_12    0.200 0.246 27   -0.305  0.70555
##  FGF2_24   -0.142 0.246 27   -0.647  0.36354
##  FGF2_48   -1.072 0.246 27   -1.577 -0.56672
##  FGF2_6    -0.496 0.246 27   -1.001  0.00966
##  PBS_0      0.000 0.246 27   -0.505  0.50523
##  PBS_12    -0.289 0.246 27   -0.795  0.21585
```

```
##  PBS_24    -0.120 0.246 27   -0.625  0.38516
##  PBS_48    -1.004 0.246 27   -1.510 -0.49925
##  PBS_6     -0.380 0.246 27   -0.885  0.12571
##
## Results are given on the log2 (not the response) scale.
## Confidence level used: 0.95
##
## $contrasts
##  contrast        estimate    SE df t.ratio p.value
##  FGF2_12 - PBS_0    0.200 0.348 27   0.575  0.9684
##  FGF2_24 - PBS_0   -0.142 0.348 27  -0.407  0.9900
##  FGF2_48 - PBS_0   -1.072 0.348 27  -3.078  0.0303
##  FGF2_6 - PBS_0    -0.496 0.348 27  -1.423  0.5902
##  PBS_12 - PBS_0    -0.289 0.348 27  -0.831  0.9002
##  PBS_24 - PBS_0    -0.120 0.348 27  -0.345  0.9943
##  PBS_48 - PBS_0    -1.004 0.348 27  -2.885  0.0471
##  PBS_6 - PBS_0     -0.380 0.348 27  -1.090  0.7849
##
## Results are given on the log2 (not the response) scale.
## P value adjustment: dunnettx method for 8 tests
```

**Analysis: Treatment Comparisons**

Here we compare the two treatments (PBS vs. FGF2) at each time point greater than 0, for the genes in both Figures 1C and 4S. Since here we will omit the "PBS at Time 0", we have two crossed factors and can fit a two-way ANOVA.

```
d1C_2 <- d1C %>%
  rename(Time=`Time.(h)`) %>%
  filter(Time != 0) %>%
  mutate(Time=as.factor(Time))
glimpse(d1C_2)
```

```
## Rows: 32
## Columns: 9
## $ Chemical <chr> "PBS", "PBS", "PBS", "PBS", "FGF2", "FGF2", "FGF2", "FGF2", "~
## $ Sample   <dbl> 1, 2, 3, 4, 1, 2, 3, 4, 1, 2, 3, 4, 1, 2, 3, 4, 1, 2, 3, 4, 1~
## $ Time     <fct> 6, 6, 6, 6, 6, 6, 6, 6, 12, 12, 12, 12, 12, 12, 12, 12, 24, 2~
## $ SOX2     <dbl> 0.9540158, 1.0775179, 1.6006179, 3.6215695, 5.6372863, 7.0325~
## $ SIX6     <dbl> 13.70090, 13.50870, 14.72723, 11.63435, 32.24528, 24.13720, 2~
## $ PAX6     <dbl> 0.7258499, 0.7009500, 0.9946392, 1.3939510, 1.2935352, 0.7592~
## $ RPE65    <dbl> 0.3257529, 0.2030872, 0.1613492, 0.9810401, 0.4461522, 0.7676~
## $ TYR      <dbl> 1.1734015, 0.7431016, 0.7364267, 1.0064826, 1.2448899, 1.3597~
## $ OTX2     <dbl> 0.9470746, 0.7333406, 1.3031499, 0.8817553, 0.7246953, 0.6984~
```

```
dS4_2 <- dS4 %>%
  rename(Time=`Time.(h)`) %>%
  filter(Time != 0) %>%
  mutate(Time=as.factor(Time))
glimpse(dS4_2)
```

```
## Rows: 32
```

```
## Columns: 9
## $ Chemical <chr> "PBS", "PBS", "PBS", "PBS", "FGF2", "FGF2", "FGF2", "FGF2", "~
## $ Sample   <dbl> 1, 2, 3, 4, 1, 2, 3, 4, 1, 2, 3, 4, 1, 2, 3, 4, 1, 2, 3, 4, 1~
## $ Time     <fct> 6, 6, 6, 6, 6, 6, 6, 6, 12, 12, 12, 12, 12, 12, 12, 12, 24, 2~
## $ HK1      <dbl> 0.5300377, 1.0827360, 0.6807349, 1.8274311, 1.4297081, 1.0479~
## $ HK2      <dbl> 1.1842694, 2.1198080, 2.3972541, 1.7568371, 2.1572973, 2.3383~
## $ GADPH    <dbl> 1.7912290, 1.0256197, 1.2311533, 1.1636124, 1.9997260, 2.3045~
## $ LDHA     <dbl> 6.022702, 4.200064, 7.415969, 9.220020, 5.578625, 2.918004, 5~
## $ LDHB     <dbl> 0.6576684, 0.6897773, 0.6461395, 1.1911589, 0.6110950, 0.3948~
## $ FBP1     <dbl> 0.7015782, 0.4250852, 1.5768726, 2.1654884, 0.9939663, 0.1093~
```

**Treatment Comparisons (Figure 1C)** In the analysis below we compare FGF2 to PBS at Time 6, 12, 24, and 48. Then, in order to account for multiple comparisons for each response, we adjust these four p-values using the Benjamini and Hochberg False Discovery Rate.

```r
a1C_2SOX2 <- aov(log2(SOX2)~Chemical*Time, data=d1C_2)
a1C_2SIX6 <- aov(log2(SIX6)~Chemical*Time, data=d1C_2)
a1C_2PAX6 <- aov(log2(PAX6)~Chemical*Time, data=d1C_2)
a1C_2RPE65 <- aov(log2(RPE65)~Chemical*Time, data=d1C_2)
a1C_2TYR <- aov(log2(TYR)~Chemical*Time, data=d1C_2)
a1C_2OTX2 <- aov(log2(OTX2)~Chemical*Time, data=d1C_2)

plot(a1C_2SOX2,which=1:2)
```

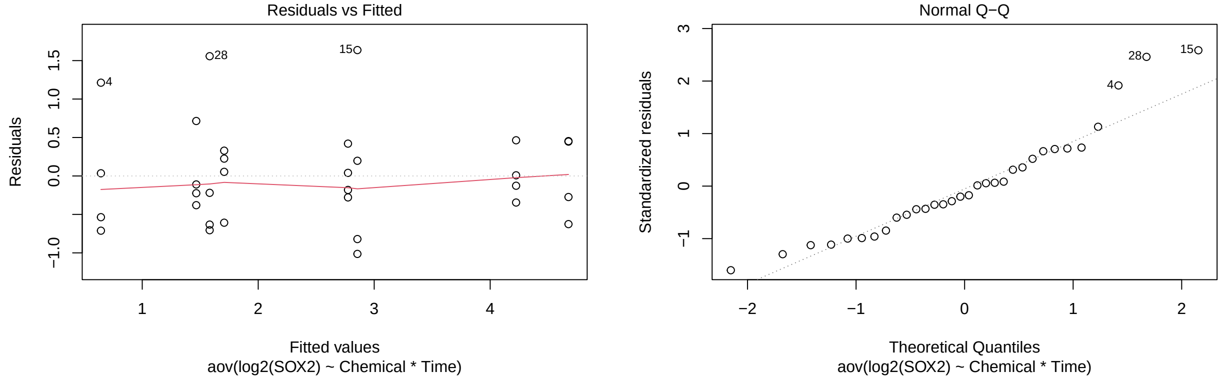

```r
plot(a1C_2SIX6,which=1:2)
```

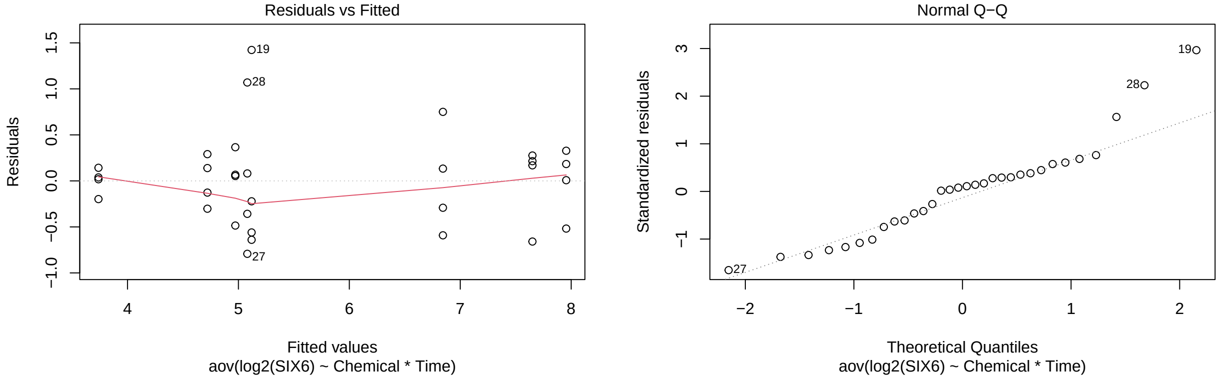

```r
plot(a1C_2PAX6,which=1:2)
```

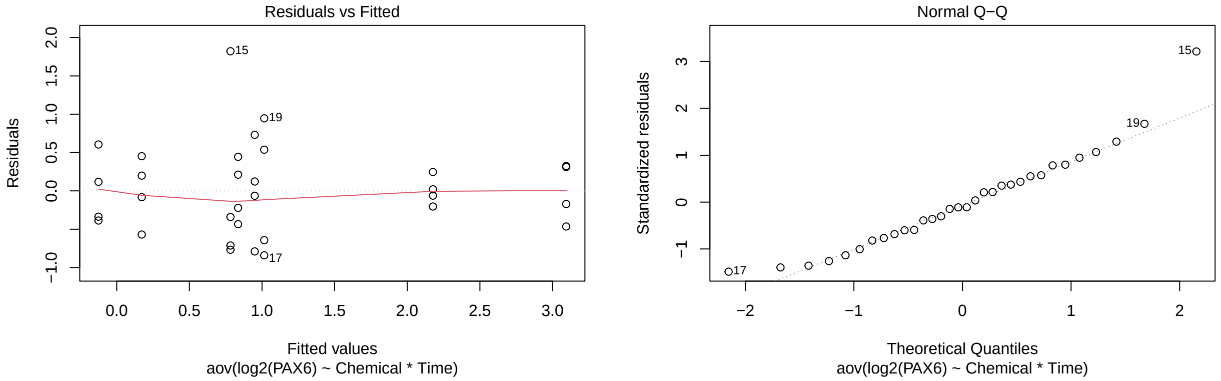

```r
plot(a1C_2RPE65,which=1:2)
```

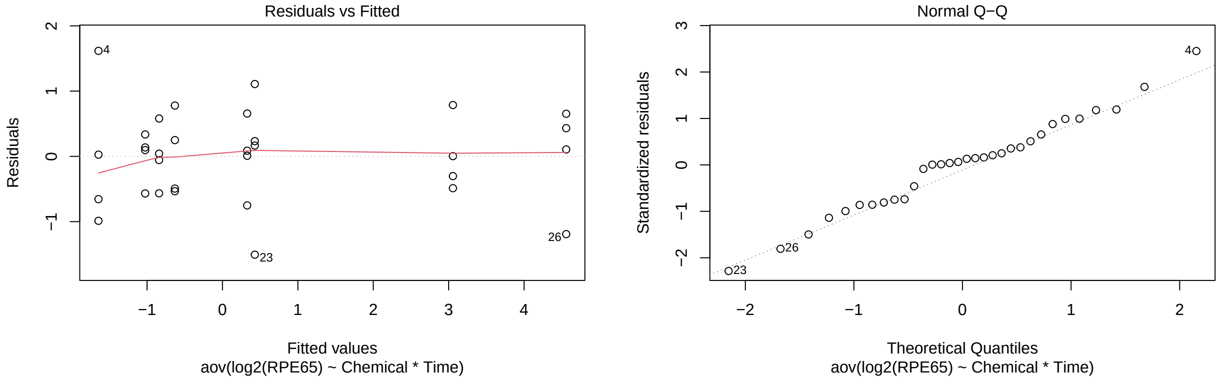

```r
plot(a1C_2TYR,which=1:2)
```

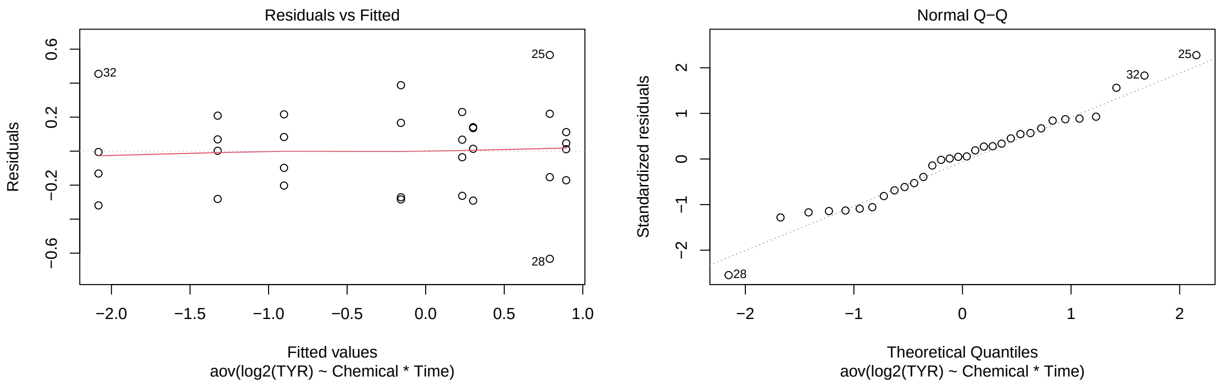

```r
plot(a1C_2OTX2,which=1:2)
```

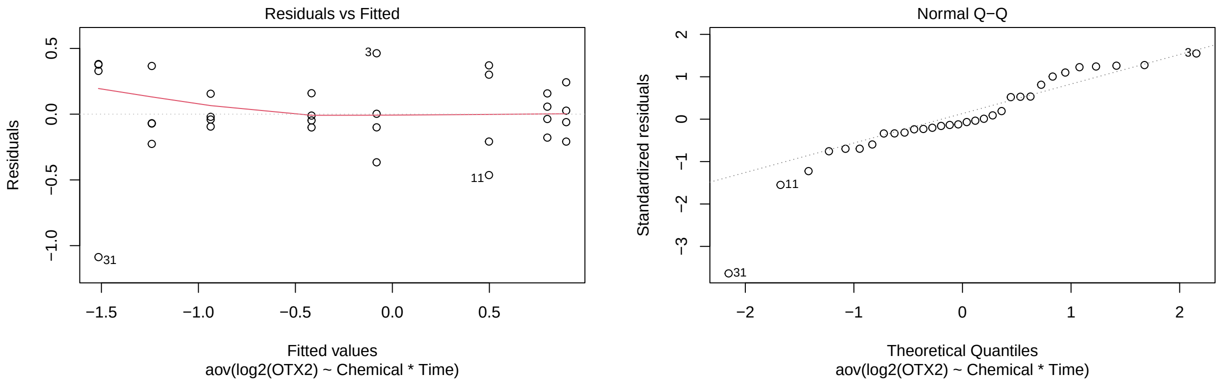

```r
a1C_2tSOX2 <- test(emmeans(a1C_2SOX2, pairwise~Chemical|Time))
a1C_2tSOX2
```

```
## $emmeans
## Time = 6:
##  Chemical emmean    SE df t.ratio p.value
##  FGF2      2.774 0.365 24   7.592  <.0001
##  PBS       0.644 0.365 24   1.762  0.0908
##
## Time = 12:
##  Chemical emmean    SE df t.ratio p.value
##  FGF2      2.855 0.365 24   7.814  <.0001
##  PBS       1.466 0.365 24   4.011  0.0005
##
## Time = 24:
##  Chemical emmean    SE df t.ratio p.value
##  FGF2      4.223 0.365 24  11.558  <.0001
##  PBS       1.706 0.365 24   4.670  0.0001
##
## Time = 48:
##  Chemical emmean    SE df t.ratio p.value
##  FGF2      4.675 0.365 24  12.794  <.0001
```

```
##  PBS       1.581 0.365 24   4.328  0.0002
##
## Results are given on the log2 (not the response) scale.
##
## $contrasts
## Time = 6:
##  contrast   estimate    SE df t.ratio p.value
##  FGF2 - PBS     2.13 0.517 24   4.122  0.0004
##
## Time = 12:
##  contrast   estimate    SE df t.ratio p.value
##  FGF2 - PBS     1.39 0.517 24   2.689  0.0128
##
## Time = 24:
##  contrast   estimate    SE df t.ratio p.value
##  FGF2 - PBS     2.52 0.517 24   4.871  0.0001
##
## Time = 48:
##  contrast   estimate    SE df t.ratio p.value
##  FGF2 - PBS     3.09 0.517 24   5.987  <.0001
##
## Results are given on the log2 (not the response) scale.
```

```
p.adjust(a1C_2tSOX2$contrasts$p.value, method="fdr")
```

```
## [1] 5.156404e-04 1.283618e-02 1.154001e-04 1.408442e-05
```

```
a1C_2tSIX6 <- test(emmeans(a1C_2SIX6, pairwise~Chemical|Time))
a1C_2tSIX6
```

```
## $emmeans
## Time = 6:
##  Chemical emmean    SE df t.ratio p.value
##  FGF2       4.72 0.277 24  17.044  <.0001
##  PBS        3.74 0.277 24  13.498  <.0001
##
## Time = 12:
##  Chemical emmean    SE df t.ratio p.value
##  FGF2       6.84 0.277 24  24.712  <.0001
##  PBS        4.97 0.277 24  17.953  <.0001
##
## Time = 24:
##  Chemical emmean    SE df t.ratio p.value
##  FGF2       7.96 0.277 24  28.725  <.0001
##  PBS        5.12 0.277 24  18.483  <.0001
##
## Time = 48:
##  Chemical emmean    SE df t.ratio p.value
##  FGF2       7.65 0.277 24  27.626  <.0001
##  PBS        5.08 0.277 24  18.344  <.0001
##
## Results are given on the log2 (not the response) scale.
##
```

```
## $contrasts
## Time = 6:
##  contrast   estimate    SE df t.ratio p.value
##  FGF2 - PBS    0.982 0.392 24   2.508  0.0193
##
## Time = 12:
##  contrast   estimate    SE df t.ratio p.value
##  FGF2 - PBS    1.872 0.392 24   4.779  0.0001
##
## Time = 24:
##  contrast   estimate    SE df t.ratio p.value
##  FGF2 - PBS    2.837 0.392 24   7.242  <.0001
##
## Time = 48:
##  contrast   estimate    SE df t.ratio p.value
##  FGF2 - PBS    2.571 0.392 24   6.563  <.0001
##
## Results are given on the log2 (not the response) scale.
```

```r
p.adjust(a1C_2tSIX6$contrasts$p.value, method="fdr")
```

```
## [1] 1.931443e-02 9.715567e-05 7.011665e-07 1.734455e-06
```

```r
a1C_2tSIX6$contrasts$p.value
```

```
## [1] 1.931443e-02 7.286675e-05 1.752916e-07 8.672275e-07
```

```r
a1C_2tPAX6 <- test(emmeans(a1C_2PAX6, pairwise~Chemical|Time))
a1C_2tPAX6
```

```
## $emmeans
## Time = 6:
##  Chemical emmean    SE df t.ratio p.value
##  FGF2      0.172 0.327 24   0.526  0.6034
##  PBS      -0.126 0.327 24  -0.385  0.7037
##
## Time = 12:
##  Chemical emmean    SE df t.ratio p.value
##  FGF2      0.783 0.327 24   2.394  0.0249
##  PBS       0.835 0.327 24   2.555  0.0174
##
## Time = 24:
##  Chemical emmean    SE df t.ratio p.value
##  FGF2      3.094 0.327 24   9.461  <.0001
##  PBS       1.015 0.327 24   3.105  0.0048
##
## Time = 48:
##  Chemical emmean    SE df t.ratio p.value
##  FGF2      2.177 0.327 24   6.656  <.0001
##  PBS       0.950 0.327 24   2.904  0.0078
##
## Results are given on the log2 (not the response) scale.
```

```
##
## $contrasts
## Time = 6:
##  contrast   estimate    SE df t.ratio p.value
##  FGF2 - PBS   0.2980 0.463 24   0.644  0.5254
##
## Time = 12:
##  contrast   estimate    SE df t.ratio p.value
##  FGF2 - PBS  -0.0526 0.463 24  -0.114  0.9103
##
## Time = 24:
##  contrast   estimate    SE df t.ratio p.value
##  FGF2 - PBS   2.0788 0.463 24   4.495  0.0002
##
## Time = 48:
##  contrast   estimate    SE df t.ratio p.value
##  FGF2 - PBS   1.2269 0.463 24   2.653  0.0139
##
## Results are given on the log2 (not the response) scale.
```

```
p.adjust(a1C_2tPAX6$contrasts$p.value, method="fdr")
```

```
## [1] 0.7005835130 0.9103188068 0.0006004952 0.0278668746
```

```
a1C_2tRPE65 <- test(emmeans(a1C_2RPE65, pairwise~Chemical|Time))
a1C_2tRPE65
```

```
## $emmeans
## Time = 6:
##  Chemical emmean    SE df t.ratio p.value
##  FGF2     -0.631 0.381 24  -1.656  0.1107
##  PBS      -1.644 0.381 24  -4.319  0.0002
##
## Time = 12:
##  Chemical emmean    SE df t.ratio p.value
##  FGF2     -1.025 0.381 24  -2.692  0.0127
##  PBS      -0.840 0.381 24  -2.207  0.0371
##
## Time = 24:
##  Chemical emmean    SE df t.ratio p.value
##  FGF2      0.429 0.381 24   1.128  0.2706
##  PBS       3.057 0.381 24   8.029  <.0001
##
## Time = 48:
##  Chemical emmean    SE df t.ratio p.value
##  FGF2      0.328 0.381 24   0.861  0.3977
##  PBS       4.559 0.381 24  11.974  <.0001
##
## Results are given on the log2 (not the response) scale.
##
## $contrasts
## Time = 6:
##  contrast   estimate    SE df t.ratio p.value
```

```
##   FGF2 - PBS    1.014 0.538 24   1.883  0.0719
##
## Time = 12:
##   contrast   estimate    SE df t.ratio p.value
##   FGF2 - PBS   -0.185 0.538 24  -0.343  0.7347
##
## Time = 24:
##   contrast   estimate    SE df t.ratio p.value
##   FGF2 - PBS   -2.627 0.538 24  -4.880  0.0001
##
## Time = 48:
##   contrast   estimate    SE df t.ratio p.value
##   FGF2 - PBS   -4.231 0.538 24  -7.858  <.0001
##
## Results are given on the log2 (not the response) scale.
```

```r
p.adjust(a1C_2tRPE65$contrasts$p.value, method="fdr")
```

```
## [1] 9.584095e-02 7.347186e-01 1.127489e-04 1.730255e-07
```

```r
a1C_2tTYR <- test(emmeans(a1C_2TYR, pairwise~Chemical|Time))
a1C_2tTYR
```

```
## $emmeans
## Time = 6:
##   Chemical emmean    SE df t.ratio p.value
##   FGF2      0.302 0.144 24   2.104  0.0460
##   PBS      -0.157 0.144 24  -1.097  0.2837
##
## Time = 12:
##   Chemical emmean    SE df t.ratio p.value
##   FGF2     -0.902 0.144 24  -6.280  <.0001
##   PBS       0.233 0.144 24   1.620  0.1183
##
## Time = 24:
##   Chemical emmean    SE df t.ratio p.value
##   FGF2     -1.324 0.144 24  -9.223  <.0001
##   PBS       0.894 0.144 24   6.231  <.0001
##
## Time = 48:
##   Chemical emmean    SE df t.ratio p.value
##   FGF2     -2.083 0.144 24 -14.509  <.0001
##   PBS       0.790 0.144 24   5.503  <.0001
##
## Results are given on the log2 (not the response) scale.
##
## $contrasts
## Time = 6:
##   contrast   estimate    SE df t.ratio p.value
##   FGF2 - PBS     0.46 0.203 24   2.263  0.0329
##
## Time = 12:
##   contrast   estimate    SE df t.ratio p.value
```

```
##  FGF2 - PBS    -1.13 0.203 24  -5.586  <.0001
##
## Time = 24:
##  contrast   estimate    SE df t.ratio p.value
##  FGF2 - PBS    -2.22 0.203 24 -10.927  <.0001
##
## Time = 48:
##  contrast   estimate    SE df t.ratio p.value
##  FGF2 - PBS    -2.87 0.203 24 -14.150  <.0001
##
## Results are given on the log2 (not the response) scale.
```

```
p.adjust(a1C_2tTYR$contrasts$p.value, method="fdr")
```

```
## [1] 3.293503e-02 1.267237e-05 1.693320e-10 1.535537e-12
```

```
a1C_2tOTX2 <- test(emmeans(a1C_2OTX2, pairwise~Chemical|Time))
a1C_2tOTX2
```

```
## $emmeans
## Time = 6:
##  Chemical  emmean    SE df t.ratio p.value
##  FGF2     -0.4168 0.173 24  -2.416  0.0237
##  PBS      -0.0814 0.173 24  -0.471  0.6416
##
## Time = 12:
##  Chemical  emmean    SE df t.ratio p.value
##  FGF2     -0.9368 0.173 24  -5.429  <.0001
##  PBS       0.4980 0.173 24   2.886  0.0081
##
## Time = 24:
##  Chemical  emmean    SE df t.ratio p.value
##  FGF2     -1.2404 0.173 24  -7.188  <.0001
##  PBS       0.8960 0.173 24   5.192  <.0001
##
## Time = 48:
##  Chemical  emmean    SE df t.ratio p.value
##  FGF2     -1.5152 0.173 24  -8.781  <.0001
##  PBS       0.7988 0.173 24   4.629  0.0001
##
## Results are given on the log2 (not the response) scale.
##
## $contrasts
## Time = 6:
##  contrast   estimate    SE df t.ratio p.value
##  FGF2 - PBS   -0.335 0.244 24  -1.375  0.1819
##
## Time = 12:
##  contrast   estimate    SE df t.ratio p.value
##  FGF2 - PBS   -1.435 0.244 24  -5.879  <.0001
##
## Time = 24:
##  contrast   estimate    SE df t.ratio p.value
```

```
##  FGF2 - PBS   -2.136 0.244 24  -8.754  <.0001
##
## Time = 48:
##  contrast   estimate    SE df t.ratio p.value
##  FGF2 - PBS   -2.314 0.244 24  -9.482  <.0001
##
## Results are given on the log2 (not the response) scale.
```

```
p.adjust(a1C_2tOTX2$contrasts$p.value, method="fdr")
```

```
## [1] 1.819386e-01 6.118899e-06 1.235680e-08 5.499195e-09
```

**Treatment Comparisons (Figure S4)** Same idea as for Figure 1C.

```
aS4_2FBP1 <- aov(log2(FBP1)~Chemical*Time, data=dS4_2)
aS4_2GADPH <- aov(log2(GADPH)~Chemical*Time, data=dS4_2)
aS4_2HK1 <- aov(log2(HK1)~Chemical*Time, data=dS4_2)
aS4_2HK2 <- aov(log2(HK2)~Chemical*Time, data=dS4_2)
aS4_2LDHA <- aov(log2(LDHA)~Chemical*Time, data=dS4_2)
aS4_2LDHB <- aov(log2(LDHB)~Chemical*Time, data=dS4_2)

plot(aS4_2FBP1,which=1:2)
```

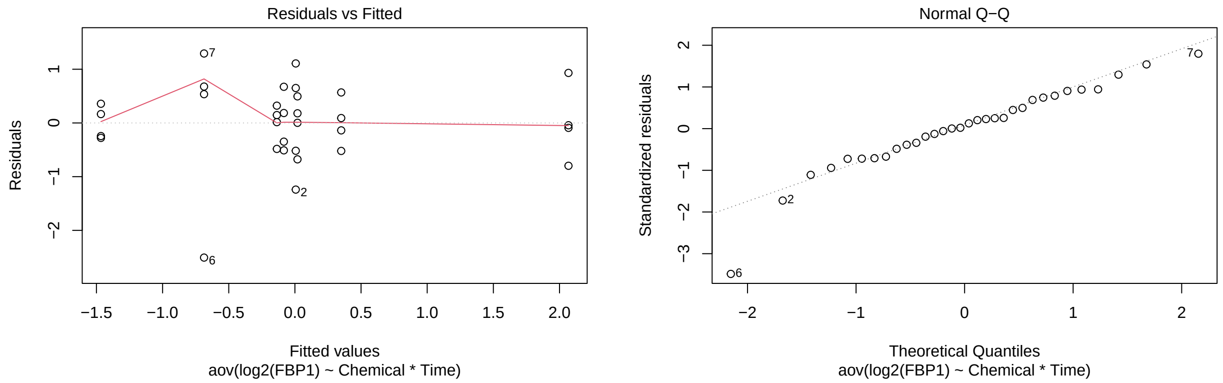

```
plot(aS4_2GADPH,which=1:2)
```

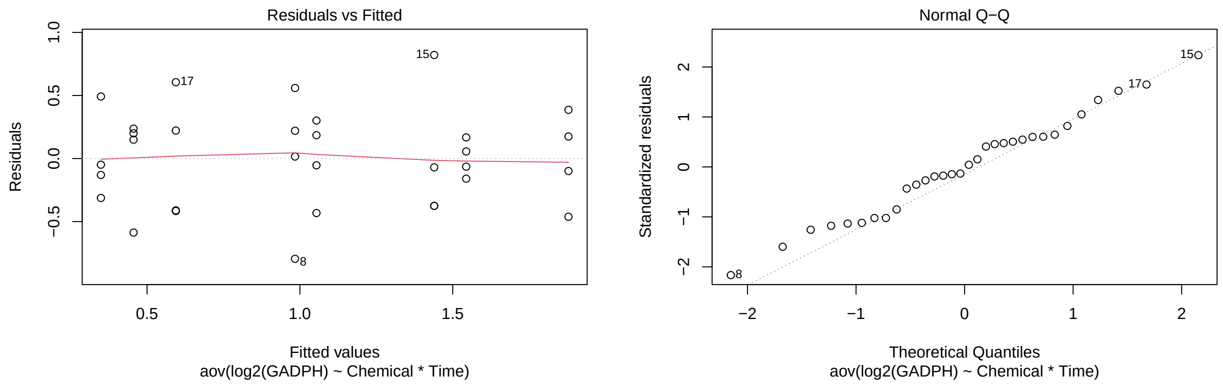

```
plot(aS4_2HK1,which=1:2)
```

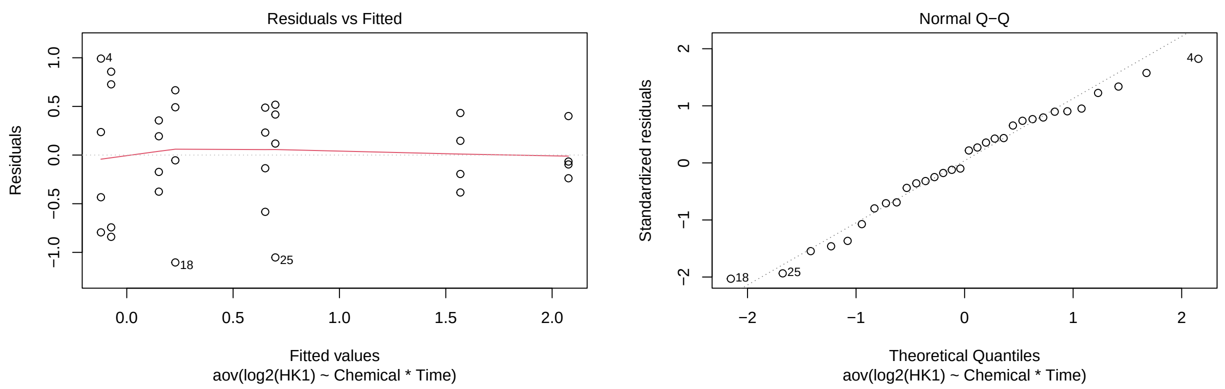

```
plot(aS4_2HK2,which=1:2)
```

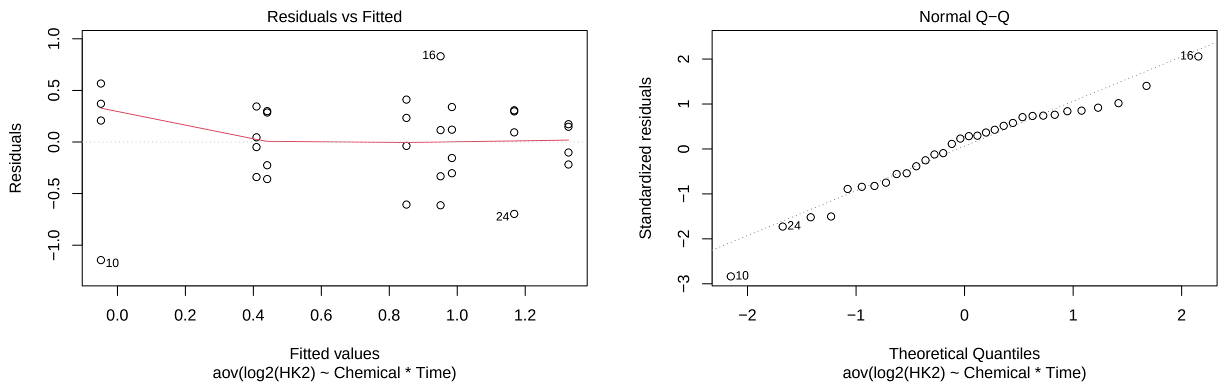

```
plot(aS4_2LDHA,which=1:2)
```

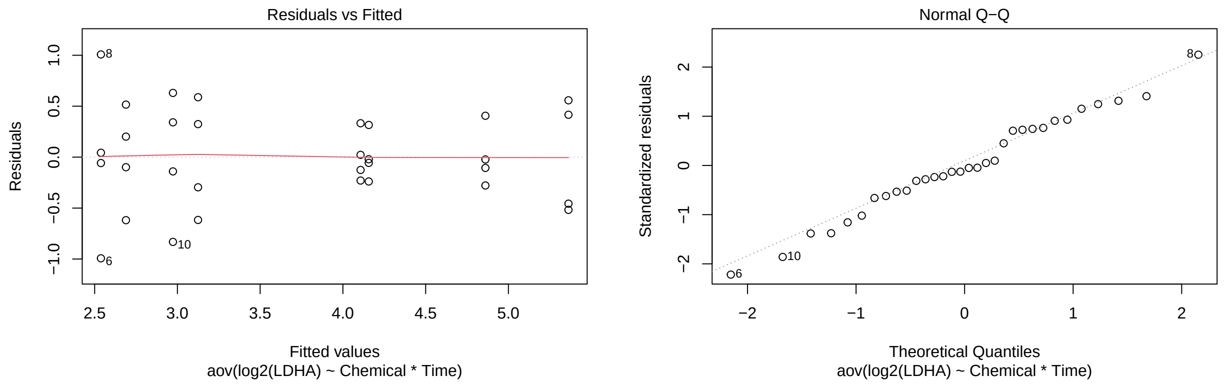

```
plot(aS4_2LDHB,which=1:2)
```

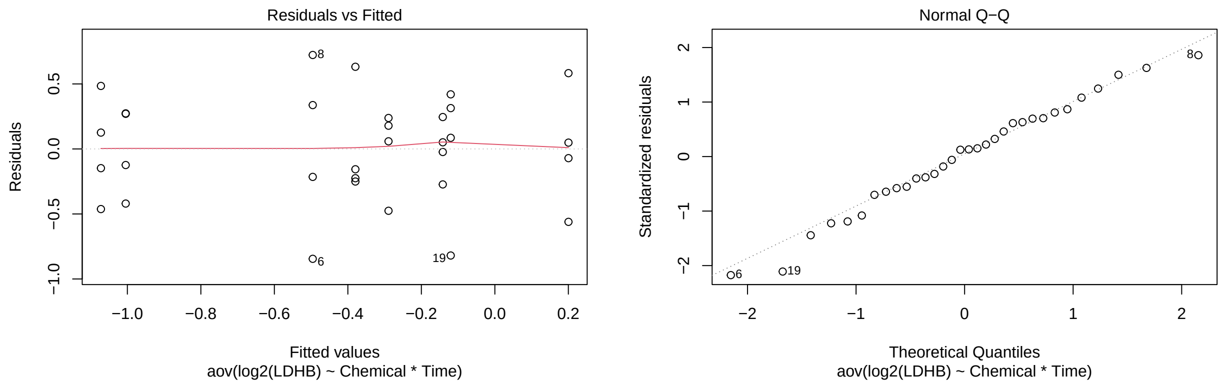

```r
aS4_2tFBP1 <- test(emmeans(aS4_2FBP1, pairwise~Chemical|Time))
aS4_2tFBP1
```

```
## $emmeans
## Time = 6:
##  Chemical   emmean    SE df t.ratio p.value
##  FGF2     -0.68665 0.415 24  -1.654  0.1112
##  PBS       0.00656 0.415 24   0.016  0.9875
##
## Time = 12:
##  Chemical   emmean    SE df t.ratio p.value
##  FGF2     -0.13646 0.415 24  -0.329  0.7453
##  PBS       0.35009 0.415 24   0.843  0.4075
##
## Time = 24:
##  Chemical   emmean    SE df t.ratio p.value
##  FGF2     -1.46610 0.415 24  -3.531  0.0017
##  PBS      -0.08360 0.415 24  -0.201  0.8421
##
## Time = 48:
##  Chemical   emmean    SE df t.ratio p.value
##  FGF2      0.01957 0.415 24   0.047  0.9628
##  PBS       2.06771 0.415 24   4.980  <.0001
##
## Results are given on the log2 (not the response) scale.
##
## $contrasts
## Time = 6:
##  contrast   estimate    SE df t.ratio p.value
##  FGF2 - PBS   -0.693 0.587 24  -1.180  0.2494
##
## Time = 12:
##  contrast   estimate    SE df t.ratio p.value
##  FGF2 - PBS   -0.487 0.587 24  -0.829  0.4155
##
## Time = 24:
##  contrast   estimate    SE df t.ratio p.value
##  FGF2 - PBS   -1.383 0.587 24  -2.354  0.0271
##
```

```
## Time = 48:
##  contrast   estimate    SE df t.ratio p.value
##  FGF2 - PBS   -2.048 0.587 24  -3.488  0.0019
##
## Results are given on the log2 (not the response) scale.
```

```r
p.adjust(aS4_2tFBP1$contrasts$p.value, method="fdr")
```

```
## [1] 0.332511725 0.415528348 0.054163497 0.007596631
```

```r
aS4_2tGADPH <- test(emmeans(aS4_2GADPH, pairwise~Chemical|Time))
aS4_2tGADPH
```

```
## $emmeans
## Time = 6:
##  Chemical emmean    SE df t.ratio p.value
##  FGF2      0.984 0.212 24   4.642  0.0001
##  PBS       0.349 0.212 24   1.646  0.1127
##
## Time = 12:
##  Chemical emmean    SE df t.ratio p.value
##  FGF2      1.439 0.212 24   6.790  <.0001
##  PBS       0.456 0.212 24   2.150  0.0419
##
## Time = 24:
##  Chemical emmean    SE df t.ratio p.value
##  FGF2      1.544 0.212 24   7.283  <.0001
##  PBS       0.594 0.212 24   2.803  0.0099
##
## Time = 48:
##  Chemical emmean    SE df t.ratio p.value
##  FGF2      1.879 0.212 24   8.862  <.0001
##  PBS       1.054 0.212 24   4.973  <.0001
##
## Results are given on the log2 (not the response) scale.
##
## $contrasts
## Time = 6:
##  contrast   estimate  SE df t.ratio p.value
##  FGF2 - PBS    0.635 0.3 24   2.118  0.0447
##
## Time = 12:
##  contrast   estimate  SE df t.ratio p.value
##  FGF2 - PBS    0.984 0.3 24   3.281  0.0032
##
## Time = 24:
##  contrast   estimate  SE df t.ratio p.value
##  FGF2 - PBS    0.950 0.3 24   3.168  0.0041
##
## Time = 48:
##  contrast   estimate  SE df t.ratio p.value
##  FGF2 - PBS    0.824 0.3 24   2.750  0.0111
##
## Results are given on the log2 (not the response) scale.
```

```r
p.adjust(aS4_2tGADPH$contrasts$p.value, method="fdr")
```

```
## [1] 0.044678173 0.008292999 0.008292999 0.014857888
```

```r
aS4_2tHK1 <- test(emmeans(aS4_2HK1, pairwise~Chemical|Time))
aS4_2tHK1
```

```
## $emmeans
## Time = 6:
##  Chemical  emmean    SE df t.ratio p.value
##  FGF2      0.6510 0.314 24   2.075  0.0489
##  PBS      -0.1215 0.314 24  -0.387  0.7019
##
## Time = 12:
##  Chemical  emmean    SE df t.ratio p.value
##  FGF2     -0.0734 0.314 24  -0.234  0.8172
##  PBS       0.1508 0.314 24   0.481  0.6351
##
## Time = 24:
##  Chemical  emmean    SE df t.ratio p.value
##  FGF2      2.0769 0.314 24   6.618  <.0001
##  PBS       0.2283 0.314 24   0.727  0.4741
##
## Time = 48:
##  Chemical  emmean    SE df t.ratio p.value
##  FGF2      1.5686 0.314 24   4.998  <.0001
##  PBS       0.6986 0.314 24   2.226  0.0357
##
## Results are given on the log2 (not the response) scale.
##
## $contrasts
## Time = 6:
##  contrast   estimate    SE df t.ratio p.value
##  FGF2 - PBS    0.773 0.444 24   1.741  0.0945
##
## Time = 12:
##  contrast   estimate    SE df t.ratio p.value
##  FGF2 - PBS   -0.224 0.444 24  -0.505  0.6181
##
## Time = 24:
##  contrast   estimate    SE df t.ratio p.value
##  FGF2 - PBS    1.849 0.444 24   4.165  0.0003
##
## Time = 48:
##  contrast   estimate    SE df t.ratio p.value
##  FGF2 - PBS    0.870 0.444 24   1.960  0.0617
##
## Results are given on the log2 (not the response) scale.
```

```r
p.adjust(aS4_2tHK1$contrasts$p.value, method="fdr")
```

```
## [1] 0.126040725 0.618068425 0.001386587 0.123336234
```

```r
aS4_2tHK2 <- test(emmeans(aS4_2HK2, pairwise~Chemical|Time))
aS4_2tHK2
```

```
## $emmeans
## Time = 6:
##  Chemical  emmean    SE df t.ratio p.value
##  FGF2      1.3274 0.233 24   5.695  <.0001
##  PBS       0.8506 0.233 24   3.649  0.0013
##
## Time = 12:
##  Chemical  emmean    SE df t.ratio p.value
##  FGF2      0.9510 0.233 24   4.080  0.0004
##  PBS      -0.0485 0.233 24  -0.208  0.8371
##
## Time = 24:
##  Chemical  emmean    SE df t.ratio p.value
##  FGF2      1.1677 0.233 24   5.010  <.0001
##  PBS       0.9843 0.233 24   4.223  0.0003
##
## Time = 48:
##  Chemical  emmean    SE df t.ratio p.value
##  FGF2      0.4093 0.233 24   1.756  0.0918
##  PBS       0.4408 0.233 24   1.891  0.0707
##
## Results are given on the log2 (not the response) scale.
##
## $contrasts
## Time = 6:
##  contrast   estimate   SE df t.ratio p.value
##  FGF2 - PBS   0.4768 0.33 24   1.447  0.1610
##
## Time = 12:
##  contrast   estimate   SE df t.ratio p.value
##  FGF2 - PBS   0.9994 0.33 24   3.032  0.0057
##
## Time = 24:
##  contrast   estimate   SE df t.ratio p.value
##  FGF2 - PBS   0.1834 0.33 24   0.556  0.5831
##
## Time = 48:
##  contrast   estimate   SE df t.ratio p.value
##  FGF2 - PBS  -0.0315 0.33 24  -0.096  0.9247
##
## Results are given on the log2 (not the response) scale.
```

```r
p.adjust(aS4_2tHK2$contrasts$p.value, method="fdr")
```

```
## [1] 0.32192564 0.02299921 0.77751835 0.92469491
```

```r
aS4_2tLDHA <- test(emmeans(aS4_2LDHA, pairwise~Chemical|Time))
aS4_2tLDHA
```

```
## $emmeans
## Time = 6:
##  Chemical emmean    SE df t.ratio p.value
##  FGF2       2.54 0.258 24   9.825  <.0001
##  PBS        2.69 0.258 24  10.411  <.0001
##
## Time = 12:
##  Chemical emmean    SE df t.ratio p.value
##  FGF2       3.12 0.258 24  12.097  <.0001
##  PBS        2.97 0.258 24  11.510  <.0001
##
## Time = 24:
##  Chemical emmean    SE df t.ratio p.value
##  FGF2       5.36 0.258 24  20.763  <.0001
##  PBS        4.16 0.258 24  16.091  <.0001
##
## Time = 48:
##  Chemical emmean    SE df t.ratio p.value
##  FGF2       4.86 0.258 24  18.819  <.0001
##  PBS        4.11 0.258 24  15.899  <.0001
##
## Results are given on the log2 (not the response) scale.
##
## $contrasts
## Time = 6:
##  contrast   estimate    SE df t.ratio p.value
##  FGF2 - PBS   -0.151 0.365 24  -0.414  0.6826
##
## Time = 12:
##  contrast   estimate    SE df t.ratio p.value
##  FGF2 - PBS    0.152 0.365 24   0.415  0.6818
##
## Time = 24:
##  contrast   estimate    SE df t.ratio p.value
##  FGF2 - PBS    1.207 0.365 24   3.303  0.0030
##
## Time = 48:
##  contrast   estimate    SE df t.ratio p.value
##  FGF2 - PBS    0.754 0.365 24   2.065  0.0499
##
## Results are given on the log2 (not the response) scale.
```

```
p.adjust(aS4_2tLDHA$contrasts$p.value, method="fdr")
```

```
## [1] 0.68259089 0.68259089 0.01194852 0.09983677
```

```
aS4_2tLDHB <- test(emmeans(aS4_2LDHB, pairwise~Chemical|Time))
aS4_2tLDHB
```

```
## $emmeans
## Time = 6:
##  Chemical emmean    SE df t.ratio p.value
##  FGF2     -0.496 0.224 24  -2.209  0.0370
```

```
##  PBS      -0.380 0.224 24  -1.692  0.1036
##
## Time = 12:
##  Chemical emmean    SE df t.ratio p.value
##  FGF2      0.200 0.224 24   0.893  0.3807
##  PBS      -0.289 0.224 24  -1.290  0.2093
##
## Time = 24:
##  Chemical emmean    SE df t.ratio p.value
##  FGF2     -0.142 0.224 24  -0.632  0.5336
##  PBS      -0.120 0.224 24  -0.535  0.5974
##
## Time = 48:
##  Chemical emmean    SE df t.ratio p.value
##  FGF2     -1.072 0.224 24  -4.779  0.0001
##  PBS      -1.004 0.224 24  -4.478  0.0002
##
## Results are given on the log2 (not the response) scale.
##
## $contrasts
## Time = 6:
##  contrast   estimate    SE df t.ratio p.value
##  FGF2 - PBS  -0.1160 0.317 24  -0.366  0.7177
##
## Time = 12:
##  contrast   estimate    SE df t.ratio p.value
##  FGF2 - PBS   0.4897 0.317 24   1.544  0.1358
##
## Time = 24:
##  contrast   estimate    SE df t.ratio p.value
##  FGF2 - PBS  -0.0216 0.317 24  -0.068  0.9462
##
## Time = 48:
##  contrast   estimate    SE df t.ratio p.value
##  FGF2 - PBS  -0.0675 0.317 24  -0.213  0.8334
##
## Results are given on the log2 (not the response) scale.
```

```r
p.adjust(aS4_2tLDHB$contrasts$p.value, method="fdr")
```

```
## [1] 0.9462344 0.5430738 0.9462344 0.9462344
```

### Experiment 2 (Figure 2E)

Two factors: Chemical (levels: PBS, FGF2) and Time (levels: 24 and 48). Measuring the ratio between EdU+/DAPI cells and pHH3/DAPI cells. Since the responses are counts, including small counts and 0's, we will analyze using the negative binomial distribution.

Comparisons for Figure 2E: PBS vs FGF2 at each time point for both EdU and pHH3

#### Data and Visualization

```
d2E <- read.xlsx(xlsxFile = "MetabolismPaperRawData_Byran_working.xlsx", colNames = TRUE, sheet="Fig 2_I
glimpse(d2E)
```

```
## Rows: 20
## Columns: 8
## $ Chemical      <chr> "PBS", "PBS", "PBS", "PBS", "PBS", "FGF2", "FGF2", "FGF2~
## $ Sample        <dbl> 1, 2, 3, 4, 5, 1, 2, 3, 4, 5, 1, 2, 3, 4, 5, 1, 2, 3, 4,~
## $ `Time.(h)`    <dbl> 24, 24, 24, 24, 24, 24, 24, 24, 24, 24, 48, 48, 48, 48, ~
## $ `Edu/DAPI`    <dbl> 0.00000000, 0.00606061, 0.08510638, 0.00000000, 0.000000~
## $ `pHH3/DAPI`   <dbl> 0.00000000, 0.00606061, 0.02127660, 0.00000000, 0.000000~
## $ `Edu+.Cells`  <dbl> 0, 1, 8, 0, 0, 37, 24, 26, 16, 21, 103, 59, 2, 4, 6, 44,~
## $ `DAPI+.Cells` <dbl> 81, 165, 94, 88, 63, 92, 61, 80, 52, 83, 5, 2, 43, 43, 5~
## $ `pHH3+.Cells` <dbl> 0, 1, 2, 0, 0, 6, 8, 4, 2, 3, 0, 0, 1, 1, 0, 16, 8, 4, 6~
```

```
d2E_long <- d2E %>%
  rename(Time=`Time.(h)`, Edu_DAPI_ratio=`Edu/DAPI`, pHH3_DAPI_ratio=`pHH3/DAPI`, Edu=`Edu+.Cells`, DAPI
  pivot_longer(cols=4:5,names_to="RatioType",values_to="Ratio") %>%
  mutate(Time=as.factor(Time))
glimpse(d2E_long)
```

```
## Rows: 40
## Columns: 8
## $ Chemical  <chr> "PBS", "PBS", "PBS", "PBS", "PBS", "PBS", "PBS", "PBS", "PBS~
## $ Sample    <dbl> 1, 1, 2, 2, 3, 3, 4, 4, 5, 5, 1, 1, 2, 2, 3, 3, 4, 4, 5, 5, ~
## $ Time      <fct> 24, 24, 24, 24, 24, 24, 24, 24, 24, 24, 24, 24, 24, 24, 24, ~
## $ Edu       <dbl> 0, 0, 1, 1, 8, 8, 0, 0, 0, 0, 37, 37, 24, 24, 26, 26, 16, 16~
## $ DAPI      <dbl> 81, 81, 165, 165, 94, 94, 88, 88, 63, 63, 92, 92, 61, 61, 80~
## $ pHH3      <dbl> 0, 0, 1, 1, 2, 2, 0, 0, 0, 0, 6, 6, 8, 8, 4, 4, 2, 2, 3, 3, ~
## $ RatioType <chr> "Edu_DAPI_ratio", "pHH3_DAPI_ratio", "Edu_DAPI_ratio", "pHH3~
## $ Ratio     <dbl> 0.00000000, 0.00000000, 0.00606061, 0.00606061, 0.08510638, ~
```

```
ggplot(d2E_long, aes(x=Time,y=Ratio,color=Chemical)) +
  geom_jitter(width=.05,height=0) +
  facet_wrap(~RatioType, nrow=1) +
  labs(title="For Figure 2E: Treatment Comparisons by Time (no log)")
```

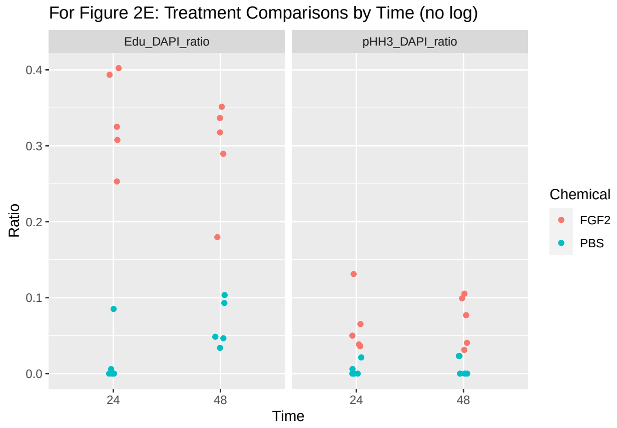

```r
ggplot(d2E_long, aes(x=Time,y=log(Ratio),color=Chemical)) +
  geom_jitter(width=.05,height=0) +
  facet_wrap(~RatioType, nrow=1) +
  labs(title="For Figure 2E: Treatment Comparisons by Time")
```

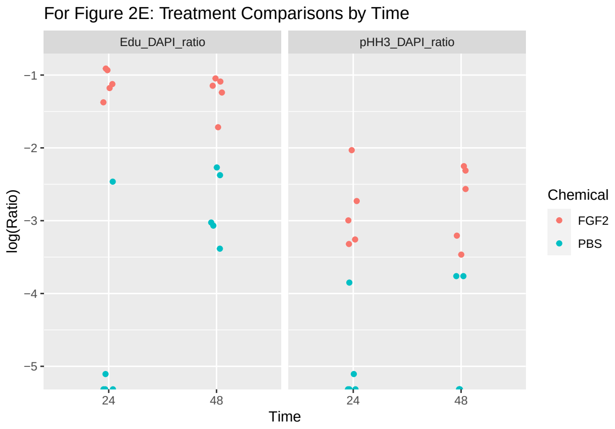

#### Data Analysis (Experiment 2)

```r
d2E <- d2E %>%
  rename(Time=`Time.(h)`, Edu_DAPI_ratio=`Edu/DAPI`, pHH3_DAPI_ratio=`pHH3/DAPI`, Edu=`Edu+.Cells`, DAPI
  mutate(Time=as.factor(Time))
glimpse(d2E)
```

```
## Rows: 20
## Columns: 8
## $ Chemical        <chr> "PBS", "PBS", "PBS", "PBS", "PBS", "FGF2", "FGF2", "FG~
## $ Sample          <dbl> 1, 2, 3, 4, 5, 1, 2, 3, 4, 5, 1, 2, 3, 4, 5, 1, 2, 3, ~
## $ Time            <fct> 24, 24, 24, 24, 24, 24, 24, 24, 24, 24, 48, 48, 48, 48~
## $ Edu_DAPI_ratio  <dbl> 0.00000000, 0.00606061, 0.08510638, 0.00000000, 0.0000~
## $ pHH3_DAPI_ratio <dbl> 0.00000000, 0.00606061, 0.02127660, 0.00000000, 0.0000~
## $ Edu             <dbl> 0, 1, 8, 0, 0, 37, 24, 26, 16, 21, 103, 59, 2, 4, 6, 4~
## $ DAPI            <dbl> 81, 165, 94, 88, 63, 92, 61, 80, 52, 83, 5, 2, 43, 43,~
## $ pHH3            <dbl> 0, 1, 2, 0, 0, 6, 8, 4, 2, 3, 0, 0, 1, 1, 0, 16, 8, 4,~
```

```r
nb.Edu <- glm.nb(Edu~Chemical + Time + Chemical:Time + offset(log(DAPI)),data=d2E)
summary(nb.Edu)
```

```
##
## Call:
## glm.nb(formula = Edu ~ Chemical + Time + Chemical:Time + offset(log(DAPI)),
##     data = d2E, init.theta = 0.6306265286, link = log)
##
## Deviance Residuals:
##      Min        1Q    Median        3Q       Max
## -2.30880  -1.17253  -0.04744   0.13016   1.44780
##
```

```
## Coefficients:
##                     Estimate Std. Error z value Pr(>|z|)
## (Intercept)          -1.0898     0.5706  -1.910 0.056140 .
## ChemicalPBS          -2.8971     0.8725  -3.320 0.000899 ***
## Time48               -0.1314     0.8051  -0.163 0.870346
## ChemicalPBS:Time48    6.4138     1.1849   5.413  6.2e-08 ***
## ---
## Signif. codes:  0 '***' 0.001 '**' 0.01 '*' 0.05 '.' 0.1 ' ' 1
##
## (Dispersion parameter for Negative Binomial(0.6306) family taken to be 1)
##
##     Null deviance: 71.177  on 19  degrees of freedom
## Residual deviance: 23.090  on 16  degrees of freedom
## AIC: 174.86
##
## Number of Fisher Scoring iterations: 1
##
##
##               Theta:  0.631
##           Std. Err.:  0.198
##
##  2 x log-likelihood:  -164.858
```

```r
nb.pHH3 <- glm.nb(pHH3~Chemical + Time + Chemical:Time + offset(log(DAPI)),data=d2E)
summary(nb.pHH3)
```

```
##
## Call:
## glm.nb(formula = pHH3 ~ Chemical + Time + Chemical:Time + offset(log(DAPI)),
##     data = d2E, init.theta = 14.76929967, link = log)
##
## Deviance Residuals:
##     Min       1Q   Median       3Q      Max
## -1.5378  -0.9427  -0.2973   0.5002   1.5996
##
## Coefficients:
##                     Estimate Std. Error z value Pr(>|z|)
## (Intercept)          -2.7658     0.2391 -11.569  < 2e-16 ***
## ChemicalPBS          -2.3349     0.6377  -3.662 0.000251 ***
## Time48                0.1089     0.3050   0.357 0.721143
## ChemicalPBS:Time48    0.6710     0.9807   0.684 0.493882
## ---
## Signif. codes:  0 '***' 0.001 '**' 0.01 '*' 0.05 '.' 0.1 ' ' 1
##
## (Dispersion parameter for Negative Binomial(14.7693) family taken to be 1)
##
##     Null deviance: 50.014  on 19  degrees of freedom
## Residual deviance: 16.852  on 16  degrees of freedom
## AIC: 75.655
##
## Number of Fisher Scoring iterations: 1
##
##
##               Theta:  14.8
```

```
##           Std. Err.:  19.9
##
##  2 x log-likelihood:  -65.655
```

Again, we make the necessary comparisons, and then adjust the p-values to account for multiple comparisons.

```r
nb.Edu.t <- test(emmeans(nb.Edu, pairwise~Chemical|Time))
nb.Edu.t
```

```
## $emmeans
## Time = 24:
##  Chemical emmean    SE  df z.ratio p.value
##  FGF2      3.018 0.571 Inf   5.290  <.0001
##  PBS       0.121 0.660 Inf   0.184  0.8543
##
## Time = 48:
##  Chemical emmean    SE  df z.ratio p.value
##  FGF2      2.887 0.568 Inf   5.083  <.0001
##  PBS       6.404 0.566 Inf  11.317  <.0001
##
## Results are given on the log (not the response) scale.
##
## $contrasts
## Time = 24:
##  contrast   estimate    SE  df z.ratio p.value
##  FGF2 - PBS     2.90 0.873 Inf   3.320  0.0009
##
## Time = 48:
##  contrast   estimate    SE  df z.ratio p.value
##  FGF2 - PBS    -3.52 0.802 Inf  -4.387  <.0001
##
## Results are given on the log (not the response) scale.
```

```r
p.adjust(nb.Edu.t$contrasts$p.value, method="fdr")
```

```
## [1] 8.99302e-04 2.30285e-05
```

```r
nb.pHH3.t <- test(emmeans(nb.pHH3, pairwise~Chemical|Time))
nb.pHH3.t
```

```
## $emmeans
## Time = 24:
##  Chemical emmean    SE  df z.ratio p.value
##  FGF2      1.342 0.239 Inf   5.615  <.0001
##  PBS      -0.993 0.591 Inf  -1.679  0.0931
##
## Time = 48:
##  Chemical emmean    SE  df z.ratio p.value
##  FGF2      1.451 0.189 Inf   7.659  <.0001
##  PBS      -0.213 0.721 Inf  -0.295  0.7678
##
## Results are given on the log (not the response) scale.
```

```
##
## $contrasts
## Time = 24:
##  contrast   estimate    SE  df z.ratio p.value
##  FGF2 - PBS     2.33 0.638 Inf   3.662  0.0003
##
## Time = 48:
##  contrast   estimate    SE  df z.ratio p.value
##  FGF2 - PBS     1.66 0.745 Inf   2.233  0.0256
##
## Results are given on the log (not the response) scale.
```

```r
p.adjust(nb.pHH3.t$contrasts$p.value, method="fdr")
```

```
## [1] 0.0005012702 0.0255513831
```

### Experiment 3 (Figures 3E, 3F, 3G, S5F)

Responses are relative expressions of a number of genes. Three factors: FGF2 (Present or Absent), 2DG (Present or Absent), and Time (24 and 48). So 8 treatments, 4 replications/treatment, for a total of 32 experimental units.

Four comparisons each for Figure 3E, 3F, 3G, S5F: PBS vs FGF2; PBS vs 2DG; FGF2 vs FGF2+2DG, and 2DG vs FGF2+2DG at each time point for each gene.

#### Data and Visualization

We will first obtain the datasets for each relevant figure (Figures 3E, 3E, 3G, S5F), visualize them, then analyze them according to the specified set of comparisons.

##### Plots to Compare the Four Treatments at each Time Point

Here we compare PBS vs. FGF2 at each time point.

```r
d3E <- read.xlsx(xlsxFile = "MetabolismPaperRawData_Byran_working.xlsx",
                 colNames = TRUE, sheet="Fig 3_E")
glimpse(d3E)
```

```
## Rows: 32
## Columns: 9
## $ Chemical   <chr> "PBS", "PBS", "PBS", "PBS", "FGF2", "FGF2", "FGF2", "FGF2",~
## $ Sample     <dbl> 1, 2, 3, 4, 1, 2, 3, 4, 1, 2, 3, 4, 1, 2, 3, 4, 1, 2, 3, 4,~
## $ `Time.(h)` <dbl> 24, 24, 24, 24, 24, 24, 24, 24, 24, 24, 24, 24, 24, 24, 24,~
## $ SOX2       <dbl> 1.0768791, 1.0895012, 1.0097666, 0.8440814, 1.3118550, 0.91~
## $ SIX6       <dbl> 0.94656814, 1.06640843, 1.06739862, 0.92810671, 5.88316680,~
## $ PAX6       <dbl> 0.9288248, 1.3186025, 0.9589969, 0.8514029, 1.1119467, 1.56~
## $ RPE65      <dbl> 0.39900204, 1.97573108, 0.89361521, 1.41953629, 0.38652829,~
## $ TYR        <dbl> 0.62262023, 1.34223370, 1.33395453, 0.89703131, 0.18453189,~
## $ OTX2       <dbl> 0.8638951, 1.2395212, 1.0506627, 0.8888362, 0.3042530, 0.31~
```

```r
d3F <- read.xlsx(xlsxFile = "MetabolismPaperRawData_Byran_working.xlsx",
                 colNames = TRUE, sheet="Fig 3_F")
glimpse(d3F)
```

```
## Rows: 32
## Columns: 7
## $ Chemical   <chr> "PBS", "PBS", "PBS", "PBS", "FGF2", "FGF2", "FGF2", "FGF2",~
## $ Sample     <dbl> 1, 2, 3, 4, 1, 2, 3, 4, 1, 2, 3, 4, 1, 2, 3, 4, 1, 2, 3, 4,~
## $ `Time.(h)` <dbl> 24, 24, 24, 24, 24, 24, 24, 24, 24, 24, 24, 24, 24, 24, 24,~
## $ HK1        <dbl> 0.9345654, 0.9719146, 1.1105429, 0.9913496, 0.9108738, 0.94~
## $ ENO1       <dbl> 0.8344102, 1.3613660, 0.9864293, 0.8924410, 0.8202481, 0.82~
## $ LDHA       <dbl> 0.9439363, 1.3617872, 1.0260627, 0.7581832, 1.0208911, 1.13~
## $ LDHB       <dbl> 0.9044622, 1.3056468, 0.9359639, 0.9047420, 1.2114672, 1.22~
```

```r
d3G <- read.xlsx(xlsxFile = "MetabolismPaperRawData_Byran_working.xlsx",
                 colNames = TRUE, sheet="Fig 3_G")
glimpse(d3G)
```

```
## Rows: 32
## Columns: 5
## $ Chemical   <chr> "PBS", "PBS", "PBS", "PBS", "FGF2", "FGF2", "FGF2", "FGF2",~
## $ Sample     <dbl> 1, 2, 3, 4, 1, 2, 3, 4, 1, 2, 3, 4, 1, 2, 3, 4, 1, 2, 3, 4,~
## $ `Time.(h)` <dbl> 24, 24, 24, 24, 24, 24, 24, 24, 24, 24, 24, 24, 24, 24, 24,~
## $ E2F1       <dbl> 0.8986053, 1.4141304, 0.9211225, 0.8543271, 0.7966359, 0.87~
## $ PCNA       <dbl> 0.8245521, 1.3507035, 0.9894602, 0.9074518, 0.8719502, 1.29~
```

```r
d5SF <- read.xlsx(xlsxFile = "MetabolismPaperRawData_Byran_working.xlsx",
                  colNames = TRUE, sheet="Fig 5S_F")
glimpse(d5SF)
```

```
## Rows: 32
## Columns: 7
## $ Chemical   <chr> "PBS", "PBS", "PBS", "PBS", "FGF2", "FGF2", "FGF2", "FGF2",~
## $ Sample     <dbl> 1, 2, 3, 4, 1, 2, 3, 4, 1, 2, 3, 4, 1, 2, 3, 4, 1, 2, 3, 4,~
## $ `Time.(h)` <dbl> 24, 24, 24, 24, 24, 24, 24, 24, 24, 24, 24, 24, 24, 24, 24,~
## $ HK2        <dbl> 0.8728273, 1.2285455, 0.9407780, 0.9912730, 1.5591569, 2.21~
## $ TPI1       <dbl> 0.8604943, 1.1873188, 0.8730974, 1.1210423, 1.1849948, 1.11~
## $ PGAM1      <dbl> 0.8995324, 1.3276418, 0.9521238, 0.8794453, 0.8828983, 1.20~
## $ PKLR       <dbl> 0.9844731, 1.2367123, 0.9637804, 0.8522154, 0.8773619, 1.07~
```

```r
# to compare the 4 treatments at each time point
d3E_long <- d3E %>%
  rename(Time=`Time.(h)`) %>%
  pivot_longer(cols=4:9,names_to="Gene",values_to="RGE") %>%
  mutate(Time=as.factor(Time))
glimpse(d3E_long)
```

```
## Rows: 192
## Columns: 5
## $ Chemical <chr> "PBS", "PBS", "PBS", "PBS", "PBS", "PBS", "PBS", "PBS", "PBS"~
## $ Sample   <dbl> 1, 1, 1, 1, 1, 1, 2, 2, 2, 2, 2, 2, 3, 3, 3, 3, 3, 3, 4, 4, 4~
```

```
## $ Time     <fct> 24, 24, 24, 24, 24, 24, 24, 24, 24, 24, 24, 24, 24, 24, 24, 2~
## $ Gene     <chr> "SOX2", "SIX6", "PAX6", "RPE65", "TYR", "OTX2", "SOX2", "SIX6~
## $ RGE      <dbl> 1.0768791, 0.9465681, 0.9288248, 0.3990020, 0.6226202, 0.8638~
```

```
d3F_long <- d3F %>%
  rename(Time=`Time.(h)`) %>%
  pivot_longer(cols=4:7,names_to="Gene",values_to="RGE") %>%
  mutate(Time=as.factor(Time))
glimpse(d3F_long)
```

```
## Rows: 128
## Columns: 5
## $ Chemical <chr> "PBS", "PBS", "PBS", "PBS", "PBS", "PBS", "PBS", "PBS", "PBS"~
## $ Sample   <dbl> 1, 1, 1, 1, 2, 2, 2, 2, 3, 3, 3, 3, 4, 4, 4, 4, 1, 1, 1, 1, 2~
## $ Time     <fct> 24, 24, 24, 24, 24, 24, 24, 24, 24, 24, 24, 24, 24, 24, 24, 2~
## $ Gene     <chr> "HK1", "ENO1", "LDHA", "LDHB", "HK1", "ENO1", "LDHA", "LDHB",~
## $ RGE      <dbl> 0.9345654, 0.8344102, 0.9439363, 0.9044622, 0.9719146, 1.3613~
```

```
d3G_long <- d3G %>%
  rename(Time=`Time.(h)`) %>%
  pivot_longer(cols=4:5,names_to="Gene",values_to="RGE") %>%
  mutate(Time=as.factor(Time))
glimpse(d3G_long)
```

```
## Rows: 64
## Columns: 5
## $ Chemical <chr> "PBS", "PBS", "PBS", "PBS", "PBS", "PBS", "PBS", "PBS", "FGF2~
## $ Sample   <dbl> 1, 1, 2, 2, 3, 3, 4, 4, 1, 1, 2, 2, 3, 3, 4, 4, 1, 1, 2, 2, 3~
## $ Time     <fct> 24, 24, 24, 24, 24, 24, 24, 24, 24, 24, 24, 24, 24, 24, 24, 2~
## $ Gene     <chr> "E2F1", "PCNA", "E2F1", "PCNA", "E2F1", "PCNA", "E2F1", "PCNA~
## $ RGE      <dbl> 0.8986053, 0.8245521, 1.4141304, 1.3507035, 0.9211225, 0.9894~
```

```
d5SF_long <- d5SF %>%
  rename(Time=`Time.(h)`) %>%
  pivot_longer(cols=4:7,names_to="Gene",values_to="RGE") %>%
  mutate(Time=as.factor(Time))
glimpse(d5SF_long)
```

```
## Rows: 128
## Columns: 5
## $ Chemical <chr> "PBS", "PBS", "PBS", "PBS", "PBS", "PBS", "PBS", "PBS", "PBS"~
## $ Sample   <dbl> 1, 1, 1, 1, 2, 2, 2, 2, 3, 3, 3, 3, 4, 4, 4, 4, 1, 1, 1, 1, 2~
## $ Time     <fct> 24, 24, 24, 24, 24, 24, 24, 24, 24, 24, 24, 24, 24, 24, 24, 2~
## $ Gene     <chr> "HK2", "TPI1", "PGAM1", "PKLR", "HK2", "TPI1", "PGAM1", "PKLR~
## $ RGE      <dbl> 0.8728273, 0.8604943, 0.8995324, 0.9844731, 1.2285455, 1.1873~
```

```
ggplot(d3E_long, aes(x=Time,y=log2(RGE),color=Chemical)) +
  geom_jitter(width=.05,height=0) +
  facet_wrap(~Gene, nrow=3) +
  labs(title="For Figure 3E: Time Comparisons")
```

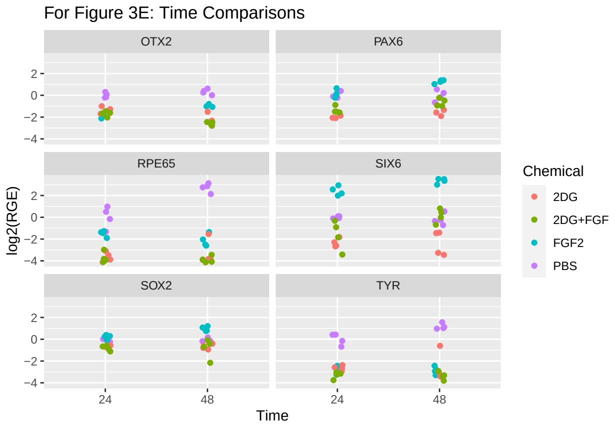

```r
ggplot(d3F_long, aes(x=Time,y=log2(RGE),color=Chemical)) +
  geom_jitter(width=.05,height=0) +
  facet_wrap(~Gene, nrow=2) +
  labs(title="For Figure 3F: Time Comparisons")
```

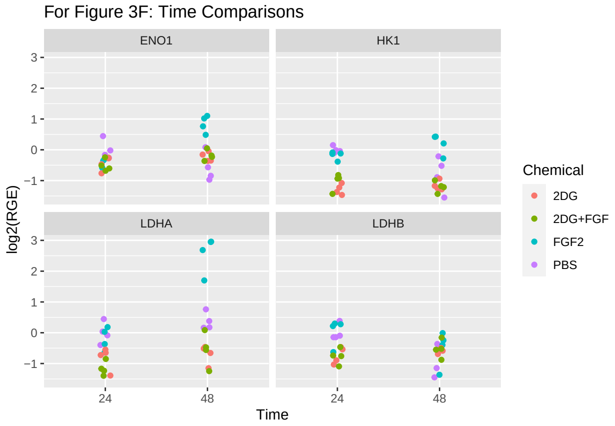

```r
ggplot(d3G_long, aes(x=Time,y=log2(RGE),color=Chemical)) +
  geom_jitter(width=.05,height=0) +
  facet_wrap(~Gene, nrow=1) +
  labs(title="For Figure 3G: Time Comparisons")
```

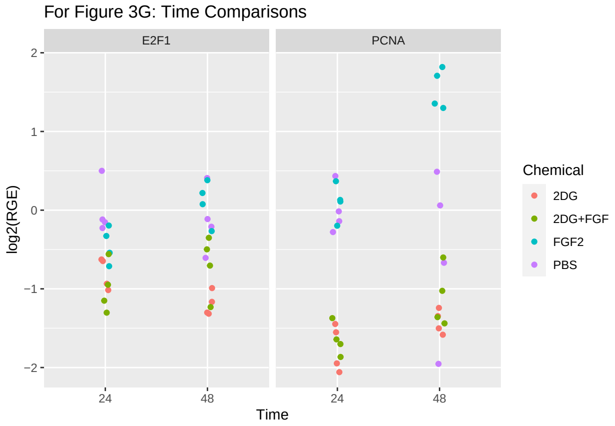

```r
ggplot(d5SF_long, aes(x=Time,y=log2(RGE),color=Chemical)) +
  geom_jitter(width=.05,height=0) +
  facet_wrap(~Gene, nrow=2) +
  labs(title="For Figure S5F: Time Comparisons")
```

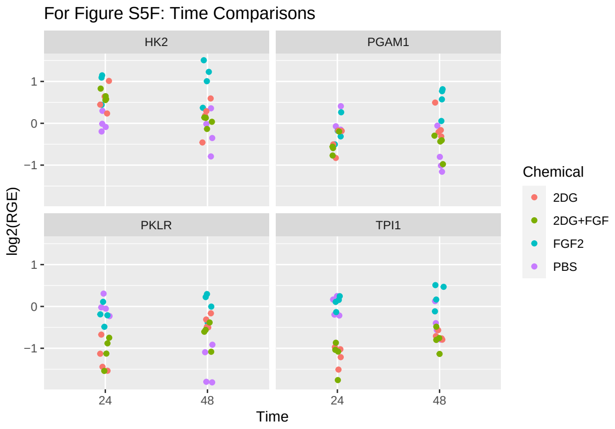

#### Data Analysis (Experiment 3)

The analysis here is to compare the four specified pairs of means across the two time points. We will treat this like a two-way ANOVA, with Treatment and Time as the factors, and in the end pull out the pairs of means of interest and adjust their p-values by controlling the False Discovery Rate across the eight pairs of means for each dataset.

```r
d3E <- d3E %>%
  rename(Time=`Time.(h)`) %>%
  mutate(Time=as.factor(Time))
glimpse(d3E)
```

```
## Rows: 32
## Columns: 9
## $ Chemical <chr> "PBS", "PBS", "PBS", "PBS", "FGF2", "FGF2", "FGF2", "FGF2", "~
## $ Sample   <dbl> 1, 2, 3, 4, 1, 2, 3, 4, 1, 2, 3, 4, 1, 2, 3, 4, 1, 2, 3, 4, 1~
## $ Time     <fct> 24, 24, 24, 24, 24, 24, 24, 24, 24, 24, 24, 24, 24, 24, 24, 2~
## $ SOX2     <dbl> 1.0768791, 1.0895012, 1.0097666, 0.8440814, 1.3118550, 0.9122~
## $ SIX6     <dbl> 0.94656814, 1.06640843, 1.06739862, 0.92810671, 5.88316680, 7~
## $ PAX6     <dbl> 0.9288248, 1.3186025, 0.9589969, 0.8514029, 1.1119467, 1.5670~
## $ RPE65    <dbl> 0.39900204, 1.97573108, 0.89361521, 1.41953629, 0.38652829, 0~
## $ TYR      <dbl> 0.62262023, 1.34223370, 1.33395453, 0.89703131, 0.18453189, 0~
## $ OTX2     <dbl> 0.8638951, 1.2395212, 1.0506627, 0.8888362, 0.3042530, 0.3164~
```

```r
d3F <- d3F %>%
  rename(Time=`Time.(h)`) %>%
  mutate(Time=as.factor(Time))
glimpse(d3F)
```

```
## Rows: 32
## Columns: 7
```

```
## $ Chemical <chr> "PBS", "PBS", "PBS", "PBS", "FGF2", "FGF2", "FGF2", "FGF2", "~
## $ Sample   <dbl> 1, 2, 3, 4, 1, 2, 3, 4, 1, 2, 3, 4, 1, 2, 3, 4, 1, 2, 3, 4, 1~
## $ Time     <fct> 24, 24, 24, 24, 24, 24, 24, 24, 24, 24, 24, 24, 24, 24, 24, 2~
## $ HK1      <dbl> 0.9345654, 0.9719146, 1.1105429, 0.9913496, 0.9108738, 0.9417~
## $ ENO1     <dbl> 0.8344102, 1.3613660, 0.9864293, 0.8924410, 0.8202481, 0.8236~
## $ LDHA     <dbl> 0.9439363, 1.3617872, 1.0260627, 0.7581832, 1.0208911, 1.1352~
## $ LDHB     <dbl> 0.9044622, 1.3056468, 0.9359639, 0.9047420, 1.2114672, 1.2290~
```

```
d3G <- d3G %>%
  rename(Time=`Time.(h)`) %>%
  mutate(Time=as.factor(Time))
glimpse(d3G)
```

```
## Rows: 32
## Columns: 5
## $ Chemical <chr> "PBS", "PBS", "PBS", "PBS", "FGF2", "FGF2", "FGF2", "FGF2", "~
## $ Sample   <dbl> 1, 2, 3, 4, 1, 2, 3, 4, 1, 2, 3, 4, 1, 2, 3, 4, 1, 2, 3, 4, 1~
## $ Time     <fct> 24, 24, 24, 24, 24, 24, 24, 24, 24, 24, 24, 24, 24, 24, 24, 2~
## $ E2F1     <dbl> 0.8986053, 1.4141304, 0.9211225, 0.8543271, 0.7966359, 0.8732~
## $ PCNA     <dbl> 0.8245521, 1.3507035, 0.9894602, 0.9074518, 0.8719502, 1.2903~
```

```
d5SF <- d5SF %>%
  rename(Time=`Time.(h)`) %>%
  mutate(Time=as.factor(Time))
glimpse(d5SF)
```

```
## Rows: 32
## Columns: 7
## $ Chemical <chr> "PBS", "PBS", "PBS", "PBS", "FGF2", "FGF2", "FGF2", "FGF2", "~
## $ Sample   <dbl> 1, 2, 3, 4, 1, 2, 3, 4, 1, 2, 3, 4, 1, 2, 3, 4, 1, 2, 3, 4, 1~
## $ Time     <fct> 24, 24, 24, 24, 24, 24, 24, 24, 24, 24, 24, 24, 24, 24, 24, 2~
## $ HK2      <dbl> 0.8728273, 1.2285455, 0.9407780, 0.9912730, 1.5591569, 2.2125~
## $ TPI1     <dbl> 0.8604943, 1.1873188, 0.8730974, 1.1210423, 1.1849948, 1.1142~
## $ PGAM1    <dbl> 0.8995324, 1.3276418, 0.9521238, 0.8794453, 0.8828983, 1.2000~
## $ PKLR     <dbl> 0.9844731, 1.2367123, 0.9637804, 0.8522154, 0.8773619, 1.0796~
```

##### Treatment Comparisons (Figure 3E)

```
a3E_SOX2 <- aov(log2(SOX2)~Chemical*Time, data=d3E)
a3E_SIX6 <- aov(log2(SIX6)~Chemical*Time, data=d3E)
a3E_PAX6 <- aov(log2(PAX6)~Chemical*Time, data=d3E)
a3E_RPE65 <- aov(log2(RPE65)~Chemical*Time, data=d3E)
a3E_TYR <- aov(log2(TYR)~Chemical*Time, data=d3E)
a3E_OTX2 <- aov(log2(OTX2)~Chemical*Time, data=d3E)

summary(a3E_SOX2)
```

```
##                Df Sum Sq Mean Sq F value  Pr(>F)
## Chemical        3  9.926   3.309  23.759 2.3e-07 ***
## Time            1  0.201   0.201   1.444   0.241
```

```
## Chemical:Time  3  0.945   0.315   2.262   0.107
## Residuals     24  3.342   0.139
## ---
## Signif. codes:  0 '***' 0.001 '**' 0.01 '*' 0.05 '.' 0.1 ' ' 1
```

```r
plot(a3E_SOX2,which=1:2)
```

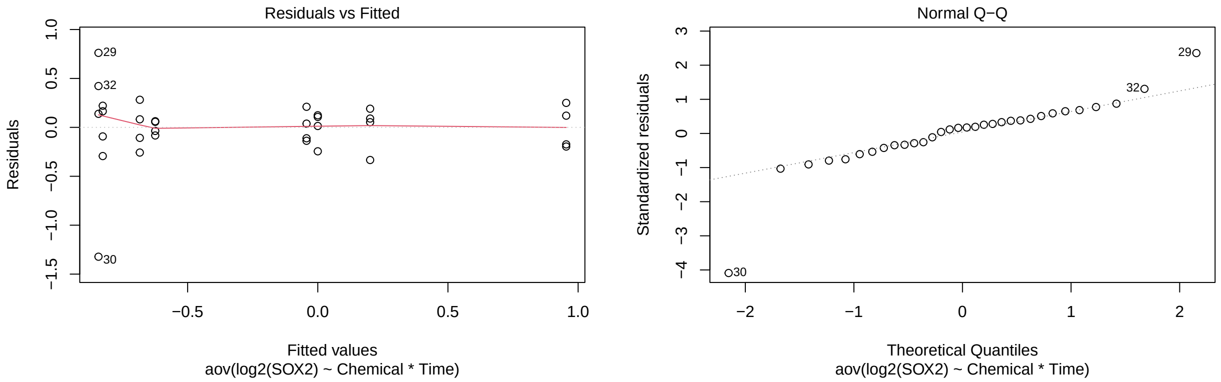

```r
plot(a3E_SIX6,which=1:2)
```

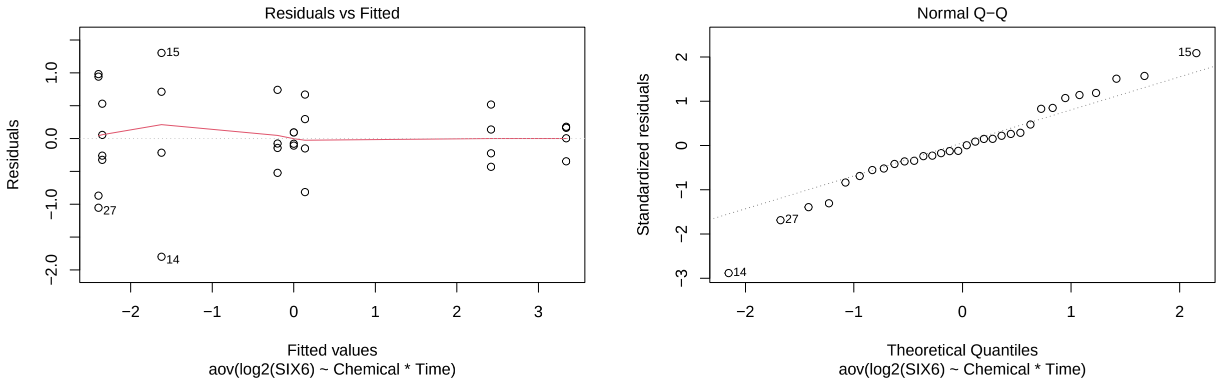

```r
plot(a3E_PAX6,which=1:2)
```

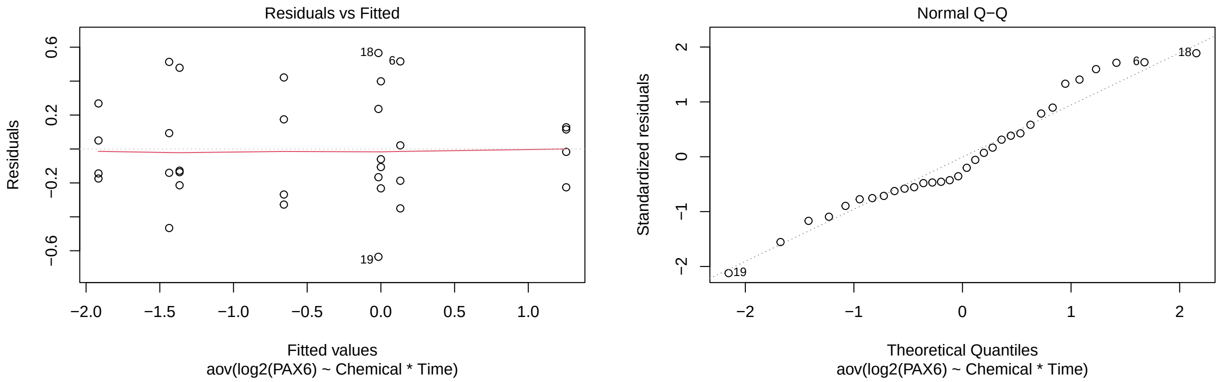

```r
plot(a3E_RPE65,which=1:2)
```

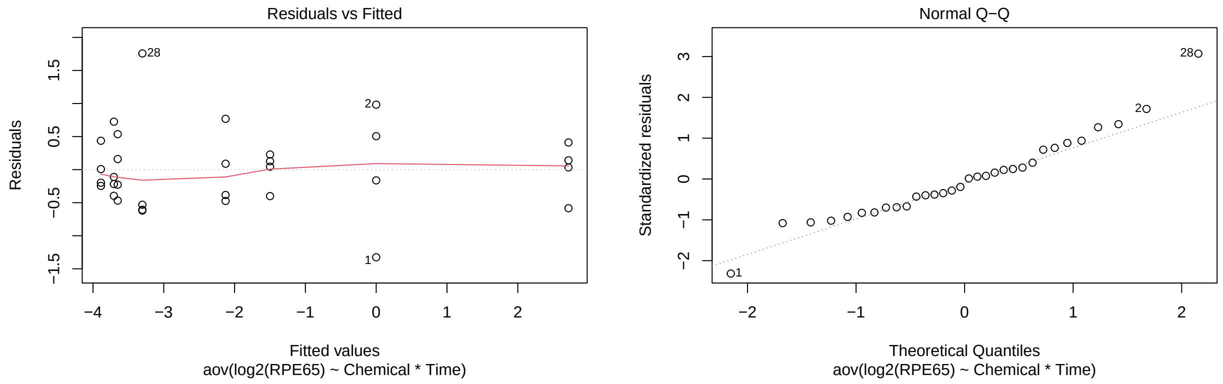

```r
plot(a3E_TYR,which=1:2)
```

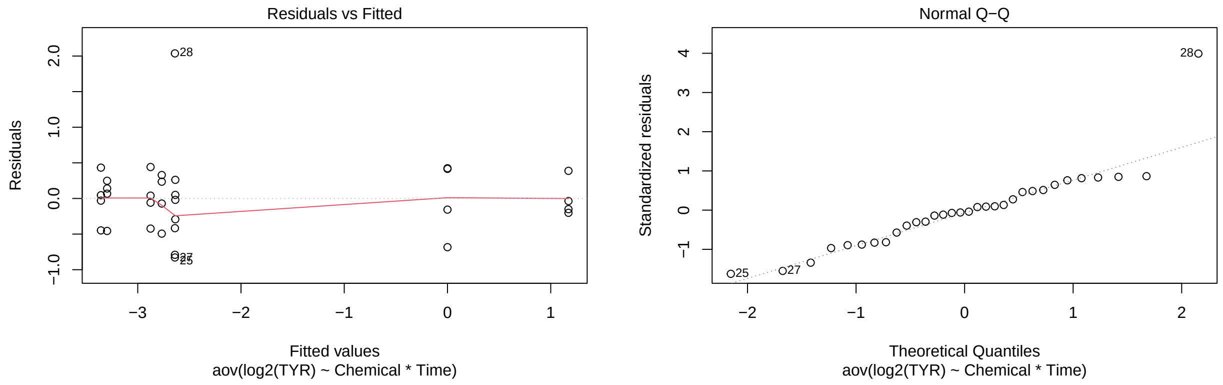

```r
plot(a3E_OTX2,which=1:2)
```

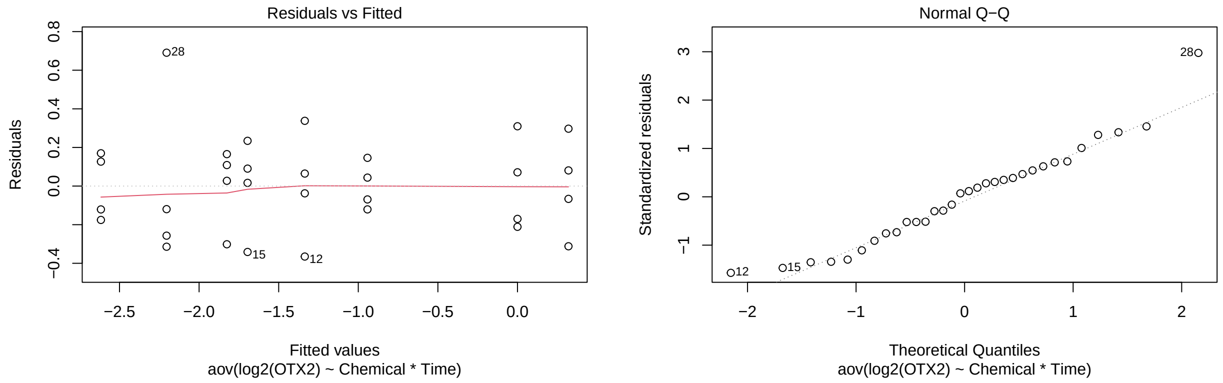

```r
a3E_tSOX2 <- test(emmeans(a3E_SOX2, pairwise~Chemical|Time), adjust="none")
a3E_tSOX2
```

```
## $emmeans
## Time = 24:
##  Chemical  emmean    SE df t.ratio p.value
##  2DG      -0.6244 0.187 24  -3.346  0.0027
##  2DG+FGF  -0.8260 0.187 24  -4.427  0.0002
##  FGF2      0.2011 0.187 24   1.078  0.2919
##  PBS       0.0000 0.187 24   0.000  1.0000
##
## Time = 48:
##  Chemical  emmean    SE df t.ratio p.value
##  2DG      -0.6837 0.187 24  -3.664  0.0012
##  2DG+FGF  -0.8425 0.187 24  -4.516  0.0001
##  FGF2      0.9543 0.187 24   5.115  <.0001
##  PBS      -0.0432 0.187 24  -0.232  0.8188
##
## Results are given on the log2 (not the response) scale.
##
## $contrasts
## Time = 24:
##  contrast         estimate    SE df t.ratio p.value
##  2DG - (2DG+FGF)     0.202 0.264 24   0.764  0.4523
##  2DG - FGF2         -0.825 0.264 24  -3.128  0.0046
##  2DG - PBS          -0.624 0.264 24  -2.366  0.0264
##  (2DG+FGF) - FGF2   -1.027 0.264 24  -3.892  0.0007
##  (2DG+FGF) - PBS    -0.826 0.264 24  -3.130  0.0045
##  FGF2 - PBS          0.201 0.264 24   0.762  0.4535
##
## Time = 48:
##  contrast         estimate    SE df t.ratio p.value
##  2DG - (2DG+FGF)     0.159 0.264 24   0.602  0.5528
##  2DG - FGF2         -1.638 0.264 24  -6.208  <.0001
##  2DG - PBS          -0.640 0.264 24  -2.427  0.0231
##  (2DG+FGF) - FGF2   -1.797 0.264 24  -6.810  <.0001
##  (2DG+FGF) - PBS    -0.799 0.264 24  -3.029  0.0058
##  FGF2 - PBS          0.998 0.264 24   3.780  0.0009
##
## Results are given on the log2 (not the response) scale.
```

```r
p.adjust(a3E_tSOX2$contrasts$p.value[c(1,3,4,6,7,9,10,12)], method="fdr")
```

```
## [1] 5.182300e-01 4.220957e-02 2.443133e-03 5.182300e-01 5.528162e-01
## [6] 4.220957e-02 3.860222e-06 2.443133e-03
```

```r
a3E_tSIX6 <- test(emmeans(a3E_SIX6, pairwise~Chemical|Time), adjust="none")
a3E_tSIX6
```

```
## $emmeans
## Time = 24:
##  Chemical emmean   SE df t.ratio p.value
##  2DG      -2.349 0.36 24  -6.516  <.0001
##  2DG+FGF  -1.622 0.36 24  -4.501  0.0001
##  FGF2      2.419 0.36 24   6.712  <.0001
##  PBS       0.000 0.36 24   0.000  1.0000
```

```
##
## Time = 48:
##  Chemical emmean   SE df t.ratio p.value
##  2DG      -2.395 0.36 24  -6.645  <.0001
##  2DG+FGF   0.138 0.36 24   0.382  0.7056
##  FGF2      3.341 0.36 24   9.269  <.0001
##  PBS      -0.199 0.36 24  -0.553  0.5855
##
## Results are given on the log2 (not the response) scale.
##
## $contrasts
## Time = 24:
##  contrast         estimate   SE df t.ratio p.value
##  2DG - (2DG+FGF)    -0.726 0.51 24  -1.425  0.1671
##  2DG - FGF2         -4.768 0.51 24  -9.353  <.0001
##  2DG - PBS          -2.349 0.51 24  -4.607  0.0001
##  (2DG+FGF) - FGF2   -4.042 0.51 24  -7.928  <.0001
##  (2DG+FGF) - PBS    -1.622 0.51 24  -3.183  0.0040
##  FGF2 - PBS          2.419 0.51 24   4.746  0.0001
##
## Time = 48:
##  contrast         estimate   SE df t.ratio p.value
##  2DG - (2DG+FGF)    -2.533 0.51 24  -4.969  <.0001
##  2DG - FGF2         -5.736 0.51 24 -11.252  <.0001
##  2DG - PBS          -2.196 0.51 24  -4.308  0.0002
##  (2DG+FGF) - FGF2   -3.203 0.51 24  -6.284  <.0001
##  (2DG+FGF) - PBS     0.337 0.51 24   0.661  0.5148
##  FGF2 - PBS          3.540 0.51 24   6.945  <.0001
##
## Results are given on the log2 (not the response) scale.
```

```r
p.adjust(a3E_tSIX6$contrasts$p.value[c(1,3,4,6,7,9,10,12)], method="fdr")
```

```
## [1] 1.671271e-01 1.502965e-04 2.959284e-07 1.268252e-04 8.995434e-05
## [6] 2.760176e-04 4.543491e-06 1.403019e-06
```

```r
a3E_tPAX6 <- test(emmeans(a3E_PAX6, pairwise~Chemical|Time), adjust="none")
a3E_tPAX6
```

```
## $emmeans
## Time = 24:
##  Chemical  emmean    SE df t.ratio p.value
##  2DG      -1.9161 0.173 24 -11.072  <.0001
##  2DG+FGF  -1.3659 0.173 24  -7.893  <.0001
##  FGF2      0.1318 0.173 24   0.761  0.4538
##  PBS       0.0000 0.173 24   0.000  1.0000
##
## Time = 48:
##  Chemical  emmean    SE df t.ratio p.value
##  2DG      -1.4364 0.173 24  -8.300  <.0001
##  2DG+FGF  -0.6582 0.173 24  -3.804  0.0009
##  FGF2      1.2572 0.173 24   7.265  <.0001
##  PBS      -0.0167 0.173 24  -0.097  0.9239
```

```
##
## Results are given on the log2 (not the response) scale.
##
## $contrasts
## Time = 24:
##  contrast          estimate    SE df t.ratio p.value
##  2DG - (2DG+FGF)     -0.550 0.245 24  -2.248  0.0340
##  2DG - FGF2          -2.048 0.245 24  -8.368  <.0001
##  2DG - PBS           -1.916 0.245 24  -7.829  <.0001
##  (2DG+FGF) - FGF2    -1.498 0.245 24  -6.120  <.0001
##  (2DG+FGF) - PBS     -1.366 0.245 24  -5.581  <.0001
##  FGF2 - PBS           0.132 0.245 24   0.538  0.5953
##
## Time = 48:
##  contrast          estimate    SE df t.ratio p.value
##  2DG - (2DG+FGF)     -0.778 0.245 24  -3.180  0.0040
##  2DG - FGF2          -2.694 0.245 24 -11.007  <.0001
##  2DG - PBS           -1.420 0.245 24  -5.801  <.0001
##  (2DG+FGF) - FGF2    -1.915 0.245 24  -7.827  <.0001
##  (2DG+FGF) - PBS     -0.642 0.245 24  -2.621  0.0150
##  FGF2 - PBS           1.274 0.245 24   5.205  <.0001
##
## Results are given on the log2 (not the response) scale.
```

```
p.adjust(a3E_tPAX6$contrasts$p.value[c(1,3,4,6,7,9,10,12)], method="fdr")
```

```
## [1] 3.887945e-02 1.855668e-07 6.776658e-06 5.952755e-01 5.377285e-03
## [6] 1.113527e-05 1.855668e-07 3.955075e-05
```

```
a3E_tRPE65 <- test(emmeans(a3E_RPE65, pairwise~Chemical|Time), adjust="none")
a3E_tRPE65
```

```
## $emmeans
## Time = 24:
##  Chemical emmean   SE df t.ratio p.value
##  2DG       -3.65 0.33 24 -11.049  <.0001
##  2DG+FGF   -3.70 0.33 24 -11.212  <.0001
##  FGF2      -1.50 0.33 24  -4.537  0.0001
##  PBS        0.00 0.33 24   0.000  1.0000
##
## Time = 48:
##  Chemical emmean   SE df t.ratio p.value
##  2DG       -3.30 0.33 24  -9.995  <.0001
##  2DG+FGF   -3.89 0.33 24 -11.767  <.0001
##  FGF2      -2.13 0.33 24  -6.438  <.0001
##  PBS        2.72 0.33 24   8.220  <.0001
##
## Results are given on the log2 (not the response) scale.
##
## $contrasts
## Time = 24:
##  contrast          estimate    SE df t.ratio p.value
##  2DG - (2DG+FGF)     0.0536 0.467 24   0.115  0.9096
```

```
##  2DG - FGF2        -2.1517 0.467 24  -4.605  0.0001
##  2DG - PBS         -3.6504 0.467 24  -7.813  <.0001
##  (2DG+FGF) - FGF2  -2.2052 0.467 24  -4.720  0.0001
##  (2DG+FGF) - PBS   -3.7040 0.467 24  -7.928  <.0001
##  FGF2 - PBS        -1.4988 0.467 24  -3.208  0.0038
##
## Time = 48:
##  contrast         estimate    SE df t.ratio p.value
##  2DG - (2DG+FGF)    0.5856 0.467 24   1.253  0.2222
##  2DG - FGF2        -1.1751 0.467 24  -2.515  0.0190
##  2DG - PBS         -6.0177 0.467 24 -12.880  <.0001
##  (2DG+FGF) - FGF2  -1.7607 0.467 24  -3.768  0.0009
##  (2DG+FGF) - PBS   -6.6032 0.467 24 -14.133  <.0001
##  FGF2 - PBS        -4.8426 0.467 24 -10.365  <.0001
##
## Results are given on the log2 (not the response) scale.
```

```
p.adjust(a3E_tRPE65$contrasts$p.value[c(1,3,4,6,7,9,10,12)], method="fdr")
```

```
## [1] 9.096275e-01 1.275374e-07 1.692766e-04 5.023599e-03 2.539010e-01
## [6] 2.279849e-11 1.510426e-03 9.723334e-10
```

```
a3E_tTYR <- test(emmeans(a3E_TYR, pairwise~Chemical|Time), adjust="none")
a3E_tTYR
```

```
## $emmeans
## Time = 24:
##  Chemical emmean    SE df t.ratio p.value
##  2DG       -2.64 0.295 24  -8.950  <.0001
##  2DG+FGF   -3.30 0.295 24 -11.192  <.0001
##  FGF2      -2.77 0.295 24  -9.395  <.0001
##  PBS        0.00 0.295 24   0.000  1.0000
##
## Time = 48:
##  Chemical emmean    SE df t.ratio p.value
##  2DG       -2.64 0.295 24  -8.963  <.0001
##  2DG+FGF   -3.36 0.295 24 -11.396  <.0001
##  FGF2      -2.88 0.295 24  -9.763  <.0001
##  PBS        1.17 0.295 24   3.978  0.0006
##
## Results are given on the log2 (not the response) scale.
##
## $contrasts
## Time = 24:
##  contrast         estimate    SE df t.ratio p.value
##  2DG - (2DG+FGF)     0.661 0.417 24   1.586  0.1258
##  2DG - FGF2          0.131 0.417 24   0.315  0.7554
##  2DG - PBS          -2.636 0.417 24  -6.328  <.0001
##  (2DG+FGF) - FGF2   -0.529 0.417 24  -1.271  0.2160
##  (2DG+FGF) - PBS    -3.297 0.417 24  -7.914  <.0001
##  FGF2 - PBS         -2.767 0.417 24  -6.643  <.0001
##
## Time = 48:
```

```
##  contrast         estimate    SE df t.ratio p.value
##  2DG - (2DG+FGF)     0.717 0.417 24   1.721  0.0982
##  2DG - FGF2          0.236 0.417 24   0.566  0.5765
##  2DG - PBS          -3.812 0.417 24  -9.151  <.0001
##  (2DG+FGF) - FGF2   -0.481 0.417 24  -1.154  0.2597
##  (2DG+FGF) - PBS    -4.528 0.417 24 -10.871  <.0001
##  FGF2 - PBS         -4.048 0.417 24  -9.717  <.0001
##
## Results are given on the log2 (not the response) scale.
```

```
p.adjust(a3E_tTYR$contrasts$p.value[c(1,3,4,6,7,9,10,12)], method="fdr")
```

```
## [1] 1.677966e-01 3.056933e-06 2.468076e-01 1.910047e-06 1.571205e-01
## [6] 1.080753e-08 2.597051e-01 6.871202e-09
```

```
a3E_tOTX2 <- test(emmeans(a3E_OTX2, pairwise~Chemical|Time), adjust="none")
a3E_tOTX2
```

```
## $emmeans
## Time = 24:
##  Chemical emmean    SE df t.ratio p.value
##  2DG      -1.336 0.134 24  -9.971  <.0001
##  2DG+FGF  -1.696 0.134 24 -12.654  <.0001
##  FGF2     -1.826 0.134 24 -13.624  <.0001
##  PBS       0.000 0.134 24   0.000  1.0000
##
## Time = 48:
##  Chemical emmean    SE df t.ratio p.value
##  2DG      -2.204 0.134 24 -16.450  <.0001
##  2DG+FGF  -2.617 0.134 24 -19.529  <.0001
##  FGF2     -0.942 0.134 24  -7.032  <.0001
##  PBS       0.320 0.134 24   2.391  0.0250
##
## Results are given on the log2 (not the response) scale.
##
## $contrasts
## Time = 24:
##  contrast         estimate    SE df t.ratio p.value
##  2DG - (2DG+FGF)     0.359 0.189 24   1.897  0.0699
##  2DG - FGF2          0.489 0.189 24   2.583  0.0163
##  2DG - PBS          -1.336 0.189 24  -7.050  <.0001
##  (2DG+FGF) - FGF2    0.130 0.189 24   0.686  0.4993
##  (2DG+FGF) - PBS    -1.696 0.189 24  -8.948  <.0001
##  FGF2 - PBS         -1.826 0.189 24  -9.634  <.0001
##
## Time = 48:
##  contrast         estimate    SE df t.ratio p.value
##  2DG - (2DG+FGF)     0.412 0.189 24   2.177  0.0396
##  2DG - FGF2         -1.262 0.189 24  -6.659  <.0001
##  2DG - PBS          -2.525 0.189 24 -13.323  <.0001
##  (2DG+FGF) - FGF2   -1.674 0.189 24  -8.836  <.0001
##  (2DG+FGF) - PBS    -2.937 0.189 24 -15.499  <.0001
##  FGF2 - PBS         -1.263 0.189 24  -6.663  <.0001
```

```
##
## Results are given on the log2 (not the response) scale.
```

```r
p.adjust(a3E_tOTX2$contrasts$p.value[c(1,3,4,6,7,9,10,12)], method="fdr")
```

```
## [1] 7.989750e-02 5.475827e-07 4.992961e-01 4.055947e-09 5.275929e-02
## [6] 1.114628e-11 1.386258e-08 1.093073e-06
```

##### Treatment Comparisons (Figure 3F)

```r
a3F_HK1 <- aov(log2(HK1)~Chemical*Time, data=d3F)
a3F_ENO1 <- aov(log2(ENO1)~Chemical*Time, data=d3F)
a3F_LDHA <- aov(log2(LDHA)~Chemical*Time, data=d3F)
a3F_LDHB <- aov(log2(LDHB)~Chemical*Time, data=d3F)

plot(a3F_HK1,which=1:2)
```

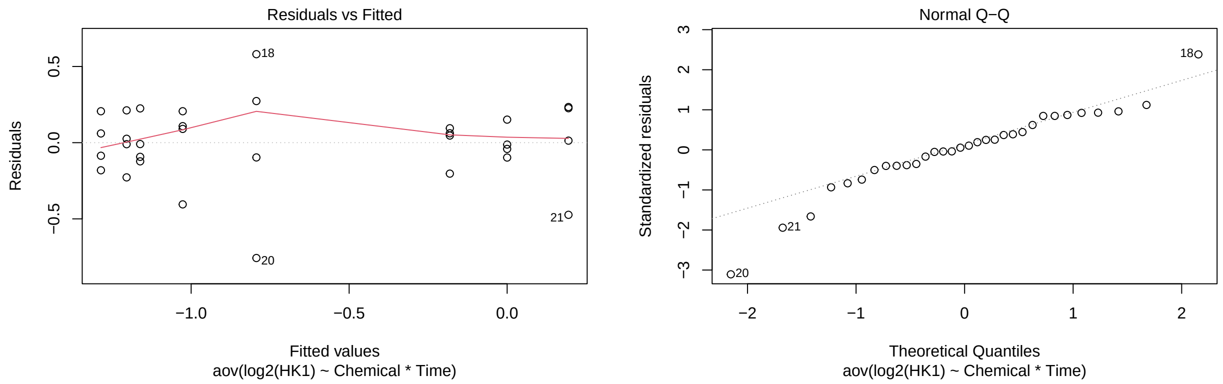

```r
plot(a3F_ENO1,which=1:2)
```

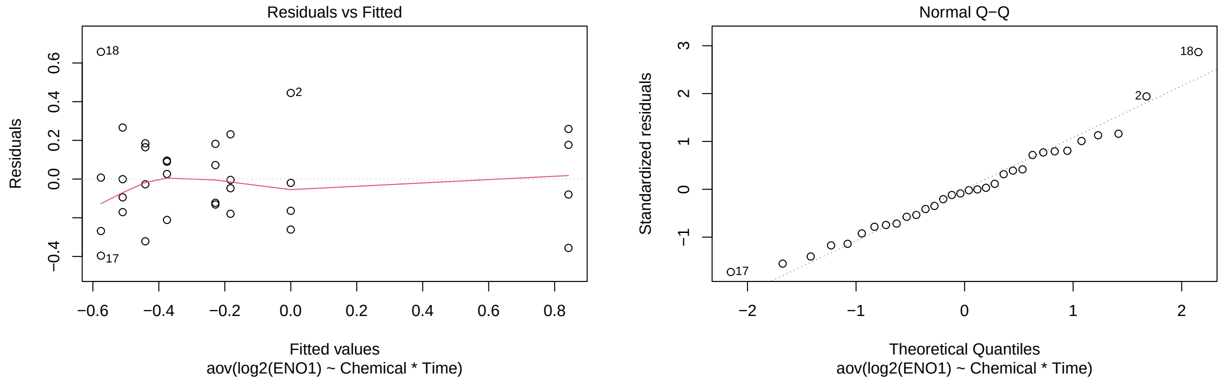

```r
plot(a3F_LDHA,which=1:2)
```

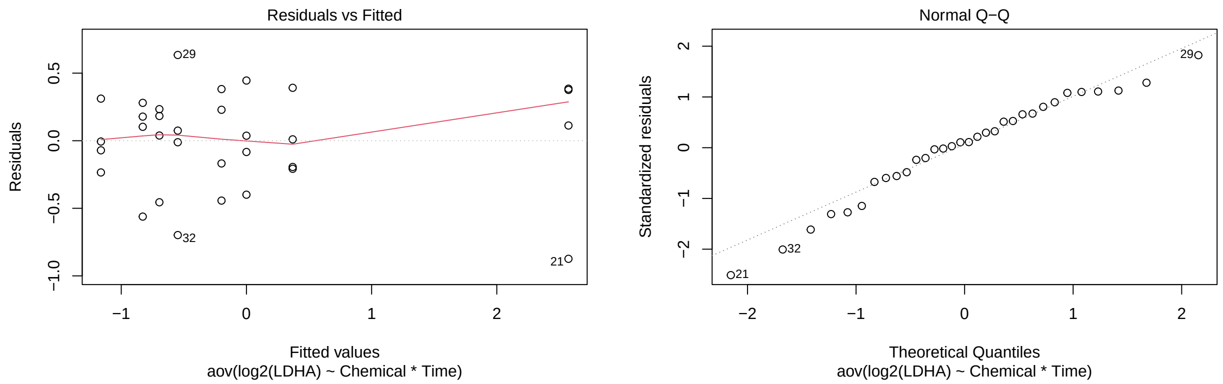

```r
plot(a3F_LDHB,which=1:2)
```

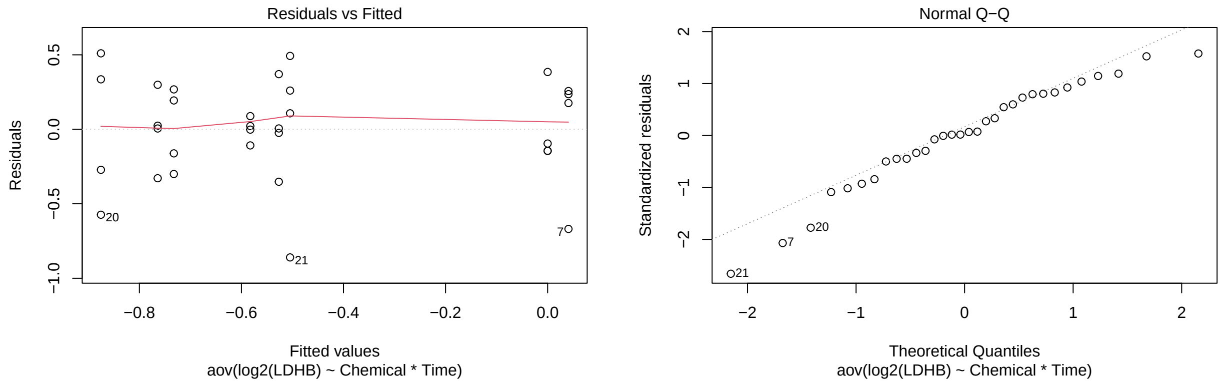

```r
a3F_tHK1 <- test(emmeans(a3F_HK1, pairwise~Chemical|Time), adjust="none")
a3F_tHK1
```

```
## $emmeans
## Time = 24:
##  Chemical emmean    SE df t.ratio p.value
##  2DG      -1.286 0.141 24  -9.138  <.0001
##  2DG+FGF  -1.027 0.141 24  -7.298  <.0001
##  FGF2     -0.181 0.141 24  -1.289  0.2097
##  PBS       0.000 0.141 24   0.000  1.0000
##
## Time = 48:
##  Chemical emmean    SE df t.ratio p.value
##  2DG      -1.161 0.141 24  -8.255  <.0001
##  2DG+FGF  -1.204 0.141 24  -8.560  <.0001
##  FGF2      0.194 0.141 24   1.380  0.1805
##  PBS      -0.794 0.141 24  -5.641  <.0001
##
## Results are given on the log2 (not the response) scale.
##
## $contrasts
## Time = 24:
```

```
##  contrast         estimate    SE df t.ratio p.value
##  2DG - (2DG+FGF)   -0.2588 0.199 24  -1.301  0.2056
##  2DG - FGF2        -1.1042 0.199 24  -5.550  <.0001
##  2DG - PBS         -1.2855 0.199 24  -6.461  <.0001
##  (2DG+FGF) - FGF2  -0.8454 0.199 24  -4.249  0.0003
##  (2DG+FGF) - PBS   -1.0267 0.199 24  -5.161  <.0001
##  FGF2 - PBS        -0.1813 0.199 24  -0.911  0.3711
##
## Time = 48:
##  contrast         estimate    SE df t.ratio p.value
##  2DG - (2DG+FGF)    0.0429 0.199 24   0.216  0.8312
##  2DG - FGF2        -1.3554 0.199 24  -6.813  <.0001
##  2DG - PBS         -0.3678 0.199 24  -1.848  0.0769
##  (2DG+FGF) - FGF2  -1.3983 0.199 24  -7.028  <.0001
##  (2DG+FGF) - PBS   -0.4107 0.199 24  -2.064  0.0500
##  FGF2 - PBS         0.9876 0.199 24   4.964  <.0001
##
## Results are given on the log2 (not the response) scale.
```

```
p.adjust(a3F_tHK1$contrasts$p.value[c(1,3,4,6,7,9,10,12)], method="fdr")
```

```
## [1] 2.741834e-01 4.432184e-06 5.606322e-04 4.241518e-01 8.311653e-01
## [6] 1.230283e-01 2.308085e-06 1.214323e-04
```

```
a3F_tENO1 <- test(emmeans(a3F_ENO1, pairwise~Chemical|Time), adjust="none")
a3F_tENO1
```

```
## $emmeans
## Time = 24:
##  Chemical emmean    SE df t.ratio p.value
##  2DG      -0.441 0.132 24  -3.331  0.0028
##  2DG+FGF  -0.510 0.132 24  -3.850  0.0008
##  FGF2     -0.375 0.132 24  -2.836  0.0091
##  PBS       0.000 0.132 24   0.000  1.0000
##
## Time = 48:
##  Chemical emmean    SE df t.ratio p.value
##  2DG      -0.228 0.132 24  -1.726  0.0972
##  2DG+FGF  -0.183 0.132 24  -1.380  0.1802
##  FGF2      0.842 0.132 24   6.361  <.0001
##  PBS      -0.576 0.132 24  -4.348  0.0002
##
## Results are given on the log2 (not the response) scale.
##
## $contrasts
## Time = 24:
##  contrast         estimate    SE df t.ratio p.value
##  2DG - (2DG+FGF)    0.0687 0.187 24   0.367  0.7169
##  2DG - FGF2        -0.0656 0.187 24  -0.350  0.7293
##  2DG - PBS         -0.4409 0.187 24  -2.355  0.0270
##  (2DG+FGF) - FGF2  -0.1342 0.187 24  -0.717  0.4803
##  (2DG+FGF) - PBS   -0.5096 0.187 24  -2.722  0.0119
##  FGF2 - PBS        -0.3754 0.187 24  -2.005  0.0563
```

```
##
## Time = 48:
##  contrast          estimate    SE df t.ratio p.value
##  2DG - (2DG+FGF)    -0.0457 0.187 24  -0.244  0.8092
##  2DG - FGF2         -1.0704 0.187 24  -5.718  <.0001
##  2DG - PBS           0.3471 0.187 24   1.854  0.0760
##  (2DG+FGF) - FGF2  -1.0247 0.187 24  -5.474  <.0001
##  (2DG+FGF) - PBS     0.3928 0.187 24   2.099  0.0466
##  FGF2 - PBS          1.4175 0.187 24   7.572  <.0001
##
## Results are given on the log2 (not the response) scale.
```

```
p.adjust(a3F_tENO1$contrasts$p.value[c(1,3,4,6,7,9,10,12)], method="fdr")
```

```
## [1] 8.091615e-01 7.202979e-02 6.403415e-01 1.126773e-01 8.091615e-01
## [6] 1.216284e-01 5.034697e-05 6.582629e-07
```

```
a3F_tLDHA <- test(emmeans(a3F_LDHA, pairwise~Chemical|Time), adjust="none")
a3F_tLDHA
```

```
## $emmeans
## Time = 24:
##  Chemical emmean    SE df t.ratio p.value
##  2DG      -0.828 0.201 24  -4.123  0.0004
##  2DG+FGF  -1.163 0.201 24  -5.786  <.0001
##  FGF2     -0.199 0.201 24  -0.991  0.3316
##  PBS       0.000 0.201 24   0.000  1.0000
##
## Time = 48:
##  Chemical emmean    SE df t.ratio p.value
##  2DG      -0.696 0.201 24  -3.462  0.0020
##  2DG+FGF  -0.549 0.201 24  -2.730  0.0117
##  FGF2      2.573 0.201 24  12.805  <.0001
##  PBS       0.369 0.201 24   1.839  0.0783
##
## Results are given on the log2 (not the response) scale.
##
## $contrasts
## Time = 24:
##  contrast          estimate    SE df t.ratio p.value
##  2DG - (2DG+FGF)      0.334 0.284 24   1.176  0.2511
##  2DG - FGF2          -0.629 0.284 24  -2.215  0.0365
##  2DG - PBS           -0.828 0.284 24  -2.915  0.0076
##  (2DG+FGF) - FGF2    -0.963 0.284 24  -3.391  0.0024
##  (2DG+FGF) - PBS     -1.163 0.284 24  -4.091  0.0004
##  FGF2 - PBS          -0.199 0.284 24  -0.701  0.4902
##
## Time = 48:
##  contrast          estimate    SE df t.ratio p.value
##  2DG - (2DG+FGF)     -0.147 0.284 24  -0.517  0.6097
##  2DG - FGF2          -3.268 0.284 24 -11.502  <.0001
##  2DG - PBS           -1.065 0.284 24  -3.748  0.0010
##  (2DG+FGF) - FGF2    -3.121 0.284 24 -10.985  <.0001
```

```
##  (2DG+FGF) - PBS    -0.918 0.284 24  -3.231  0.0036
##  FGF2 - PBS          2.203 0.284 24   7.754  <.0001
##
## Results are given on the log2 (not the response) scale.
```

```
p.adjust(a3F_tLDHA$contrasts$p.value[c(1,3,4,6,7,9,10,12)], method="fdr")
```

```
## [1] 3.347569e-01 1.213106e-02 4.823482e-03 5.602475e-01 6.097095e-01
## [6] 2.647672e-03 6.087243e-10 2.183768e-07
```

```
a3F_tLDHB <- test(emmeans(a3F_LDHB, pairwise~Chemical|Time), adjust="none")
a3F_tLDHB
```

```
## $emmeans
## Time = 24:
##  Chemical  emmean    SE df t.ratio p.value
##  2DG      -0.7324 0.187 24  -3.926  0.0006
##  2DG+FGF  -0.7641 0.187 24  -4.096  0.0004
##  FGF2      0.0408 0.187 24   0.219  0.8286
##  PBS       0.0000 0.187 24   0.000  1.0000
##
## Time = 48:
##  Chemical  emmean    SE df t.ratio p.value
##  2DG      -0.5827 0.187 24  -3.124  0.0046
##  2DG+FGF  -0.5267 0.187 24  -2.824  0.0094
##  FGF2     -0.5047 0.187 24  -2.706  0.0123
##  PBS      -0.8753 0.187 24  -4.693  0.0001
##
## Results are given on the log2 (not the response) scale.
##
## $contrasts
## Time = 24:
##  contrast          estimate    SE df t.ratio p.value
##  2DG - (2DG+FGF)     0.0317 0.264 24   0.120  0.9055
##  2DG - FGF2         -0.7733 0.264 24  -2.931  0.0073
##  2DG - PBS          -0.7324 0.264 24  -2.776  0.0105
##  (2DG+FGF) - FGF2  -0.8049 0.264 24  -3.051  0.0055
##  (2DG+FGF) - PBS   -0.7641 0.264 24  -2.896  0.0079
##  FGF2 - PBS          0.0408 0.264 24   0.155  0.8783
##
## Time = 48:
##  contrast          estimate    SE df t.ratio p.value
##  2DG - (2DG+FGF)    -0.0559 0.264 24  -0.212  0.8339
##  2DG - FGF2         -0.0780 0.264 24  -0.296  0.7701
##  2DG - PBS           0.2927 0.264 24   1.109  0.2782
##  (2DG+FGF) - FGF2  -0.0220 0.264 24  -0.083  0.9342
##  (2DG+FGF) - PBS    0.3486 0.264 24   1.321  0.1988
##  FGF2 - PBS          0.3706 0.264 24   1.405  0.1729
##
## Results are given on the log2 (not the response) scale.
```

```r
p.adjust(a3F_tLDHB$contrasts$p.value[c(1,3,4,6,7,9,10,12)], method="fdr")
```

```
## [1] 0.93417479 0.04194626 0.04194626 0.93417479 0.93417479 0.55648114 0.93417479
## [8] 0.46094209
```

##### Treatment Comparisons (Figure 3G)

```r
a3G_E2F1 <- aov(log2(E2F1)~Chemical*Time, data=d3G)
a3G_PCNA <- aov(log2(PCNA)~Chemical*Time, data=d3G)

plot(a3G_E2F1,which=1:2)
```

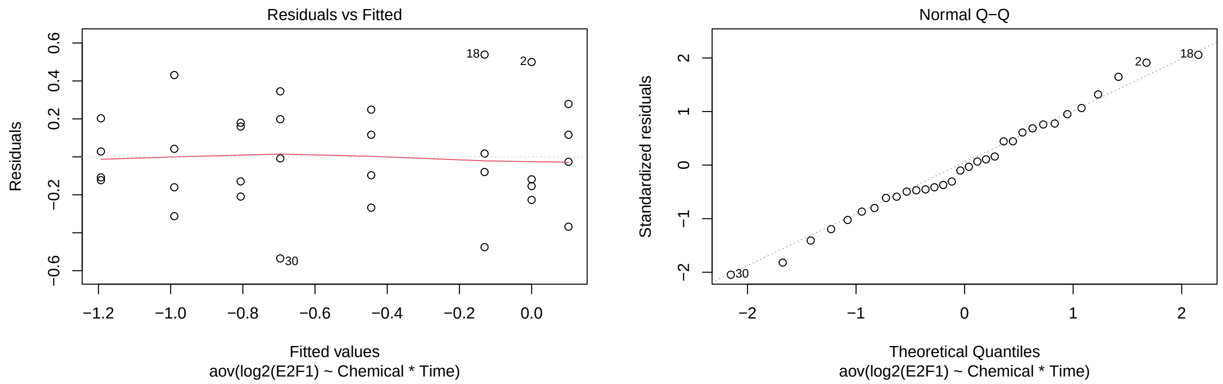

```r
plot(a3G_PCNA,which=1:2)
```

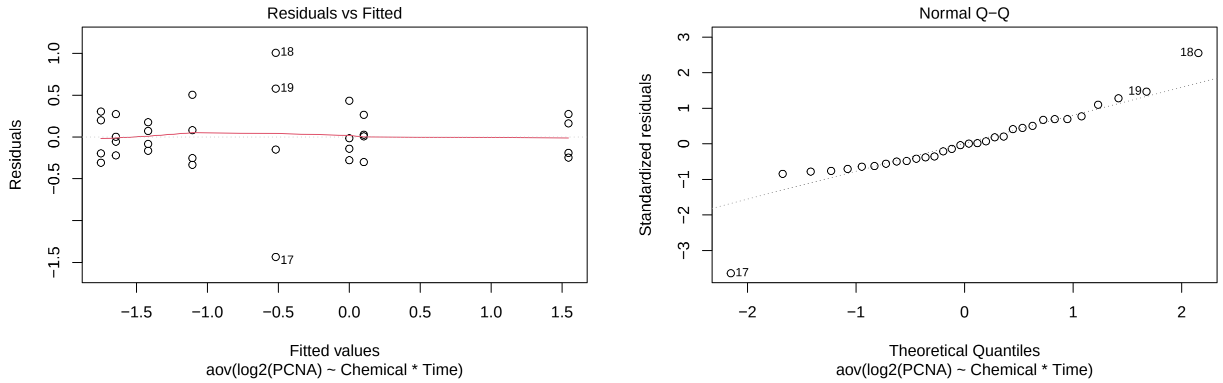

```r
a3G_tE2F1 <- test(emmeans(a3G_E2F1, pairwise~Chemical|Time), adjust="none")
a3G_tE2F1
```

```
## $emmeans
## Time = 24:
##  Chemical emmean    SE df t.ratio p.value
```

```
##  2DG      -0.806 0.151 24  -5.336  <.0001
##  2DG+FGF  -0.990 0.151 24  -6.554  <.0001
##  FGF2     -0.444 0.151 24  -2.942  0.0071
##  PBS       0.000 0.151 24   0.000  1.0000
##
## Time = 48:
##  Chemical emmean    SE df t.ratio p.value
##  2DG      -1.193 0.151 24  -7.899  <.0001
##  2DG+FGF  -0.696 0.151 24  -4.610  0.0001
##  FGF2      0.102 0.151 24   0.676  0.5055
##  PBS      -0.130 0.151 24  -0.863  0.3966
##
## Results are given on the log2 (not the response) scale.
##
## $contrasts
## Time = 24:
##  contrast          estimate    SE df t.ratio p.value
##  2DG - (2DG+FGF)      0.184 0.214 24   0.861  0.3978
##  2DG - FGF2          -0.362 0.214 24  -1.693  0.1034
##  2DG - PBS           -0.806 0.214 24  -3.773  0.0009
##  (2DG+FGF) - FGF2    -0.546 0.214 24  -2.554  0.0174
##  (2DG+FGF) - PBS     -0.990 0.214 24  -4.634  0.0001
##  FGF2 - PBS          -0.444 0.214 24  -2.080  0.0484
##
## Time = 48:
##  contrast          estimate    SE df t.ratio p.value
##  2DG - (2DG+FGF)     -0.497 0.214 24  -2.326  0.0288
##  2DG - FGF2          -1.295 0.214 24  -6.064  <.0001
##  2DG - PBS           -1.063 0.214 24  -4.975  <.0001
##  (2DG+FGF) - FGF2    -0.798 0.214 24  -3.738  0.0010
##  (2DG+FGF) - PBS     -0.566 0.214 24  -2.650  0.0140
##  FGF2 - PBS           0.232 0.214 24   1.088  0.2873
##
## Results are given on the log2 (not the response) scale.
```

```
p.adjust(a3G_tE2F1$contrasts$p.value[c(1,3,4,6,7,9,10,12)], method="fdr")
```

```
## [1] 0.3978466072 0.0027180318 0.0348349604 0.0644869500 0.0460779248
## [6] 0.0003539377 0.0027180318 0.3283423181
```

```
a3G_tPCNA <- test(emmeans(a3G_PCNA, pairwise~Chemical|Time), adjust="none")
a3G_tPCNA
```

```
## $emmeans
## Time = 24:
##  Chemical emmean    SE df t.ratio p.value
##  2DG      -1.751 0.228 24  -7.692  <.0001
##  2DG+FGF  -1.645 0.228 24  -7.229  <.0001
##  FGF2      0.102 0.228 24   0.450  0.6567
##  PBS       0.000 0.228 24   0.000  1.0000
##
## Time = 48:
##  Chemical emmean    SE df t.ratio p.value
```

```
##  2DG      -1.418 0.228 24  -6.231  <.0001
##  2DG+FGF  -1.106 0.228 24  -4.859  0.0001
##  FGF2      1.545 0.228 24   6.788  <.0001
##  PBS      -0.518 0.228 24  -2.277  0.0320
##
## Results are given on the log2 (not the response) scale.
##
## $contrasts
## Time = 24:
##  contrast          estimate    SE df t.ratio p.value
##  2DG - (2DG+FGF)     -0.106 0.322 24  -0.328  0.7458
##  2DG - FGF2          -1.853 0.322 24  -5.758  <.0001
##  2DG - PBS           -1.751 0.322 24  -5.439  <.0001
##  (2DG+FGF) - FGF2    -1.748 0.322 24  -5.430  <.0001
##  (2DG+FGF) - PBS     -1.645 0.322 24  -5.111  <.0001
##  FGF2 - PBS           0.102 0.322 24   0.318  0.7530
##
## Time = 48:
##  contrast          estimate    SE df t.ratio p.value
##  2DG - (2DG+FGF)     -0.312 0.322 24  -0.970  0.3419
##  2DG - FGF2          -2.963 0.322 24  -9.205  <.0001
##  2DG - PBS           -0.900 0.322 24  -2.795  0.0100
##  (2DG+FGF) - FGF2    -2.651 0.322 24  -8.236  <.0001
##  (2DG+FGF) - PBS     -0.588 0.322 24  -1.826  0.0804
##  FGF2 - PBS           2.063 0.322 24   6.410  <.0001
##
## Results are given on the log2 (not the response) scale.
```

```r
p.adjust(a3G_tPCNA$contrasts$p.value[c(1,3,4,6,7,9,10,12)], method="fdr")
```

```
## [1] 7.530313e-01 2.812003e-05 2.812003e-05 7.530313e-01 4.558465e-01
## [6] 1.605679e-02 1.504044e-07 5.016836e-06
```

##### Treatment Comparisons (Figure S5F)

```r
a5SF_HK2 <- aov(log2(HK2)~Chemical*Time, data=d5SF)
a5SF_TPI1 <- aov(log2(TPI1)~Chemical*Time, data=d5SF)
a5SF_PGAM1 <- aov(log2(PGAM1)~Chemical*Time, data=d5SF)
a5SF_PKLR <- aov(log2(PKLR)~Chemical*Time, data=d5SF)

plot(a5SF_HK2,which=1:2)
```

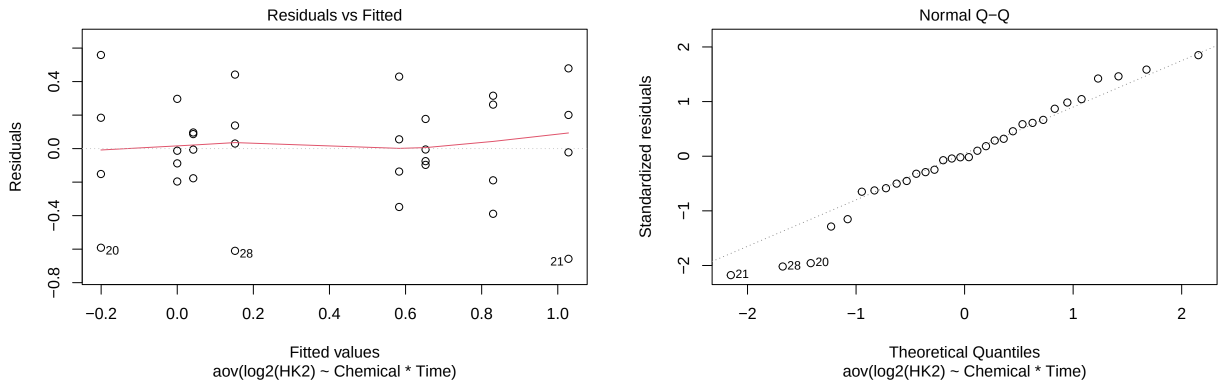

```
plot(a5SF_TPI1,which=1:2)
```

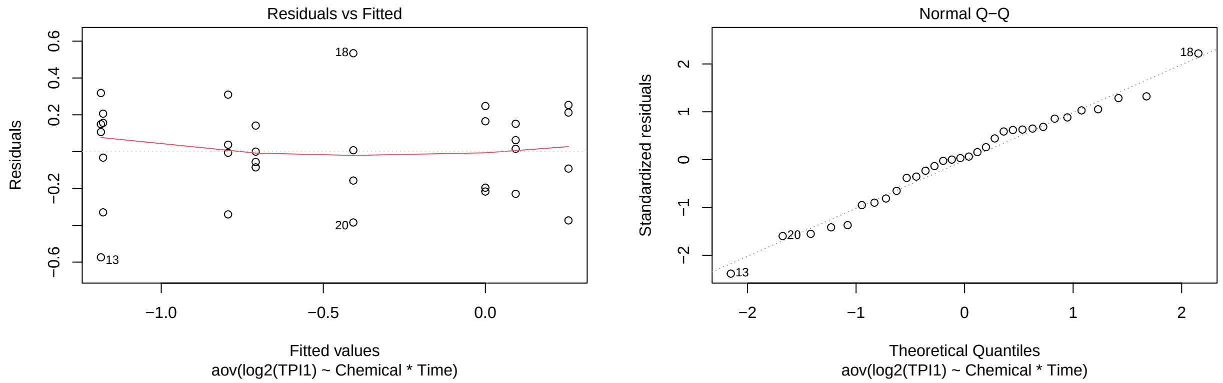

```
plot(a5SF_PGAM1,which=1:2)
```

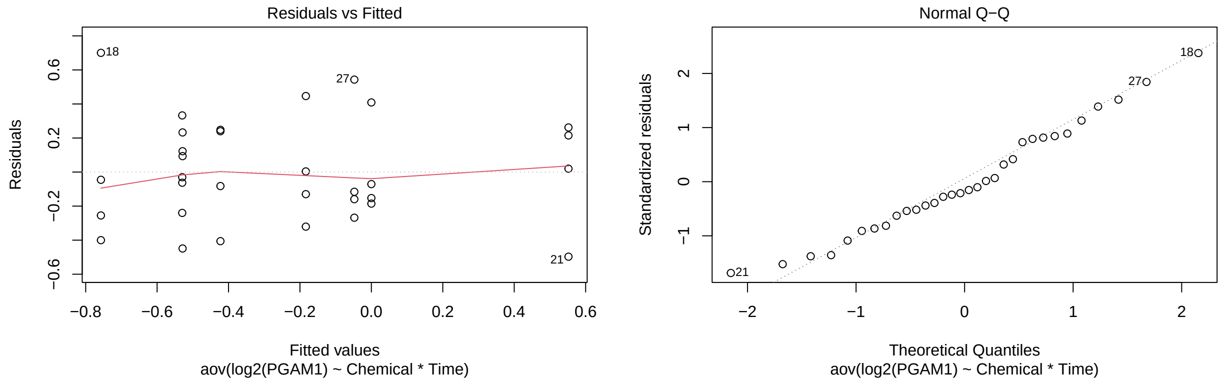

```
plot(a5SF_PKLR,which=1:2)
```

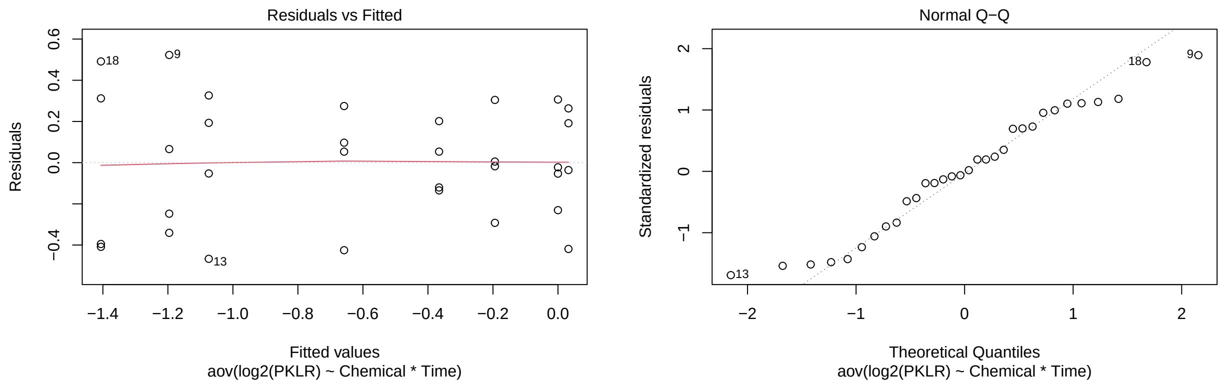

```
a5SF_tHK2 <- test(emmeans(a5SF_HK2, pairwise~Chemical|Time), adjust="none")
a5SF_tHK2
```

```
## $emmeans
## Time = 24:
##  Chemical emmean    SE df t.ratio p.value
##  2DG       0.583 0.174 24   3.342  0.0027
##  2DG+FGF   0.653 0.174 24   3.741  0.0010
##  FGF2      0.830 0.174 24   4.758  0.0001
##  PBS       0.000 0.174 24   0.000  1.0000
##
## Time = 48:
##  Chemical emmean    SE df t.ratio p.value
##  2DG       0.152 0.174 24   0.870  0.3928
##  2DG+FGF   0.042 0.174 24   0.241  0.8117
##  FGF2      1.028 0.174 24   5.893  <.0001
##  PBS      -0.201 0.174 24  -1.150  0.2616
##
## Results are given on the log2 (not the response) scale.
##
## $contrasts
## Time = 24:
##  contrast         estimate    SE df t.ratio p.value
##  2DG - (2DG+FGF)   -0.0695 0.247 24  -0.282  0.7807
##  2DG - FGF2        -0.2469 0.247 24  -1.001  0.3269
##  2DG - PBS          0.5831 0.247 24   2.363  0.0265
##  (2DG+FGF) - FGF2  -0.1775 0.247 24  -0.719  0.4789
##  (2DG+FGF) - PBS    0.6526 0.247 24   2.645  0.0142
##  FGF2 - PBS         0.8300 0.247 24   3.364  0.0026
##
## Time = 48:
##  contrast         estimate    SE df t.ratio p.value
##  2DG - (2DG+FGF)    0.1098 0.247 24   0.445  0.6603
##  2DG - FGF2        -0.8763 0.247 24  -3.552  0.0016
##  2DG - PBS          0.3524 0.247 24   1.428  0.1661
##  (2DG+FGF) - FGF2  -0.9861 0.247 24  -3.997  0.0005
##  (2DG+FGF) - PBS    0.2426 0.247 24   0.983  0.3352
##  FGF2 - PBS         1.2287 0.247 24   4.980  <.0001
##
```

```
## Results are given on the log2 (not the response) scale.
```

```r
p.adjust(a5SF_tHK2$contrasts$p.value[c(1,3,4,6,7,9,10,12)], method="fdr")
```

```
## [1] 0.7807148217 0.0530905117 0.6385181131 0.0068627585 0.7546161184
## [6] 0.2657026853 0.0021248021 0.0003497017
```

```r
a5SF_tTPI1 <- test(emmeans(a5SF_TPI1, pairwise~Chemical|Time), adjust="none")
a5SF_tTPI1
```

```
## $emmeans
## Time = 24:
##  Chemical  emmean    SE df t.ratio p.value
##  2DG      -1.1794 0.139 24  -8.486  <.0001
##  2DG+FGF  -1.1861 0.139 24  -8.534  <.0001
##  FGF2      0.0936 0.139 24   0.673  0.5071
##  PBS       0.0000 0.139 24   0.000  1.0000
##
## Time = 48:
##  Chemical  emmean    SE df t.ratio p.value
##  2DG      -0.7085 0.139 24  -5.098  <.0001
##  2DG+FGF  -0.7937 0.139 24  -5.711  <.0001
##  FGF2      0.2564 0.139 24   1.845  0.0774
##  PBS      -0.4073 0.139 24  -2.931  0.0073
##
## Results are given on the log2 (not the response) scale.
##
## $contrasts
## Time = 24:
##  contrast          estimate    SE df t.ratio p.value
##  2DG - (2DG+FGF)    0.00673 0.197 24   0.034  0.9730
##  2DG - FGF2        -1.27300 0.197 24  -6.476  <.0001
##  2DG - PBS         -1.17940 0.197 24  -6.000  <.0001
##  (2DG+FGF) - FGF2 -1.27973 0.197 24  -6.511  <.0001
##  (2DG+FGF) - PBS  -1.18613 0.197 24  -6.034  <.0001
##  FGF2 - PBS         0.09360 0.197 24   0.476  0.6383
##
## Time = 48:
##  contrast          estimate    SE df t.ratio p.value
##  2DG - (2DG+FGF)    0.08525 0.197 24   0.434  0.6684
##  2DG - FGF2        -0.96494 0.197 24  -4.909  0.0001
##  2DG - PBS         -0.30115 0.197 24  -1.532  0.1386
##  (2DG+FGF) - FGF2 -1.05019 0.197 24  -5.343  <.0001
##  (2DG+FGF) - PBS  -0.38640 0.197 24  -1.966  0.0610
##  FGF2 - PBS         0.66379 0.197 24   3.377  0.0025
##
## Results are given on the log2 (not the response) scale.
```

```r
p.adjust(a5SF_tTPI1$contrasts$p.value[c(1,3,4,6,7,9,10,12)], method="fdr")
```

```
## [1] 9.729617e-01 1.362131e-05 7.874096e-06 7.638649e-01 7.638649e-01
## [6] 2.217066e-01 4.663110e-05 4.988606e-03
```

```r
a5SF_tPGAM1 <- test(emmeans(a5SF_PGAM1, pairwise~Chemical|Time), adjust="none")
a5SF_tPGAM1
```

```
## $emmeans
## Time = 24:
##  Chemical  emmean   SE df t.ratio p.value
##  2DG      -0.4227 0.17 24  -2.485  0.0203
##  2DG+FGF  -0.5296 0.17 24  -3.114  0.0047
##  FGF2     -0.1835 0.17 24  -1.079  0.2914
##  PBS       0.0000 0.17 24   0.000  1.0000
##
## Time = 48:
##  Chemical  emmean   SE df t.ratio p.value
##  2DG      -0.0477 0.17 24  -0.280  0.7816
##  2DG+FGF  -0.5287 0.17 24  -3.108  0.0048
##  FGF2      0.5519 0.17 24   3.245  0.0034
##  PBS      -0.7573 0.17 24  -4.453  0.0002
##
## Results are given on the log2 (not the response) scale.
##
## $contrasts
## Time = 24:
##  contrast         estimate    SE df t.ratio p.value
##  2DG - (2DG+FGF)     0.107 0.241 24   0.445  0.6605
##  2DG - FGF2         -0.239 0.241 24  -0.994  0.3300
##  2DG - PBS          -0.423 0.241 24  -1.757  0.0916
##  (2DG+FGF) - FGF2   -0.346 0.241 24  -1.439  0.1630
##  (2DG+FGF) - PBS    -0.530 0.241 24  -2.202  0.0375
##  FGF2 - PBS         -0.183 0.241 24  -0.763  0.4530
##
## Time = 48:
##  contrast         estimate    SE df t.ratio p.value
##  2DG - (2DG+FGF)     0.481 0.241 24   2.000  0.0570
##  2DG - FGF2         -0.600 0.241 24  -2.493  0.0200
##  2DG - PBS           0.710 0.241 24   2.950  0.0070
##  (2DG+FGF) - FGF2   -1.081 0.241 24  -4.493  0.0002
##  (2DG+FGF) - PBS     0.229 0.241 24   0.951  0.3513
##  FGF2 - PBS          1.309 0.241 24   5.443  <.0001
##
## Results are given on the log2 (not the response) scale.
```

```r
p.adjust(a5SF_tPGAM1$contrasts$p.value[c(1,3,4,6,7,9,10,12)], method="fdr")
```

```
## [1] 0.6605114926 0.1466149148 0.2173999110 0.5176897594 0.1139568035
## [6] 0.0186160155 0.0006036999 0.0001087183
```

```r
a5SF_tPKLR <- test(emmeans(a5SF_PKLR, pairwise~Chemical|Time), adjust="none")
a5SF_tPKLR
```

```
## $emmeans
## Time = 24:
##  Chemical  emmean    SE df t.ratio p.value
```

```
##  2DG      -1.1959 0.159 24  -7.504  <.0001
##  2DG+FGF  -1.0742 0.159 24  -6.741  <.0001
##  FGF2     -0.1940 0.159 24  -1.217  0.2353
##  PBS       0.0000 0.159 24   0.000  1.0000
##
## Time = 48:
##  Chemical  emmean    SE df t.ratio p.value
##  2DG      -0.3657 0.159 24  -2.295  0.0308
##  2DG+FGF  -0.6579 0.159 24  -4.128  0.0004
##  FGF2      0.0323 0.159 24   0.203  0.8410
##  PBS      -1.4060 0.159 24  -8.823  <.0001
##
## Results are given on the log2 (not the response) scale.
##
## $contrasts
## Time = 24:
##  contrast         estimate    SE df t.ratio p.value
##  2DG - (2DG+FGF)    -0.122 0.225 24  -0.540  0.5942
##  2DG - FGF2         -1.002 0.225 24  -4.446  0.0002
##  2DG - PBS          -1.196 0.225 24  -5.306  <.0001
##  (2DG+FGF) - FGF2   -0.880 0.225 24  -3.906  0.0007
##  (2DG+FGF) - PBS    -1.074 0.225 24  -4.766  0.0001
##  FGF2 - PBS         -0.194 0.225 24  -0.861  0.3979
##
## Time = 48:
##  contrast         estimate    SE df t.ratio p.value
##  2DG - (2DG+FGF)     0.292 0.225 24   1.296  0.2072
##  2DG - FGF2         -0.398 0.225 24  -1.766  0.0901
##  2DG - PBS           1.040 0.225 24   4.616  0.0001
##  (2DG+FGF) - FGF2   -0.690 0.225 24  -3.063  0.0053
##  (2DG+FGF) - PBS     0.748 0.225 24   3.319  0.0029
##  FGF2 - PBS          1.438 0.225 24   6.382  <.0001
##
## Results are given on the log2 (not the response) scale.
```

```r
p.adjust(a5SF_tPKLR$contrasts$p.value[c(1,3,4,6,7,9,10,12)], method="fdr")
```

```
## [1] 5.941914e-01 7.668667e-05 1.337401e-03 4.547122e-01 2.762295e-01
## [6] 2.941973e-04 8.552993e-03 1.073797e-05
```

### Experiment 4 (Figures 4C, 5F, S6E)

The responses are relative expressions of several genes. Same treatment setup as Experiment 3, except chemical DGA instead of 2DG. We have 8 treatments and 4 replications/treatment.

Comparisons for Figure 4C, 5F, S6E: PBS vs FGF2; PBS vs DCA; FGF2 vs FGF2+DCA, and DCA vs FGF2+DCA at each time point for each gene

#### Data and Visualization

##### Plots to Compare the Four Treatments at each Time Point

```
d4C <- read.xlsx(xlsxFile = "MetabolismPaperRawData_Byran_working.xlsx",
                 colNames = TRUE, sheet="Fig 4_C")
glimpse(d4C)
```

```
## Rows: 32
## Columns: 9
## $ Chemical   <chr> "PBS", "PBS", "PBS", "PBS", "FGF2", "FGF2", "FGF2", "FGF2",~
## $ Sample     <dbl> 1, 2, 3, 4, 1, 2, 3, 4, 1, 2, 3, 4, 1, 2, 3, 4, 1, 2, 3, 4,~
## $ `Time.(h)` <dbl> 24, 24, 24, 24, 24, 24, 24, 24, 24, 24, 24, 24, 24, 24, 24,~
## $ SOX2       <dbl> 0.6683726, 1.7839131, 1.0073875, 0.8325515, 2.0972087, 1.74~
## $ SIX6       <dbl> 1.2109944, 1.2081256, 0.5674564, 1.2045180, 9.4502289, 10.4~
## $ PAX6       <dbl> 0.8869278, 0.8325113, 1.1858234, 1.1420932, 3.2961737, 4.79~
## $ RPE65      <dbl> 1.0168114, 1.5935640, 0.6206805, 0.9943105, 0.9160471, 0.73~
## $ TYR        <dbl> 1.1903880, 1.0445896, 0.6141800, 1.3093932, 0.3771100, 0.91~
## $ OTX2       <dbl> 0.9992054, 0.9078300, 0.6148917, 1.7928421, 0.5696953, 0.77~
```

```
d5F <- read.xlsx(xlsxFile = "MetabolismPaperRawData_Byran_working.xlsx",
                 colNames = TRUE, sheet="Fig 5_F")
glimpse(d5F)
```

```
## Rows: 32
## Columns: 8
## $ Chemical   <chr> "PBS", "PBS", "PBS", "PBS", "FGF2", "FGF2", "FGF2", "FGF2",~
## $ Sample     <dbl> 1, 2, 3, 4, 1, 2, 3, 4, 1, 2, 3, 4, 1, 2, 3, 4, 1, 2, 3, 4,~
## $ `Time.(h)` <dbl> 24, 24, 24, 24, 24, 24, 24, 24, 24, 24, 24, 24, 24, 24, 24,~
## $ TGFb2      <dbl> 1.0408138, 0.9427125, 1.1536739, 0.8834147, 2.7705353, 2.20~
## $ VIM        <dbl> 0.8836757, 1.0358740, 1.1462509, 0.9530605, 0.7944647, 0.69~
## $ SNAI1      <dbl> 1.0273824, 1.0213613, 1.1766059, 0.8099486, 0.8444048, 0.66~
## $ PITX1      <dbl> 1.6556716, 1.1297966, 1.2822236, 0.4169287, 0.3300321, 0.54~
## $ aSMA       <dbl> 1.51305862, 0.92738579, 0.78124815, 0.91220996, 0.40508463,~
```

```
d6SE <- read.xlsx(xlsxFile = "MetabolismPaperRawData_Byran_working.xlsx",
                  colNames = TRUE, sheet="Fig S6_E")
glimpse(d6SE)
```

```
## Rows: 32
## Columns: 5
## $ Chemical   <chr> "PBS", "PBS", "PBS", "PBS", "FGF2", "FGF2", "FGF2", "FGF2",~
## $ Sample     <dbl> 1, 2, 3, 4, 1, 2, 3, 4, 1, 2, 3, 4, 1, 2, 3, 4, 1, 2, 3, 4,~
## $ `Time.(h)` <dbl> 24, 24, 24, 24, 24, 24, 24, 24, 24, 24, 24, 24, 24, 24, 24,~
## $ PCNA       <dbl> 0.7240973, 1.5207968, 0.9943427, 1.3700884, 1.5356894, 4.72~
## $ E2F1       <dbl> 0.5978008, 1.3172125, 1.1166214, 1.1373175, 1.9053047, 3.67~
```

```
# to compare the 4 treatments at each time point
d4C_long <- d4C %>%
  rename(Time=`Time.(h)`) %>%
```

```r
  pivot_longer(cols=4:9,names_to="Gene",values_to="RGE") %>%
  mutate(Time=as.factor(Time))
glimpse(d4C_long)
```

```
## Rows: 192
## Columns: 5
## $ Chemical <chr> "PBS", "PBS", "PBS", "PBS", "PBS", "PBS", "PBS", "PBS", "PBS"~
## $ Sample   <dbl> 1, 1, 1, 1, 1, 1, 2, 2, 2, 2, 2, 2, 3, 3, 3, 3, 3, 3, 4, 4, 4~
## $ Time     <fct> 24, 24, 24, 24, 24, 24, 24, 24, 24, 24, 24, 24, 24, 24, 24, 2~
## $ Gene     <chr> "SOX2", "SIX6", "PAX6", "RPE65", "TYR", "OTX2", "SOX2", "SIX6~
## $ RGE      <dbl> 0.6683726, 1.2109944, 0.8869278, 1.0168114, 1.1903880, 0.9992~
```

```r
d5F_long <- d5F %>%
  rename(Time=`Time.(h)`) %>%
  pivot_longer(cols=4:8,names_to="Gene",values_to="RGE") %>%
  mutate(Time=as.factor(Time))
glimpse(d5F_long)
```

```
## Rows: 160
## Columns: 5
## $ Chemical <chr> "PBS", "PBS", "PBS", "PBS", "PBS", "PBS", "PBS", "PBS", "PBS"~
## $ Sample   <dbl> 1, 1, 1, 1, 1, 2, 2, 2, 2, 2, 3, 3, 3, 3, 3, 4, 4, 4, 4, 4, 1~
## $ Time     <fct> 24, 24, 24, 24, 24, 24, 24, 24, 24, 24, 24, 24, 24, 24, 24, 2~
## $ Gene     <chr> "TGFb2", "VIM", "SNAI1", "PITX1", "aSMA", "TGFb2", "VIM", "SN~
## $ RGE      <dbl> 1.0408138, 0.8836757, 1.0273824, 1.6556716, 1.5130586, 0.9427~
```

```r
d6SE_long <- d6SE %>%
  rename(Time=`Time.(h)`) %>%
  pivot_longer(cols=4:5,names_to="Gene",values_to="RGE") %>%
  mutate(Time=as.factor(Time))
glimpse(d6SE_long)
```

```
## Rows: 64
## Columns: 5
## $ Chemical <chr> "PBS", "PBS", "PBS", "PBS", "PBS", "PBS", "PBS", "PBS", "FGF2~
## $ Sample   <dbl> 1, 1, 2, 2, 3, 3, 4, 4, 1, 1, 2, 2, 3, 3, 4, 4, 1, 1, 2, 2, 3~
## $ Time     <fct> 24, 24, 24, 24, 24, 24, 24, 24, 24, 24, 24, 24, 24, 24, 24, 2~
## $ Gene     <chr> "PCNA", "E2F1", "PCNA", "E2F1", "PCNA", "E2F1", "PCNA", "E2F1~
## $ RGE      <dbl> 0.7240973, 0.5978008, 1.5207968, 1.3172125, 0.9943427, 1.1166~
```

```r
ggplot(d4C_long, aes(x=Time,y=log2(RGE),color=Chemical)) +
  geom_jitter(width=.05,height=0) +
  facet_wrap(~Gene, nrow=3) +
  labs(title="For Figure 4C: Time Comparisons")
```

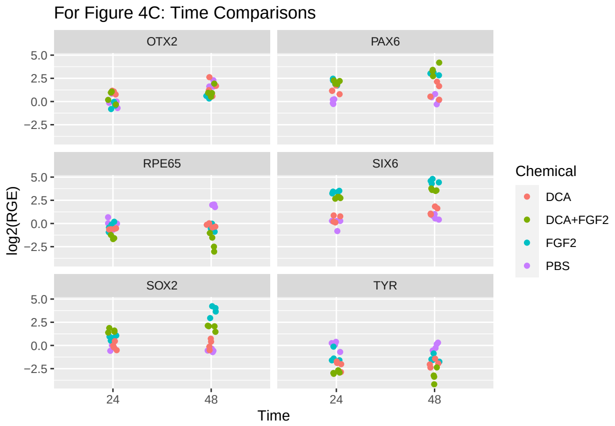

```r
ggplot(d5F_long, aes(x=Time,y=log2(RGE),color=Chemical)) +
  geom_jitter(width=.05,height=0) +
  facet_wrap(~Gene, nrow=2) +
  labs(title="For Figure 5F: Time Comparisons")
```

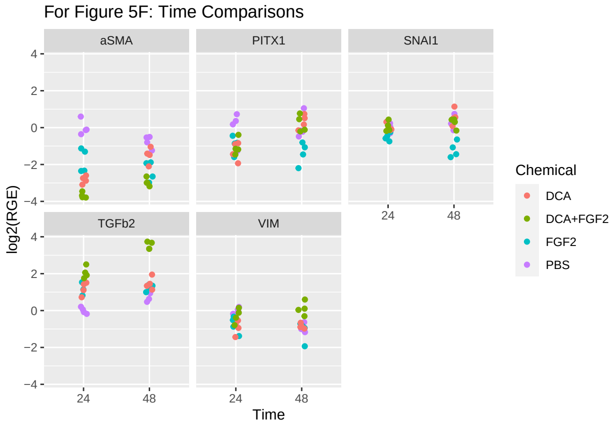

```r
ggplot(d6SE_long, aes(x=Time,y=log2(RGE),color=Chemical)) +
  geom_jitter(width=.05,height=0) +
  facet_wrap(~Gene, nrow=1) +
  labs(title="For Figure S6E: Time Comparisons")
```

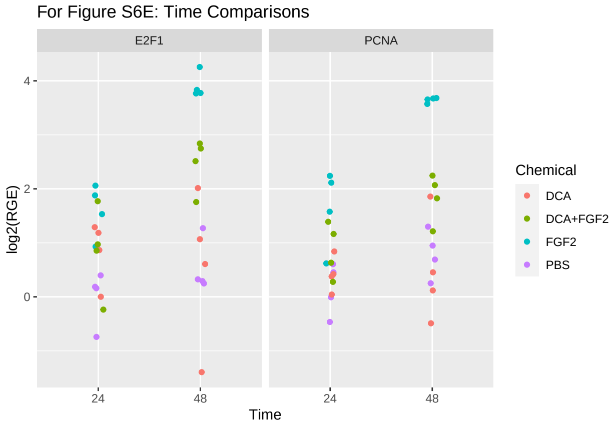

#### Data Analysis (Experiment 4)

The analysis here is to compare the four specified pairs of means across the two time points. We will treat this like a two-way ANOVA, with Treatment and Time as the factors, and in the end pull out the pairs of means of interest and adjust their p-values by controlling the False Discovery Rate across the eight pairs of means for each dataset.

```
d4C <- d4C %>%
  rename(Time=`Time.(h)`) %>%
  mutate(Time=as.factor(Time))
glimpse(d4C)
```

```
## Rows: 32
## Columns: 9
## $ Chemical <chr> "PBS", "PBS", "PBS", "PBS", "FGF2", "FGF2", "FGF2", "FGF2", "~
## $ Sample   <dbl> 1, 2, 3, 4, 1, 2, 3, 4, 1, 2, 3, 4, 1, 2, 3, 4, 1, 2, 3, 4, 1~
## $ Time     <fct> 24, 24, 24, 24, 24, 24, 24, 24, 24, 24, 24, 24, 24, 24, 24, 2~
## $ SOX2     <dbl> 0.6683726, 1.7839131, 1.0073875, 0.8325515, 2.0972087, 1.7403~
## $ SIX6     <dbl> 1.2109944, 1.2081256, 0.5674564, 1.2045180, 9.4502289, 10.430~
## $ PAX6     <dbl> 0.8869278, 0.8325113, 1.1858234, 1.1420932, 3.2961737, 4.7995~
## $ RPE65    <dbl> 1.0168114, 1.5935640, 0.6206805, 0.9943105, 0.9160471, 0.7381~
## $ TYR      <dbl> 1.1903880, 1.0445896, 0.6141800, 1.3093932, 0.3771100, 0.9148~
## $ OTX2     <dbl> 0.9992054, 0.9078300, 0.6148917, 1.7928421, 0.5696953, 0.7700~
```

```
d5F <- d5F %>%
  rename(Time=`Time.(h)`) %>%
  mutate(Time=as.factor(Time))
glimpse(d5F)
```

```
## Rows: 32
## Columns: 8
## $ Chemical <chr> "PBS", "PBS", "PBS", "PBS", "FGF2", "FGF2", "FGF2", "FGF2", "~
## $ Sample   <dbl> 1, 2, 3, 4, 1, 2, 3, 4, 1, 2, 3, 4, 1, 2, 3, 4, 1, 2, 3, 4, 1~
## $ Time     <fct> 24, 24, 24, 24, 24, 24, 24, 24, 24, 24, 24, 24, 24, 24, 24, 2~
## $ TGFb2    <dbl> 1.0408138, 0.9427125, 1.1536739, 0.8834147, 2.7705353, 2.2084~
## $ VIM      <dbl> 0.8836757, 1.0358740, 1.1462509, 0.9530605, 0.7944647, 0.6990~
## $ SNAI1    <dbl> 1.0273824, 1.0213613, 1.1766059, 0.8099486, 0.8444048, 0.6676~
## $ PITX1    <dbl> 1.6556716, 1.1297966, 1.2822236, 0.4169287, 0.3300321, 0.5494~
## $ aSMA     <dbl> 1.51305862, 0.92738579, 0.78124815, 0.91220996, 0.40508463, 0~
```

```
d6SE <- d6SE %>%
  rename(Time=`Time.(h)`) %>%
  mutate(Time=as.factor(Time))
glimpse(d6SE)
```

```
## Rows: 32
## Columns: 5
## $ Chemical <chr> "PBS", "PBS", "PBS", "PBS", "FGF2", "FGF2", "FGF2", "FGF2", "~
## $ Sample   <dbl> 1, 2, 3, 4, 1, 2, 3, 4, 1, 2, 3, 4, 1, 2, 3, 4, 1, 2, 3, 4, 1~
## $ Time     <fct> 24, 24, 24, 24, 24, 24, 24, 24, 24, 24, 24, 24, 24, 24, 24, 2~
## $ PCNA     <dbl> 0.7240973, 1.5207968, 0.9943427, 1.3700884, 1.5356894, 4.7239~
## $ E2F1     <dbl> 0.5978008, 1.3172125, 1.1166214, 1.1373175, 1.9053047, 3.6773~
```

#### Treatment Comparisons (Figure 4C)

```r
a4C_SOX2 <- aov(log2(SOX2)~Chemical*Time, data=d4C)
a4C_SIX6 <- aov(log2(SIX6)~Chemical*Time, data=d4C)
a4C_PAX6 <- aov(log2(PAX6)~Chemical*Time, data=d4C)
a4C_RPE65 <- aov(log2(RPE65)~Chemical*Time, data=d4C)
a4C_TYR <- aov(log2(TYR)~Chemical*Time, data=d4C)
a4C_OTX2 <- aov(log2(OTX2)~Chemical*Time, data=d4C)

plot(a4C_SOX2,which=1:2)
```

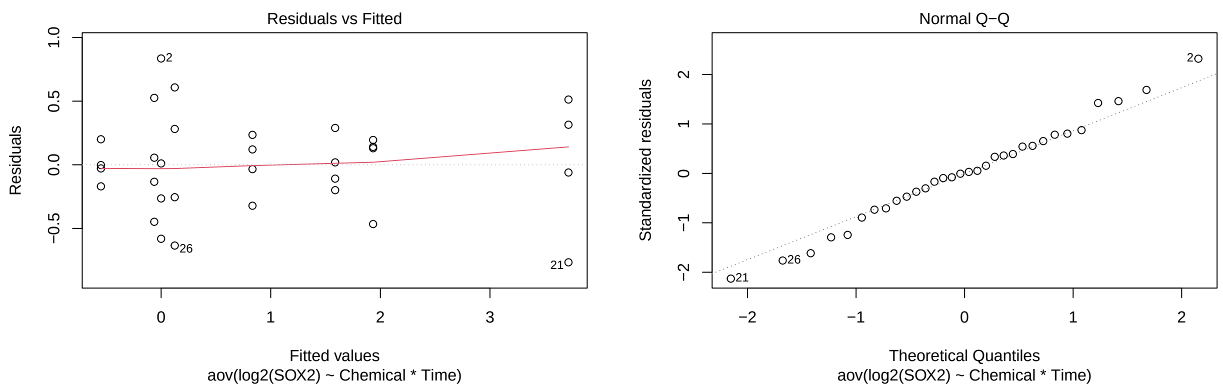

```r
plot(a4C_SIX6,which=1:2)
```

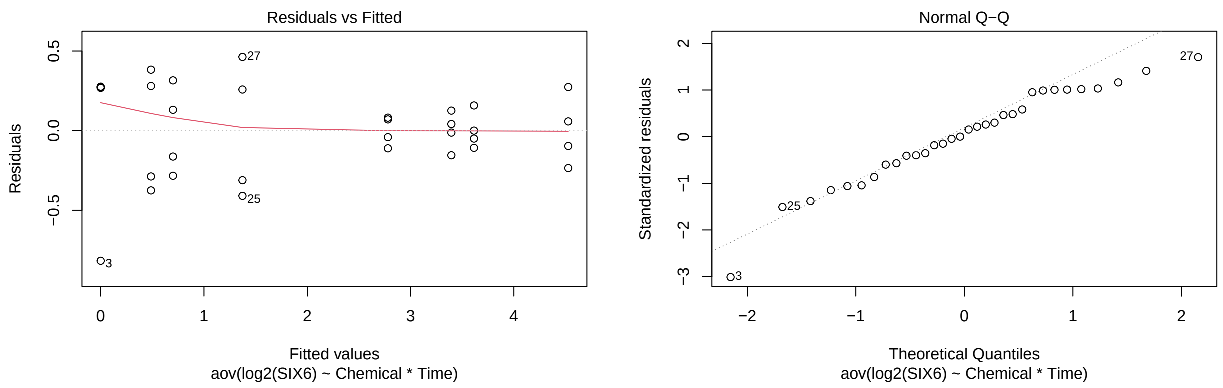

```r
plot(a4C_PAX6,which=1:2)
```

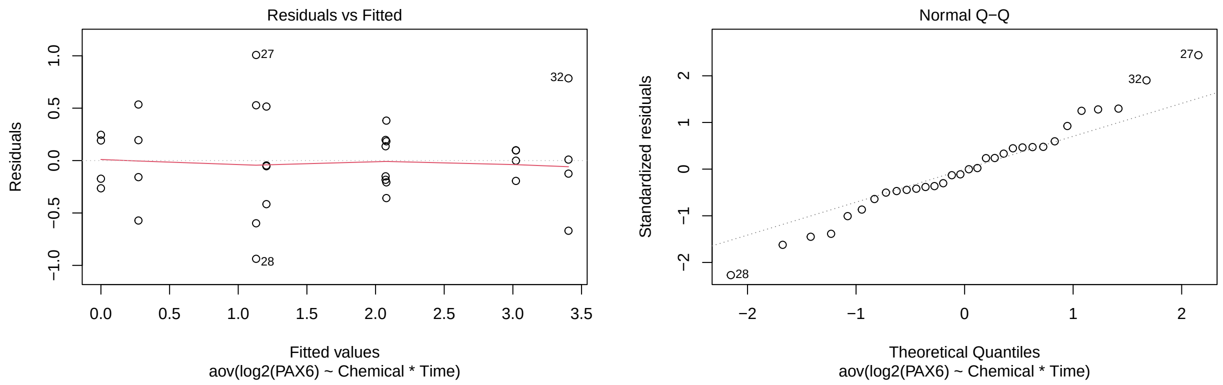

```r
plot(a4C_RPE65,which=1:2)
```

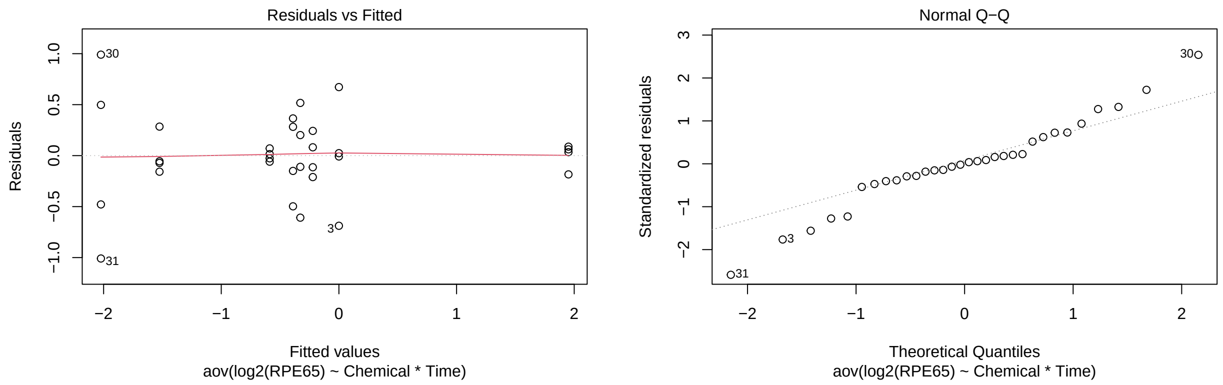

```r
plot(a4C_TYR,which=1:2)
```

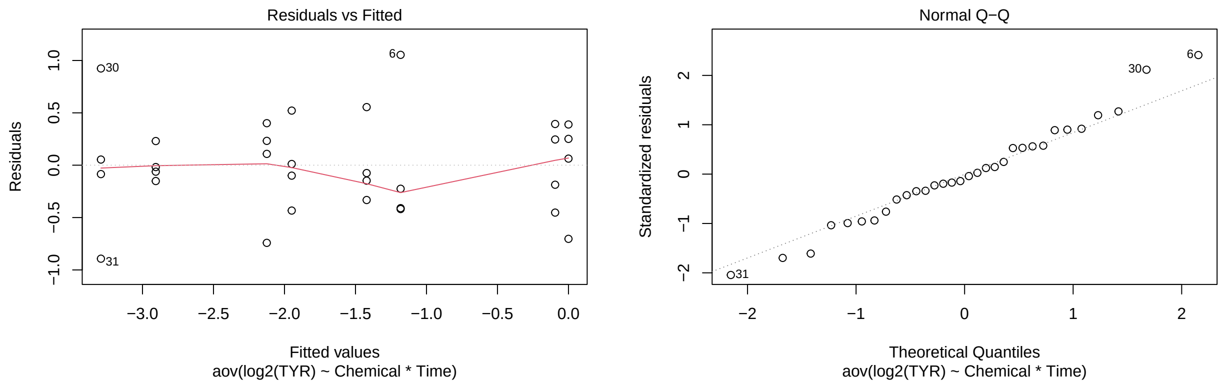

```r
plot(a4C_OTX2,which=1:2)
```

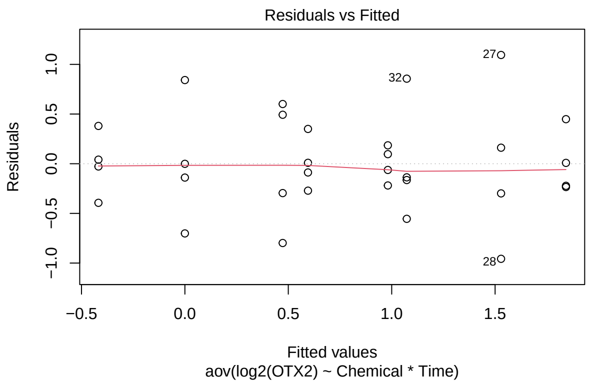

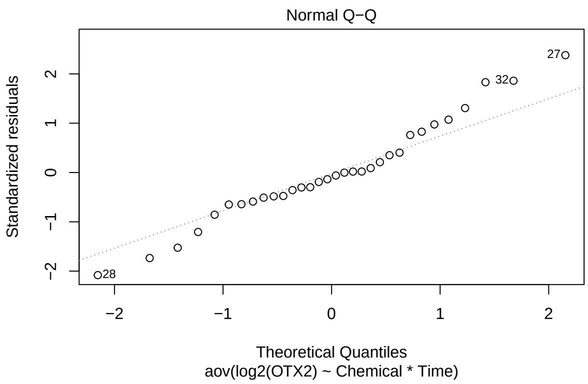

```
a4C_tSOX2 <- test(emmeans(a4C_SOX2, pairwise~Chemical|Time), adjust="none")
a4C_tSOX2
```

```
## $emmeans
## Time = 24:
##  Chemical  emmean    SE df t.ratio p.value
##  DCA      -0.0624 0.208 24  -0.301  0.7664
##  DCA+FGF2  1.5877 0.208 24   7.643  <.0001
##  FGF2      0.8334 0.208 24   4.012  0.0005
##  PBS       0.0000 0.208 24   0.000  1.0000
##
## Time = 48:
##  Chemical  emmean    SE df t.ratio p.value
##  DCA       0.1244 0.208 24   0.599  0.5550
##  DCA+FGF2  1.9339 0.208 24   9.309  <.0001
##  FGF2      3.7161 0.208 24  17.888  <.0001
##  PBS      -0.5492 0.208 24  -2.644  0.0142
##
## Results are given on the log2 (not the response) scale.
##
## $contrasts
## Time = 24:
##  contrast          estimate    SE df t.ratio p.value
##  DCA - (DCA+FGF2)   -1.6501 0.294 24  -5.617  <.0001
##  DCA - FGF2         -0.8959 0.294 24  -3.049  0.0055
##  DCA - PBS          -0.0624 0.294 24  -0.212  0.8335
##  (DCA+FGF2) - FGF2   0.7542 0.294 24   2.567  0.0169
##  (DCA+FGF2) - PBS    1.5877 0.294 24   5.404  <.0001
##  FGF2 - PBS          0.8334 0.294 24   2.837  0.0091
##
## Time = 48:
##  contrast          estimate    SE df t.ratio p.value
##  DCA - (DCA+FGF2)   -1.8095 0.294 24  -6.159  <.0001
##  DCA - FGF2         -3.5918 0.294 24 -12.226  <.0001
##  DCA - PBS           0.6735 0.294 24   2.293  0.0309
##  (DCA+FGF2) - FGF2  -1.7823 0.294 24  -6.066  <.0001
##  (DCA+FGF2) - PBS    2.4831 0.294 24   8.452  <.0001
##  FGF2 - PBS          4.2653 0.294 24  14.518  <.0001
##
```

```
## Results are given on the log2 (not the response) scale.
```

```r
p.adjust(a4C_tSOX2$contrasts$p.value[c(1,3,4,6,7,9,10,12)], method="fdr")
```

```
## [1] 1.761356e-05 8.335146e-01 2.254190e-02 1.457946e-02 7.717556e-06
## [6] 3.535914e-02 7.717556e-06 1.763962e-12
```

```r
a4C_tSIX6 <- test(emmeans(a4C_SIX6, pairwise~Chemical|Time), adjust="none")
a4C_tSIX6
```

```
## $emmeans
## Time = 24:
##  Chemical emmean    SE df t.ratio p.value
##  DCA       0.487 0.157 24   3.108  0.0048
##  DCA+FGF2  2.781 0.157 24  17.741  <.0001
##  FGF2      3.395 0.157 24  21.660  <.0001
##  PBS       0.000 0.157 24   0.000  1.0000
##
## Time = 48:
##  Chemical emmean    SE df t.ratio p.value
##  DCA       1.373 0.157 24   8.757  <.0001
##  DCA+FGF2  3.614 0.157 24  23.058  <.0001
##  FGF2      4.527 0.157 24  28.880  <.0001
##  PBS       0.700 0.157 24   4.466  0.0002
##
## Results are given on the log2 (not the response) scale.
##
## $contrasts
## Time = 24:
##  contrast          estimate    SE df t.ratio p.value
##  DCA - (DCA+FGF2)    -2.294 0.222 24 -10.347  <.0001
##  DCA - FGF2          -2.908 0.222 24 -13.119  <.0001
##  DCA - PBS            0.487 0.222 24   2.197  0.0379
##  (DCA+FGF2) - FGF2   -0.614 0.222 24  -2.771  0.0106
##  (DCA+FGF2) - PBS     2.781 0.222 24  12.545  <.0001
##  FGF2 - PBS           3.395 0.222 24  15.316  <.0001
##
## Time = 48:
##  contrast          estimate    SE df t.ratio p.value
##  DCA - (DCA+FGF2)    -2.242 0.222 24 -10.112  <.0001
##  DCA - FGF2          -3.154 0.222 24 -14.229  <.0001
##  DCA - PBS            0.673 0.222 24   3.034  0.0057
##  (DCA+FGF2) - FGF2   -0.912 0.222 24  -4.116  0.0004
##  (DCA+FGF2) - PBS     2.914 0.222 24  13.147  <.0001
##  FGF2 - PBS           3.827 0.222 24  17.263  <.0001
##
## Results are given on the log2 (not the response) scale.
```

```r
p.adjust(a4C_tSIX6$contrasts$p.value[c(1,3,4,6,7,9,10,12)], method="fdr")
```

```
## [1] 6.700047e-10 3.787782e-02 1.213227e-02 2.755549e-13 7.900621e-10
## [6] 7.626534e-03 6.278479e-04 3.925907e-14
```

```r
a4C_tPAX6 <- test(emmeans(a4C_PAX6, pairwise~Chemical|Time), adjust="none")
a4C_tPAX6
```

```
## $emmeans
## Time = 24:
##  Chemical emmean    SE df t.ratio p.value
##  DCA       1.205 0.238 24   5.059  <.0001
##  DCA+FGF2  2.073 0.238 24   8.702  <.0001
##  FGF2      2.079 0.238 24   8.725  <.0001
##  PBS       0.000 0.238 24   0.000  1.0000
##
## Time = 48:
##  Chemical emmean    SE df t.ratio p.value
##  DCA       1.130 0.238 24   4.744  0.0001
##  DCA+FGF2  3.405 0.238 24  14.292  <.0001
##  FGF2      3.022 0.238 24  12.686  <.0001
##  PBS       0.273 0.238 24   1.146  0.2629
##
## Results are given on the log2 (not the response) scale.
##
## $contrasts
## Time = 24:
##  contrast          estimate    SE df t.ratio p.value
##  DCA - (DCA+FGF2)   -0.8680 0.337 24  -2.576  0.0166
##  DCA - FGF2         -0.8734 0.337 24  -2.592  0.0160
##  DCA - PBS           1.2052 0.337 24   3.577  0.0015
##  (DCA+FGF2) - FGF2  -0.0054 0.337 24  -0.016  0.9873
##  (DCA+FGF2) - PBS    2.0732 0.337 24   6.153  <.0001
##  FGF2 - PBS          2.0786 0.337 24   6.169  <.0001
##
## Time = 48:
##  contrast          estimate    SE df t.ratio p.value
##  DCA - (DCA+FGF2)   -2.2747 0.337 24  -6.751  <.0001
##  DCA - FGF2         -1.8920 0.337 24  -5.616  <.0001
##  DCA - PBS           0.8571 0.337 24   2.544  0.0178
##  (DCA+FGF2) - FGF2   0.3827 0.337 24   1.136  0.2672
##  (DCA+FGF2) - PBS    3.1318 0.337 24   9.295  <.0001
##  FGF2 - PBS          2.7491 0.337 24   8.159  <.0001
##
## Results are given on the log2 (not the response) scale.
```

```r
p.adjust(a4C_tPAX6$contrasts$p.value[c(1,3,4,6,7,9,10,12)], method="fdr")
```

```
## [1] 2.376368e-02 3.043947e-03 9.873491e-01 5.998985e-06 2.215033e-06
## [6] 2.376368e-02 3.054044e-01 1.777588e-07
```

```r
a4C_tRPE65 <- test(emmeans(a4C_RPE65, pairwise~Chemical|Time), adjust="none")
a4C_tRPE65
```

```
## $emmeans
## Time = 24:
##  Chemical emmean    SE df t.ratio p.value
```

```
##  DCA      -0.588 0.225 24  -2.613  0.0152
##  DCA+FGF2 -1.525 0.225 24  -6.772  <.0001
##  FGF2     -0.328 0.225 24  -1.457  0.1581
##  PBS       0.000 0.225 24   0.000  1.0000
##
## Time = 48:
##  Chemical emmean    SE df t.ratio p.value
##  DCA      -0.222 0.225 24  -0.986  0.3342
##  DCA+FGF2 -2.023 0.225 24  -8.984  <.0001
##  FGF2     -0.390 0.225 24  -1.733  0.0960
##  PBS       1.951 0.225 24   8.666  <.0001
##
## Results are given on the log2 (not the response) scale.
##
## $contrasts
## Time = 24:
##  contrast          estimate    SE df t.ratio p.value
##  DCA - (DCA+FGF2)     0.936 0.318 24   2.941  0.0071
##  DCA - FGF2          -0.260 0.318 24  -0.817  0.4217
##  DCA - PBS           -0.588 0.318 24  -1.848  0.0770
##  (DCA+FGF2) - FGF2   -1.197 0.318 24  -3.758  0.0010
##  (DCA+FGF2) - PBS    -1.525 0.318 24  -4.789  0.0001
##  FGF2 - PBS          -0.328 0.318 24  -1.030  0.3132
##
## Time = 48:
##  contrast          estimate    SE df t.ratio p.value
##  DCA - (DCA+FGF2)     1.801 0.318 24   5.655  <.0001
##  DCA - FGF2           0.168 0.318 24   0.528  0.6021
##  DCA - PBS           -2.173 0.318 24  -6.824  <.0001
##  (DCA+FGF2) - FGF2   -1.633 0.318 24  -5.127  <.0001
##  (DCA+FGF2) - PBS    -3.974 0.318 24 -12.480  <.0001
##  FGF2 - PBS          -2.341 0.318 24  -7.353  <.0001
##
## Results are given on the log2 (not the response) scale.
```

```
p.adjust(a4C_tRPE65$contrasts$p.value[c(1,3,4,6,7,9,10,12)], method="fdr")
```

```
## [1] 9.519048e-03 8.800521e-02 1.548993e-03 3.131738e-01 2.132246e-05
## [6] 1.862928e-06 6.027908e-05 1.086607e-06
```

```
a4C_tTYR <- test(emmeans(a4C_TYR, pairwise~Chemical|Time), adjust="none")
a4C_tTYR
```

```
## $emmeans
## Time = 24:
##  Chemical  emmean    SE df t.ratio p.value
##  DCA      -2.1245 0.252 24  -8.425  <.0001
##  DCA+FGF2 -2.9064 0.252 24 -11.525  <.0001
##  FGF2     -1.1821 0.252 24  -4.688  0.0001
##  PBS       0.0000 0.252 24   0.000  1.0000
##
## Time = 48:
##  Chemical  emmean    SE df t.ratio p.value
```

```
##  DCA      -1.9490 0.252 24  -7.729  <.0001
##  DCA+FGF2 -3.2927 0.252 24 -13.057  <.0001
##  FGF2     -1.4213 0.252 24  -5.636  <.0001
##  PBS      -0.0939 0.252 24  -0.372  0.7130
##
## Results are given on the log2 (not the response) scale.
##
## $contrasts
## Time = 24:
##  contrast          estimate    SE df t.ratio p.value
##  DCA - (DCA+FGF2)     0.782 0.357 24   2.192  0.0383
##  DCA - FGF2          -0.942 0.357 24  -2.642  0.0143
##  DCA - PBS           -2.125 0.357 24  -5.957  <.0001
##  (DCA+FGF2) - FGF2   -1.724 0.357 24  -4.835  0.0001
##  (DCA+FGF2) - PBS    -2.906 0.357 24  -8.149  <.0001
##  FGF2 - PBS          -1.182 0.357 24  -3.315  0.0029
##
## Time = 48:
##  contrast          estimate    SE df t.ratio p.value
##  DCA - (DCA+FGF2)     1.344 0.357 24   3.768  0.0009
##  DCA - FGF2          -0.528 0.357 24  -1.480  0.1519
##  DCA - PBS           -1.855 0.357 24  -5.202  <.0001
##  (DCA+FGF2) - FGF2   -1.871 0.357 24  -5.247  <.0001
##  (DCA+FGF2) - PBS    -3.199 0.357 24  -8.969  <.0001
##  FGF2 - PBS          -1.327 0.357 24  -3.722  0.0011
##
## Results are given on the log2 (not the response) scale.
```

```
p.adjust(a4C_tTYR$contrasts$p.value[c(1,3,4,6,7,9,10,12)], method="fdr")
```

```
## [1] 3.829636e-02 3.029543e-05 1.264807e-04 3.320746e-03 1.413502e-03
## [6] 6.653059e-05 6.653059e-05 1.413502e-03
```

```
a4C_tOTX2 <- test(emmeans(a4C_OTX2, pairwise~Chemical|Time), adjust="none")
a4C_tOTX2
```

```
## $emmeans
## Time = 24:
##  Chemical emmean    SE df t.ratio p.value
##  DCA        0.981 0.266 24   3.695  0.0011
##  DCA+FGF2   0.473 0.266 24   1.781  0.0876
##  FGF2      -0.418 0.266 24  -1.574  0.1285
##  PBS        0.000 0.266 24   0.000  1.0000
##
## Time = 48:
##  Chemical emmean    SE df t.ratio p.value
##  DCA        1.529 0.266 24   5.756  <.0001
##  DCA+FGF2   1.073 0.266 24   4.039  0.0005
##  FGF2       0.595 0.266 24   2.241  0.0345
##  PBS        1.843 0.266 24   6.938  <.0001
##
## Results are given on the log2 (not the response) scale.
##
```

```
## $contrasts
## Time = 24:
##  contrast           estimate    SE df t.ratio p.value
##  DCA - (DCA+FGF2)      0.508 0.376 24   1.354  0.1885
##  DCA - FGF2            1.400 0.376 24   3.726  0.0011
##  DCA - PBS             0.981 0.376 24   2.613  0.0153
##  (DCA+FGF2) - FGF2     0.891 0.376 24   2.372  0.0260
##  (DCA+FGF2) - PBS      0.473 0.376 24   1.259  0.2201
##  FGF2 - PBS           -0.418 0.376 24  -1.113  0.2767
##
## Time = 48:
##  contrast           estimate    SE df t.ratio p.value
##  DCA - (DCA+FGF2)      0.456 0.376 24   1.214  0.2365
##  DCA - FGF2            0.934 0.376 24   2.485  0.0203
##  DCA - PBS            -0.314 0.376 24  -0.836  0.4116
##  (DCA+FGF2) - FGF2     0.478 0.376 24   1.271  0.2158
##  (DCA+FGF2) - PBS     -0.770 0.376 24  -2.050  0.0515
##  FGF2 - PBS           -1.247 0.376 24  -3.321  0.0029
##
## Results are given on the log2 (not the response) scale.
```

```
p.adjust(a4C_tOTX2$contrasts$p.value[c(1,3,4,6,7,9,10,12)], method="fdr")
```

```
## [1] 0.31539093 0.06104258 0.06945511 0.31621250 0.31539093 0.41164082 0.31539093
## [8] 0.02288881
```

##### Treatment Comparisons (Figure 5F)

```
a5F_TGFb2 <- aov(log2(TGFb2)~Chemical*Time, data=d5F)
a5F_VIM <- aov(log2(VIM)~Chemical*Time, data=d5F)
a5F_SNAI1 <- aov(log2(SNAI1)~Chemical*Time, data=d5F)
a5F_PITX1 <- aov(log2(PITX1)~Chemical*Time, data=d5F)
a5F_aSMA <- aov(log2(aSMA)~Chemical*Time, data=d5F)

plot(a5F_TGFb2,which=1:2)
```

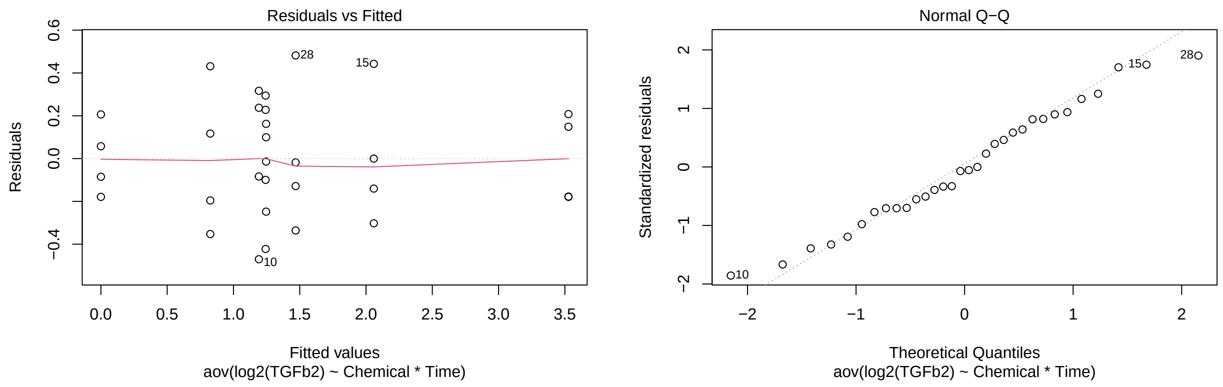

```
plot(a5F_VIM,which=1:2)
```

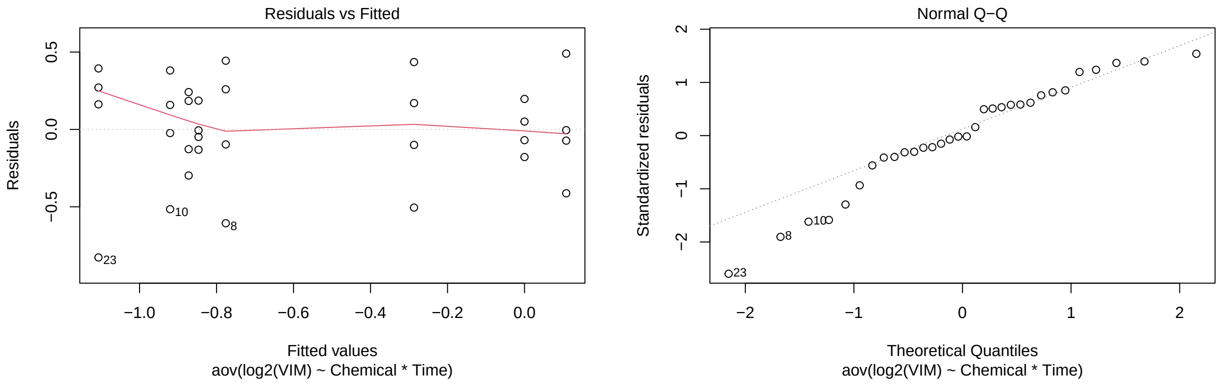

```
plot(a5F_SNAI1,which=1:2)
```

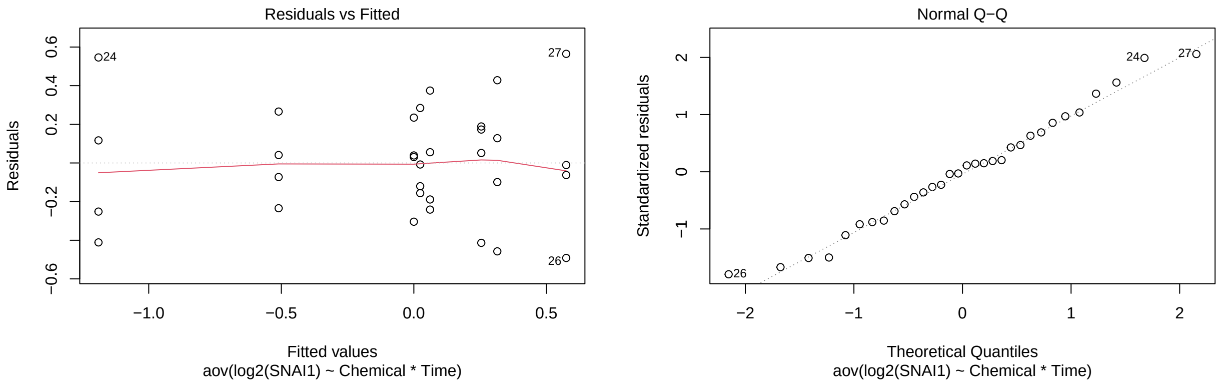

```
plot(a5F_PITX1,which=1:2)
```

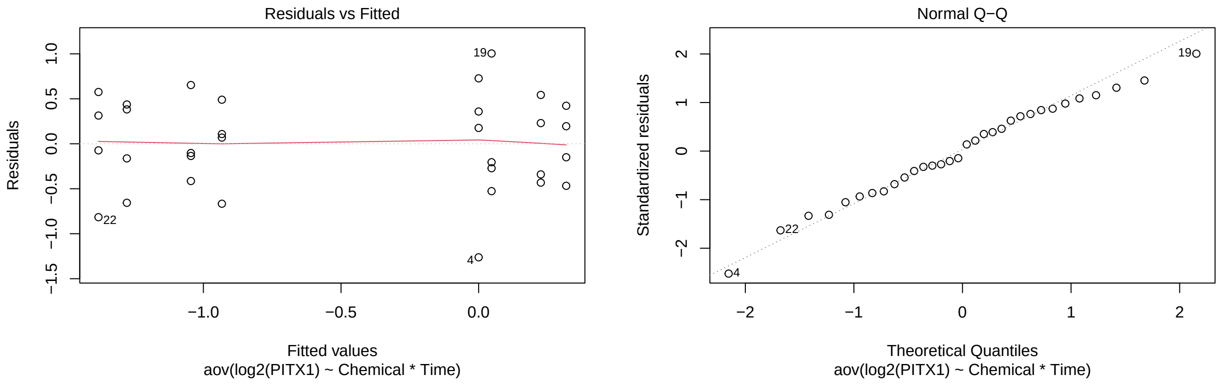

```
plot(a5F_aSMA,which=1:2)
```

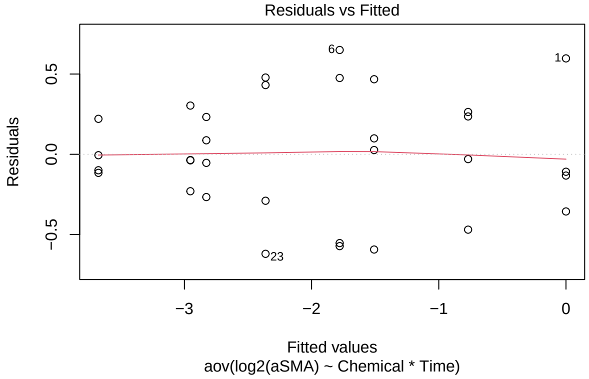

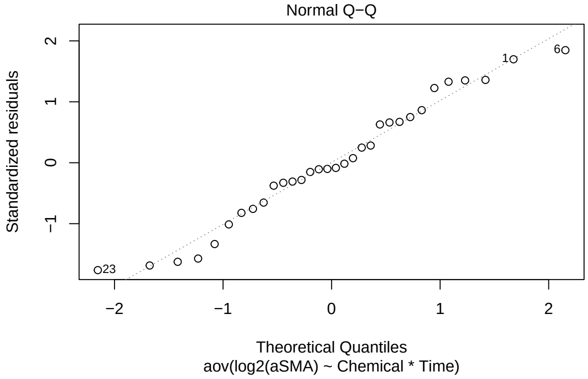

```r
a5F_tTGFb2 <- test(emmeans(a5F_TGFb2, pairwise~Chemical|Time), adjust="none")
a5F_tTGFb2
```

```
## $emmeans
## Time = 24:
##  Chemical emmean    SE df t.ratio p.value
##  DCA       1.192 0.146 24   8.146  <.0001
##  DCA+FGF2  2.059 0.146 24  14.073  <.0001
##  FGF2      1.243 0.146 24   8.493  <.0001
##  PBS       0.000 0.146 24   0.000  1.0000
##
## Time = 48:
##  Chemical emmean    SE df t.ratio p.value
##  DCA       1.469 0.146 24  10.042  <.0001
##  DCA+FGF2  3.527 0.146 24  24.110  <.0001
##  FGF2      1.246 0.146 24   8.518  <.0001
##  PBS       0.825 0.146 24   5.639  <.0001
##
## Results are given on the log2 (not the response) scale.
##
## $contrasts
## Time = 24:
##  contrast          estimate    SE df t.ratio p.value
##  DCA - (DCA+FGF2)   -0.8672 0.207 24  -4.191  0.0003
##  DCA - FGF2         -0.0508 0.207 24  -0.246  0.8081
##  DCA - PBS           1.1917 0.207 24   5.760  <.0001
##  (DCA+FGF2) - FGF2   0.8164 0.207 24   3.946  0.0006
##  (DCA+FGF2) - PBS    2.0589 0.207 24   9.951  <.0001
##  FGF2 - PBS          1.2425 0.207 24   6.006  <.0001
##
## Time = 48:
##  contrast          estimate    SE df t.ratio p.value
##  DCA - (DCA+FGF2)   -2.0582 0.207 24  -9.948  <.0001
##  DCA - FGF2          0.2229 0.207 24   1.078  0.2920
##  DCA - PBS           0.6442 0.207 24   3.113  0.0047
##  (DCA+FGF2) - FGF2   2.2811 0.207 24  11.025  <.0001
##  (DCA+FGF2) - PBS    2.7023 0.207 24  13.061  <.0001
##  FGF2 - PBS          0.4212 0.207 24   2.036  0.0529
##
```

```
## Results are given on the log2 (not the response) scale.
```

```r
p.adjust(a5F_tTGFb2$contrasts$p.value[c(1,3,4,6,7,9,10,12)], method="fdr")
```

```
## [1] 5.193613e-04 1.232797e-05 8.057257e-04 8.961252e-06 2.177526e-09
## [6] 5.408879e-03 5.654277e-10 5.294552e-02
```

```r
a5F_tVIM <- test(emmeans(a5F_VIM, pairwise~Chemical|Time), adjust="none")
a5F_tVIM
```

```
## $emmeans
## Time = 24:
##  Chemical emmean    SE df t.ratio p.value
##  DCA      -0.921 0.184 24  -5.006  <.0001
##  DCA+FGF2 -0.287 0.184 24  -1.560  0.1317
##  FGF2     -0.776 0.184 24  -4.220  0.0003
##  PBS       0.000 0.184 24   0.000  1.0000
##
## Time = 48:
##  Chemical emmean    SE df t.ratio p.value
##  DCA      -0.847 0.184 24  -4.605  0.0001
##  DCA+FGF2  0.108 0.184 24   0.589  0.5614
##  FGF2     -1.107 0.184 24  -6.018  <.0001
##  PBS      -0.872 0.184 24  -4.744  0.0001
##
## Results are given on the log2 (not the response) scale.
##
## $contrasts
## Time = 24:
##  contrast          estimate   SE df t.ratio p.value
##  DCA - (DCA+FGF2)   -0.6336 0.26 24  -2.436  0.0226
##  DCA - FGF2         -0.1446 0.26 24  -0.556  0.5834
##  DCA - PBS          -0.9206 0.26 24  -3.540  0.0017
##  (DCA+FGF2) - FGF2   0.4890 0.26 24   1.880  0.0723
##  (DCA+FGF2) - PBS   -0.2870 0.26 24  -1.103  0.2808
##  FGF2 - PBS         -0.7760 0.26 24  -2.984  0.0065
##
## Time = 48:
##  contrast          estimate   SE df t.ratio p.value
##  DCA - (DCA+FGF2)   -0.9551 0.26 24  -3.672  0.0012
##  DCA - FGF2          0.2599 0.26 24   0.999  0.3276
##  DCA - PBS           0.0256 0.26 24   0.099  0.9223
##  (DCA+FGF2) - FGF2   1.2150 0.26 24   4.672  0.0001
##  (DCA+FGF2) - PBS    0.9808 0.26 24   3.771  0.0009
##  FGF2 - PBS         -0.2343 0.26 24  -0.901  0.3766
##
## Results are given on the log2 (not the response) scale.
```

```r
p.adjust(a5F_tVIM$contrasts$p.value[c(1,3,4,6,7,9,10,12)], method="fdr")
```

```
## [1] 0.0362149639 0.0044533335 0.0963551619 0.0129024994 0.0044533335
## [6] 0.9223288999 0.0007654871 0.4304553400
```

```r
a5F_tSNAI1 <- test(emmeans(a5F_SNAI1, pairwise~Chemical|Time), adjust="none")
a5F_tSNAI1
```

```
## $emmeans
## Time = 24:
##  Chemical  emmean    SE df t.ratio p.value
##  DCA       0.0239 0.158 24   0.151  0.8812
##  DCA+FGF2  0.0611 0.158 24   0.386  0.7031
##  FGF2     -0.5101 0.158 24  -3.221  0.0036
##  PBS       0.0000 0.158 24   0.000  1.0000
##
## Time = 48:
##  Chemical  emmean    SE df t.ratio p.value
##  DCA       0.5746 0.158 24   3.629  0.0013
##  DCA+FGF2  0.2543 0.158 24   1.606  0.1214
##  FGF2     -1.1896 0.158 24  -7.513  <.0001
##  PBS       0.3148 0.158 24   1.988  0.0583
##
## Results are given on the log2 (not the response) scale.
##
## $contrasts
## Time = 24:
##  contrast          estimate    SE df t.ratio p.value
##  DCA - (DCA+FGF2)   -0.0372 0.224 24  -0.166  0.8696
##  DCA - FGF2          0.5340 0.224 24   2.385  0.0253
##  DCA - PBS           0.0239 0.224 24   0.107  0.9159
##  (DCA+FGF2) - FGF2   0.5711 0.224 24   2.551  0.0176
##  (DCA+FGF2) - PBS    0.0611 0.224 24   0.273  0.7874
##  FGF2 - PBS         -0.5101 0.224 24  -2.278  0.0319
##
## Time = 48:
##  contrast          estimate    SE df t.ratio p.value
##  DCA - (DCA+FGF2)    0.3203 0.224 24   1.431  0.1654
##  DCA - FGF2          1.7643 0.224 24   7.879  <.0001
##  DCA - PBS           0.2599 0.224 24   1.161  0.2572
##  (DCA+FGF2) - FGF2   1.4439 0.224 24   6.448  <.0001
##  (DCA+FGF2) - PBS   -0.0605 0.224 24  -0.270  0.7894
##  FGF2 - PBS         -1.5044 0.224 24  -6.718  <.0001
##
## Results are given on the log2 (not the response) scale.
```

```r
p.adjust(a5F_tSNAI1$contrasts$p.value[c(1,3,4,6,7,9,10,12)], method="fdr")
```

```
## [1] 9.158527e-01 9.158527e-01 4.680553e-02 6.385639e-02 2.646950e-01
## [6] 3.429954e-01 4.574759e-06 4.574759e-06
```

```r
a5F_tPITX1 <- test(emmeans(a5F_PITX1, pairwise~Chemical|Time), adjust="none")
a5F_tPITX1
```

```
## $emmeans
## Time = 24:
##  Chemical emmean    SE df t.ratio p.value
```

```
##  DCA      -1.278 0.289 24  -4.424  0.0002
##  DCA+FGF2 -1.045 0.289 24  -3.618  0.0014
##  FGF2     -0.933 0.289 24  -3.228  0.0036
##  PBS       0.000 0.289 24   0.000  1.0000
##
## Time = 48:
##  Chemical emmean    SE df t.ratio p.value
##  DCA       0.318 0.289 24   1.101  0.2817
##  DCA+FGF2  0.226 0.289 24   0.782  0.4418
##  FGF2     -1.381 0.289 24  -4.781  0.0001
##  PBS       0.047 0.289 24   0.163  0.8722
##
## Results are given on the log2 (not the response) scale.
##
## $contrasts
## Time = 24:
##  contrast          estimate    SE df t.ratio p.value
##  DCA - (DCA+FGF2)   -0.2327 0.409 24  -0.570  0.5742
##  DCA - FGF2         -0.3453 0.409 24  -0.845  0.4063
##  DCA - PBS          -1.2780 0.409 24  -3.128  0.0046
##  (DCA+FGF2) - FGF2  -0.1126 0.409 24  -0.276  0.7852
##  (DCA+FGF2) - PBS   -1.0453 0.409 24  -2.558  0.0172
##  FGF2 - PBS         -0.9327 0.409 24  -2.283  0.0316
##
## Time = 48:
##  contrast          estimate    SE df t.ratio p.value
##  DCA - (DCA+FGF2)    0.0922 0.409 24   0.226  0.8233
##  DCA - FGF2          1.6993 0.409 24   4.159  0.0004
##  DCA - PBS           0.2712 0.409 24   0.664  0.5132
##  (DCA+FGF2) - FGF2   1.6071 0.409 24   3.934  0.0006
##  (DCA+FGF2) - PBS    0.1790 0.409 24   0.438  0.6653
##  FGF2 - PBS         -1.4281 0.409 24  -3.495  0.0019
##
## Results are given on the log2 (not the response) scale.
```

```
p.adjust(a5F_tPITX1$contrasts$p.value[c(1,3,4,6,7,9,10,12)], method="fdr")
```

```
## [1] 0.765654800 0.012182496 0.823330619 0.063177319 0.823330619 0.765654800
## [7] 0.004985168 0.007452103
```

```
a5F_taSMA <- test(emmeans(a5F_aSMA, pairwise~Chemical|Time), adjust="none")
a5F_taSMA
```

```
## $emmeans
## Time = 24:
##  Chemical emmean    SE df t.ratio p.value
##  DCA      -2.828 0.203 24 -13.920  <.0001
##  DCA+FGF2 -3.676 0.203 24 -18.096  <.0001
##  FGF2     -1.779 0.203 24  -8.758  <.0001
##  PBS       0.000 0.203 24   0.000  1.0000
##
## Time = 48:
##  Chemical emmean    SE df t.ratio p.value
```

```
##  DCA      -1.508 0.203 24  -7.426  <.0001
##  DCA+FGF2 -2.953 0.203 24 -14.538  <.0001
##  FGF2     -2.362 0.203 24 -11.628  <.0001
##  PBS      -0.769 0.203 24  -3.788  0.0009
##
## Results are given on the log2 (not the response) scale.
##
## $contrasts
## Time = 24:
##  contrast          estimate    SE df t.ratio p.value
##  DCA - (DCA+FGF2)     0.848 0.287 24   2.953  0.0069
##  DCA - FGF2          -1.048 0.287 24  -3.650  0.0013
##  DCA - PBS           -2.828 0.287 24  -9.843  <.0001
##  (DCA+FGF2) - FGF2   -1.897 0.287 24  -6.603  <.0001
##  (DCA+FGF2) - PBS    -3.676 0.287 24 -12.796  <.0001
##  FGF2 - PBS          -1.779 0.287 24  -6.193  <.0001
##
## Time = 48:
##  contrast          estimate    SE df t.ratio p.value
##  DCA - (DCA+FGF2)     1.445 0.287 24   5.029  <.0001
##  DCA - FGF2           0.854 0.287 24   2.971  0.0066
##  DCA - PBS           -0.739 0.287 24  -2.572  0.0167
##  (DCA+FGF2) - FGF2   -0.591 0.287 24  -2.058  0.0507
##  (DCA+FGF2) - PBS    -2.184 0.287 24  -7.601  <.0001
##  FGF2 - PBS          -1.593 0.287 24  -5.544  <.0001
##
## Results are given on the log2 (not the response) scale.
```

```r
p.adjust(a5F_taSMA$contrasts$p.value[c(1,3,4,6,7,9,10,12)], method="fdr")
```

```
## [1] 9.247238e-03 5.352789e-09 3.156119e-06 5.662158e-06 6.181404e-05
## [6] 1.910980e-02 5.065439e-02 2.114312e-05
```

##### Treatment Comparisons (Figure S6E)

```r
a6SE_PCNA <- aov(log2(PCNA)~Chemical*Time, data=d6SE)
a6SE_E2F1 <- aov(log2(E2F1)~Chemical*Time, data=d6SE)

plot(a6SE_PCNA,which=1:2)
```

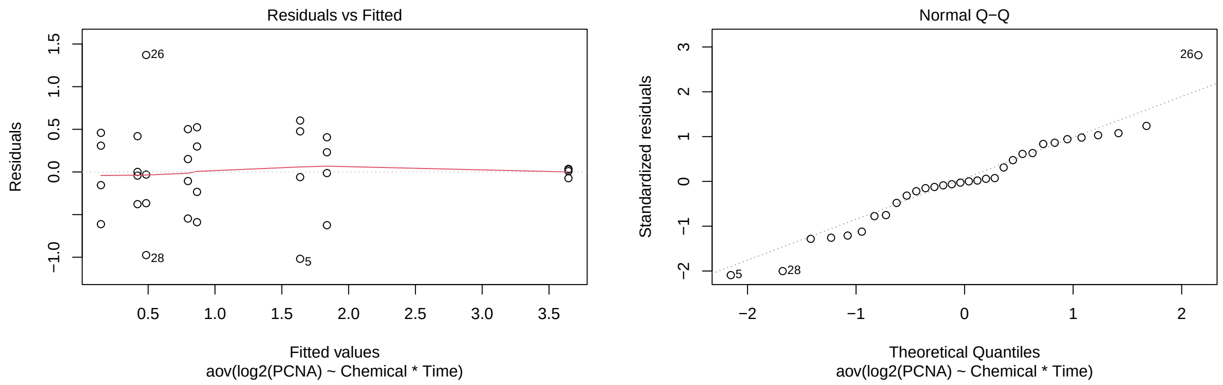

```r
plot(a6SE_E2F1,which=1:2)
```

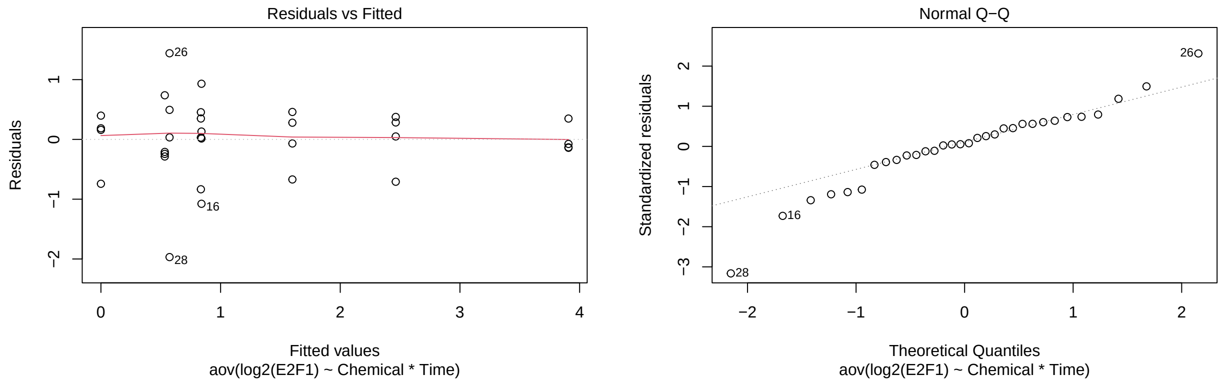

```r
a6SE_tPCNA <- test(emmeans(a6SE_PCNA, pairwise~Chemical|Time), adjust="none")
a6SE_tPCNA
```

```
## $emmeans
## Time = 24:
##  Chemical emmean    SE df t.ratio p.value
##  DCA        0.420 0.281 24   1.495  0.1480
##  DCA+FGF2   0.866 0.281 24   3.079  0.0051
##  FGF2       1.637 0.281 24   5.824  <.0001
##  PBS        0.146 0.281 24   0.520  0.6076
##
## Time = 48:
##  Chemical emmean    SE df t.ratio p.value
##  DCA        0.485 0.281 24   1.725  0.0975
##  DCA+FGF2   1.838 0.281 24   6.537  <.0001
##  FGF2       3.645 0.281 24  12.968  <.0001
##  PBS        0.797 0.281 24   2.837  0.0091
##
## Results are given on the log2 (not the response) scale.
##
## $contrasts
## Time = 24:
```

```
##  contrast           estimate    SE df t.ratio p.value
##  DCA - (DCA+FGF2)    -0.445 0.398 24  -1.120  0.2737
##  DCA - FGF2          -1.217 0.398 24  -3.061  0.0054
##  DCA - PBS            0.274 0.398 24   0.689  0.4974
##  (DCA+FGF2) - FGF2   -0.772 0.398 24  -1.941  0.0641
##  (DCA+FGF2) - PBS     0.719 0.398 24   1.809  0.0829
##  FGF2 - PBS           1.491 0.398 24   3.750  0.0010
##
## Time = 48:
##  contrast           estimate    SE df t.ratio p.value
##  DCA - (DCA+FGF2)    -1.353 0.398 24  -3.403  0.0023
##  DCA - FGF2          -3.161 0.398 24  -7.950  <.0001
##  DCA - PBS           -0.313 0.398 24  -0.786  0.4393
##  (DCA+FGF2) - FGF2   -1.808 0.398 24  -4.547  0.0001
##  (DCA+FGF2) - PBS     1.040 0.398 24   2.617  0.0151
##  FGF2 - PBS           2.848 0.398 24   7.164  <.0001
##
## Results are given on the log2 (not the response) scale.
```

```
p.adjust(a6SE_tPCNA$contrasts$p.value[c(1,3,4,6,7,9,10,12)], method="fdr")
```

```
## [1] 3.648890e-01 4.974168e-01 1.025604e-01 2.634396e-03 4.677964e-03
## [6] 4.974168e-01 5.259494e-04 1.682570e-06
```

```
a6SE_tE2F1 <- test(emmeans(a6SE_E2F1, pairwise~Chemical|Time), adjust="none")
a6SE_tE2F1
```

```
## $emmeans
## Time = 24:
##  Chemical emmean    SE df t.ratio p.value
##  DCA       0.835 0.359 24   2.325  0.0289
##  DCA+FGF2  0.840 0.359 24   2.339  0.0280
##  FGF2      1.600 0.359 24   4.453  0.0002
##  PBS       0.000 0.359 24   0.000  1.0000
##
## Time = 48:
##  Chemical emmean    SE df t.ratio p.value
##  DCA       0.573 0.359 24   1.596  0.1235
##  DCA+FGF2  2.464 0.359 24   6.857  <.0001
##  FGF2      3.907 0.359 24  10.875  <.0001
##  PBS       0.534 0.359 24   1.485  0.1505
##
## Results are given on the log2 (not the response) scale.
##
## $contrasts
## Time = 24:
##  contrast           estimate    SE df t.ratio p.value
##  DCA - (DCA+FGF2)  -0.00493 0.508 24  -0.010  0.9923
##  DCA - FGF2        -0.76451 0.508 24  -1.505  0.1455
##  DCA - PBS          0.83533 0.508 24   1.644  0.1132
##  (DCA+FGF2) - FGF2 -0.75958 0.508 24  -1.495  0.1480
##  (DCA+FGF2) - PBS   0.84026 0.508 24   1.654  0.1112
##  FGF2 - PBS         1.59984 0.508 24   3.149  0.0043
```

```
##
## Time = 48:
##  contrast           estimate    SE df t.ratio p.value
##  DCA - (DCA+FGF2)  -1.89020 0.508 24  -3.720  0.0011
##  DCA - FGF2        -3.33351 0.508 24  -6.561  <.0001
##  DCA - PBS          0.03983 0.508 24   0.078  0.9382
##  (DCA+FGF2) - FGF2 -1.44331 0.508 24  -2.841  0.0090
##  (DCA+FGF2) - PBS   1.93003 0.508 24   3.799  0.0009
##  FGF2 - PBS         3.37334 0.508 24   6.639  <.0001
##
## Results are given on the log2 (not the response) scale.
```

```
p.adjust(a6SE_tE2F1$contrasts$p.value[c(1,3,4,6,7,9,10,12)], method="fdr")
```

```
## [1] 9.923415e-01 1.811331e-01 1.972829e-01 1.159376e-02 4.261042e-03
## [6] 9.923415e-01 1.806711e-02 5.788118e-06
```

### Experiment 5 (Figures 4E, S6D)

The same sort of responses as Experiment 2, with the same treatment structure as Experiment 4. We have 8 treatments, 6 replications/treatment. However, we analyze at 48h (Fig 4E) and 24h (Fig S6D) separately.

Comparisons for Figure 4E, S6D: PBS vs FGF2; PBS vs DCA; FGF2 vs FGF2+DCA, and DCA vs FGF2+DCA for EdU and pHH3.

#### Data and Visualization

```
d4E <- read.xlsx(xlsxFile = "MetabolismPaperRawData_Byran_working.xlsx",
                 colNames = TRUE, sheet="Fig 4_E")
glimpse(d4E)
```

```
## Rows: 24
## Columns: 8
## $ Chemical      <chr> "PBS", "PBS", "PBS", "PBS", "PBS", "PBS", "FGF2", "FGF2"~
## $ Sample        <dbl> 1, 2, 3, 4, 5, 6, 1, 2, 3, 4, 5, 6, 1, 2, 3, 4, 5, 6, 1,~
## $ `Time.(h)`    <dbl> 48, 48, 48, 48, 48, 48, 48, 48, 48, 48, 48, 48, 48, 48, ~
## $ `Edu/DAPI`    <dbl> 0.04411765, 0.03508772, 0.02564103, 0.02702703, 0.041666~
## $ `pHH3/DAPI`   <dbl> 0.05882353, 0.03508772, 0.02564103, 0.02702703, 0.027777~
## $ `Edu+.Cells`  <dbl> 3, 2, 2, 2, 3, 1, 14, 34, 23, 25, 17, 18, 3, 0, 1, 3, 0,~
## $ `DAPI+.Cells` <dbl> 68, 57, 78, 74, 72, 119, 134, 270, 140, 127, 136, 176, 9~
## $ `pHH3+.Cells` <dbl> 4, 2, 2, 2, 2, 1, 13, 30, 16, 11, 5, 16, 0, 0, 0, 0, 0, ~
```

```
dS6D <- read.xlsx(xlsxFile = "MetabolismPaperRawData_Byran_working.xlsx",
                  colNames = TRUE, sheet="Fig S6_D")
glimpse(dS6D)
```

```
## Rows: 24
## Columns: 8
## $ Chemical      <chr> "PBS", "PBS", "PBS", "PBS", "PBS", "PBS", "FGF2", "FGF2"~
```

```
## $ Sample        <dbl> 1, 2, 3, 4, 5, 6, 1, 2, 3, 4, 5, 6, 1, 2, 3, 4, 5, 6, 1,~
## $ `Time.(h)`    <dbl> 24, 24, 24, 24, 24, 24, 24, 24, 24, 24, 24, 24, 24, 24, ~
## $ `Edu/DAPI`    <dbl> 0.00000000, 0.01282051, 0.00000000, 0.00000000, 0.021052~
## $ `pHH3/DAPI`   <dbl> 0.00000000, 0.00000000, 0.00000000, 0.00000000, 0.000000~
## $ `Edu+.Cells`  <dbl> 0, 1, 0, 0, 2, 2, 10, 18, 10, 13, 16, 19, 0, 1, 1, 0, 0,~
## $ `DAPI+.Cells` <dbl> 99, 78, 52, 87, 95, 91, 119, 97, 85, 93, 105, 99, 71, 68~
## $ `pHH3+.Cells` <dbl> 0, 0, 0, 0, 0, 0, 3, 2, 9, 2, 6, 1, 0, 0, 0, 0, 0, 1, 0,~
```

```
d4E_long <- d4E %>%
  rename(Time=`Time.(h)`, Edu_DAPI_ratio=`Edu/DAPI`, pHH3_DAPI_ratio=`pHH3/DAPI`,
         Edu=`Edu+.Cells`, DAPI=`DAPI+.Cells`, pHH3=`pHH3+.Cells`) %>%
  pivot_longer(cols=4:5,names_to="RatioType",values_to="Ratio") %>%
  mutate(Time=as.factor(Time))
glimpse(d4E_long)
```

```
## Rows: 48
## Columns: 8
## $ Chemical  <chr> "PBS", "PBS", "PBS", "PBS", "PBS", "PBS", "PBS", "PBS", "PBS~
## $ Sample    <dbl> 1, 1, 2, 2, 3, 3, 4, 4, 5, 5, 6, 6, 1, 1, 2, 2, 3, 3, 4, 4, ~
## $ Time      <fct> 48, 48, 48, 48, 48, 48, 48, 48, 48, 48, 48, 48, 48, 48, 48, ~
## $ Edu       <dbl> 3, 3, 2, 2, 2, 2, 2, 2, 3, 3, 1, 1, 14, 14, 34, 34, 23, 23, ~
## $ DAPI      <dbl> 68, 68, 57, 57, 78, 78, 74, 74, 72, 72, 119, 119, 134, 134, ~
## $ pHH3      <dbl> 4, 4, 2, 2, 2, 2, 2, 2, 2, 2, 1, 1, 13, 13, 30, 30, 16, 16, ~
## $ RatioType <chr> "Edu_DAPI_ratio", "pHH3_DAPI_ratio", "Edu_DAPI_ratio", "pHH3~
## $ Ratio     <dbl> 0.04411765, 0.05882353, 0.03508772, 0.03508772, 0.02564103, ~
```

```
dS6D_long <- dS6D %>%
  rename(Time=`Time.(h)`, Edu_DAPI_ratio=`Edu/DAPI`, pHH3_DAPI_ratio=`pHH3/DAPI`,
         Edu=`Edu+.Cells`, DAPI=`DAPI+.Cells`, pHH3=`pHH3+.Cells`) %>%
  pivot_longer(cols=4:5,names_to="RatioType",values_to="Ratio") %>%
  mutate(Time=as.factor(Time))
glimpse(dS6D_long)
```

```
## Rows: 48
## Columns: 8
## $ Chemical  <chr> "PBS", "PBS", "PBS", "PBS", "PBS", "PBS", "PBS", "PBS", "PBS~
## $ Sample    <dbl> 1, 1, 2, 2, 3, 3, 4, 4, 5, 5, 6, 6, 1, 1, 2, 2, 3, 3, 4, 4, ~
## $ Time      <fct> 24, 24, 24, 24, 24, 24, 24, 24, 24, 24, 24, 24, 24, 24, 24, ~
## $ Edu       <dbl> 0, 0, 1, 1, 0, 0, 0, 0, 2, 2, 2, 2, 10, 10, 18, 18, 10, 10, ~
## $ DAPI      <dbl> 99, 99, 78, 78, 52, 52, 87, 87, 95, 95, 91, 91, 119, 119, 97~
## $ pHH3      <dbl> 0, 0, 0, 0, 0, 0, 0, 0, 0, 0, 0, 0, 3, 3, 2, 2, 9, 9, 2, 2, ~
## $ RatioType <chr> "Edu_DAPI_ratio", "pHH3_DAPI_ratio", "Edu_DAPI_ratio", "pHH3~
## $ Ratio     <dbl> 0.00000000, 0.00000000, 0.01282051, 0.00000000, 0.00000000, ~
```

```
ggplot(d4E_long, aes(x=Chemical,y=log(Ratio),color=Chemical)) +
   geom_jitter(width=.05,height=0) +
   facet_wrap(~RatioType, nrow=1) +
   labs(title="For Figure 4E: Treatment Comparisons by Time")
```

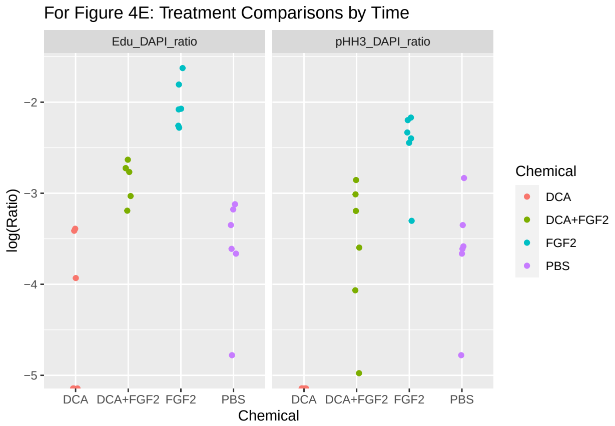

```r
ggplot(dS6D_long, aes(x=Chemical,y=log(Ratio),color=Chemical)) +
   geom_jitter(width=.05,height=0) +
   facet_wrap(~RatioType, nrow=1) +
   labs(title="For Figure S6D: Treatment Comparisons by Time")
```

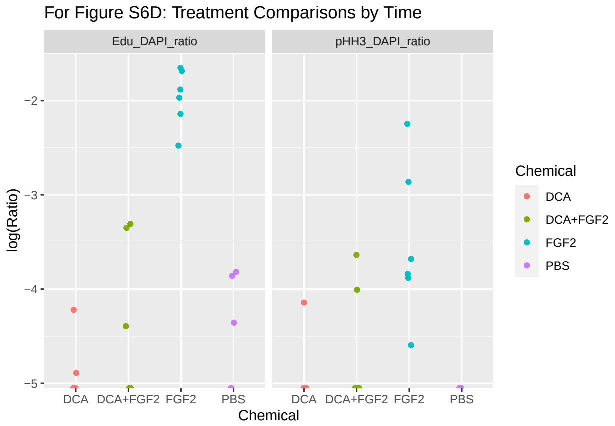

#### Data Analysis (Experiment 5)

```r
d4E <- d4E %>%
  rename(Time=`Time.(h)`, Edu_DAPI_ratio=`Edu/DAPI`, pHH3_DAPI_ratio=`pHH3/DAPI`,
         Edu=`Edu+.Cells`, DAPI=`DAPI+.Cells`, pHH3=`pHH3+.Cells`) %>%
  mutate(Time=as.factor(Time))
glimpse(d4E)
```

```
## Rows: 24
## Columns: 8
## $ Chemical        <chr> "PBS", "PBS", "PBS", "PBS", "PBS", "PBS", "FGF2", "FGF~
## $ Sample          <dbl> 1, 2, 3, 4, 5, 6, 1, 2, 3, 4, 5, 6, 1, 2, 3, 4, 5, 6, ~
## $ Time            <fct> 48, 48, 48, 48, 48, 48, 48, 48, 48, 48, 48, 48, 48, 48~
## $ Edu_DAPI_ratio  <dbl> 0.04411765, 0.03508772, 0.02564103, 0.02702703, 0.0416~
## $ pHH3_DAPI_ratio <dbl> 0.05882353, 0.03508772, 0.02564103, 0.02702703, 0.0277~
## $ Edu             <dbl> 3, 2, 2, 2, 3, 1, 14, 34, 23, 25, 17, 18, 3, 0, 1, 3, ~
## $ DAPI            <dbl> 68, 57, 78, 74, 72, 119, 134, 270, 140, 127, 136, 176,~
## $ pHH3            <dbl> 4, 2, 2, 2, 2, 1, 13, 30, 16, 11, 5, 16, 0, 0, 0, 0, 0~
```

```r
# because pHH3 has all 0 counts for DCA
d4E.p <- d4E %>%
  filter(Chemical != "DCA")
  glimpse(d4E.p)
```

```
## Rows: 18
## Columns: 8
## $ Chemical        <chr> "PBS", "PBS", "PBS", "PBS", "PBS", "PBS", "FGF2", "FGF~
## $ Sample          <dbl> 1, 2, 3, 4, 5, 6, 1, 2, 3, 4, 5, 6, 1, 2, 3, 4, 5, 6
## $ Time            <fct> 48, 48, 48, 48, 48, 48, 48, 48, 48, 48, 48, 48, 48, 48~
## $ Edu_DAPI_ratio  <dbl> 0.04411765, 0.03508772, 0.02564103, 0.02702703, 0.0416~
## $ pHH3_DAPI_ratio <dbl> 0.05882353, 0.03508772, 0.02564103, 0.02702703, 0.0277~
## $ Edu             <dbl> 3, 2, 2, 2, 3, 1, 14, 34, 23, 25, 17, 18, 3, 7, 10, 11~
## $ DAPI            <dbl> 68, 57, 78, 74, 72, 119, 134, 270, 140, 127, 136, 176,~
## $ pHH3            <dbl> 4, 2, 2, 2, 2, 1, 13, 30, 16, 11, 5, 16, 2, 1, 8, 3, 5~
```

```r
dS6D <- dS6D %>%
  rename(Time=`Time.(h)`, Edu_DAPI_ratio=`Edu/DAPI`, pHH3_DAPI_ratio=`pHH3/DAPI`,
         Edu=`Edu+.Cells`, DAPI=`DAPI+.Cells`, pHH3=`pHH3+.Cells`) %>%
  mutate(Time=as.factor(Time))
glimpse(dS6D)
```

```
## Rows: 24
## Columns: 8
## $ Chemical        <chr> "PBS", "PBS", "PBS", "PBS", "PBS", "PBS", "FGF2", "FGF~
## $ Sample          <dbl> 1, 2, 3, 4, 5, 6, 1, 2, 3, 4, 5, 6, 1, 2, 3, 4, 5, 6, ~
## $ Time            <fct> 24, 24, 24, 24, 24, 24, 24, 24, 24, 24, 24, 24, 24, 24~
## $ Edu_DAPI_ratio  <dbl> 0.00000000, 0.01282051, 0.00000000, 0.00000000, 0.0210~
## $ pHH3_DAPI_ratio <dbl> 0.00000000, 0.00000000, 0.00000000, 0.00000000, 0.0000~
## $ Edu             <dbl> 0, 1, 0, 0, 2, 2, 10, 18, 10, 13, 16, 19, 0, 1, 1, 0, ~
## $ DAPI            <dbl> 99, 78, 52, 87, 95, 91, 119, 97, 85, 93, 105, 99, 71, ~
## $ pHH3            <dbl> 0, 0, 0, 0, 0, 0, 3, 2, 9, 2, 6, 1, 0, 0, 0, 0, 0, 1, ~
```

```r
# because pHH3 has all 0 counts for PBS
dS6D.p <- dS6D %>%
  filter(Chemical != "PBS")
  glimpse(dS6D.p)
```

```
## Rows: 18
## Columns: 8
## $ Chemical        <chr> "FGF2", "FGF2", "FGF2", "FGF2", "FGF2", "FGF2", "DCA",~
## $ Sample          <dbl> 1, 2, 3, 4, 5, 6, 1, 2, 3, 4, 5, 6, 1, 2, 3, 4, 5, 6
## $ Time            <fct> 24, 24, 24, 24, 24, 24, 24, 24, 24, 24, 24, 24, 24, 24~
## $ Edu_DAPI_ratio  <dbl> 0.08403361, 0.18556701, 0.11764706, 0.13978495, 0.1523~
## $ pHH3_DAPI_ratio <dbl> 0.02521008, 0.02061856, 0.10588235, 0.02150538, 0.0571~
## $ Edu             <dbl> 10, 18, 10, 13, 16, 19, 0, 1, 1, 0, 0, 0, 0, 3, 1, 0, ~
## $ DAPI            <dbl> 119, 97, 85, 93, 105, 99, 71, 68, 133, 83, 77, 63, 97,~
## $ pHH3            <dbl> 3, 2, 9, 2, 6, 1, 0, 0, 0, 0, 0, 1, 0, 0, 0, 2, 3, 0
```

```r
nb.Edu.4E <- glm(Edu~Chemical + offset(log(DAPI)), family=poisson(), data=d4E)
summary(nb.Edu.4E) # poisson because theta estimate is large
```

```
##
## Call:
## glm(formula = Edu ~ Chemical + offset(log(DAPI)), family = poisson(),
##     data = d4E)
##
## Deviance Residuals:
##      Min        1Q    Median        3Q       Max
## -2.96117  -0.71253   0.05329   0.67940   2.29686
##
## Coefficients:
##                   Estimate Std. Error z value Pr(>|z|)
## (Intercept)        -5.0858     0.3779 -13.457  < 2e-16 ***
## ChemicalDCA+FGF2   2.2747     0.4076   5.581 2.38e-08 ***
## ChemicalFGF2       3.0704     0.3879   7.915 2.47e-15 ***
## ChemicalPBS        1.5023     0.4688   3.205  0.00135 **
## ---
## Signif. codes:  0 '***' 0.001 '**' 0.01 '*' 0.05 '.' 0.1 ' ' 1
##
## (Dispersion parameter for poisson family taken to be 1)
##
##     Null deviance: 198.999  on 23  degrees of freedom
## Residual deviance:  33.876  on 20  degrees of freedom
## AIC: 117.42
##
## Number of Fisher Scoring iterations: 6
```

```
nb.pHH3.4E <- glm.nb(pHH3~Chemical + offset(log(DAPI)), data=d4E.p)
summary(nb.pHH3.4E) # without DCA because all 0 counts
```

```
##
## Call:
## glm.nb(formula = pHH3 ~ Chemical + offset(log(DAPI)), data = d4E.p,
##     init.theta = 79.43628861, link = log)
##
## Deviance Residuals:
##      Min        1Q    Median        3Q       Max
## -2.28299  -0.18269  -0.02288   0.70109   1.54480
##
## Coefficients:
##               Estimate Std. Error z value Pr(>|z|)
## (Intercept)    -3.4779     0.2182 -15.936  < 2e-16 ***
## ChemicalFGF2    1.0913     0.2469   4.420 9.86e-06 ***
## ChemicalPBS    -0.1024     0.3557  -0.288    0.773
## ---
## Signif. codes:  0 '***' 0.001 '**' 0.01 '*' 0.05 '.' 0.1 ' ' 1
##
## (Dispersion parameter for Negative Binomial(79.4363) family taken to be 1)
##
##     Null deviance: 52.145  on 17  degrees of freedom
## Residual deviance: 18.958  on 15  degrees of freedom
## AIC: 88.79
##
## Number of Fisher Scoring iterations: 1
##
```

```
##
##                Theta:  79
##            Std. Err.:  338
##
##  2 x log-likelihood:  -80.79
```

```
nb.Edu.S6D <- glm.nb(Edu~Chemical + offset(log(DAPI)),data=dS6D)
summary(nb.Edu.S6D)
```

```
##
## Call:
## glm.nb(formula = Edu ~ Chemical + offset(log(DAPI)), data = dS6D,
##     init.theta = 73.33304892, link = log)
##
## Deviance Residuals:
##     Min       1Q   Median       3Q      Max
## -1.7245  -1.0899  -0.3596   0.9346   1.6076
##
## Coefficients:
##                   Estimate Std. Error z value Pr(>|z|)
## (Intercept)        -5.5120     0.7091  -7.774 7.62e-15 ***
## ChemicalDCA+FGF2    1.2184     0.7938   1.535    0.125
## ChemicalFGF2        3.5748     0.7188   4.973 6.58e-07 ***
## ChemicalPBS         0.9024     0.8398   1.075    0.283
## ---
## Signif. codes:  0 '***' 0.001 '**' 0.01 '*' 0.05 '.' 0.1 ' ' 1
##
## (Dispersion parameter for Negative Binomial(73.333) family taken to be 1)
##
##     Null deviance: 169.742  on 23  degrees of freedom
## Residual deviance:  28.885  on 20  degrees of freedom
## AIC: 86.52
##
## Number of Fisher Scoring iterations: 1
##
##
##                Theta:  73
##            Std. Err.:  271
##
##  2 x log-likelihood:  -76.52
```

```
nb.pHH3.S6D <- glm.nb(pHH3~Chemical + offset(log(DAPI)),data=dS6D.p)
summary(nb.pHH3.S6D) # without PBS because all 0 counts
```

```
##
## Call:
## glm.nb(formula = pHH3 ~ Chemical + offset(log(DAPI)), data = dS6D.p,
##     init.theta = 2.636715005, link = log)
##
## Deviance Residuals:
##     Min       1Q   Median       3Q      Max
## -1.2419  -0.9998  -0.5652   0.2555   1.5245
##
```

```
## Coefficients:
##                   Estimate Std. Error z value Pr(>|z|)
## (Intercept)         -6.186      1.024  -6.039 1.55e-09 ***
## ChemicalDCA+FGF2     1.390      1.149   1.210  0.22625
## ChemicalFGF2         2.951      1.075   2.746  0.00603 **
## ---
## Signif. codes:  0 '***' 0.001 '**' 0.01 '*' 0.05 '.' 0.1 ' ' 1
##
## (Dispersion parameter for Negative Binomial(2.6367) family taken to be 1)
##
##     Null deviance: 33.571  on 17  degrees of freedom
## Residual deviance: 16.879  on 15  degrees of freedom
## AIC: 57.217
##
## Number of Fisher Scoring iterations: 1
##
##
##               Theta:  2.64
##           Std. Err.:  2.53
##
##  2 x log-likelihood:  -49.217
```

```
# for nb.Edu.4E, with all four treatments
K <- matrix(c(0,0,-1,1, # PBS vs. FGF2
              0,0,0,1, # PBS vs. DCA
              0,-1,1,0, # FGF2 vs. FGF2+DCA
              0,-1,0,0),nrow=4, byrow=TRUE) # DCA vs. FGF2+DCA

t <- glht(nb.Edu.4E, linfct=K)
summary(t, test=adjusted("fdr"))
```

```
##
##   Simultaneous Tests for General Linear Hypotheses
##
## Fit: glm(formula = Edu ~ Chemical + offset(log(DAPI)), family = poisson(),
##     data = d4E)
##
## Linear Hypotheses:
##        Estimate Std. Error z value Pr(>|z|)
## 1 == 0  -1.5681     0.2908  -5.393 1.39e-07 ***
## 2 == 0   1.5023     0.4688   3.205  0.00135 **
## 3 == 0   0.7957     0.1758   4.527 7.97e-06 ***
## 4 == 0  -2.2747     0.4076  -5.581 9.54e-08 ***
## ---
## Signif. codes:  0 '***' 0.001 '**' 0.01 '*' 0.05 '.' 0.1 ' ' 1
## (Adjusted p values reported -- fdr method)
```

```
t <- glht(nb.Edu.S6D, linfct=K)
summary(t, test=adjusted("fdr"))
```

```
##
##   Simultaneous Tests for General Linear Hypotheses
##
```

```
## Fit: glm.nb(formula = Edu ~ Chemical + offset(log(DAPI)), data = dS6D,
##     init.theta = 73.33304892, link = log)
##
## Linear Hypotheses:
##        Estimate Std. Error z value Pr(>|z|)
## 1 == 0  -2.6724     0.4651  -5.746 1.83e-08 ***
## 2 == 0   0.9024     0.8398   1.075    0.283
## 3 == 0   2.3564     0.3757   6.271 1.43e-09 ***
## 4 == 0  -1.2184     0.7938  -1.535    0.166
## ---
## Signif. codes:  0 '***' 0.001 '**' 0.01 '*' 0.05 '.' 0.1 ' ' 1
## (Adjusted p values reported -- fdr method)
```

```
# For pHH3 Fig 4E only can compare the first and third contrasts, because no DCA
K2 <- matrix(c(0,-1,1, # PBS vs. FGF2
              0,1,0 # FGF2 vs. FGF2+DCA
              ),nrow=2, byrow=TRUE)
t <- glht(nb.pHH3.4E, linfct=K2)
summary(t, test=adjusted("fdr"))
```

```
##
##   Simultaneous Tests for General Linear Hypotheses
##
## Fit: glm.nb(formula = pHH3 ~ Chemical + offset(log(DAPI)), data = d4E.p,
##     init.theta = 79.43628861, link = log)
##
## Linear Hypotheses:
##        Estimate Std. Error z value Pr(>|z|)
## 1 == 0  -1.1937     0.3037  -3.931 8.47e-05 ***
## 2 == 0   1.0913     0.2469   4.420 1.97e-05 ***
## ---
## Signif. codes:  0 '***' 0.001 '**' 0.01 '*' 0.05 '.' 0.1 ' ' 1
## (Adjusted p values reported -- fdr method)
```

```
# For pHH3 Fig S6D only can compare the first and third contrasts, because no PBS
K3 <- matrix(c(0,-1,1, # FGF2 vs. FGF2+DCA
              0,-1,0),nrow=2, byrow=TRUE) # DCA vs. FGF2+DCA

t <- glht(nb.pHH3.S6D, linfct=K3)
summary(t, test=adjusted("fdr"))
```

```
##
##   Simultaneous Tests for General Linear Hypotheses
##
## Fit: glm.nb(formula = pHH3 ~ Chemical + offset(log(DAPI)), data = dS6D.p,
##     init.theta = 2.636715005, link = log)
##
## Linear Hypotheses:
##        Estimate Std. Error z value Pr(>|z|)
## 1 == 0   1.5612     0.6135   2.545   0.0219 *
## 2 == 0  -1.3901     1.1488  -1.210   0.2263
## ---
## Signif. codes:  0 '***' 0.001 '**' 0.01 '*' 0.05 '.' 0.1 ' ' 1
## (Adjusted p values reported -- fdr method)
```

#### Experiment 6 (Figures 4G, S6G)

The responses are ratios that measure the intensity of the pixels of images in Fig 4F and S6F, using the same treatment structure as Experiments 4 and 5. We have 8 treatments and 3 reps/treatment.

Comparisons for Figure 4G: PBS vs FGF2; PBS vs DCA; FGF2 vs FGF2+DCA, and DCA vs FGF2+DCA for pPDH/PDH, Tubulin/actin, and pERK/ERK

Comparisons for Figure S6G: PBS vs FGF2; PBS vs DCA; FGF2 vs FGF2+DCA, and DCA vs FGF2+DCA for pPDH/PDH and pERK/ERK

The analysis is done for each time period, and the data is split into two figures. Figure 4G shows the results at 48 hours, whereas Figure S6G shows the results at 24 hours. This is because there were different Western Blots for each time point, and there could be significant variation across the different blots. Furthermore, in both parts they measured pPDH/PDH and pERK/ERK, but they only measured Tubulin/Actin in the 48h part (Tubulin wasn't detectable at 24h).

##### Data and Visualization

```
d4G <- read.xlsx(xlsxFile = "MetabolismPaperRawData_Byran_working.xlsx",
                 colNames = TRUE, sheet="Fig 4_G")
glimpse(d4G)
```

```
## Rows: 12
## Columns: 14
## $ Chemical        <chr> "PBS", "PBS", "PBS", "DCA", "DCA", "DCA", "FGF2", "FGF~
## $ FGF2            <dbl> 0, 0, 0, 0, 0, 0, 1, 1, 1, 1, 1, 1
## $ DCA             <dbl> 0, 0, 0, 1, 1, 1, 0, 0, 0, 1, 1, 1
## $ Sample          <dbl> 1, 2, 3, 1, 2, 3, 1, 2, 3, 1, 2, 3
## $ `Time.(h)`      <dbl> 48, 48, 48, 48, 48, 48, 48, 48, 48, 48, 48, 48
## $ `pPDH/PDH`      <dbl> 0.42006887, 0.71284963, 1.00000000, 0.03587164, 0.0640~
## $ `Tubulin/Actin` <dbl> 1.0000000, 0.5995649, 0.6949761, 0.7805529, 0.4193759,~
## $ `pERK/ERK`      <dbl> 0.3119960, 0.7643253, 1.0000000, 1.7416681, 1.4339798,~
## $ pPDH            <dbl> 3750400, 7592576, 4084096, 294784, 459648, 609920, 582~
## $ PHD             <dbl> 69423915, 82821526, 31757620, 63900558, 55760874, 3041~
## $ Tubulin         <dbl> 25296, 16244, 13144, 19840, 12896, 5828, 90272, 58280,~
## $ Actin           <dbl> 43809684, 46921843, 32754873, 44020768, 53256153, 5262~
## $ pERK            <dbl> 5337779, 9165888, 4707572, 23706720, 26409715, 2179515~
## $ ERK             <dbl> 23196500, 16259500, 6382750, 18455125, 24970750, 17717~
```

```
dS6G <- read.xlsx(xlsxFile = "MetabolismPaperRawData_Byran_working.xlsx",
                  colNames = TRUE, sheet="Fig S6_G")
glimpse(dS6G)
```

```
## Rows: 12
## Columns: 11
## $ Chemical   <chr> "PBS", "PBS", "PBS", "DCA", "DCA", "DCA", "FGF2", "FGF2", "~
## $ FGF2       <dbl> 0, 0, 0, 0, 0, 0, 1, 1, 1, 1, 1, 1
## $ DCA        <dbl> 0, 0, 0, 1, 1, 1, 0, 0, 0, 1, 1, 1
## $ Sample     <dbl> 1, 2, 3, 1, 2, 3, 1, 2, 3, 1, 2, 3
## $ `Time.(h)` <dbl> 24, 24, 24, 24, 24, 24, 24, 24, 24, 24, 24, 24
## $ `pPDH/PDH` <dbl> 0.210653346, 0.576908267, 1.000000000, 0.078264713, 0.00988~
```

```
## $ `pERK/ERK` <dbl> 1.0000000, 0.9128367, 0.7796046, 1.3157399, 0.6913616, 2.14~
## $ pPDH       <dbl> 37536247, 71371841, 188181488, 20036409, 1422781, 3343402, ~
## $ PHD        <dbl> 108053504, 75019904, 114112512, 155242368, 87260160, 176947~
## $ pERK       <dbl> 53455697, 61614177, 69263387, 95630746, 39240460, 127646938~
## $ ERK        <dbl> 6019785, 7601067, 10004982, 8184921, 6391692, 6687489, 1306~
```

```
# d4G_S6G <- bind_rows(d4G,dS6G)
# kable(d4G_S6G)
```

```
d4G_long <- d4G %>%
  rename(Time=`Time.(h)`, pPDH_PDH_ratio=`pPDH/PDH`, pERK_ERK_ratio=`pERK/ERK`, Tubulin_Actin_ratio=`Tu
  pivot_longer(cols=6:8,names_to="Type",values_to="Ratio") %>%
  mutate(Time=as.factor(Time))
glimpse(d4G_long)
```

```
## Rows: 36
## Columns: 13
## $ Chemical <chr> "PBS", "PBS", "PBS", "PBS", "PBS", "PBS", "PBS", "PBS", "PBS"~
## $ FGF2     <dbl> 0, 0, 0, 0, 0, 0, 0, 0, 0, 0, 0, 0, 0, 0, 0, 0, 0, 0, 1, 1, 1~
## $ DCA      <dbl> 0, 0, 0, 0, 0, 0, 0, 0, 0, 1, 1, 1, 1, 1, 1, 1, 1, 1, 0, 0, 0~
## $ Sample   <dbl> 1, 1, 1, 2, 2, 2, 3, 3, 3, 1, 1, 1, 2, 2, 2, 3, 3, 3, 1, 1, 1~
## $ Time     <fct> 48, 48, 48, 48, 48, 48, 48, 48, 48, 48, 48, 48, 48, 48, 48, 4~
## $ pPDH     <dbl> 3750400, 3750400, 3750400, 7592576, 7592576, 7592576, 4084096~
## $ PHD      <dbl> 69423915, 69423915, 69423915, 82821526, 82821526, 82821526, 3~
## $ Tubulin  <dbl> 25296, 25296, 25296, 16244, 16244, 16244, 13144, 13144, 13144~
## $ Actin    <dbl> 43809684, 43809684, 43809684, 46921843, 46921843, 46921843, 3~
## $ pERK     <dbl> 5337779, 5337779, 5337779, 9165888, 9165888, 9165888, 4707572~
## $ ERK      <dbl> 23196500, 23196500, 23196500, 16259500, 16259500, 16259500, 6~
## $ Type     <chr> "pPDH_PDH_ratio", "Tubulin_Actin_ratio", "pERK_ERK_ratio", "p~
## $ Ratio    <dbl> 0.42006887, 1.00000000, 0.31199601, 0.71284963, 0.59956488, 0~
```

```
dS6G_long <- dS6G %>%
  rename(Time=`Time.(h)`, pPDH_PDH_ratio=`pPDH/PDH`, pERK_ERK_ratio=`pERK/ERK`) %>%
  pivot_longer(cols=6:7,names_to="Type",values_to="Ratio") %>%
  mutate(Time=as.factor(Time))
glimpse(dS6G_long)
```

```
## Rows: 24
## Columns: 11
## $ Chemical <chr> "PBS", "PBS", "PBS", "PBS", "PBS", "PBS", "DCA", "DCA", "DCA"~
## $ FGF2     <dbl> 0, 0, 0, 0, 0, 0, 0, 0, 0, 0, 0, 0, 1, 1, 1, 1, 1, 1, 1, 1, 1~
## $ DCA      <dbl> 0, 0, 0, 0, 0, 0, 1, 1, 1, 1, 1, 1, 0, 0, 0, 0, 0, 0, 1, 1, 1~
## $ Sample   <dbl> 1, 1, 2, 2, 3, 3, 1, 1, 2, 2, 3, 3, 1, 1, 2, 2, 3, 3, 1, 1, 2~
## $ Time     <fct> 24, 24, 24, 24, 24, 24, 24, 24, 24, 24, 24, 24, 24, 24, 24, 2~
## $ pPDH     <dbl> 37536247, 37536247, 71371841, 71371841, 188181488, 188181488,~
## $ PHD      <dbl> 108053504, 108053504, 75019904, 75019904, 114112512, 11411251~
## $ pERK     <dbl> 53455697, 53455697, 61614177, 61614177, 69263387, 69263387, 9~
## $ ERK      <dbl> 6019785, 6019785, 7601067, 7601067, 10004982, 10004982, 81849~
## $ Type     <chr> "pPDH_PDH_ratio", "pERK_ERK_ratio", "pPDH_PDH_ratio", "pERK_E~
## $ Ratio    <dbl> 0.210653346, 1.000000000, 0.576908267, 0.912836664, 1.0000000~
```

```
ggplot(d4G_long, aes(x=Chemical,y=Ratio,color=Chemical)) +
  geom_jitter(width=.05,height=0) +
  facet_wrap(~Type, nrow=2) +
  labs(title="For Figure 4G: Treatment Comparisons (no log2)")
```

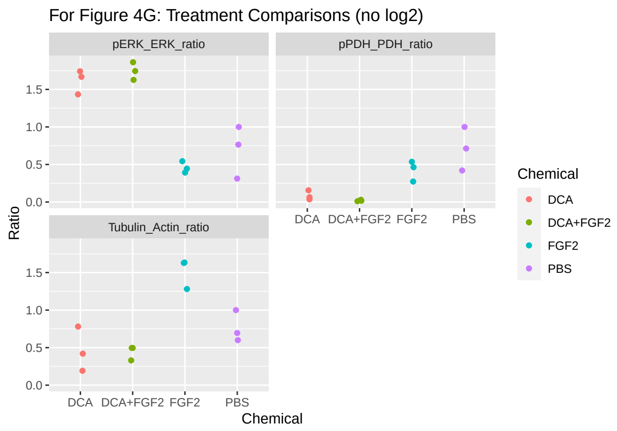

```
ggplot(dS6G_long, aes(x=Chemical,y=Ratio,color=Chemical)) +
  geom_jitter(width=.05,height=0) +
  facet_wrap(~Type, nrow=1) +
  labs(title="For Figure S6G: Treatment Comparisons (no log2)")
```

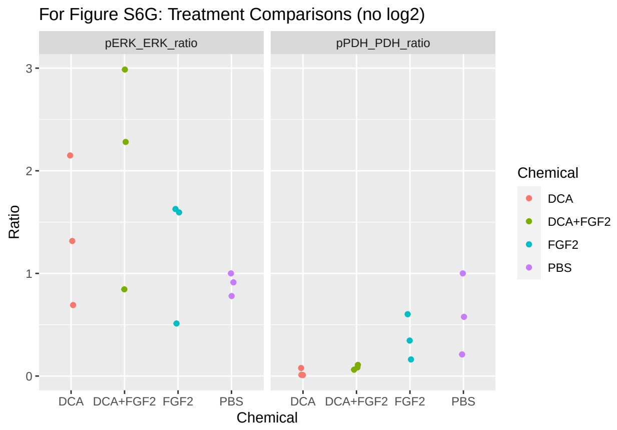

```
ggplot(d4G_long, aes(x=Chemical,y=log2(Ratio),color=Chemical)) +
  geom_jitter(width=.05,height=0) +
  facet_wrap(~Type, nrow=2) +
  labs(title="For Figure 4G: Treatment Comparisons")
```

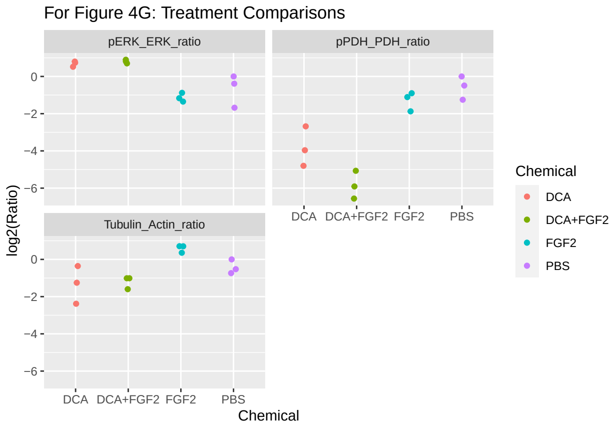

```r
ggplot(dS6G_long, aes(x=Chemical,y=log2(Ratio),color=Chemical)) +
  geom_jitter(width=.05,height=0) +
  facet_wrap(~Type, nrow=1) +
  labs(title="For Figure S6G: Treatment Comparisons")
```

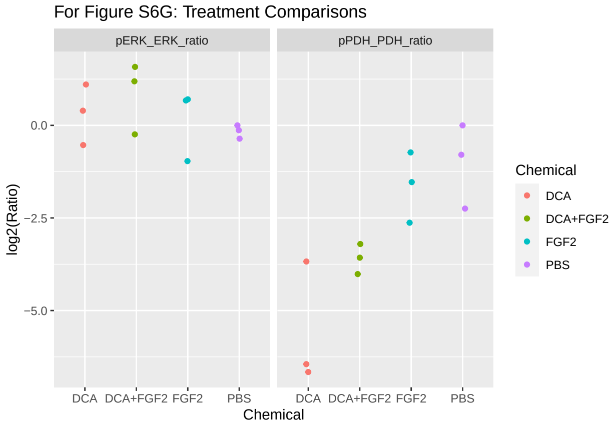

### Data Analysis (Experiment 6)

##### Analyzing $\log_2$ of the Ratios

```r
d4G <- d4G %>%
  rename(Time=`Time.(h)`, pPDH_PDH_ratio=`pPDH/PDH`, pERK_ERK_ratio=`pERK/ERK`,
         Tubulin_Actin_ratio=`Tubulin/Actin`) %>%
  mutate(Time=as.factor(Time))
glimpse(d4G)
```

```
## Rows: 12
## Columns: 14
## $ Chemical                 <chr> "PBS", "PBS", "PBS", "DCA", "DCA", "DCA", "FGF2", ~
## $ FGF2                     <dbl> 0, 0, 0, 0, 0, 0, 1, 1, 1, 1, 1, 1
## $ DCA                      <dbl> 0, 0, 0, 1, 1, 1, 0, 0, 0, 1, 1, 1
## $ Sample                   <dbl> 1, 2, 3, 1, 2, 3, 1, 2, 3, 1, 2, 3
## $ Time                     <fct> 48, 48, 48, 48, 48, 48, 48, 48, 48, 48, 48, 48
```

```
## $ pPDH_PDH_ratio      <dbl> 0.42006887, 0.71284963, 1.00000000, 0.03587164, 0.~
## $ Tubulin_Actin_ratio <dbl> 1.0000000, 0.5995649, 0.6949761, 0.7805529, 0.4193~
## $ pERK_ERK_ratio      <dbl> 0.3119960, 0.7643253, 1.0000000, 1.7416681, 1.4339~
## $ pPDH                <dbl> 3750400, 7592576, 4084096, 294784, 459648, 609920,~
## $ PHD                 <dbl> 69423915, 82821526, 31757620, 63900558, 55760874, ~
## $ Tubulin             <dbl> 25296, 16244, 13144, 19840, 12896, 5828, 90272, 58~
## $ Actin               <dbl> 43809684, 46921843, 32754873, 44020768, 53256153, ~
## $ pERK                <dbl> 5337779, 9165888, 4707572, 23706720, 26409715, 217~
## $ ERK                 <dbl> 23196500, 16259500, 6382750, 18455125, 24970750, 1~
```

```
dS6G <- dS6G %>%
  rename(Time=`Time.(h)`, pPDH_PDH_ratio=`pPDH/PDH`, pERK_ERK_ratio=`pERK/ERK`) %>%
  mutate(Time=as.factor(Time))
glimpse(dS6G)
```

```
## Rows: 12
## Columns: 11
## $ Chemical       <chr> "PBS", "PBS", "PBS", "DCA", "DCA", "DCA", "FGF2", "FGF2~
## $ FGF2           <dbl> 0, 0, 0, 0, 0, 0, 1, 1, 1, 1, 1, 1
## $ DCA            <dbl> 0, 0, 0, 1, 1, 1, 0, 0, 0, 1, 1, 1
## $ Sample         <dbl> 1, 2, 3, 1, 2, 3, 1, 2, 3, 1, 2, 3
## $ Time           <fct> 24, 24, 24, 24, 24, 24, 24, 24, 24, 24, 24, 24
## $ pPDH_PDH_ratio <dbl> 0.210653346, 0.576908267, 1.000000000, 0.078264713, 0.0~
## $ pERK_ERK_ratio <dbl> 1.0000000, 0.9128367, 0.7796046, 1.3157399, 0.6913616, ~
## $ pPDH           <dbl> 37536247, 71371841, 188181488, 20036409, 1422781, 33434~
## $ PHD            <dbl> 108053504, 75019904, 114112512, 155242368, 87260160, 17~
## $ pERK           <dbl> 53455697, 61614177, 69263387, 95630746, 39240460, 12764~
## $ ERK            <dbl> 6019785, 7601067, 10004982, 8184921, 6391692, 6687489, ~
```

```
a4G_PDH <- lm(log2(pPDH_PDH_ratio) ~ DCA*FGF2, data=d4G)
a4G_ERK <- lm(log2(pERK_ERK_ratio) ~ DCA*FGF2, data=d4G)
a4G_Tub <- lm(log2(Tubulin_Actin_ratio) ~ DCA*FGF2, data=d4G)

aS6G_PDH <- lm(log2(pPDH_PDH_ratio) ~ DCA*FGF2, data=dS6G)
aS6G_ERK <- lm(log2(pERK_ERK_ratio) ~ DCA*FGF2, data=dS6G)

summary(a4G_PDH)
```

```
##
## Call:
## lm(formula = log2(pPDH_PDH_ratio) ~ DCA * FGF2, data = d4G)
##
## Residuals:
##      Min       1Q   Median       3Q      Max
## -0.98581 -0.60272  0.01404  0.44039  1.13417
##
## Coefficients:
##             Estimate Std. Error t value Pr(>|t|)
## (Intercept)  -0.5799     0.4435  -1.307 0.227411
## DCA          -3.2353     0.6273  -5.158 0.000866 ***
## FGF2         -0.7150     0.6273  -1.140 0.287336
## DCA:FGF2     -1.3157     0.8871  -1.483 0.176310
## ---
```

```
## Signif. codes:  0 '***' 0.001 '**' 0.01 '*' 0.05 '.' 0.1 ' ' 1
##
## Residual standard error: 0.7682 on 8 degrees of freedom
## Multiple R-squared:  0.9174, Adjusted R-squared:  0.8864
## F-statistic: 29.61 on 3 and 8 DF,  p-value: 0.0001108
```

```r
plot(a4G_PDH,which=1:2)
```

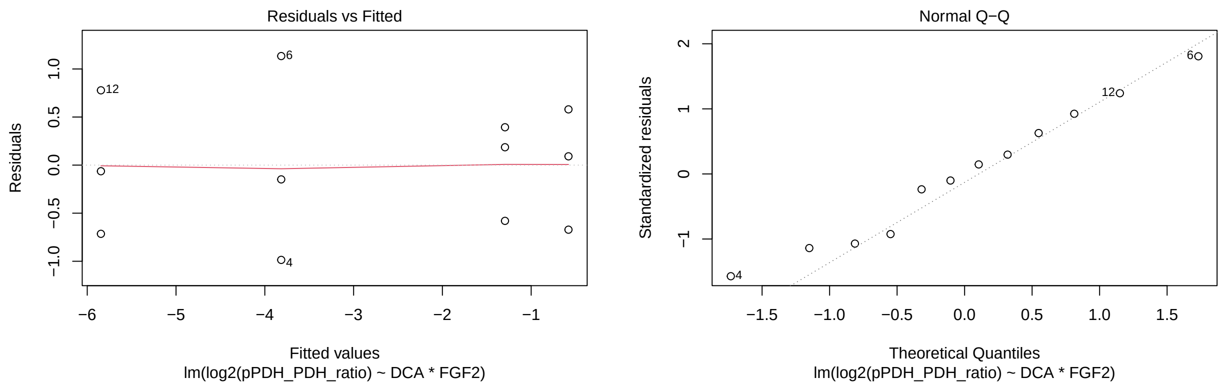

```r
plot(a4G_ERK,which=1:2)
```

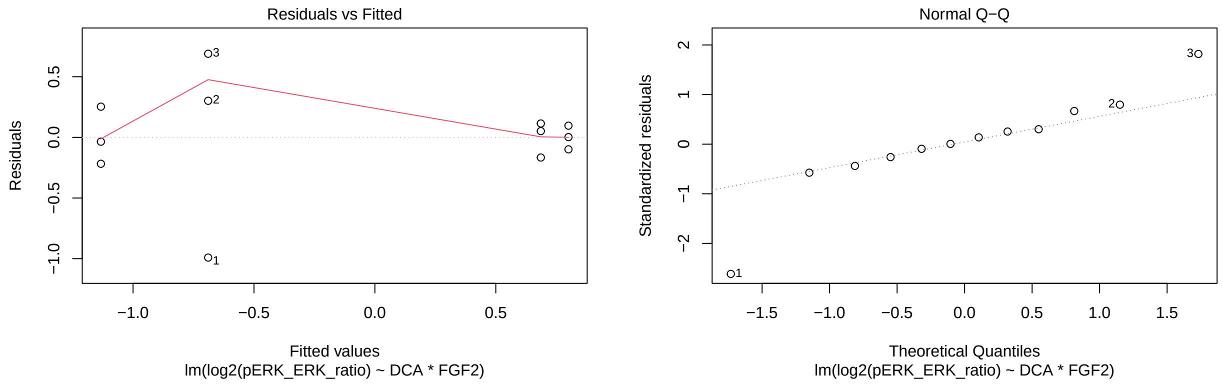

```r
plot(a4G_Tub,which=1:2)
```

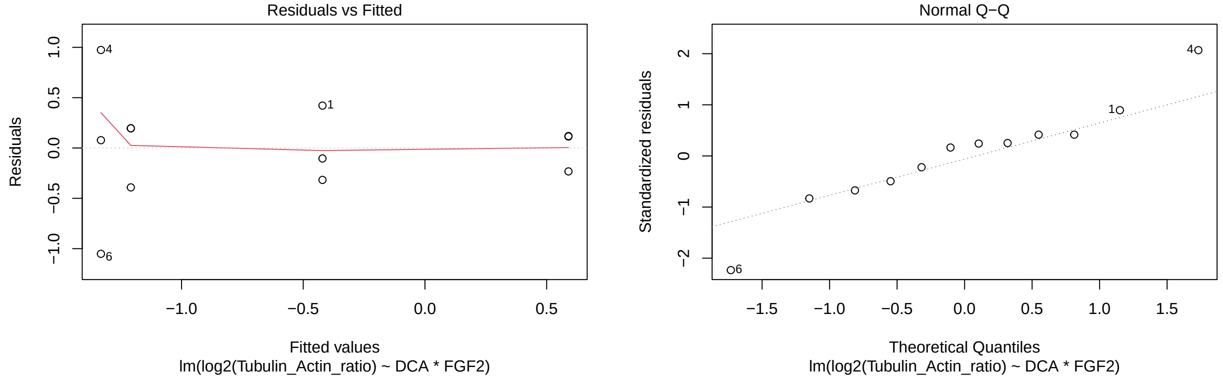

```r
plot(aS6G_PDH,which=1:2)
```

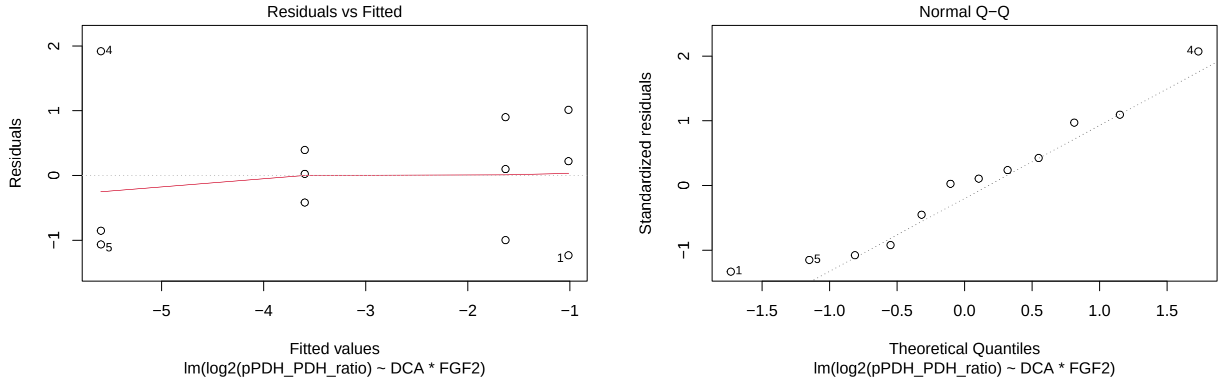

```r
plot(aS6G_ERK,which=1:2)
```

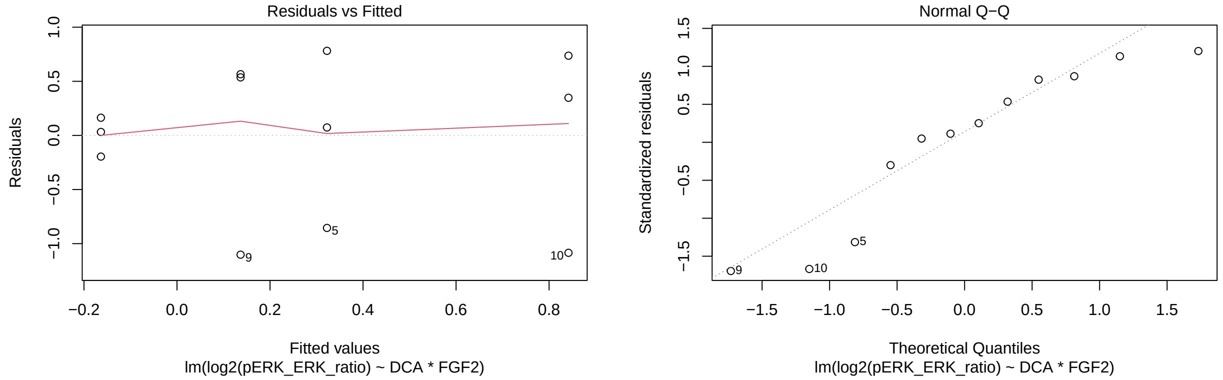

Above are the results from an analysis of this experiment where the responses are transformed using $\log_2$. Though some residual plots for the $\log_2$ are not ideal (the variance of the groups sometimes seems to vary), they are preferred over the analysis without the log transformation (below), where three of the five models exhibit residuals with a cone shape, which is a tell-tale sign that a transformation (or some other procedure) is necessary.

```r
a4G_PDH_nl <- lm(pPDH_PDH_ratio ~ DCA*FGF2, data=d4G)
a4G_ERK_nl <- lm(pERK_ERK_ratio ~ DCA*FGF2, data=d4G)
a4G_Tub_nl <- lm(Tubulin_Actin_ratio ~ DCA*FGF2, data=d4G)

aS6G_PDH_nl <- lm(pPDH_PDH_ratio ~ DCA*FGF2, data=dS6G)
aS6G_ERK_nl <- lm(pERK_ERK_ratio ~ DCA*FGF2, data=dS6G)

plot(a4G_PDH_nl,which=1:2)
```

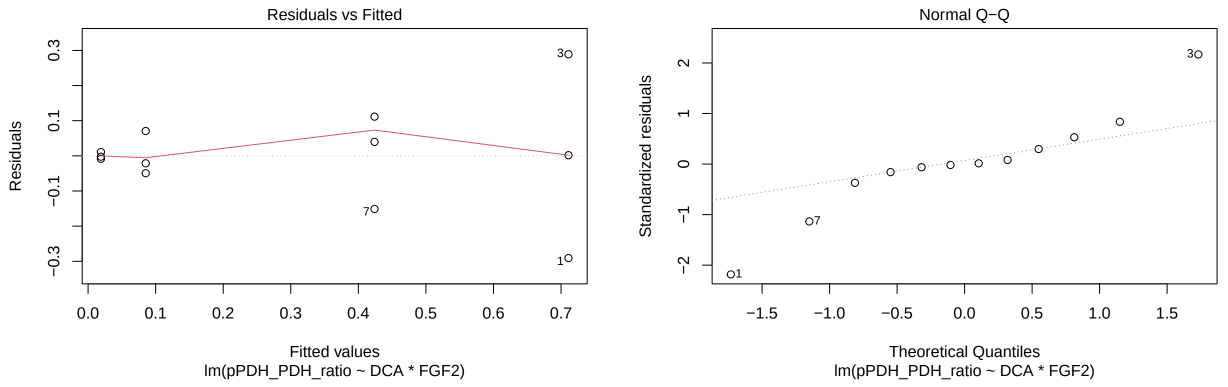

```r
plot(a4G_ERK_nl,which=1:2)
```

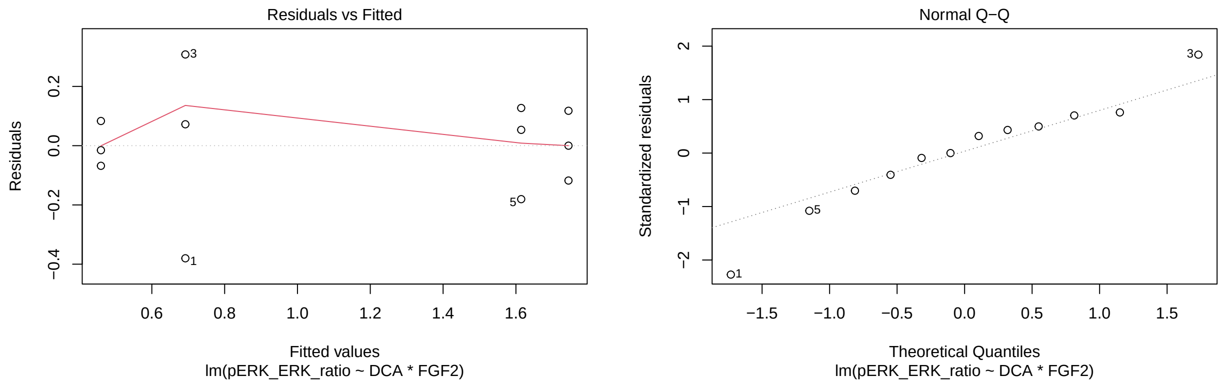

```r
plot(a4G_Tub_nl,which=1:2)
```

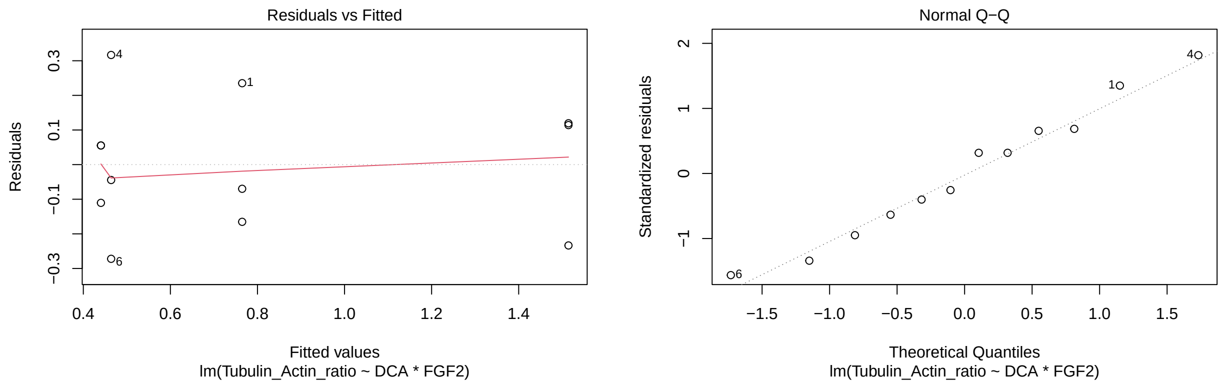

```r
plot(aS6G_PDH_nl,which=1:2)
```

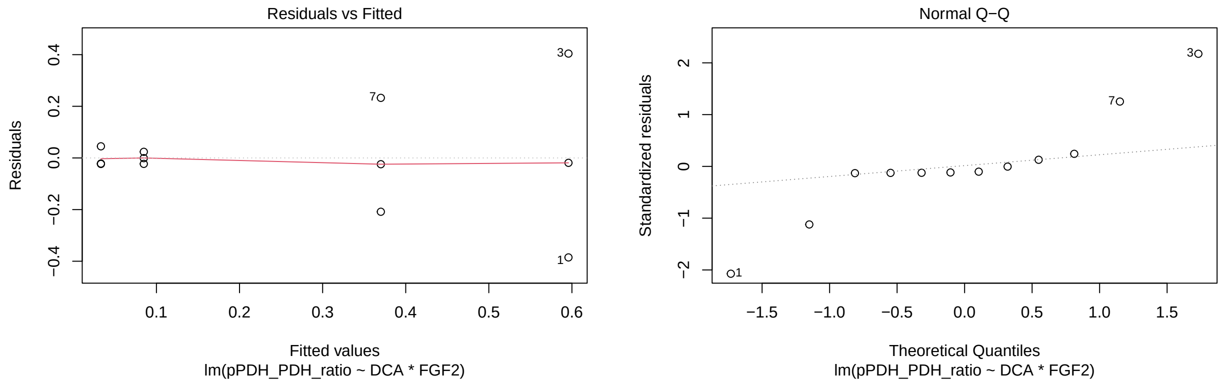

```r
plot(aS6G_ERK_nl,which=1:2)
```

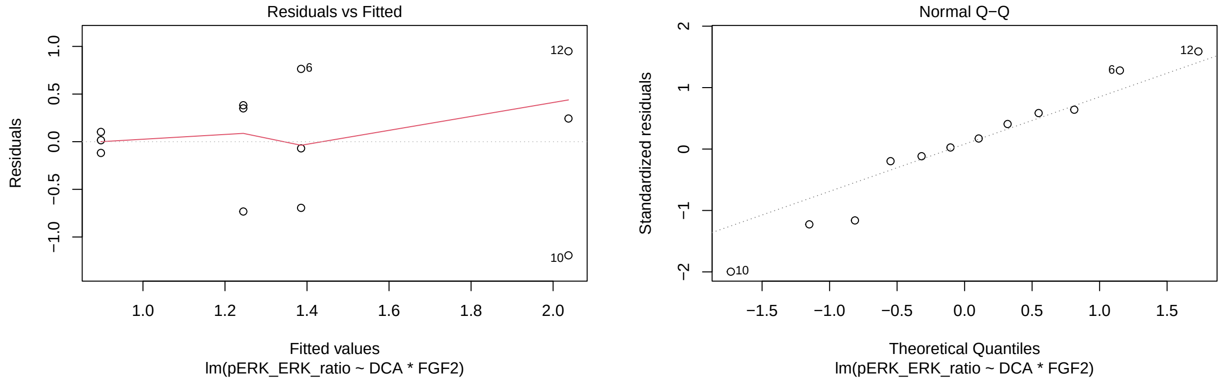

##### The Four Comparisons (based on $\log_2$ analysis)

The models we've fit above are of the form:

$$E(\log_2(Y)) = \beta_0 + \beta_1 D + \beta_2 F + \beta_3 D * F,$$

where $Y$ represents the response of interest (e.g. pERK/ERK), $D$ is an indicator variable that is 1 if the unit is DCA, 0 otherwise; and $F$ is an indicator for FGF2. Thus, when neither chemical is included (PBS treatment), the mean $\log_2$ response is $\beta_0$, while if only DCA is included, the mean is $\beta_0 + \beta_1$, if only FGF2 is included, the mean is $\beta_0 + \beta_2$, and if both are included, the mean is $\beta_0 + \beta_1 + \beta_2 + \beta_3$. Thus, to represent the four comparisons of interest, we have:

1. PBS vs. FGF2: $\beta_0 + \beta_2 - \beta_0 = \beta_2$
2. PBS vs. DCA: $\beta_1$
3. FGF2 vs. DCA+FGF2: $(\beta_0 + \beta_1 + \beta_2 + \beta_3) - (\beta_0 + \beta_2) = \beta_1 + \beta_3$.
4. DCA vs. DCA+FGF2: $(\beta_0 + \beta_1 + \beta_2 + \beta_3) - (\beta_0 + \beta_1) = \beta_2 + \beta_3$.

Below, we perform these statistical comparisons.

```
K <- matrix(c(0,0,1,0,
              0,1,0,0,
              0,1,0,1,
              0,0,1,1),nrow=4, byrow=TRUE)

t <- glht(a4G_PDH, linfct=K)
summary(t, test=adjusted("fdr"))
```

```
##
##   Simultaneous Tests for General Linear Hypotheses
##
## Fit: lm(formula = log2(pPDH_PDH_ratio) ~ DCA * FGF2, data = d4G)
##
## Linear Hypotheses:
##        Estimate Std. Error t value Pr(>|t|)
## 1 == 0  -0.7150     0.6273  -1.140  0.28734
## 2 == 0  -3.2353     0.6273  -5.158  0.00173 **
## 3 == 0  -4.5511     0.6273  -7.255  0.00035 ***
## 4 == 0  -2.0307     0.6273  -3.237  0.01590 *
## ---
## Signif. codes:  0 '***' 0.001 '**' 0.01 '*' 0.05 '.' 0.1 ' ' 1
## (Adjusted p values reported -- fdr method)
```

```
t <- glht(a4G_ERK, linfct=K)
summary(t, test=adjusted("fdr"))
```

```
##
##   Simultaneous Tests for General Linear Hypotheses
##
## Fit: lm(formula = log2(pERK_ERK_ratio) ~ DCA * FGF2, data = d4G)
##
## Linear Hypotheses:
##        Estimate Std. Error t value Pr(>|t|)
## 1 == 0  -0.4435     0.3790  -1.170  0.36743
## 2 == 0   1.3756     0.3790   3.630  0.01338 *
## 3 == 0   1.9334     0.3790   5.102  0.00371 **
## 4 == 0   0.1143     0.3790   0.302  0.77063
## ---
## Signif. codes:  0 '***' 0.001 '**' 0.01 '*' 0.05 '.' 0.1 ' ' 1
## (Adjusted p values reported -- fdr method)
```

```r
t <- glht(a4G_Tub, linfct=K)
summary(t, test=adjusted("fdr"))
```

```
##
##   Simultaneous Tests for General Linear Hypotheses
##
## Fit: lm(formula = log2(Tubulin_Actin_ratio) ~ DCA * FGF2, data = d4G)
##
## Linear Hypotheses:
##        Estimate Std. Error t value Pr(>|t|)
## 1 == 0   1.0106     0.4707   2.147   0.1190
## 2 == 0  -0.9102     0.4707  -1.934   0.1190
## 3 == 0  -1.7979     0.4707  -3.819   0.0204 *
## 4 == 0   0.1229     0.4707   0.261   0.8006
## ---
## Signif. codes:  0 '***' 0.001 '**' 0.01 '*' 0.05 '.' 0.1 ' ' 1
## (Adjusted p values reported -- fdr method)
```

```r
t <- glht(aS6G_PDH, linfct=K)
summary(t, test=adjusted("fdr"))
```

```
##
##   Simultaneous Tests for General Linear Hypotheses
##
## Fit: lm(formula = log2(pPDH_PDH_ratio) ~ DCA * FGF2, data = dS6G)
##
## Linear Hypotheses:
##        Estimate Std. Error t value Pr(>|t|)
## 1 == 0  -0.6169     0.9260  -0.666   0.5240
## 2 == 0  -4.5809     0.9260  -4.947   0.0045 **
## 3 == 0  -1.9667     0.9260  -2.124   0.0886 .
## 4 == 0   1.9972     0.9260   2.157   0.0886 .
## ---
## Signif. codes:  0 '***' 0.001 '**' 0.01 '*' 0.05 '.' 0.1 ' ' 1
## (Adjusted p values reported -- fdr method)
```

```r
t <- glht(aS6G_ERK, linfct=K)
summary(t, test=adjusted("fdr"))
```

```
##
##   Simultaneous Tests for General Linear Hypotheses
##
## Fit: lm(formula = log2(pERK_ERK_ratio) ~ DCA * FGF2, data = dS6G)
##
## Linear Hypotheses:
##        Estimate Std. Error t value Pr(>|t|)
## 1 == 0   0.3003     0.6504   0.462    0.657
## 2 == 0   0.4860     0.6504   0.747    0.635
## 3 == 0   0.7050     0.6504   1.084    0.635
## 4 == 0   0.5192     0.6504   0.798    0.635
## (Adjusted p values reported -- fdr method)
```

### Experiment 7 (Figure 5B)

Responses are normalized counts of mRNAs. We have 4 Treatments (PBS, FGF2, DCA, and FGF2+DCA), all at Time=48 and three replications per treatment, so just 12 experimental units here.

Comparisons for Figure 5B: PBS vs FGF2; PBS vs DCA; FGF2 vs FGF2+DCA, and DCA vs FGF2+DCA for each gene. See Experiment 6 for details regarding how these statistical comparisons were performed.

#### Data and Visualization

```
d5B <- read.xlsx(xlsxFile = "MetabolismPaperRawData_Byran_working.xlsx",
                 colNames = TRUE, sheet="Fig 5_B")
glimpse(d5B)
```

```
## Rows: 12
## Columns: 10
## $ Chemical   <chr> "PBS", "PBS", "PBS", "FGF2", "FGF2", "FGF2", "DCA", "DCA", ~
## $ Sample     <dbl> 1, 2, 3, 1, 2, 3, 1, 2, 3, 1, 2, 3
## $ DCA        <dbl> 0, 0, 0, 0, 0, 0, 1, 1, 1, 1, 1, 1
## $ FGF2       <dbl> 0, 0, 0, 1, 1, 1, 0, 0, 0, 1, 1, 1
## $ `Time.(h)` <dbl> 48, 48, 48, 48, 48, 48, 48, 48, 48, 48, 48, 48
## $ FGFR2      <dbl> 13171.33, 12881.48, 13568.77, 19956.18, 21749.10, 20666.79,~
## $ ASCL1      <dbl> 708.49470, 805.60599, 1287.87622, 2058.61433, 2302.85761, 2~
## $ MAP1LC3B   <dbl> 6994.995, 6992.594, 7137.578, 6254.725, 5957.808, 5874.725,~
## $ LDHA       <dbl> 89555.35, 114012.55, 92654.40, 209667.51, 234715.66, 217070~
## $ TYRP1      <dbl> 45103.116, 47988.988, 34990.716, 19532.185, 11687.241, 1120~
```

```
d5B_long <- d5B %>%
  rename(Time=`Time.(h)`) %>%
  pivot_longer(cols=6:10,names_to="Gene",values_to="mRNA_norm_count") %>%
  mutate(Time=as.factor(Time))
glimpse(d5B_long)
```

```
## Rows: 60
## Columns: 7
## $ Chemical        <chr> "PBS", "PBS", "PBS", "PBS", "PBS", "PBS", "PBS", "PBS"~
## $ Sample          <dbl> 1, 1, 1, 1, 1, 2, 2, 2, 2, 2, 3, 3, 3, 3, 3, 1, 1, 1, ~
## $ DCA             <dbl> 0, 0, 0, 0, 0, 0, 0, 0, 0, 0, 0, 0, 0, 0, 0, 0, 0, 0, ~
## $ FGF2            <dbl> 0, 0, 0, 0, 0, 0, 0, 0, 0, 0, 0, 0, 0, 0, 0, 1, 1, 1, ~
## $ Time            <fct> 48, 48, 48, 48, 48, 48, 48, 48, 48, 48, 48, 48, 48, 48~
## $ Gene            <chr> "FGFR2", "ASCL1", "MAP1LC3B", "LDHA", "TYRP1", "FGFR2"~
## $ mRNA_norm_count <dbl> 13171.3308, 708.4947, 6994.9955, 89555.3471, 45103.116~
```

```
ggplot(d5B_long, aes(x=Chemical,y=log2(mRNA_norm_count))) +
  geom_jitter(width=.05,height=0) +
  facet_wrap(~Gene, nrow=3) +
  theme(axis.text.x = element_text(angle = 90, vjust = 0.5, hjust=1)) +
  labs(title="For Figure 5B: Treatment Comparisons")
```

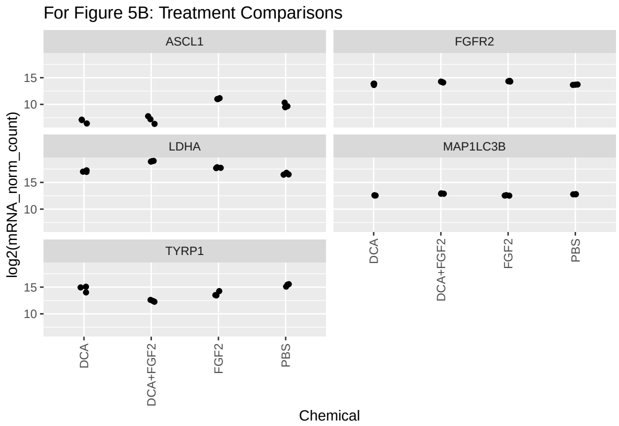

### Data Analysis (Experiment 7)

Analysis both with one-way ANOVA (4 levels) with the four required comparisons, but also with two-way ANOVA (DCA: present/absent; FGF2: present/absent) with interaction plots.

```
a5B_FGFR2 <- lm(log2(FGFR2)~DCA*FGF2, data=d5B)
a5B_ASCL1 <- lm(log2(ASCL1)~DCA*FGF2, data=d5B)
a5B_MAP1LC3B <- lm(log2(MAP1LC3B)~DCA*FGF2, data=d5B)
a5B_LDHA <- lm(log2(LDHA)~DCA*FGF2, data=d5B)
a5B_TYRP1 <- lm(log2(TYRP1)~DCA*FGF2, data=d5B)

plot(a5B_FGFR2, which=1:2)
```

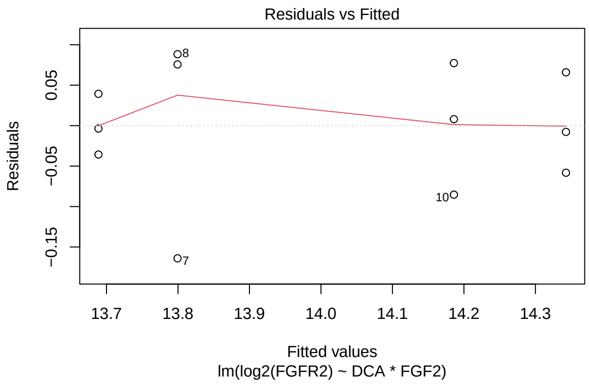

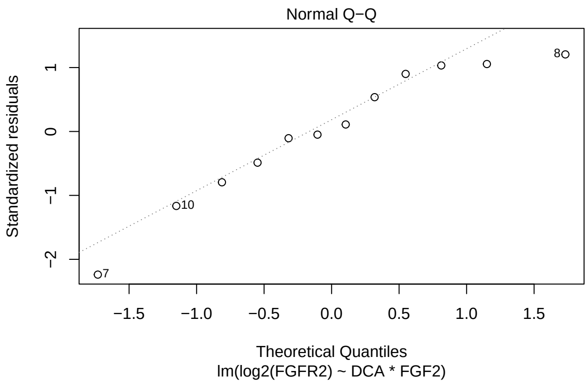

```
plot(a5B_ASCL1, which=1:2)
```

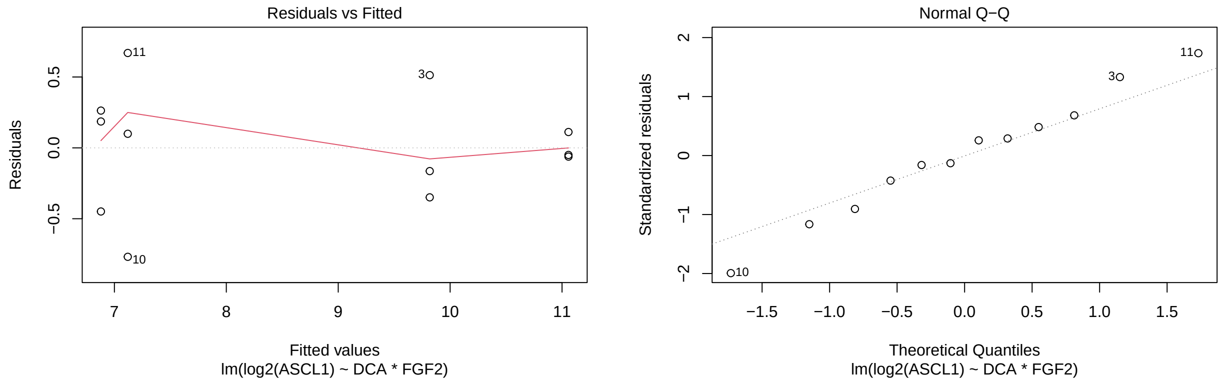

```r
plot(a5B_MAP1LC3B, which=1:2)
```

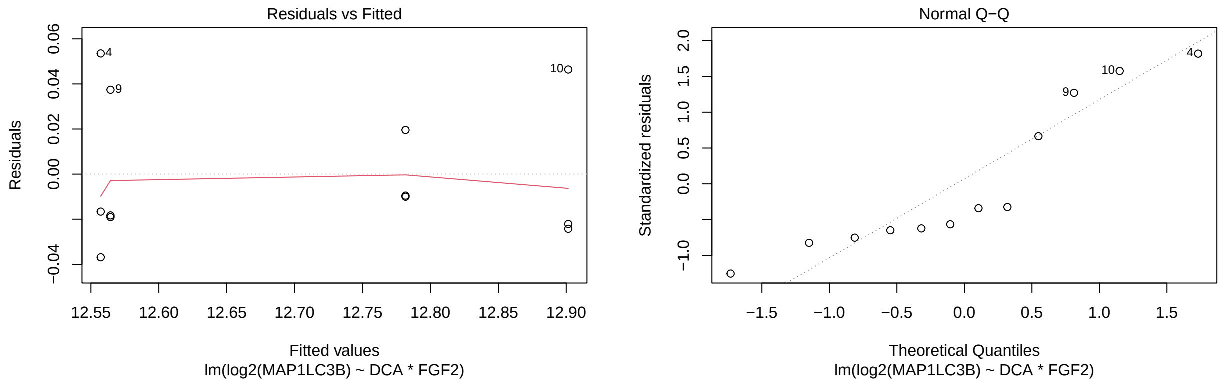

```r
plot(a5B_LDHA, which=1:2)
```

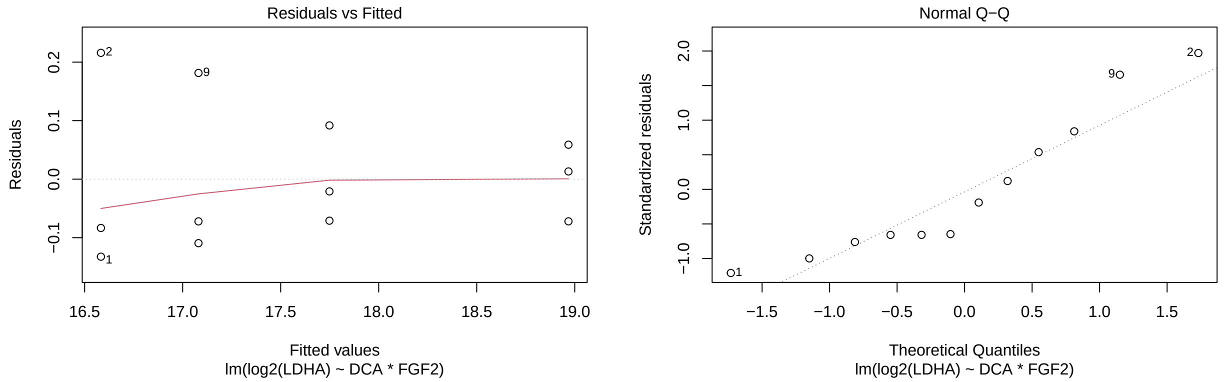

```r
plot(a5B_TYRP1, which=1:2)
```

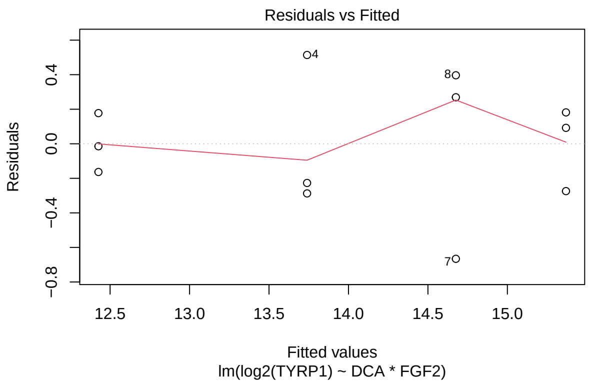

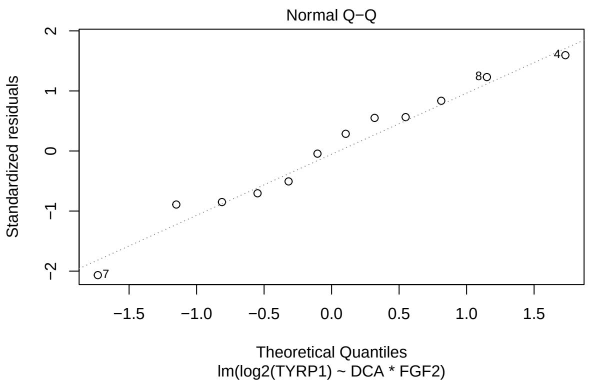

##### The Four Main Comparisons

```
K <- matrix(c(0,0,1,0,
              0,1,0,0,
              0,1,0,1,
              0,0,1,1),nrow=4, byrow=TRUE)

t <- glht(a5B_FGFR2, linfct=K)
summary(t, test=adjusted("fdr"))
```

```
##
##   Simultaneous Tests for General Linear Hypotheses
##
## Fit: lm(formula = log2(FGFR2) ~ DCA * FGF2, data = d5B)
##
## Linear Hypotheses:
##        Estimate Std. Error t value Pr(>|t|)
## 1 == 0  0.65404    0.07321   8.933 7.83e-05 ***
## 2 == 0  0.11069    0.07321   1.512  0.16903
## 3 == 0 -0.15676    0.07321  -2.141  0.08623 .
## 4 == 0  0.38659    0.07321   5.280  0.00149 **
## ---
## Signif. codes:  0 '***' 0.001 '**' 0.01 '*' 0.05 '.' 0.1 ' ' 1
## (Adjusted p values reported -- fdr method)
```

```
t <- glht(a5B_ASCL1, linfct=K)
summary(t, test=adjusted("fdr"))
```

```
##
##   Simultaneous Tests for General Linear Hypotheses
##
## Fit: lm(formula = log2(ASCL1) ~ DCA * FGF2, data = d5B)
##
## Linear Hypotheses:
##        Estimate Std. Error t value Pr(>|t|)
## 1 == 0   1.2397     0.3857   3.214 0.016459 *
```

```
## 2 == 0  -2.9383     0.3857  -7.619 0.000124 ***
## 3 == 0  -3.9379     0.3857 -10.211 2.91e-05 ***
## 4 == 0   0.2400     0.3857   0.622 0.551038
## ---
## Signif. codes:  0 '***' 0.001 '**' 0.01 '*' 0.05 '.' 0.1 ' ' 1
## (Adjusted p values reported -- fdr method)
```

```
t <- glht(a5B_MAP1LC3B, linfct=K)
summary(t, test=adjusted("fdr"))
```

```
##
##   Simultaneous Tests for General Linear Hypotheses
##
## Fit: lm(formula = log2(MAP1LC3B) ~ DCA * FGF2, data = d5B)
##
## Linear Hypotheses:
##        Estimate Std. Error t value Pr(>|t|)
## 1 == 0 -0.22445    0.02945  -7.621 7.80e-05 ***
## 2 == 0 -0.21724    0.02945  -7.376 7.80e-05 ***
## 3 == 0  0.34431    0.02945  11.691 6.14e-06 ***
## 4 == 0  0.33710    0.02945  11.446 6.14e-06 ***
## ---
## Signif. codes:  0 '***' 0.001 '**' 0.01 '*' 0.05 '.' 0.1 ' ' 1
## (Adjusted p values reported -- fdr method)
```

```
t <- glht(a5B_LDHA, linfct=K)
summary(t, test=adjusted("fdr"))
```

```
##
##   Simultaneous Tests for General Linear Hypotheses
##
## Fit: lm(formula = log2(LDHA) ~ DCA * FGF2, data = d5B)
##
## Linear Hypotheses:
##        Estimate Std. Error t value Pr(>|t|)
## 1 == 0   1.1657     0.1095  10.643 7.09e-06 ***
## 2 == 0   0.4977     0.1095   4.544  0.00189 **
## 3 == 0   1.2193     0.1095  11.132 7.09e-06 ***
## 4 == 0   1.8873     0.1095  17.231 5.24e-07 ***
## ---
## Signif. codes:  0 '***' 0.001 '**' 0.01 '*' 0.05 '.' 0.1 ' ' 1
## (Adjusted p values reported -- fdr method)
```

```
t <- glht(a5B_TYRP1, linfct=K)
summary(t, test=adjusted("fdr"))
```

```
##
##   Simultaneous Tests for General Linear Hypotheses
##
## Fit: lm(formula = log2(TYRP1) ~ DCA * FGF2, data = d5B)
##
## Linear Hypotheses:
```

```
##        Estimate Std. Error t value Pr(>|t|)
## 1 == 0  -1.6292     0.3223  -5.055  0.00197 **
## 2 == 0  -0.6929     0.3223  -2.150  0.06380 .
## 3 == 0  -1.3130     0.3223  -4.074  0.00475 **
## 4 == 0  -2.2494     0.3223  -6.979  0.00046 ***
## ---
## Signif. codes:  0 '***' 0.001 '**' 0.01 '*' 0.05 '.' 0.1 ' ' 1
## (Adjusted p values reported -- fdr method)
```

```r
# https://rcompanion.org/rcompanion/h_01.html
```

##### Interaction / Main Effects Plots

Here we identify strong DCA and/or FGF2 main effects, along with prominent interactions between the two factors. We do not use these results explicitly in the paper, but they provide some insight.

```r
summary(a5B_FGFR2)
```

```
##
## Call:
## lm(formula = log2(FGFR2) ~ DCA * FGF2, data = d5B)
##
## Residuals:
##       Min        1Q    Median        3Q       Max
## -0.163940 -0.041323  0.002214  0.068350  0.088304
##
## Coefficients:
##             Estimate Std. Error t value Pr(>|t|)
## (Intercept) 13.68871    0.05177 264.411  < 2e-16 ***
## DCA          0.11069    0.07321   1.512   0.1690
## FGF2         0.65404    0.07321   8.933 1.96e-05 ***
## DCA:FGF2    -0.26744    0.10354  -2.583   0.0325 *
## ---
## Signif. codes:  0 '***' 0.001 '**' 0.01 '*' 0.05 '.' 0.1 ' ' 1
##
## Residual standard error: 0.08967 on 8 degrees of freedom
## Multiple R-squared:  0.931,  Adjusted R-squared:  0.9051
## F-statistic: 35.96 on 3 and 8 DF,  p-value: 5.433e-05
```

```r
ggplot(d5B) +
  aes(x = FGF2, color = as.factor(DCA), group = as.factor(DCA), y = log2(FGFR2)) +
  stat_summary(fun = mean, geom = "point") +
  stat_summary(fun = mean, geom = "line") +
  geom_point()
```

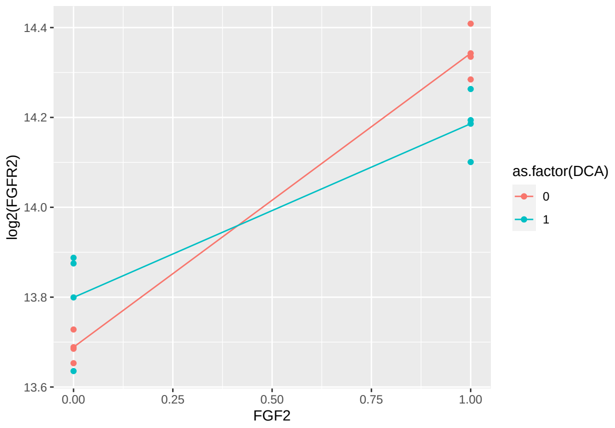

For $\log_2(FGFR2)$, the FGF2 effect is larger when DCA is present.

```
summary(a5B_ASCL1)
```

```
##
## Call:
## lm(formula = log2(ASCL1) ~ DCA * FGF2, data = d5B)
##
## Residuals:
##     Min      1Q  Median      3Q     Max
## -0.7687 -0.2102  0.0247  0.2053  0.6694
##
## Coefficients:
##             Estimate Std. Error t value Pr(>|t|)
## (Intercept)   9.8178     0.2727  36.001 3.88e-10 ***
## DCA          -2.9383     0.3857  -7.619 6.20e-05 ***
## FGF2          1.2397     0.3857   3.214   0.0123 *
## DCA:FGF2     -0.9996     0.5454  -1.833   0.1042
## ---
## Signif. codes:  0 '***' 0.001 '**' 0.01 '*' 0.05 '.' 0.1 ' ' 1
##
## Residual standard error: 0.4723 on 8 degrees of freedom
## Multiple R-squared:  0.955,  Adjusted R-squared:  0.9381
## F-statistic: 56.55 on 3 and 8 DF,  p-value: 9.934e-06
```

```
ggplot(d5B) +
  aes(x = FGF2, y = log2(ASCL1)) +
  stat_summary(fun = mean, geom = "point") +
  stat_summary(fun = mean, geom = "line") +
  geom_point()
```

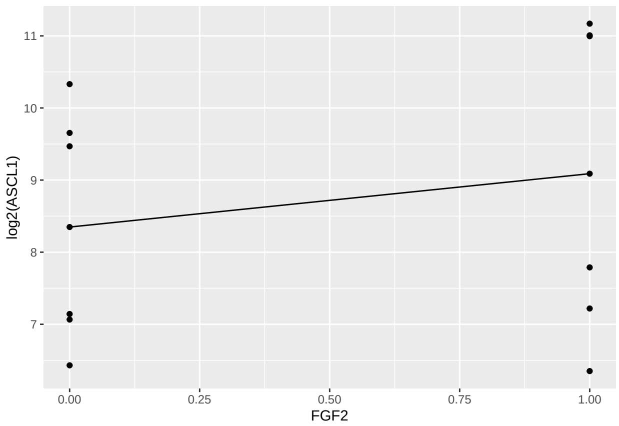

```r
ggplot(d5B) +
  aes(x = DCA, y = log2(ASCL1)) +
  stat_summary(fun = mean, geom = "point") +
  stat_summary(fun = mean, geom = "line") +
  geom_point()
```

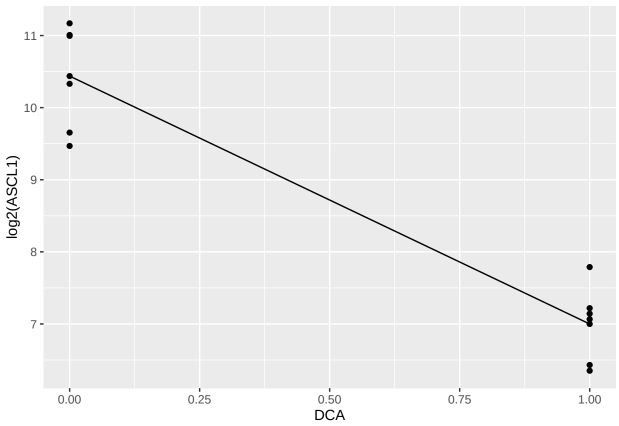

```r
summary(a5B_MAP1LC3B)
```

```
##
## Call:
## lm(formula = log2(MAP1LC3B) ~ DCA * FGF2, data = d5B)
##
## Residuals:
##      Min       1Q   Median       3Q      Max
## -0.03689 -0.01985 -0.01333  0.02403  0.05353
##
## Coefficients:
##             Estimate Std. Error t value Pr(>|t|)
## (Intercept) 12.78165    0.02083 613.751  < 2e-16 ***
## DCA         -0.21724    0.02945  -7.376 7.80e-05 ***
## FGF2        -0.22445    0.02945  -7.621 6.18e-05 ***
## DCA:FGF2     0.56155    0.04165  13.482 8.79e-07 ***
## ---
## Signif. codes:  0 '***' 0.001 '**' 0.01 '*' 0.05 '.' 0.1 ' ' 1
```

```
##
## Residual standard error: 0.03607 on 8 degrees of freedom
## Multiple R-squared:  0.9612, Adjusted R-squared:  0.9467
## F-statistic: 66.13 on 3 and 8 DF,  p-value: 5.468e-06
```

```
ggplot(d5B) +
  aes(x = FGF2, color = as.factor(DCA), group = as.factor(DCA), y = log2(MAP1LC3B)) +
  stat_summary(fun = mean, geom = "point") +
  stat_summary(fun = mean, geom = "line") +
  geom_point()
```

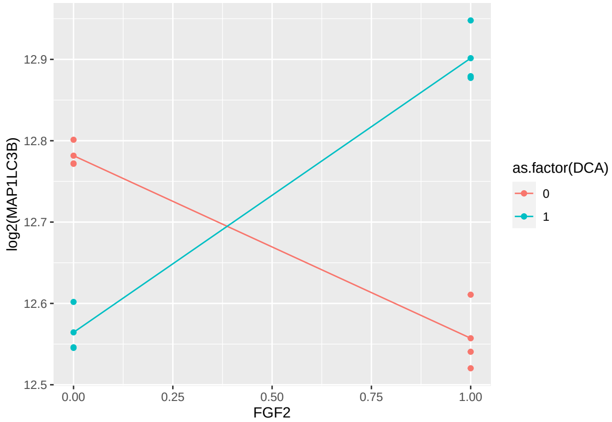

$\log_2(MAP1LC3B)$, the FGF2 effect is clearly positive when DCA is present and clearly negative when DCA is absent.

```
summary(a5B_LDHA)
```

```
##
## Call:
## lm(formula = log2(LDHA) ~ DCA * FGF2, data = d5B)
##
## Residuals:
##      Min       1Q   Median       3Q      Max
## -0.13247 -0.07495 -0.04593  0.06714  0.21587
##
## Coefficients:
##             Estimate Std. Error t value Pr(>|t|)
## (Intercept) 16.58297    0.07745 214.110 2.53e-16 ***
## DCA          0.49767    0.10953   4.544  0.00189 **
## FGF2         1.16573    0.10953  10.643 5.32e-06 ***
## DCA:FGF2     0.72160    0.15490   4.658  0.00163 **
## ---
## Signif. codes:  0 '***' 0.001 '**' 0.01 '*' 0.05 '.' 0.1 ' ' 1
##
## Residual standard error: 0.1341 on 8 degrees of freedom
## Multiple R-squared:  0.9852, Adjusted R-squared:  0.9797
## F-statistic: 177.7 on 3 and 8 DF,  p-value: 1.169e-07
```

```r
ggplot(d5B) +
  aes(x = FGF2, color = as.factor(DCA), group = as.factor(DCA), y = log2(LDHA)) +
  stat_summary(fun = mean, geom = "point") +
  stat_summary(fun = mean, geom = "line") +
  geom_point()
```

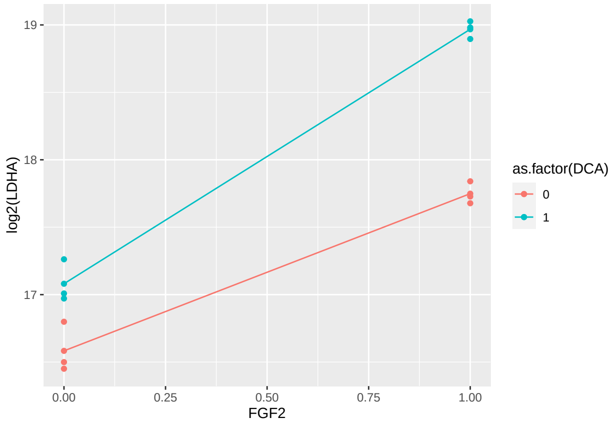

For $\log_2(LDHA)$, the FGF2 effect is slightly larger when DCA is present.

```r
summary(a5B_TYRP1)
```

```
##
## Call:
## lm(formula = log2(TYRP1) ~ DCA * FGF2, data = d5B)
##
## Residuals:
##      Min       1Q   Median       3Q      Max
## -0.66595 -0.23861  0.03898  0.20363  0.51410
##
## Coefficients:
##             Estimate Std. Error t value Pr(>|t|)
## (Intercept)  15.3687     0.2279  67.434   2.6e-12 ***
## DCA          -0.6929     0.3223  -2.150 0.063805 .
## FGF2         -1.6292     0.3223  -5.055 0.000983 ***
## DCA:FGF2     -0.6202     0.4558  -1.361 0.210746
## ---
## Signif. codes:  0 '***' 0.001 '**' 0.01 '*' 0.05 '.' 0.1 ' ' 1
##
## Residual standard error: 0.3947 on 8 degrees of freedom
## Multiple R-squared:  0.9213, Adjusted R-squared:  0.8918
## F-statistic: 31.21 on 3 and 8 DF,  p-value: 9.149e-05
```

```r
ggplot(d5B) +
  aes(x = FGF2, y = log2(TYRP1)) +
  stat_summary(fun = mean, geom = "point") +
  stat_summary(fun = mean, geom = "line") +
  geom_point()
```

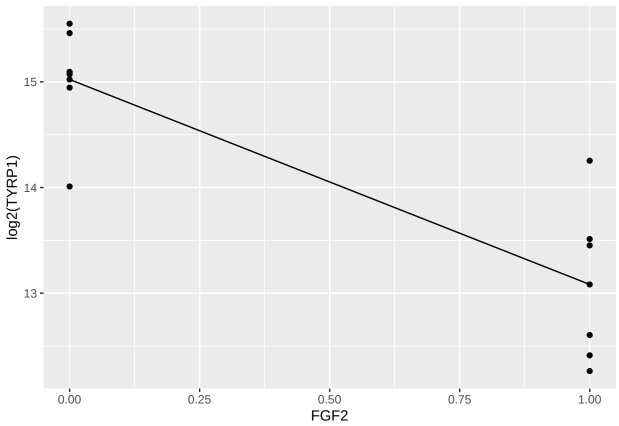

```r
ggplot(d5B) +
  aes(x = DCA, y = log2(TYRP1)) +
  stat_summary(fun = mean, geom = "point") +
  stat_summary(fun = mean, geom = "line") +
  geom_point()
```

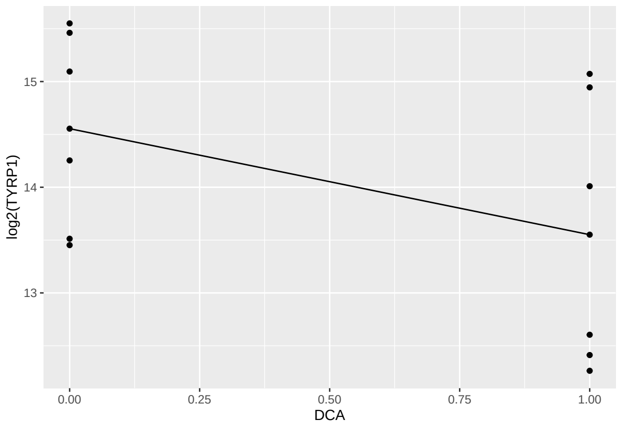

### Experiment 8 (Figures 7C, 7D, and 7E)

Response is relative expression of several genes. There are three Chemical factors, all at Time=48: FGF2 (present/absent); DCA (present/absent); NAC (present/absent), so 8 treatments with 4 replications/treatment.

Comparisons for Figure 7C, 7D, 7E: PBS vs DCA; PBS vs NAC; PBS vs DCA+NAC; DCA vs DCA+NAC; PBS vs FGF2; FGF2 vs FGF2+DCA; FGF2 vs FGF2+NAC; FGF2 vs FGF2+DCA+NAC; FGF2+DCA vs FGF2+DCA+NAC for each gene.

#### Data and Visualization

We will first obtain the datasets for each relevant figure (Figures 7C, 7D, 7E), visualize them, then analyze them according to the specified set of comparisons.

###### Plots to Compare the Treatments

```
d7C <- read.xlsx(xlsxFile = "MetabolismPaperRawData_Byran_working.xlsx", colNames = TRUE, sheet="Fig 7_(
glimpse(d7C)
```

```
## Rows: 32
## Columns: 12
## $ Chemical    <chr> "PBS", "PBS", "PBS", "PBS", "DCA", "DCA", "DCA", "DCA", "NA~
## $ DCA         <dbl> 0, 0, 0, 0, 1, 1, 1, 1, 0, 0, 0, 0, 1, 1, 1, 1, 0, 0, 0, 0,~
## $ NAC         <dbl> 0, 0, 0, 0, 0, 0, 0, 0, 1, 1, 1, 1, 1, 1, 1, 1, 0, 0, 0, 0,~
## $ FGF2        <dbl> 0, 0, 0, 0, 0, 0, 0, 0, 0, 0, 0, 0, 0, 0, 0, 0, 1, 1, 1, 1,~
## $ Sample      <dbl> 1, 2, 3, 4, 1, 2, 3, 4, 1, 2, 3, 4, 1, 2, 3, 4, 1, 2, 3, 4,~
## $ `Time.(h)` <dbl> 48, 48, 48, 48, 48, 48, 48, 48, 48, 48, 48, 48, 48, 48, 48,~
## $ SOX2        <dbl> 1.32, 0.89, 0.96, 0.89, 1.42, 1.45, 1.03, 1.10, 1.30, 1.08,~
## $ SIX6        <dbl> 1.5510985, 1.5032274, 0.5927657, 0.7235239, 1.3989732, 1.36~
## $ PAX6        <dbl> 0.9964433, 1.1834182, 0.8755977, 0.9685109, 1.6571503, 1.24~
## $ RPE65       <dbl> 0.72, 0.60, 2.02, 1.16, 0.37, 1.04, 0.38, 0.45, 0.39, 0.32,~
## $ TYR         <dbl> 1.21134227, 1.00178043, 0.90325050, 0.91233088, 0.37832580,~
## $ OTX2        <dbl> 1.1549546, 1.0504189, 0.8678014, 0.9498438, 0.8081995, 0.98~
```

```
d7D <- read.xlsx(xlsxFile = "MetabolismPaperRawData_Byran_working.xlsx", colNames = TRUE, sheet="Fig 7_I
glimpse(d7D)
```

```
## Rows: 32
## Columns: 10
## $ Chemical    <chr> "PBS", "PBS", "PBS", "PBS", "DCA", "DCA", "DCA", "DCA", "NA~
## $ DCA         <dbl> 0, 0, 0, 0, 1, 1, 1, 1, 0, 0, 0, 0, 1, 1, 1, 1, 0, 0, 0, 0,~
## $ NAC         <dbl> 0, 0, 0, 0, 0, 0, 0, 0, 1, 1, 1, 1, 1, 1, 1, 1, 0, 0, 0, 0,~
## $ FGF2        <dbl> 0, 0, 0, 0, 0, 0, 0, 0, 0, 0, 0, 0, 0, 0, 0, 0, 1, 1, 1, 1,~
## $ Sample      <dbl> 1, 2, 3, 4, 1, 2, 3, 4, 1, 2, 3, 4, 1, 2, 3, 4, 1, 2, 3, 4,~
## $ `Time.(h)` <dbl> 48, 48, 48, 48, 48, 48, 48, 48, 48, 48, 48, 48, 48, 48, 48,~
## $ TGFB2       <dbl> 1.5028399, 1.2524987, 0.7551985, 0.7034754, 1.9804227, 2.22~
## $ VIM         <dbl> 0.7755005, 1.1842007, 1.0866119, 1.0021162, 0.5370472, 0.96~
## $ SNAI1       <dbl> 0.62117728, 1.32374560, 1.08294839, 1.12298032, 0.47897417,~
## $ aSMA        <dbl> 0.77262692, 1.01501266, 1.42089100, 0.89742454, 0.61866813,~
```

```
d7E <- read.xlsx(xlsxFile = "MetabolismPaperRawData_Byran_working.xlsx", colNames = TRUE, sheet="Fig 7_I
glimpse(d7E)
```

```
## Rows: 32
## Columns: 8
## $ Chemical    <chr> "PBS", "PBS", "PBS", "PBS", "DCA", "DCA", "DCA", "DCA", "NA~
## $ DCA         <dbl> 0, 0, 0, 0, 1, 1, 1, 1, 0, 0, 0, 0, 1, 1, 1, 1, 0, 0, 0, 0,~
## $ NAC         <dbl> 0, 0, 0, 0, 0, 0, 0, 0, 1, 1, 1, 1, 1, 1, 1, 1, 0, 0, 0, 0,~
## $ FGF2        <dbl> 0, 0, 0, 0, 0, 0, 0, 0, 0, 0, 0, 0, 0, 0, 0, 0, 1, 1, 1, 1,~
## $ Sample      <dbl> 1, 2, 3, 4, 1, 2, 3, 4, 1, 2, 3, 4, 1, 2, 3, 4, 1, 2, 3, 4,~
## $ `Time.(h)` <dbl> 48, 48, 48, 48, 48, 48, 48, 48, 48, 48, 48, 48, 48, 48, 48,~
## $ E2F1        <dbl> 0.8876454, 1.0116427, 0.9026918, 1.2336554, 0.7502307, 0.86~
## $ PCNA        <dbl> 1.1070841, 1.1398686, 0.8395889, 0.9438390, 0.7853186, 0.91~
```

```r
d7C_long <- d7C %>%
  rename(Time=`Time.(h)`) %>%
  pivot_longer(cols=7:12,names_to="Gene",values_to="RGE") %>%
  mutate(Time=as.factor(Time))
glimpse(d7C_long)
```

```
## Rows: 192
## Columns: 8
## $ Chemical <chr> "PBS", "PBS", "PBS", "PBS", "PBS", "PBS", "PBS", "PBS", "PBS"~
## $ DCA      <dbl> 0, 0, 0, 0, 0, 0, 0, 0, 0, 0, 0, 0, 0, 0, 0, 0, 0, 0, 0, 0, 0~
## $ NAC      <dbl> 0, 0, 0, 0, 0, 0, 0, 0, 0, 0, 0, 0, 0, 0, 0, 0, 0, 0, 0, 0, 0~
## $ FGF2     <dbl> 0, 0, 0, 0, 0, 0, 0, 0, 0, 0, 0, 0, 0, 0, 0, 0, 0, 0, 0, 0, 0~
## $ Sample   <dbl> 1, 1, 1, 1, 1, 1, 2, 2, 2, 2, 2, 2, 3, 3, 3, 3, 3, 3, 4, 4, 4~
## $ Time     <fct> 48, 48, 48, 48, 48, 48, 48, 48, 48, 48, 48, 48, 48, 48, 48, 4~
## $ Gene     <chr> "SOX2", "SIX6", "PAX6", "RPE65", "TYR", "OTX2", "SOX2", "SIX6~
## $ RGE      <dbl> 1.3200000, 1.5510985, 0.9964433, 0.7200000, 1.2113423, 1.1549~
```

```r
d7D_long <- d7D %>%
  rename(Time=`Time.(h)`) %>%
  pivot_longer(cols=7:10,names_to="Gene",values_to="RGE") %>%
  mutate(Time=as.factor(Time))
glimpse(d7D_long)
```

```
## Rows: 128
## Columns: 8
## $ Chemical <chr> "PBS", "PBS", "PBS", "PBS", "PBS", "PBS", "PBS", "PBS", "PBS"~
## $ DCA      <dbl> 0, 0, 0, 0, 0, 0, 0, 0, 0, 0, 0, 0, 0, 0, 0, 0, 1, 1, 1, 1, 1~
## $ NAC      <dbl> 0, 0, 0, 0, 0, 0, 0, 0, 0, 0, 0, 0, 0, 0, 0, 0, 0, 0, 0, 0, 0~
## $ FGF2     <dbl> 0, 0, 0, 0, 0, 0, 0, 0, 0, 0, 0, 0, 0, 0, 0, 0, 0, 0, 0, 0, 0~
## $ Sample   <dbl> 1, 1, 1, 1, 2, 2, 2, 2, 3, 3, 3, 3, 4, 4, 4, 4, 1, 1, 1, 1, 2~
## $ Time     <fct> 48, 48, 48, 48, 48, 48, 48, 48, 48, 48, 48, 48, 48, 48, 48, 4~
## $ Gene     <chr> "TGFB2", "VIM", "SNAI1", "aSMA", "TGFB2", "VIM", "SNAI1", "aS~
## $ RGE      <dbl> 1.5028399, 0.7755005, 0.6211773, 0.7726269, 1.2524987, 1.1842~
```

```r
d7E_long <- d7E %>%
  rename(Time=`Time.(h)`) %>%
  pivot_longer(cols=7:8,names_to="Gene",values_to="RGE") %>%
  mutate(Time=as.factor(Time))
glimpse(d7E_long)
```

```
## Rows: 64
## Columns: 8
## $ Chemical <chr> "PBS", "PBS", "PBS", "PBS", "PBS", "PBS", "PBS", "PBS", "DCA"~
## $ DCA      <dbl> 0, 0, 0, 0, 0, 0, 0, 0, 1, 1, 1, 1, 1, 1, 1, 1, 0, 0, 0, 0, 0~
## $ NAC      <dbl> 0, 0, 0, 0, 0, 0, 0, 0, 0, 0, 0, 0, 0, 0, 0, 0, 1, 1, 1, 1, 1~
## $ FGF2     <dbl> 0, 0, 0, 0, 0, 0, 0, 0, 0, 0, 0, 0, 0, 0, 0, 0, 0, 0, 0, 0, 0~
## $ Sample   <dbl> 1, 1, 2, 2, 3, 3, 4, 4, 1, 1, 2, 2, 3, 3, 4, 4, 1, 1, 2, 2, 3~
## $ Time     <fct> 48, 48, 48, 48, 48, 48, 48, 48, 48, 48, 48, 48, 48, 48, 48, 4~
## $ Gene     <chr> "E2F1", "PCNA", "E2F1", "PCNA", "E2F1", "PCNA", "E2F1", "PCNA~
## $ RGE      <dbl> 0.8876454, 1.1070841, 1.0116427, 1.1398686, 0.9026918, 0.8395~
```

```r
ggplot(d7C_long, aes(x=Chemical,y=log2(RGE))) +
  geom_jitter(width=.05,height=0) +
  facet_wrap(~Gene, nrow=3) +
  theme(axis.text.x = element_text(angle = 90, vjust = 0.5, hjust=1)) +
  labs(title="For Figure 7C: Treatment Comparisons")
```

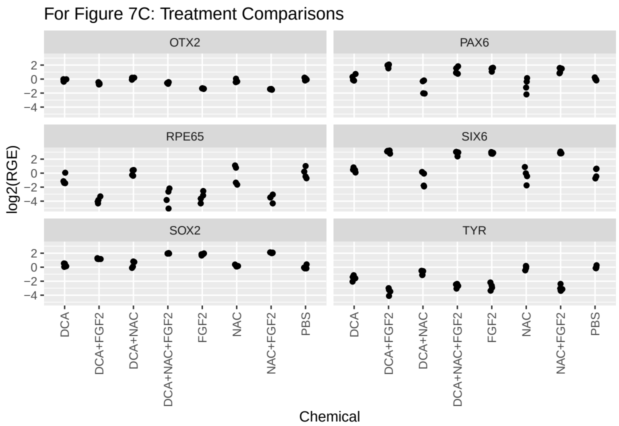

```r
ggplot(d7D_long, aes(x=Chemical,y=log2(RGE))) +
  geom_jitter(width=.05,height=0) +
  facet_wrap(~Gene, nrow=3) +
  theme(axis.text.x = element_text(angle = 90, vjust = 0.5, hjust=1)) +
  labs(title="For Figure 7D: Treatment Comparisons")
```

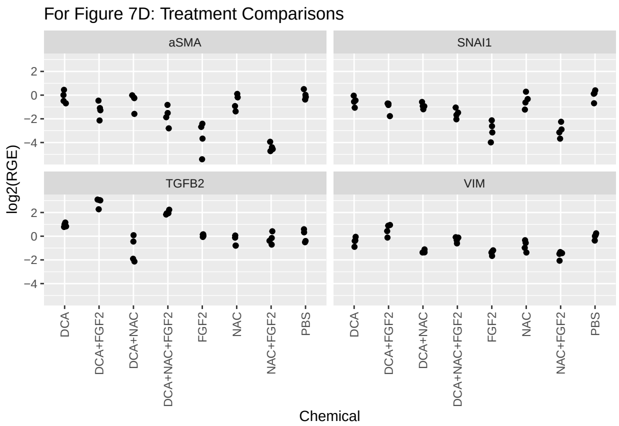

```r
ggplot(d7E_long, aes(x=Chemical,y=log2(RGE))) +
  geom_jitter(width=.05,height=0) +
  facet_wrap(~Gene, nrow=3) +
  theme(axis.text.x = element_text(angle = 90, vjust = 0.5, hjust=1)) +
  labs(title="For Figure 7E: Treatment Comparisons")
```

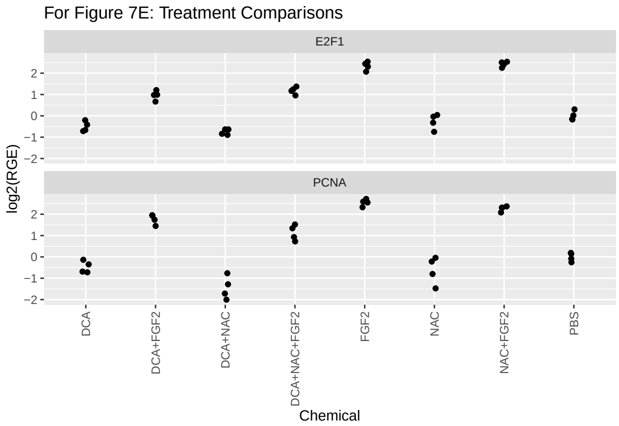

#### Data Analysis (Experiment 8)

The main analysis here is to compare nine particular pairs of treatments.

##### Analysis (Figure 7C)

**Treatment Comparisons (Figure 7C)** First the one-way ANOVAs.

```
a7C_SOX2 <- lm(log2(SOX2)~Chemical, data=d7C)
a7C_SIX6 <- lm(log2(SIX6)~Chemical, data=d7C)
a7C_PAX6 <- lm(log2(PAX6)~Chemical, data=d7C)
a7C_RPE65 <- lm(log2(RPE65)~Chemical, data=d7C)
a7C_TYR <- lm(log2(TYR)~Chemical, data=d7C)
a7C_OTX2 <- lm(log2(OTX2)~Chemical, data=d7C)

summary(a7C_SOX2)
```

```
##
## Call:
## lm(formula = log2(SOX2) ~ Chemical, data = d7C)
##
## Residuals:
##      Min       1Q   Median       3Q      Max
## -0.48839 -0.07999 -0.02009  0.06803  0.43188
##
## Coefficients:
##                        Estimate Std. Error t value Pr(>|t|)
## (Intercept)             0.30552    0.11061   2.762   0.0108 *
## ChemicalDCA+FGF2        0.89201    0.15643   5.702 7.12e-06 ***
## ChemicalDCA+NAC         0.07817    0.15643   0.500   0.6218
## ChemicalDCA+NAC+FGF2    1.65943    0.15643  10.608 1.53e-10 ***
## ChemicalFGF2            1.53236    0.15643   9.796 7.35e-10 ***
## ChemicalNAC            -0.11786    0.15643  -0.753   0.4585
## ChemicalNAC+FGF2        1.75125    0.15643  11.195 5.19e-11 ***
## ChemicalPBS            -0.30417    0.15643  -1.944   0.0637 .
## ---
## Signif. codes:  0 '***' 0.001 '**' 0.01 '*' 0.05 '.' 0.1 ' ' 1
```

```
##
## Residual standard error: 0.2212 on 24 degrees of freedom
## Multiple R-squared:  0.9476, Adjusted R-squared:  0.9323
## F-statistic: 61.97 on 7 and 24 DF,  p-value: 8.008e-14
```

```r
plot(a7C_SOX2,which=1:2)
```

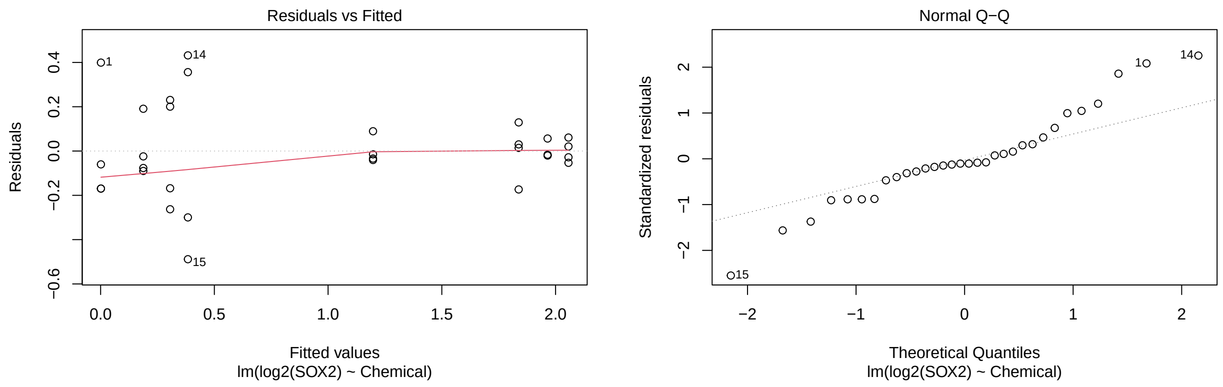

```r
plot(a7C_SIX6,which=1:2)
```

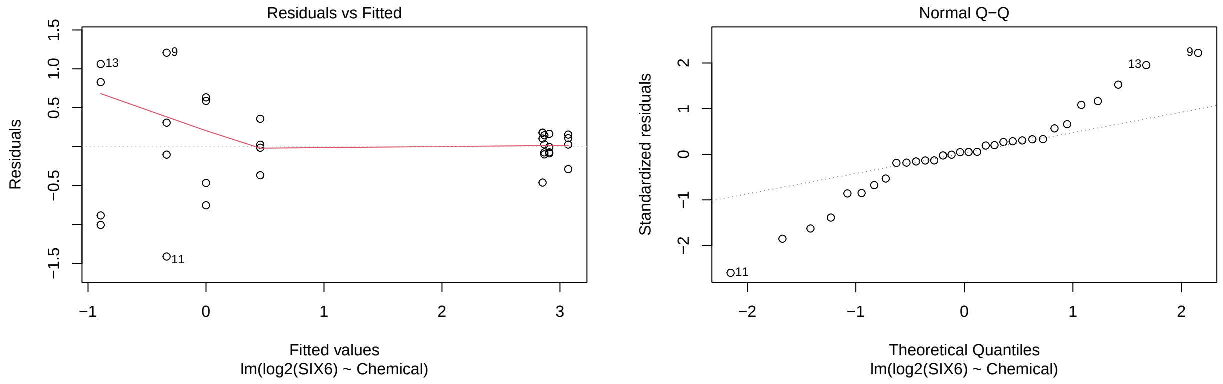

```r
plot(a7C_PAX6,which=1:2)
```

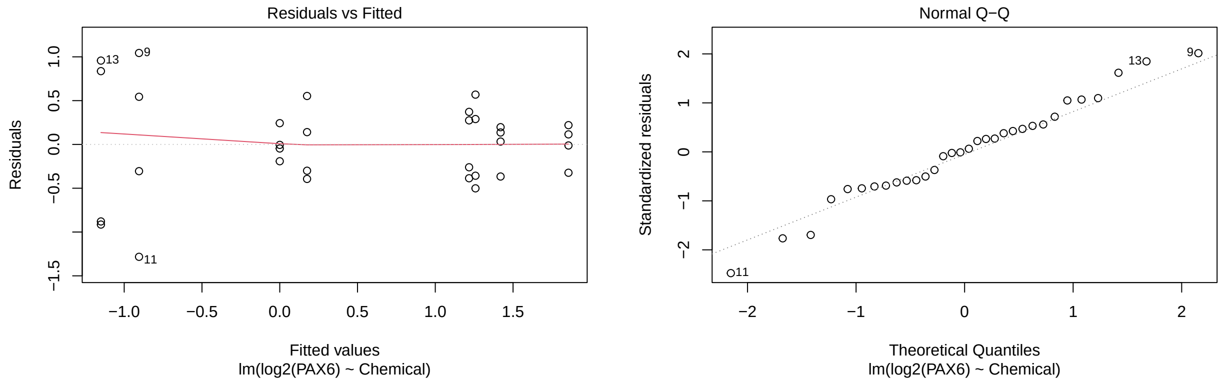

```r
plot(a7C_RPE65,which=1:2)
```

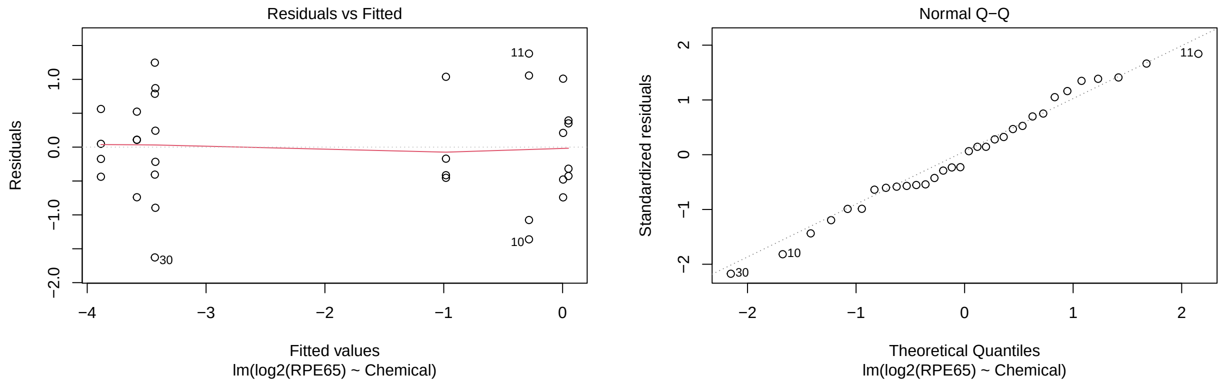

```r
plot(a7C_TYR,which=1:2)
```

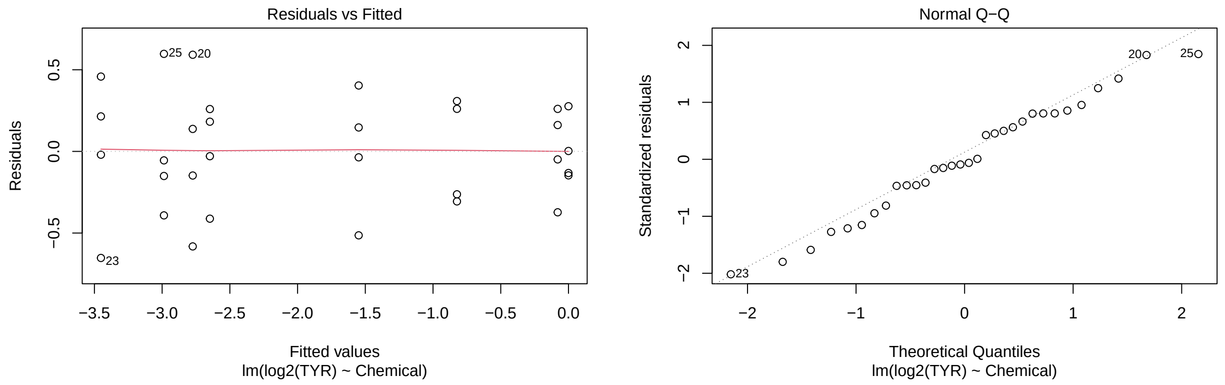

```r
plot(a7C_OTX2,which=1:2)
```

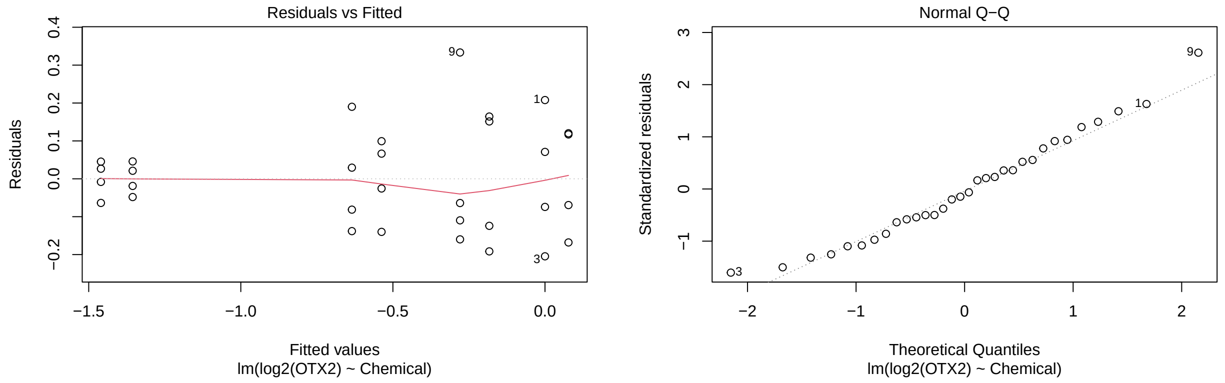

Comparisons for Figure 7C, 7D, 7E: PBS vs DCA; PBS vs NAC; PBS vs DCA+NAC; DCA vs DCA+NAC; PBS vs FGF2; FGF2 vs FGF2+DCA; FGF2 vs FGF2+NAC; FGF2 vs FGF2+DCA+NAC; FGF2+DCA vs FGF2+DCA+NAC for each gene. Assuming, as in the parameterization of `lm()`, a linear function of parameters of:

$$E(Y) = \beta_0 + \beta_1 DF + \beta_2 DN + \beta_3 DNF + \beta_4 F + \beta_5 N + \beta_6 NF + \beta_7 P,$$

where each variable represents an indicator variable for `DCA+FGF2`, `DCA+NAC`, etc. The baseline is the `DCA` treatment. Then the nine requested comparisons can be specified as below:

```
K <- matrix(c(0,0,0,0,0,0,0,1, # PBS vs. DCA
              0,0,0,0,0,-1,0,1, # PBS vs. NAC
              0,0,-1,0,0,0,0,1, # PBS vs. DCA+NAC
              0,0,-1,0,0,0,0,0, # DCA vs. DCA+NAC
              0,0,0,0,-1,0,0,1, # PBS vs. FGF2
              0,-1,0,0,1,0,0,0, # FGF2 vs. FGF2+DCA
              0,0,0,0,1,0,-1,0, # FGF2 vs. FGF2+NAC
              0,0,0,-1,1,0,0,0, # FGF2 vs. FGF2+DCA+NAC
              0,1,0,-1,0,0,0,0),nrow=9, byrow=TRUE) # FGF2+DCA vs. FGF2+DCA+NAC

t7C_SOX2 <- glht(a7C_SOX2, linfct=K)
summary(t7C_SOX2, test=adjusted("fdr"))
```

```
##
##   Simultaneous Tests for General Linear Hypotheses
##
## Fit: lm(formula = log2(SOX2) ~ Chemical, data = d7C)
##
## Linear Hypotheses:
##        Estimate Std. Error t value Pr(>|t|)
## 1 == 0 -0.30417    0.15643  -1.944 0.114576
## 2 == 0 -0.18631    0.15643  -1.191 0.315379
## 3 == 0 -0.38235    0.15643  -2.444 0.050046 .
## 4 == 0 -0.07817    0.15643  -0.500 0.621813
## 5 == 0 -1.83654    0.15643 -11.740 1.77e-10 ***
## 6 == 0  0.64036    0.15643   4.094 0.001248 **
## 7 == 0 -0.21889    0.15643  -1.399 0.261789
## 8 == 0 -0.12706    0.15643  -0.812 0.477710
## 9 == 0 -0.76742    0.15643  -4.906 0.000238 ***
## ---
## Signif. codes:  0 '***' 0.001 '**' 0.01 '*' 0.05 '.' 0.1 ' ' 1
## (Adjusted p values reported -- fdr method)
```

```
t7C_SIX6 <- glht(a7C_SIX6, linfct=K)
summary(t7C_SIX6, test=adjusted("fdr"))
```

```
##
##   Simultaneous Tests for General Linear Hypotheses
##
## Fit: lm(formula = log2(SIX6) ~ Chemical, data = d7C)
##
## Linear Hypotheses:
##        Estimate Std. Error t value Pr(>|t|)
## 1 == 0 -0.45971    0.44387  -1.036    0.699
## 2 == 0  0.33388    0.44387   0.752    0.827
```

```
## 3 == 0  0.89268    0.44387   2.011    0.167
## 4 == 0  1.35239    0.44387   3.047    0.025 *
## 5 == 0 -2.86768    0.44387  -6.461 9.99e-06 ***
## 6 == 0 -0.20244    0.44387  -0.456    0.839
## 7 == 0 -0.04173    0.44387  -0.094    0.973
## 8 == 0  0.01506    0.44387   0.034    0.973
## 9 == 0  0.21751    0.44387   0.490    0.839
## ---
## Signif. codes:  0 '***' 0.001 '**' 0.01 '*' 0.05 '.' 0.1 ' ' 1
## (Adjusted p values reported -- fdr method)
```

```
t7C_PAX6 <- glht(a7C_PAX6, linfct=K)
summary(t7C_PAX6, test=adjusted("fdr"))
```

```
##
##   Simultaneous Tests for General Linear Hypotheses
##
## Fit: lm(formula = log2(PAX6) ~ Chemical, data = d7C)
##
## Linear Hypotheses:
##         Estimate Std. Error t value Pr(>|t|)
## 1 == 0  -0.1756     0.4231  -0.415   0.7062
## 2 == 0   0.9047     0.4231   2.138   0.0965 .
## 3 == 0   1.1497     0.4231   2.717   0.0361 *
## 4 == 0   1.3253     0.4231   3.132   0.0204 *
## 5 == 0  -1.4198     0.4231  -3.355   0.0204 *
## 6 == 0  -0.4366     0.4231  -1.032   0.4687
## 7 == 0   0.2024     0.4231   0.478   0.7062
## 8 == 0   0.1614     0.4231   0.381   0.7062
## 9 == 0   0.5980     0.4231   1.413   0.3068
## ---
## Signif. codes:  0 '***' 0.001 '**' 0.01 '*' 0.05 '.' 0.1 ' ' 1
## (Adjusted p values reported -- fdr method)
```

```
t7C_RPE65 <- glht(a7C_RPE65, linfct=K)
summary(t7C_RPE65, test=adjusted("fdr"))
```

```
##
##   Simultaneous Tests for General Linear Hypotheses
##
## Fit: lm(formula = log2(RPE65) ~ Chemical, data = d7C)
##
## Linear Hypotheses:
##          Estimate Std. Error t value Pr(>|t|)
## 1 == 0  0.985834   0.611648   1.612    0.360
## 2 == 0  0.287072   0.611648   0.469    0.965
## 3 == 0 -0.045169   0.611648  -0.074    0.994
## 4 == 0 -1.031002   0.611648  -1.686    0.360
## 5 == 0  3.431046   0.611648   5.610 8.07e-05 ***
## 6 == 0  0.458162   0.611648   0.749    0.837
## 7 == 0  0.155520   0.611648   0.254    0.994
## 8 == 0  0.004268   0.611648   0.007    0.994
## 9 == 0 -0.453894   0.611648  -0.742    0.837
```

```
## ---
## Signif. codes:  0 '***' 0.001 '**' 0.01 '*' 0.05 '.' 0.1 ' ' 1
## (Adjusted p values reported -- fdr method)
```

```
t7C_TYR <- glht(a7C_TYR, linfct=K)
summary(t7C_TYR, test=adjusted("fdr"))
```

```
##
##   Simultaneous Tests for General Linear Hypotheses
##
## Fit: lm(formula = log2(TYR) ~ Chemical, data = d7C)
##
## Linear Hypotheses:
##        Estimate Std. Error t value Pr(>|t|)
## 1 == 0  1.54895    0.26402   5.867 2.13e-05 ***
## 2 == 0  0.07963    0.26402   0.302   0.7655
## 3 == 0  0.82324    0.26402   3.118   0.0125 *
## 4 == 0 -0.72572    0.26402  -2.749   0.0201 *
## 5 == 0  2.77406    0.26402  10.507 1.67e-09 ***
## 6 == 0  0.67789    0.26402   2.568   0.0253 *
## 7 == 0  0.21260    0.26402   0.805   0.5510
## 8 == 0 -0.12657    0.26402  -0.479   0.7155
## 9 == 0 -0.80445    0.26402  -3.047   0.0125 *
## ---
## Signif. codes:  0 '***' 0.001 '**' 0.01 '*' 0.05 '.' 0.1 ' ' 1
## (Adjusted p values reported -- fdr method)
```

```
t7C_OTX2 <- glht(a7C_OTX2, linfct=K)
summary(t7C_OTX2, test=adjusted("fdr"))
```

```
##
##   Simultaneous Tests for General Linear Hypotheses
##
## Fit: lm(formula = log2(OTX2) ~ Chemical, data = d7C)
##
## Linear Hypotheses:
##        Estimate Std. Error t value Pr(>|t|)
## 1 == 0  0.18307    0.10418   1.757   0.1375
## 2 == 0  0.27927    0.10418   2.681   0.0294 *
## 3 == 0 -0.07721    0.10418  -0.741   0.4658
## 4 == 0 -0.26028    0.10418  -2.498   0.0355 *
## 5 == 0  1.35551    0.10418  13.011 2.07e-11 ***
## 6 == 0 -0.72073    0.10418  -6.918 1.12e-06 ***
## 7 == 0  0.10436    0.10418   1.002   0.4034
## 8 == 0 -0.81825    0.10418  -7.854 1.96e-07 ***
## 9 == 0 -0.09752    0.10418  -0.936   0.4034
## ---
## Signif. codes:  0 '***' 0.001 '**' 0.01 '*' 0.05 '.' 0.1 ' ' 1
## (Adjusted p values reported -- fdr method)
```

**Three-way ANOVA (Figure 7C)** Here is a three-way ANOVA which allows us to investigate interactions. We did not use this analysis explicitly in the paper, with the exception of the comparison used to test the hypothesis that DCA and NAC cancel out the effect of FGF2; this can be found below.

```r
af7C_SOX2 <- lm(log2(SOX2)~DCA*NAC*FGF2, data=d7C)
af7C_SIX6 <- lm(log2(SIX6)~DCA*NAC*FGF2, data=d7C)
af7C_PAX6 <- lm(log2(PAX6)~DCA*NAC*FGF2, data=d7C)
af7C_RPE65 <- lm(log2(RPE65)~DCA*NAC*FGF2, data=d7C)
af7C_TYR <- lm(log2(TYR)~DCA*NAC*FGF2, data=d7C)
af7C_OTX2 <- lm(log2(OTX2)~DCA*NAC*FGF2, data=d7C)

summary(af7C_SOX2)
```

```
##
## Call:
## lm(formula = log2(SOX2) ~ DCA * NAC * FGF2, data = d7C)
##
## Residuals:
##      Min       1Q   Median       3Q      Max
## -0.48839 -0.07999 -0.02009  0.06803  0.43188
##
## Coefficients:
##              Estimate Std. Error t value Pr(>|t|)
## (Intercept)   0.00135    0.11062   0.012 0.990366
## DCA           0.30417    0.15643   1.944 0.063653 .
## NAC           0.18631    0.15643   1.191 0.245295
## FGF2          1.83654    0.15643  11.740 1.96e-11 ***
## DCA:NAC      -0.10814    0.22123  -0.489 0.629414
## DCA:FGF2     -0.94453    0.22123  -4.269 0.000266 ***
## NAC:FGF2      0.03258    0.22123   0.147 0.884160
## DCA:NAC:FGF2  0.65667    0.31287   2.099 0.046528 *
## ---
## Signif. codes:  0 '***' 0.001 '**' 0.01 '*' 0.05 '.' 0.1 ' ' 1
##
## Residual standard error: 0.2212 on 24 degrees of freedom
## Multiple R-squared:  0.9476, Adjusted R-squared:  0.9323
## F-statistic: 61.97 on 7 and 24 DF,  p-value: 8.008e-14
```

```r
ggplot(d7C) +
  aes(x = FGF2, color = as.factor(DCA), group = as.factor(DCA), y = log2(SOX2)) +
  stat_summary(fun = mean, geom = "point") +
  stat_summary(fun = mean, geom = "line") +
  geom_point()
```

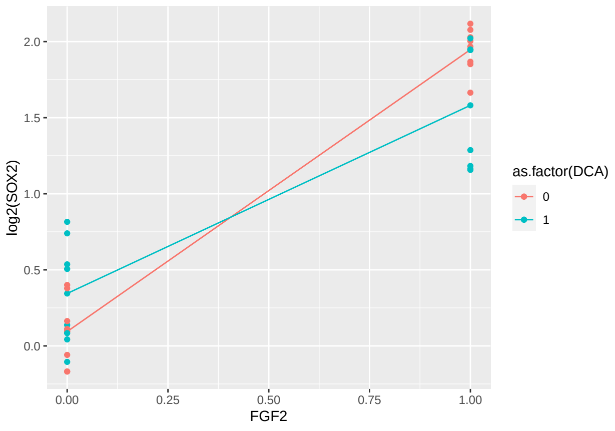

```r
summary(af7C_SIX6)
```

```
##
## Call:
## lm(formula = log2(SIX6) ~ DCA * NAC * FGF2, data = d7C)
##
## Residuals:
##      Min       1Q   Median       3Q      Max
## -1.41305 -0.14957  0.02561  0.17875  1.20789
##
## Coefficients:
##                Estimate Std. Error t value Pr(>|t|)
## (Intercept)  -1.676e-09  3.139e-01   0.000    1.000
## DCA           4.597e-01  4.439e-01   1.036    0.311
## NAC          -3.339e-01  4.439e-01  -0.752    0.459
## FGF2          2.868e+00  4.439e-01   6.461 1.11e-06 ***
## DCA:NAC      -1.019e+00  6.277e-01  -1.623    0.118
## DCA:FGF2     -2.573e-01  6.277e-01  -0.410    0.686
## NAC:FGF2      3.756e-01  6.277e-01   0.598    0.555
## DCA:NAC:FGF2  7.593e-01  8.877e-01   0.855    0.401
## ---
## Signif. codes:  0 '***' 0.001 '**' 0.01 '*' 0.05 '.' 0.1 ' ' 1
##
## Residual standard error: 0.6277 on 24 degrees of freedom
## Multiple R-squared:  0.8963, Adjusted R-squared:  0.866
## F-statistic: 29.63 on 7 and 24 DF,  p-value: 2.537e-10
```

```r
ggplot(d7C) +
  aes(x = FGF2, y = log2(SIX6)) +
  stat_summary(fun = mean, geom = "point") +
  stat_summary(fun = mean, geom = "line") +
  geom_point()
```

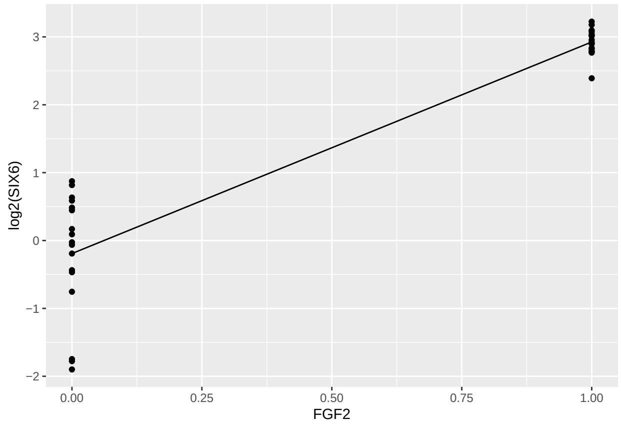

```r
summary(af7C_PAX6)
```

```
##
## Call:
```

```
## lm(formula = log2(PAX6) ~ DCA * NAC * FGF2, data = d7C)
##
## Residuals:
##      Min       1Q   Median       3Q      Max
## -1.28283 -0.33171  0.01379  0.27857  1.04398
##
## Coefficients:
##                Estimate Std. Error t value Pr(>|t|)
## (Intercept)  -2.993e-09  2.992e-01   0.000  1.00000
## DCA           1.756e-01  4.231e-01   0.415  0.68189
## NAC          -9.047e-01  4.231e-01  -2.138  0.04290 *
## FGF2          1.420e+00  4.231e-01   3.355  0.00263 **
## DCA:NAC      -4.206e-01  5.984e-01  -0.703  0.48891
## DCA:FGF2      2.610e-01  5.984e-01   0.436  0.66661
## NAC:FGF2      7.023e-01  5.984e-01   1.174  0.25204
## DCA:NAC:FGF2  2.497e-02  8.463e-01   0.030  0.97671
## ---
## Signif. codes:  0 '***' 0.001 '**' 0.01 '*' 0.05 '.' 0.1 ' ' 1
##
## Residual standard error: 0.5984 on 24 degrees of freedom
## Multiple R-squared:  0.8042, Adjusted R-squared:  0.7471
## F-statistic: 14.08 on 7 and 24 DF,  p-value: 4.065e-07
```

```r
ggplot(d7C) +
  aes(x = FGF2, y = log2(PAX6)) +
  stat_summary(fun = mean, geom = "point") +
  stat_summary(fun = mean, geom = "line") +
  geom_point()
```

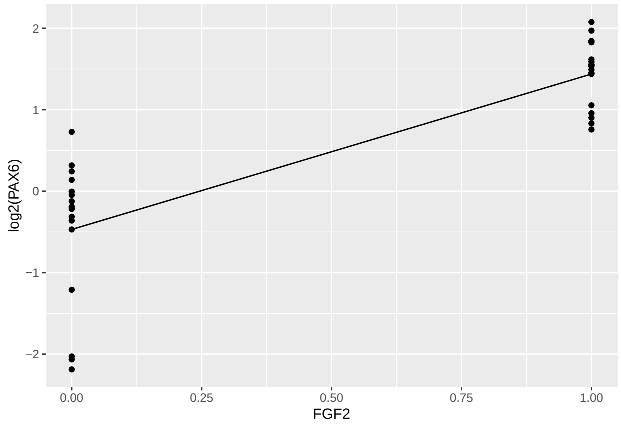

```r
ggplot(d7C) +
  aes(x = NAC, y = log2(PAX6)) +
  stat_summary(fun = mean, geom = "point") +
  stat_summary(fun = mean, geom = "line") +
  geom_point()
```

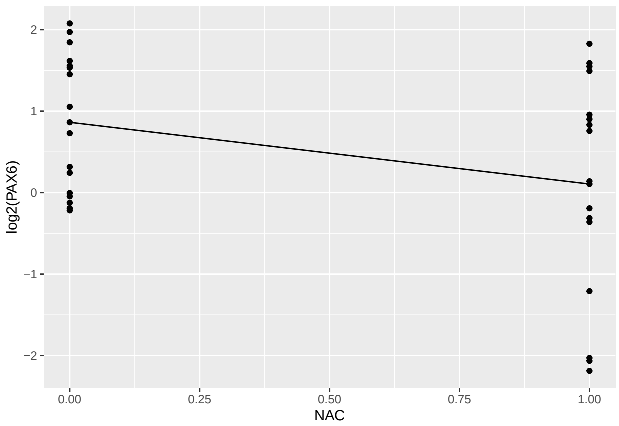

```r
summary(af7C_RPE65)
```

```
##
## Call:
## lm(formula = log2(RPE65) ~ DCA * NAC * FGF2, data = d7C)
##
## Residuals:
##      Min       1Q   Median       3Q      Max
## -1.62797 -0.44108 -0.06113  0.53318  1.38029
##
## Coefficients:
##               Estimate Std. Error t value Pr(>|t|)
## (Intercept)   0.004396   0.432501   0.010    0.992
## DCA          -0.985834   0.611648  -1.612    0.120
## NAC          -0.287072   0.611648  -0.469    0.643
## FGF2         -3.431046   0.611648  -5.610 8.97e-06 ***
## DCA:NAC       1.318074   0.865001   1.524    0.141
## DCA:FGF2      0.527671   0.865001   0.610    0.548
## NAC:FGF2      0.131551   0.865001   0.152    0.880
## DCA:NAC:FGF2 -0.708660   1.223296  -0.579    0.568
## ---
## Signif. codes:  0 '***' 0.001 '**' 0.01 '*' 0.05 '.' 0.1 ' ' 1
##
## Residual standard error: 0.865 on 24 degrees of freedom
## Multiple R-squared:  0.8325, Adjusted R-squared:  0.7837
## F-statistic: 17.04 on 7 and 24 DF,  p-value: 6.733e-08
```

```r
ggplot(d7C) +
  aes(x = FGF2, y = log2(RPE65)) +
  stat_summary(fun = mean, geom = "point") +
  stat_summary(fun = mean, geom = "line") +
  geom_point()
```

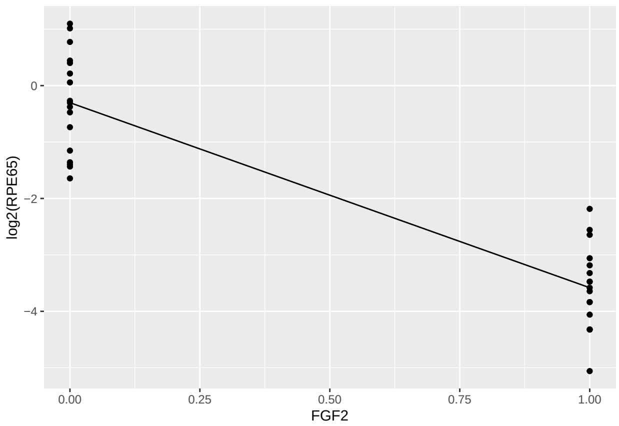

```r
summary(af7C_TYR)
```

```
##
## Call:
## lm(formula = log2(TYR) ~ DCA * NAC * FGF2, data = d7C)
##
## Residuals:
##      Min       1Q   Median       3Q      Max
## -0.65252 -0.17867 -0.02479  0.25967  0.59740
##
## Coefficients:
##                Estimate Std. Error t value Pr(>|t|)
## (Intercept)   9.560e-10  1.867e-01   0.000   1.0000
## DCA          -1.549e+00  2.640e-01  -5.867 4.73e-06 ***
## NAC          -7.963e-02  2.640e-01  -0.302   0.7655
## FGF2         -2.774e+00  2.640e-01 -10.507 1.85e-10 ***
## DCA:NAC       8.053e-01  3.734e-01   2.157   0.0412 *
## DCA:FGF2      8.711e-01  3.734e-01   2.333   0.0284 *
## NAC:FGF2     -1.330e-01  3.734e-01  -0.356   0.7249
## DCA:NAC:FGF2  2.117e-01  5.280e-01   0.401   0.6920
## ---
## Signif. codes:  0 '***' 0.001 '**' 0.01 '*' 0.05 '.' 0.1 ' ' 1
##
## Residual standard error: 0.3734 on 24 degrees of freedom
## Multiple R-squared:  0.9396, Adjusted R-squared:  0.922
## F-statistic: 53.37 on 7 and 24 DF,  p-value: 4.269e-13
```

```r
ggplot(d7C) +
  aes(x = FGF2, color = as.factor(DCA), group = as.factor(DCA), y = log2(TYR)) +
  stat_summary(fun = mean, geom = "point") +
  stat_summary(fun = mean, geom = "line") +
  geom_point()
```

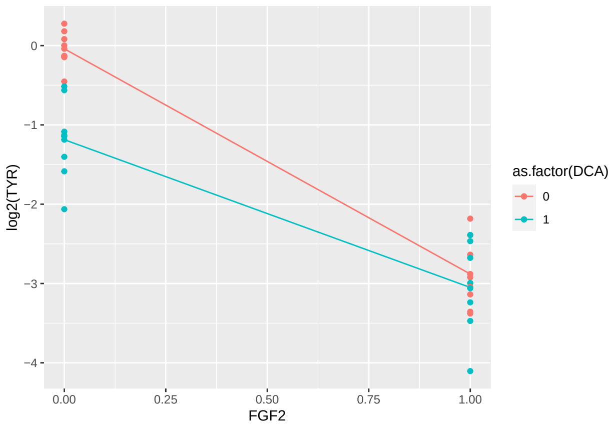

```r
ggplot(d7C) +
  aes(x = DCA, color = as.factor(NAC), group = as.factor(NAC), y = log2(TYR)) +
  stat_summary(fun = mean, geom = "point") +
  stat_summary(fun = mean, geom = "line") +
  geom_point()
```

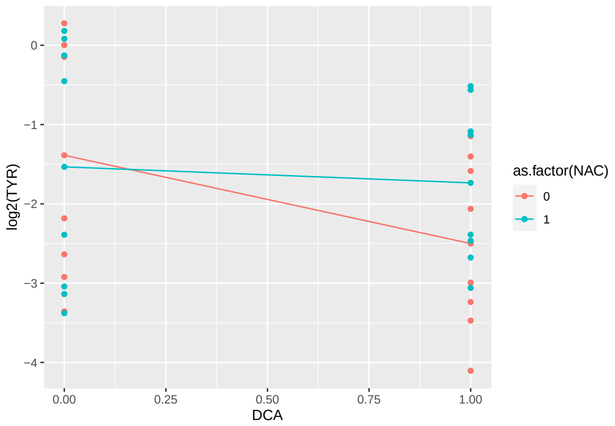

```r
summary(af7C_OTX2)
```

```
##
## Call:
## lm(formula = log2(OTX2) ~ DCA * NAC * FGF2, data = d7C)
##
## Residuals:
##      Min       1Q   Median       3Q      Max
## -0.20456 -0.08847 -0.01354  0.07804  0.33342
##
## Coefficients:
##                Estimate Std. Error t value Pr(>|t|)
## (Intercept)  -2.977e-09  7.367e-02   0.000  1.00000
## DCA          -1.831e-01  1.042e-01  -1.757  0.09164 .
## NAC          -2.793e-01  1.042e-01  -2.681  0.01307 *
## FGF2         -1.356e+00  1.042e-01 -13.011 2.30e-12 ***
## DCA:NAC       5.395e-01  1.473e-01   3.662  0.00123 **
## DCA:FGF2      9.038e-01  1.473e-01   6.134 2.45e-06 ***
```

```
## NAC:FGF2      1.749e-01  1.473e-01   1.187  0.24680
## DCA:NAC:FGF2 -3.377e-01  2.084e-01  -1.621  0.11818
## ---
## Signif. codes:  0 '***' 0.001 '**' 0.01 '*' 0.05 '.' 0.1 ' ' 1
##
## Residual standard error: 0.1473 on 24 degrees of freedom
## Multiple R-squared:  0.9483, Adjusted R-squared:  0.9332
## F-statistic: 62.86 on 7 and 24 DF,  p-value: 6.825e-14
```

```r
ggplot(d7C) +
  aes(x = FGF2, color = as.factor(DCA), group = as.factor(DCA), y = log2(OTX2)) +
  stat_summary(fun = mean, geom = "point") +
  stat_summary(fun = mean, geom = "line") +
  geom_point()
```

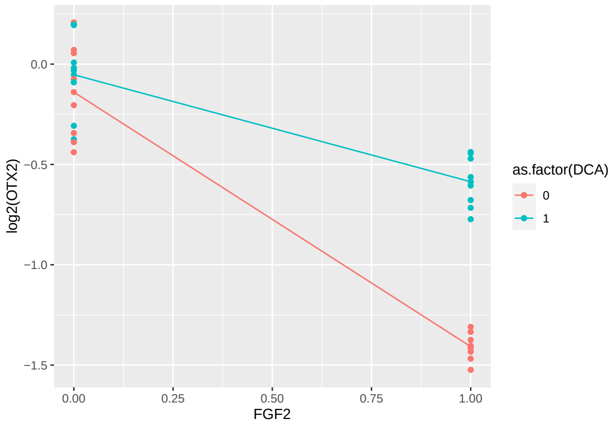

```r
ggplot(d7C) +
  aes(x = DCA, color = as.factor(NAC), group = as.factor(NAC), y = log2(OTX2)) +
  stat_summary(fun = mean, geom = "point") +
  stat_summary(fun = mean, geom = "line") +
  geom_point()
```

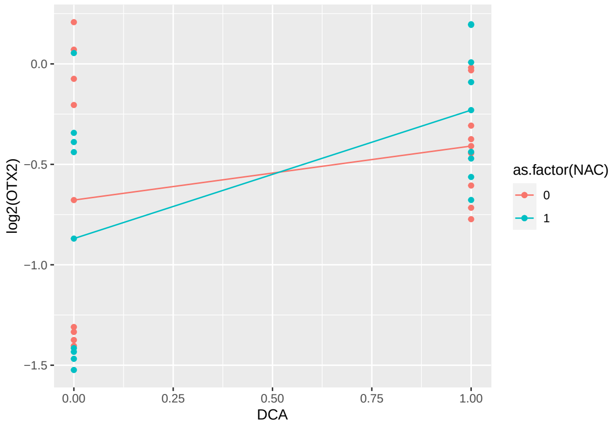

We will do one additional specific analysis, based on the three-way ANOVA. The hypothesis is that for SOX2, DCA and NAC "cancel out" the effect of FGF2. If $T_{DNF}$ represents the treatment which includes the presence of all three of the Chemicals, while $T_F$ is the treatment with FGF2 only. Assuming the following model:

$$E(\log_2[SOX2]) = \beta_0 + \beta_1 D + \beta_2 N + \beta_3 F + \beta_4 DN + \beta_5 DF + \beta_6 NF + \beta_7 DNF,$$

where SOX2 is the relative gene expression for the SOX2 gene; and $D$, $N$, and $F$ are indicators for DCA, NAC, and FGF2, respectively. Then, the treatment with all chemicals present is

$$E(\log_2[SOX2]|T_{DNF}) = \sum_{i=0}^{7} \beta_i,$$

whereas for the FGF2-only treatment we have

$$E(\log_2[SOX2]|T_F) = \beta_0 + \beta_3.$$

Thus, the difference between the two treatments is $C = \beta_1 + \beta_2 + \beta_4 + \beta_5 + \beta_6 + \beta_7$ and we can test the hypothesis of whether $H_0 : C = 0$.

```
K <- matrix(c(0,1,1,0,1,1,1,1),nrow=1, byrow=TRUE)

t <- glht(af7C_SOX2, linfct=K)
summary(t)
```

```
##
##   Simultaneous Tests for General Linear Hypotheses
##
## Fit: lm(formula = log2(SOX2) ~ DCA * NAC * FGF2, data = d7C)
##
## Linear Hypotheses:
##        Estimate Std. Error t value Pr(>|t|)
## 1 == 0   0.1271     0.1564   0.812    0.425
## (Adjusted p values reported -- single-step method)
```

There is no evidence of a difference between the two treatments. (Though the p-value is given as "adjusted", it is unadjusted since there is only one comparison.)

##### Analysis (Figure 7D)

**Treatment Comparisons (Figure 7D)** First the one-way ANOVAs.

```
a7D_TGFB2 <- lm(log2(TGFB2)~Chemical, data=d7D)
a7D_VIM <- lm(log2(VIM)~Chemical, data=d7D)
a7D_SNAI1 <- lm(log2(SNAI1)~Chemical, data=d7D)
a7D_aSMA <- lm(log2(aSMA)~Chemical, data=d7D)

summary(a7D_TGFB2)
```

```
##
## Call:
## lm(formula = log2(TGFB2) ~ Chemical, data = d7D)
##
## Residuals:
##      Min       1Q   Median       3Q      Max
## -1.03278 -0.22892  0.00443  0.23609  1.18572
```

```
##
## Coefficients:
##                      Estimate Std. Error t value Pr(>|t|)
## (Intercept)            0.9373     0.2571   3.645  0.00129 **
## ChemicalDCA+FGF2       1.9223     0.3637   5.286 2.02e-05 ***
## ChemicalDCA+NAC       -2.0386     0.3637  -5.606 9.05e-06 ***
## ChemicalDCA+NAC+FGF2   1.0456     0.3637   2.875  0.00833 **
## ChemicalFGF2          -0.8748     0.3637  -2.406  0.02422 *
## ChemicalNAC           -1.3496     0.3637  -3.711  0.00109 **
## ChemicalNAC+FGF2      -1.1463     0.3637  -3.152  0.00431 **
## ChemicalPBS           -0.9373     0.3637  -2.577  0.01653 *
## ---
## Signif. codes:  0 '***' 0.001 '**' 0.01 '*' 0.05 '.' 0.1 ' ' 1
##
## Residual standard error: 0.5143 on 24 degrees of freedom
## Multiple R-squared:  0.8857, Adjusted R-squared:  0.8524
## F-statistic: 26.57 on 7 and 24 DF,  p-value: 7.922e-10
```

```r
plot(a7D_TGFB2,which=1:2)
```

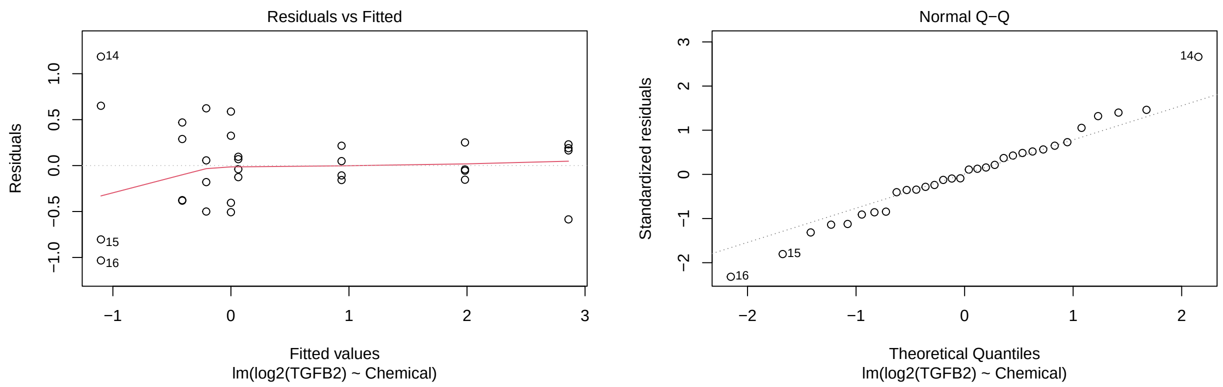

```r
plot(a7D_VIM,which=1:2)
```

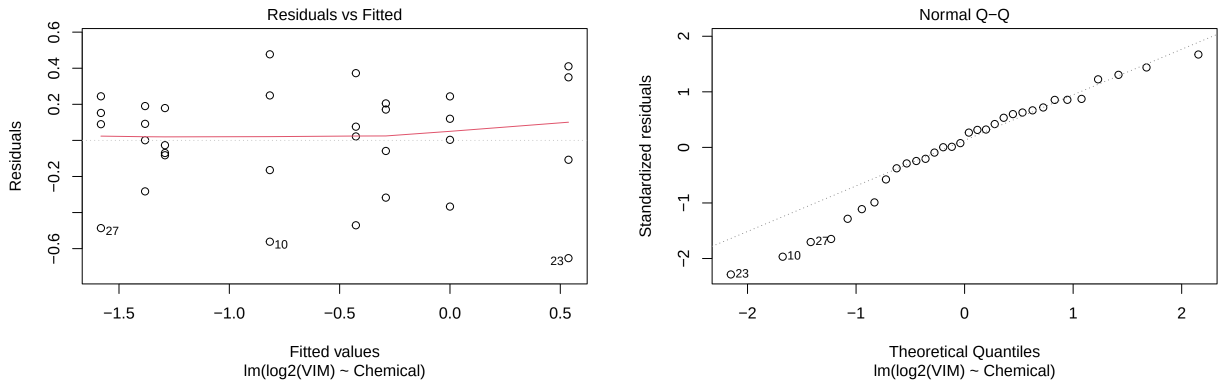

```r
plot(a7D_SNAI1,which=1:2)
```

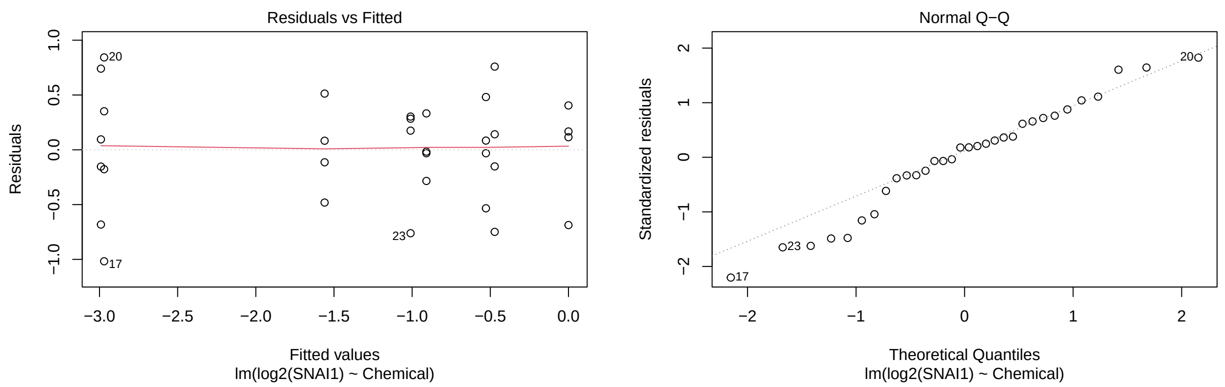

```r
plot(a7D_aSMA,which=1:2)
```

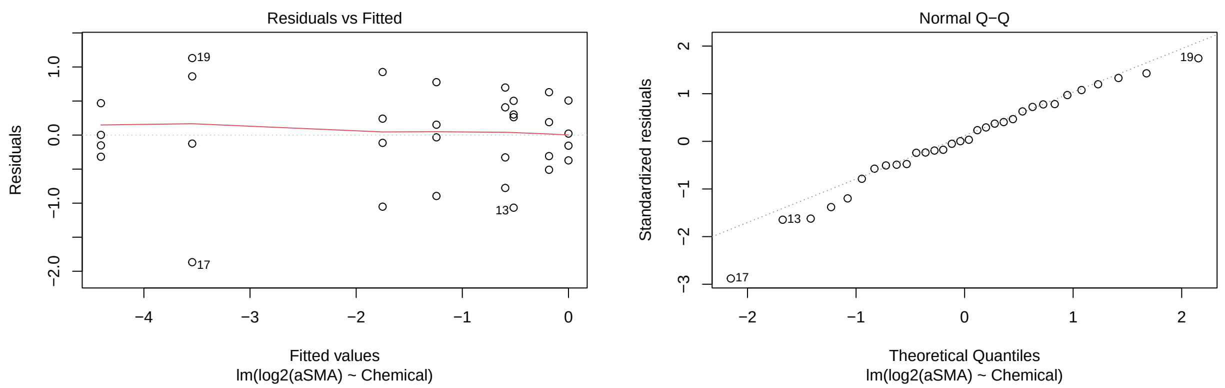

```r
K <- matrix(c(0,0,0,0,0,0,0,1, # PBS vs. DCA
              0,0,0,0,0,-1,0,1, # PBS vs. NAC
              0,0,-1,0,0,0,0,1, # PBS vs. DCA+NAC
              0,0,-1,0,0,0,0,0, # DCA vs. DCA+NAC
              0,0,0,0,-1,0,0,1, # PBS vs. FGF2
              0,-1,0,0,1,0,0,0, # FGF2 vs. FGF2+DCA
              0,0,0,0,1,0,-1,0, # FGF2 vs. FGF2+NAC
              0,0,0,-1,1,0,0,0, # FGF2 vs. FGF2+DCA+NAC
              0,1,0,-1,0,0,0,0),nrow=9, byrow=TRUE) # FGF2+DCA vs. FGF2+DCA+NAC

t7D_TGFB2 <- glht(a7D_TGFB2, linfct=K)
summary(t7D_TGFB2, test=adjusted("fdr"))
```

```
##
##   Simultaneous Tests for General Linear Hypotheses
##
## Fit: lm(formula = log2(TGFB2) ~ Chemical, data = d7D)
##
## Linear Hypotheses:
```

```
##        Estimate Std. Error t value Pr(>|t|)
## 1 == 0 -0.93727    0.36366  -2.577   0.0298 *
## 2 == 0  0.41229    0.36366   1.134   0.3447
## 3 == 0  1.10133    0.36366   3.028   0.0130 *
## 4 == 0  2.03860    0.36366   5.606 4.07e-05 ***
## 5 == 0 -0.06248    0.36366  -0.172   0.8650
## 6 == 0 -2.79705    0.36366  -7.691 5.66e-07 ***
## 7 == 0  0.27151    0.36366   0.747   0.5204
## 8 == 0 -1.92036    0.36366  -5.281 6.14e-05 ***
## 9 == 0  0.87668    0.36366   2.411   0.0359 *
## ---
## Signif. codes:  0 '***' 0.001 '**' 0.01 '*' 0.05 '.' 0.1 ' ' 1
## (Adjusted p values reported -- fdr method)
```

```r
t7D_VIM <- glht(a7D_VIM, linfct=K)
summary(t7D_VIM, test=adjusted("fdr"))
```

```
##
##   Simultaneous Tests for General Linear Hypotheses
##
## Fit: lm(formula = log2(VIM) ~ Chemical, data = d7D)
##
## Linear Hypotheses:
##        Estimate Std. Error t value Pr(>|t|)
## 1 == 0   0.4264     0.2330   1.830  0.08970 .
## 2 == 0   0.8163     0.2330   3.503  0.00235 **
## 3 == 0   1.2917     0.2330   5.543 3.18e-05 ***
## 4 == 0   0.8652     0.2330   3.713  0.00195 **
## 5 == 0   1.3818     0.2330   5.929 1.83e-05 ***
## 6 == 0  -1.9188     0.2330  -8.234 1.70e-07 ***
## 7 == 0   0.2001     0.2330   0.859  0.39895
## 8 == 0  -1.0912     0.2330  -4.682  0.00021 ***
## 9 == 0   0.8276     0.2330   3.551  0.00235 **
## ---
## Signif. codes:  0 '***' 0.001 '**' 0.01 '*' 0.05 '.' 0.1 ' ' 1
## (Adjusted p values reported -- fdr method)
```

```r
t7D_SNAI1 <- glht(a7D_SNAI1, linfct=K)
summary(t7D_SNAI1, test=adjusted("fdr"))
```

```
##
##   Simultaneous Tests for General Linear Hypotheses
##
## Fit: lm(formula = log2(SNAI1) ~ Chemical, data = d7D)
##
## Linear Hypotheses:
##        Estimate Std. Error t value Pr(>|t|)
## 1 == 0  0.52800    0.37690   1.401 0.261069
## 2 == 0  0.47187    0.37690   1.252 0.286255
## 3 == 0  0.90893    0.37690   2.412 0.053765 .
## 4 == 0  0.38093    0.37690   1.011 0.362523
## 5 == 0  2.97037    0.37690   7.881  3.7e-07 ***
## 6 == 0 -1.96031    0.37690  -5.201 0.000112 ***
```

```
## 7 == 0  0.02053    0.37690   0.054 0.957010
## 8 == 0 -1.41044    0.37690  -3.742 0.003024 **
## 9 == 0  0.54987    0.37690   1.459 0.261069
## ---
## Signif. codes:  0 '***' 0.001 '**' 0.01 '*' 0.05 '.' 0.1 ' ' 1
## (Adjusted p values reported -- fdr method)
```

```
t7D_aSMA <- glht(a7D_aSMA, linfct=K)
summary(t7D_aSMA, test=adjusted("fdr"))
```

```
##
##   Simultaneous Tests for General Linear Hypotheses
##
## Fit: lm(formula = log2(aSMA) ~ Chemical, data = d7D)
##
## Linear Hypotheses:
##        Estimate Std. Error t value Pr(>|t|)
## 1 == 0   0.1831     0.5294   0.346 0.732439
## 2 == 0   0.5957     0.5294   1.125 0.446835
## 3 == 0   0.5166     0.5294   0.976 0.446835
## 4 == 0   0.3335     0.5294   0.630 0.601463
## 5 == 0   3.5447     0.5294   6.696 5.69e-06 ***
## 6 == 0  -2.3009     0.5294  -4.346 0.000985 ***
## 7 == 0   0.8601     0.5294   1.625 0.263824
## 8 == 0  -1.7937     0.5294  -3.388 0.007278 **
## 9 == 0   0.5072     0.5294   0.958 0.446835
## ---
## Signif. codes:  0 '***' 0.001 '**' 0.01 '*' 0.05 '.' 0.1 ' ' 1
## (Adjusted p values reported -- fdr method)
```

**Three-way ANOVA (Figure 7D)** Again, analysis not used in the paper; provided to provide additional insight, particularly when interactions are important.

```
af7D_TGFB2 <- lm(log2(TGFB2)~DCA*NAC*FGF2, data=d7D)
af7D_VIM <- lm(log2(VIM)~DCA*NAC*FGF2, data=d7D)
af7D_SNAI1 <- lm(log2(SNAI1)~DCA*NAC*FGF2, data=d7D)
af7D_aSMA <- lm(log2(aSMA)~DCA*NAC*FGF2, data=d7D)

summary(af7D_TGFB2)
```

```
##
## Call:
## lm(formula = log2(TGFB2) ~ DCA * NAC * FGF2, data = d7D)
##
## Residuals:
##      Min       1Q   Median       3Q      Max
## -1.03278 -0.22892  0.00443  0.23609  1.18572
##
## Coefficients:
##                  Estimate Std. Error t value Pr(>|t|)
## (Intercept)    1.756e-10  2.571e-01   0.000  1.00000
## DCA            9.373e-01  3.637e-01   2.577  0.01653 *
```

```
## NAC          -4.123e-01  3.637e-01  -1.134  0.26811
## FGF2          6.248e-02  3.637e-01   0.172  0.86502
## DCA:NAC      -1.626e+00  5.143e-01  -3.162  0.00421 **
## DCA:FGF2      1.860e+00  5.143e-01   3.616  0.00138 **
## NAC:FGF2      1.408e-01  5.143e-01   0.274  0.78663
## DCA:NAC:FGF2  1.021e+00  7.273e-01   1.404  0.17313
## ---
## Signif. codes:  0 '***' 0.001 '**' 0.01 '*' 0.05 '.' 0.1 ' ' 1
##
## Residual standard error: 0.5143 on 24 degrees of freedom
## Multiple R-squared:  0.8857, Adjusted R-squared:  0.8524
## F-statistic: 26.57 on 7 and 24 DF,  p-value: 7.922e-10
```

```r
ggplot(d7D) +
  aes(x = FGF2, color = as.factor(DCA), group = as.factor(DCA), y = log2(TGFB2)) +
  stat_summary(fun = mean, geom = "point") +
  stat_summary(fun = mean, geom = "line") +
  geom_point()
```

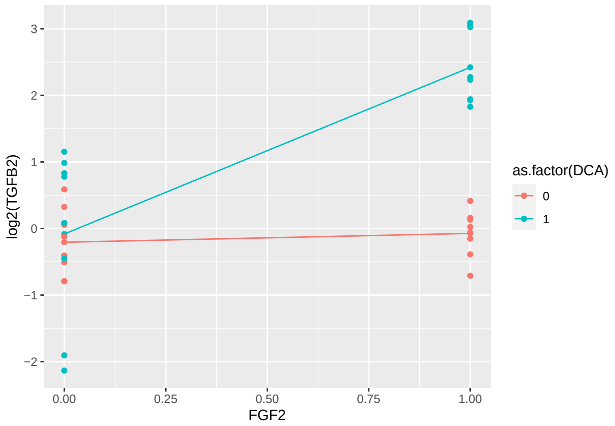

```r
ggplot(d7D) +
  aes(x = DCA, color = as.factor(NAC), group = as.factor(NAC), y = log2(TGFB2)) +
  stat_summary(fun = mean, geom = "point") +
  stat_summary(fun = mean, geom = "line") +
  geom_point()
```

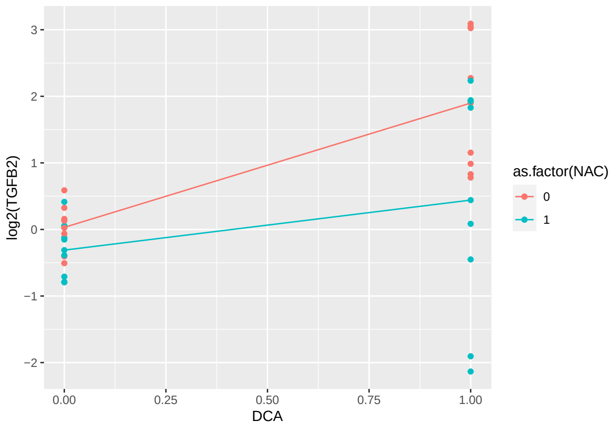

```r
summary(af7D_VIM)
```

```
##
## Call:
## lm(formula = log2(VIM) ~ DCA * NAC * FGF2, data = d7D)
##
## Residuals:
##      Min       1Q   Median       3Q      Max
## -0.65271 -0.12174  0.04891  0.19387  0.47704
##
## Coefficients:
##                Estimate Std. Error t value Pr(>|t|)
## (Intercept)  -1.824e-09  1.648e-01   0.000  1.00000
## DCA          -4.264e-01  2.330e-01  -1.830  0.07973 .
## NAC          -8.163e-01  2.330e-01  -3.503  0.00183 **
## FGF2         -1.382e+00  2.330e-01  -5.929 4.06e-06 ***
## DCA:NAC      -4.893e-02  3.296e-01  -0.148  0.88322
## DCA:FGF2      2.345e+00  3.296e-01   7.116 2.35e-07 ***
## NAC:FGF2      6.162e-01  3.296e-01   1.870  0.07378 .
## DCA:NAC:FGF2 -5.786e-01  4.661e-01  -1.241  0.22648
## ---
## Signif. codes:  0 '***' 0.001 '**' 0.01 '*' 0.05 '.' 0.1 ' ' 1
##
## Residual standard error: 0.3296 on 24 degrees of freedom
## Multiple R-squared:  0.8554, Adjusted R-squared:  0.8132
## F-statistic: 20.27 on 7 and 24 DF,  p-value: 1.234e-08
```

```r
ggplot(d7D) +
  aes(x = FGF2, color = as.factor(DCA), group = as.factor(DCA), y = log2(VIM)) +
  stat_summary(fun = mean, geom = "point") +
  stat_summary(fun = mean, geom = "line") +
  geom_point()
```

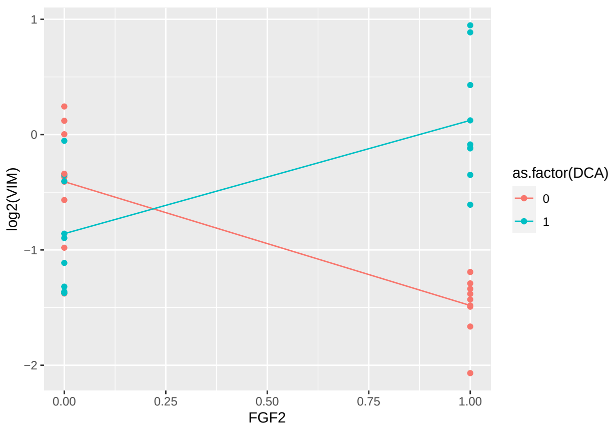

```r
ggplot(d7D) +
  aes(x = FGF2, color = as.factor(NAC), group = as.factor(NAC), y = log2(VIM)) +
  stat_summary(fun = mean, geom = "point") +
  stat_summary(fun = mean, geom = "line") +
  geom_point()
```

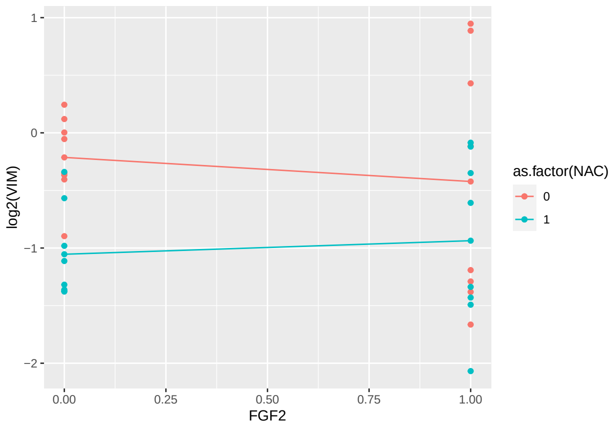

```r
summary(af7D_SNAI1)
```

```
##
## Call:
## lm(formula = log2(SNAI1) ~ DCA * NAC * FGF2, data = d7D)
##
## Residuals:
##      Min       1Q   Median       3Q      Max
## -1.01724 -0.20401  0.08337  0.31057  0.84287
##
## Coefficients:
##                 Estimate Std. Error t value Pr(>|t|)
## (Intercept)   -1.939e-09  2.665e-01   0.000    1.000
## DCA           -5.280e-01  3.769e-01  -1.401    0.174
## NAC           -4.719e-01  3.769e-01  -1.252    0.223
## FGF2          -2.970e+00  3.769e-01  -7.881 4.11e-08 ***
## DCA:NAC        9.094e-02  5.330e-01   0.171    0.866
## DCA:FGF2       2.488e+00  5.330e-01   4.668 9.65e-05 ***
## NAC:FGF2       4.513e-01  5.330e-01   0.847    0.405
## DCA:NAC:FGF2 -6.203e-01  7.538e-01  -0.823    0.419
## ---
## Signif. codes:  0 '***' 0.001 '**' 0.01 '*' 0.05 '.' 0.1 ' ' 1
##
## Residual standard error: 0.533 on 24 degrees of freedom
## Multiple R-squared:  0.8396, Adjusted R-squared:  0.7929
## F-statistic: 17.95 on 7 and 24 DF,  p-value: 4.077e-08
```

```r
ggplot(d7D) +
  aes(x = FGF2, color = as.factor(DCA), group = as.factor(DCA), y = log2(SNAI1)) +
  stat_summary(fun = mean, geom = "point") +
  stat_summary(fun = mean, geom = "line") +
  geom_point()
```

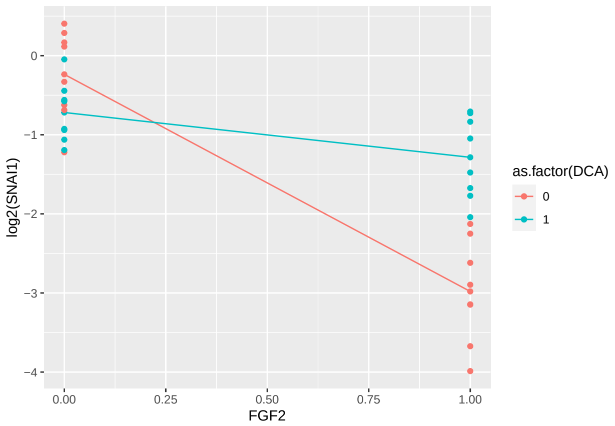

```r
summary(af7D_aSMA)
```

```
##
## Call:
## lm(formula = log2(aSMA) ~ DCA * NAC * FGF2, data = d7D)
##
## Residuals:
##      Min       1Q   Median       3Q      Max
## -1.86704 -0.32076  0.01172  0.47678  1.13044
##
## Coefficients:
##                Estimate Std. Error t value Pr(>|t|)
## (Intercept)  -4.217e-10  3.743e-01   0.000  1.00000
## DCA          -1.831e-01  5.294e-01  -0.346  0.73244
## NAC          -5.957e-01  5.294e-01  -1.125  0.27157
## FGF2         -3.545e+00  5.294e-01  -6.696 6.32e-07 ***
## DCA:NAC       2.622e-01  7.486e-01   0.350  0.72920
## DCA:FGF2      2.484e+00  7.486e-01   3.318  0.00288 **
## NAC:FGF2     -2.644e-01  7.486e-01  -0.353  0.72702
## DCA:NAC:FGF2  9.072e-02  1.059e+00   0.086  0.93243
## ---
## Signif. codes:  0 '***' 0.001 '**' 0.01 '*' 0.05 '.' 0.1 ' ' 1
##
## Residual standard error: 0.7486 on 24 degrees of freedom
## Multiple R-squared:  0.8463, Adjusted R-squared:  0.8014
## F-statistic: 18.87 on 7 and 24 DF,  p-value: 2.504e-08
```

```r
ggplot(d7D) +
  aes(x = FGF2, color = as.factor(DCA), group = as.factor(DCA), y = log2(aSMA)) +
  stat_summary(fun = mean, geom = "point") +
  stat_summary(fun = mean, geom = "line") +
  geom_point()
```

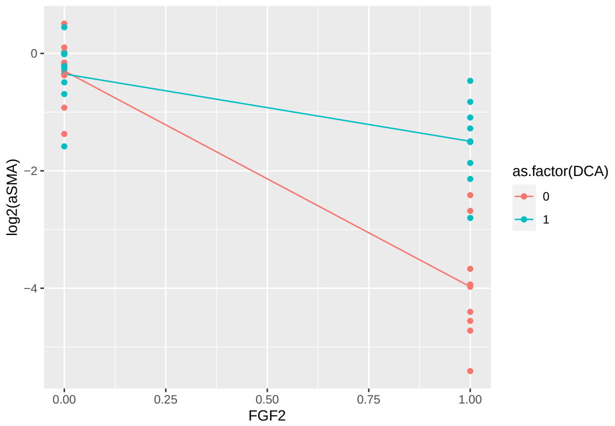

##### Analysis (Figure 7E)

```r
a7E_E2F1 <- lm(log2(E2F1)~Chemical, data=d7E)
a7E_PCNA <- lm(log2(PCNA)~Chemical, data=d7E)

summary(a7E_E2F1)
```

##### Treatment Comparisons (Figure 7E)

```
##
## Call:
## lm(formula = log2(E2F1) ~ Chemical, data = d7E)
##
## Residuals:
##      Min       1Q   Median       3Q      Max
## -0.48147 -0.15211  0.01664  0.11593  0.30658
##
## Coefficients:
##                        Estimate Std. Error t value Pr(>|t|)
## (Intercept)             -0.4999     0.1099  -4.548 0.000131 ***
## ChemicalDCA+FGF2         1.4600     0.1554   9.392 1.65e-09 ***
## ChemicalDCA+NAC         -0.2508     0.1554  -1.614 0.119678
## ChemicalDCA+NAC+FGF2     1.6854     0.1554  10.842 9.90e-11 ***
## ChemicalFGF2             2.8392     0.1554  18.265 1.39e-15 ***
## ChemicalNAC              0.2320     0.1554   1.493 0.148517
## ChemicalNAC+FGF2         2.9220     0.1554  18.797 7.28e-16 ***
## ChemicalPBS              0.4999     0.1554   3.216 0.003694 **
## ---
## Signif. codes:  0 '***' 0.001 '**' 0.01 '*' 0.05 '.' 0.1 ' ' 1
##
## Residual standard error: 0.2198 on 24 degrees of freedom
## Multiple R-squared:  0.9741, Adjusted R-squared:  0.9666
## F-statistic: 129.1 on 7 and 24 DF,  p-value: < 2.2e-16
```

```r
plot(a7E_E2F1,which=1:2)
```

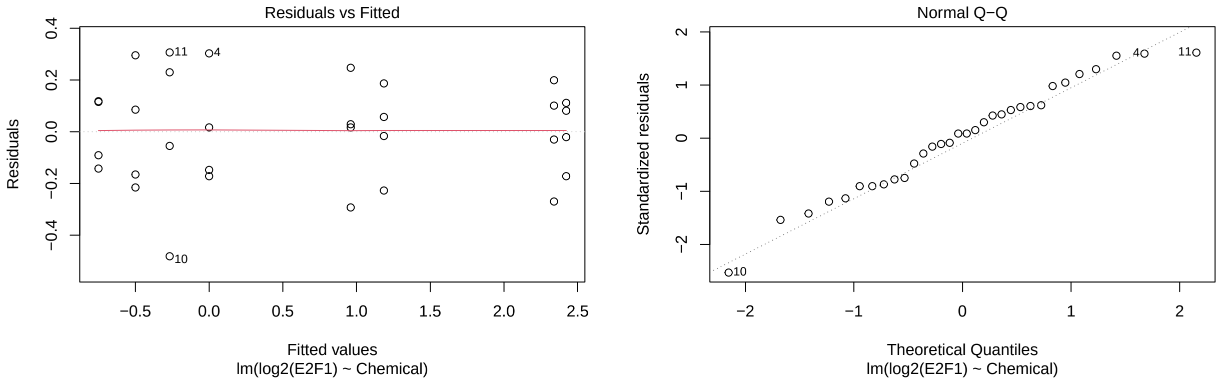

```r
plot(a7E_PCNA,which=1:2)
```

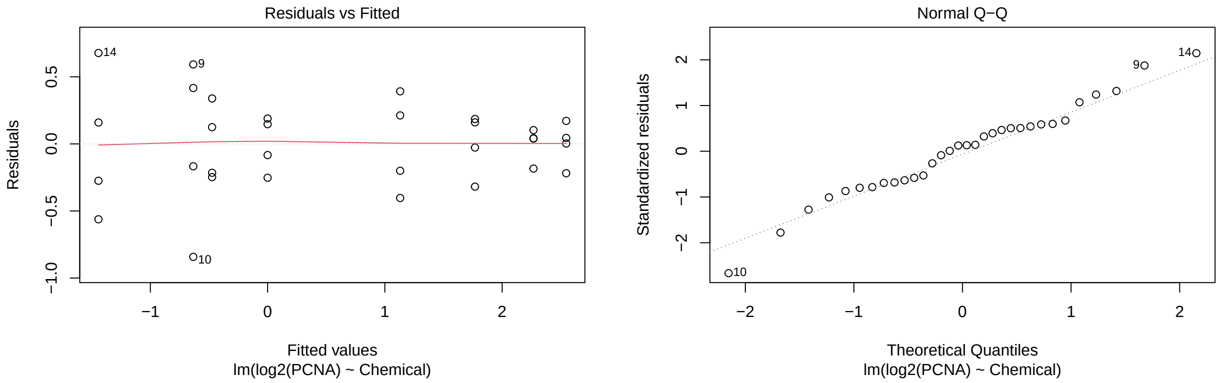

```r
K <- matrix(c(0,0,0,0,0,0,0,1, # PBS vs. DCA
              0,0,0,0,0,-1,0,1, # PBS vs. NAC
              0,0,-1,0,0,0,0,1, # PBS vs. DCA+NAC
              0,0,-1,0,0,0,0,0, # DCA vs. DCA+NAC
              0,0,0,0,-1,0,0,1, # PBS vs. FGF2
              0,-1,0,0,1,0,0,0, # FGF2 vs. FGF2+DCA
              0,0,0,0,1,0,-1,0, # FGF2 vs. FGF2+NAC
              0,0,0,-1,1,0,0,0, # FGF2 vs. FGF2+DCA+NAC
              0,1,0,-1,0,0,0,0),nrow=9, byrow=TRUE) # FGF2+DCA vs. FGF2+DCA+NAC

t7E_E2F1 <- glht(a7E_E2F1, linfct=K)
summary(t7E_E2F1, test=adjusted("fdr"))
```

```
##
##   Simultaneous Tests for General Linear Hypotheses
##
## Fit: lm(formula = log2(E2F1) ~ Chemical, data = d7E)
##
## Linear Hypotheses:
```

```
##        Estimate Std. Error t value Pr(>|t|)
## 1 == 0  0.49992    0.15544   3.216 0.006650 **
## 2 == 0  0.26787    0.15544   1.723 0.146563
## 3 == 0  0.75075    0.15544   4.830 0.000144 ***
## 4 == 0  0.25083    0.15544   1.614 0.153871
## 5 == 0 -2.33925    0.15544 -15.049 9.11e-13 ***
## 6 == 0  1.37916    0.15544   8.872 2.17e-08 ***
## 7 == 0 -0.08279    0.15544  -0.533 0.599225
## 8 == 0  1.15381    0.15544   7.423 3.47e-07 ***
## 9 == 0 -0.22535    0.15544  -1.450 0.180082
## ---
## Signif. codes:  0 '***' 0.001 '**' 0.01 '*' 0.05 '.' 0.1 ' ' 1
## (Adjusted p values reported -- fdr method)
```

```r
t7E_PCNA <- glht(a7E_PCNA, linfct=K)
summary(t7E_PCNA, test=adjusted("fdr"))
```

```
##
##   Simultaneous Tests for General Linear Hypotheses
##
## Fit: lm(formula = log2(PCNA) ~ Chemical, data = d7E)
##
## Linear Hypotheses:
##        Estimate Std. Error t value Pr(>|t|)
## 1 == 0   0.4732     0.2579   1.835  0.08882 .
## 2 == 0   0.6338     0.2579   2.457  0.02778 *
## 3 == 0   1.4426     0.2579   5.593 3.62e-05 ***
## 4 == 0   0.9693     0.2579   3.758  0.00218 **
## 5 == 0  -2.5464     0.2579  -9.873 5.68e-09 ***
## 6 == 0   0.7779     0.2579   3.016  0.01076 *
## 7 == 0   0.2789     0.2579   1.081  0.29027
## 8 == 0   1.4161     0.2579   5.490 3.62e-05 ***
## 9 == 0   0.6382     0.2579   2.475  0.02778 *
## ---
## Signif. codes:  0 '***' 0.001 '**' 0.01 '*' 0.05 '.' 0.1 ' ' 1
## (Adjusted p values reported -- fdr method)
```

```r
af7E_E2F1 <- lm(log2(E2F1)~DCA*NAC*FGF2, data=d7E)
af7E_PCNA <- lm(log2(PCNA)~DCA*NAC*FGF2, data=d7E)

summary(af7E_E2F1)
```

##### Three-way ANOVA (Figure 7E)

```
##
## Call:
## lm(formula = log2(E2F1) ~ DCA * NAC * FGF2, data = d7E)
##
## Residuals:
##      Min       1Q   Median       3Q      Max
```

```
## -0.48147 -0.15211  0.01664  0.11593  0.30658
##
## Coefficients:
##                Estimate Std. Error t value Pr(>|t|)
## (Intercept)  -1.496e-09  1.099e-01   0.000 1.000000
## DCA          -4.999e-01  1.554e-01  -3.216 0.003694 **
## NAC          -2.679e-01  1.554e-01  -1.723 0.097709 .
## FGF2          2.339e+00  1.554e-01  15.049 1.01e-13 ***
## DCA:NAC       1.703e-02  2.198e-01   0.077 0.938877
## DCA:FGF2     -8.792e-01  2.198e-01  -4.000 0.000527 ***
## NAC:FGF2      3.507e-01  2.198e-01   1.595 0.123777
## DCA:NAC:FGF2  1.255e-01  3.109e-01   0.404 0.689941
## ---
## Signif. codes:  0 '***' 0.001 '**' 0.01 '*' 0.05 '.' 0.1 ' ' 1
##
## Residual standard error: 0.2198 on 24 degrees of freedom
## Multiple R-squared:  0.9741, Adjusted R-squared:  0.9666
## F-statistic: 129.1 on 7 and 24 DF,  p-value: < 2.2e-16
```

```
ggplot(d7E) +
  aes(x = FGF2, color = as.factor(DCA), group = as.factor(DCA), y = log2(E2F1)) +
  stat_summary(fun = mean, geom = "point") +
  stat_summary(fun = mean, geom = "line") +
  geom_point()
```

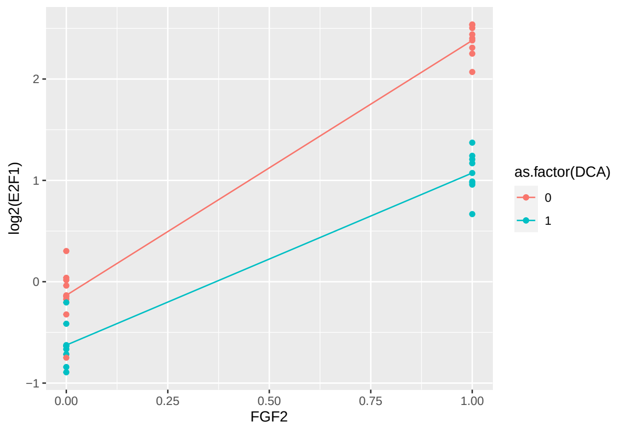

```
ggplot(d7E) +
  aes(x = DCA, color = as.factor(FGF2), group = as.factor(FGF2), y = log2(E2F1)) +
  stat_summary(fun = mean, geom = "point") +
  stat_summary(fun = mean, geom = "line") +
  geom_point()
```

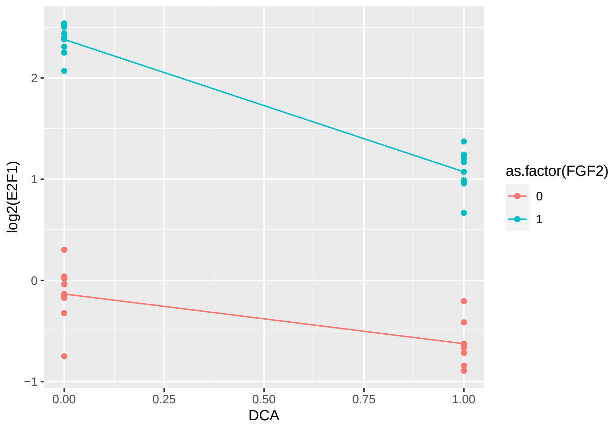

```r
summary(af7E_PCNA)
```

```
##
## Call:
## lm(formula = log2(PCNA) ~ DCA * NAC * FGF2, data = d7E)
##
## Residuals:
##      Min       1Q   Median       3Q      Max
## -0.84194 -0.21622  0.04073  0.17501  0.67736
##
## Coefficients:
##                 Estimate Std. Error t value Pr(>|t|)
## (Intercept)  -3.203e-10  1.824e-01   0.000   1.0000
## DCA          -4.732e-01  2.579e-01  -1.835   0.0790 .
## NAC          -6.338e-01  2.579e-01  -2.457   0.0216 *
## FGF2          2.546e+00  2.579e-01   9.873 6.31e-10 ***
## DCA:NAC      -3.355e-01  3.648e-01  -0.920   0.3668
## DCA:FGF2     -3.046e-01  3.648e-01  -0.835   0.4119
## NAC:FGF2      3.549e-01  3.648e-01   0.973   0.3403
## DCA:NAC:FGF2 -2.380e-02  5.158e-01  -0.046   0.9636
## ---
## Signif. codes:  0 '***' 0.001 '**' 0.01 '*' 0.05 '.' 0.1 ' ' 1
##
## Residual standard error: 0.3648 on 24 degrees of freedom
## Multiple R-squared:  0.9507, Adjusted R-squared:  0.9364
## F-statistic: 66.17 on 7 and 24 DF,  p-value: 3.826e-14
```

```r
ggplot(d7E) +
  aes(x = FGF2, y = log2(PCNA)) +
  stat_summary(fun = mean, geom = "point") +
  stat_summary(fun = mean, geom = "line") +
  geom_point()
```

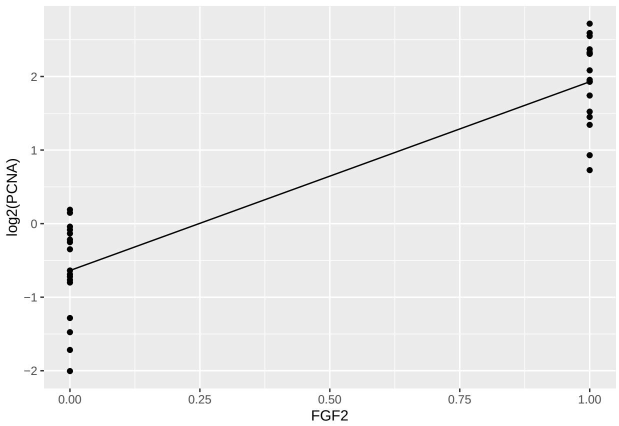

```r
ggplot(d7E) +
  aes(x = NAC, y = log2(PCNA)) +
  stat_summary(fun = mean, geom = "point") +
  stat_summary(fun = mean, geom = "line") +
  geom_point()
```

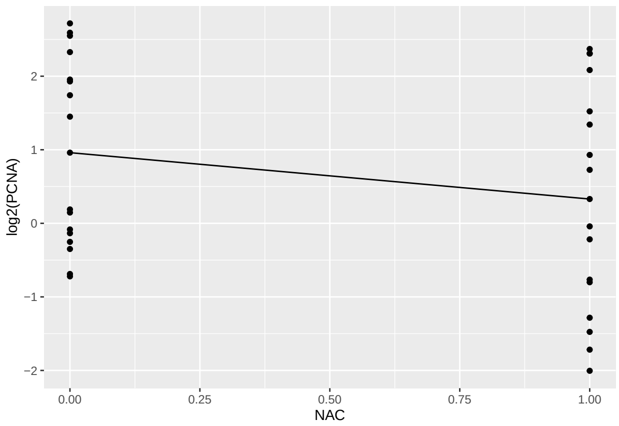

```r
ggplot(d7E) +
  aes(x = DCA, y = log2(PCNA)) +
  stat_summary(fun = mean, geom = "point") +
  stat_summary(fun = mean, geom = "line") +
  geom_point()
```

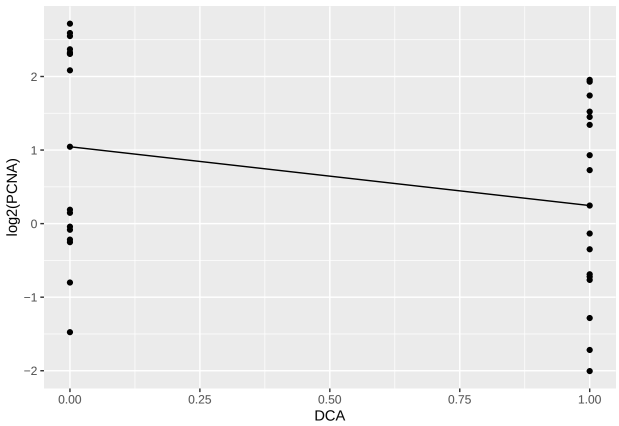

### Experiment 9 (Figure S3A)

Responses are relative expressions for several genes. There is just one Chemical factor (PBS vs. FGF2) and a Time factor (24 and 48).

Comparisons: PBS vs. FGF2 at each time point for each gene.

#### Data and Visualization

```
dS3A <- read.xlsx(xlsxFile = "MetabolismPaperRawData_Byran_working.xlsx",
                  colNames = TRUE, sheet="Fig 3S_A")
glimpse(dS3A)
```

```
## Rows: 12
## Columns: 9
## $ Chemical   <chr> "PBS", "PBS", "PBS", "FGF2", "FGF2", "FGF2", "PBS", "PBS", ~
## $ Sample     <dbl> 1, 2, 3, 1, 2, 3, 1, 2, 3, 1, 2, 3
## $ ‘Time.(h)‘ <dbl> 24, 24, 24, 24, 24, 24, 48, 48, 48, 48, 48, 48
## $ SOX2       <dbl> 99.25088, 65.32159, 92.18506, 952.79871, 619.50004, 481.604~
## $ SIX6       <dbl> 2255.023, 2176.349, 2242.821, 24150.727, 24789.424, 22717.0~
## $ PAX6       <dbl> 8205.806, 6594.370, 7890.203, 17391.511, 13764.443, 17093.4~
## $ RPE65      <dbl> 566.6905, 792.1539, 509.1130, 901.5844, 936.3166, 700.2499,~
## $ TYR        <dbl> 17588.749, 16847.786, 16566.075, 8368.197, 11459.573, 10106~
## $ OTX2       <dbl> 16101.053, 15180.530, 14995.786, 5948.323, 7796.750, 7060.8~
```

```
dS3A_long <- dS3A %>%
  rename(Time=`Time.(h)`) %>%
  pivot_longer(cols=4:9,names_to="Gene",values_to="RGE") %>%
  mutate(Time=as.factor(Time))
glimpse(dS3A_long)
```

```
## Rows: 72
## Columns: 5
## $ Chemical <chr> "PBS", "PBS", "PBS", "PBS", "PBS", "PBS", "PBS", "PBS", "PBS"~
## $ Sample   <dbl> 1, 1, 1, 1, 1, 1, 2, 2, 2, 2, 2, 2, 3, 3, 3, 3, 3, 3, 1, 1, 1~
```

```
## $ Time      <fct> 24, 24, 24, 24, 24, 24, 24, 24, 24, 24, 24, 24, 24, 24, 24, 2~
## $ Gene      <chr> "SOX2", "SIX6", "PAX6", "RPE65", "TYR", "OTX2", "SOX2", "SIX6~
## $ RGE       <dbl> 99.25088, 2255.02257, 8205.80623, 566.69048, 17588.74918, 161~
```

```
ggplot(dS3A_long, aes(x=Time,y=log2(RGE),color=Chemical)) +
  geom_jitter(width=.05,height=0) +
  facet_wrap(~Gene, nrow=3) +
  #theme(axis.text.x = element_text(angle = 90, vjust = 0.5, hjust=1)) +
  labs(title="For Figure 3SA: Time Comparisons")
```

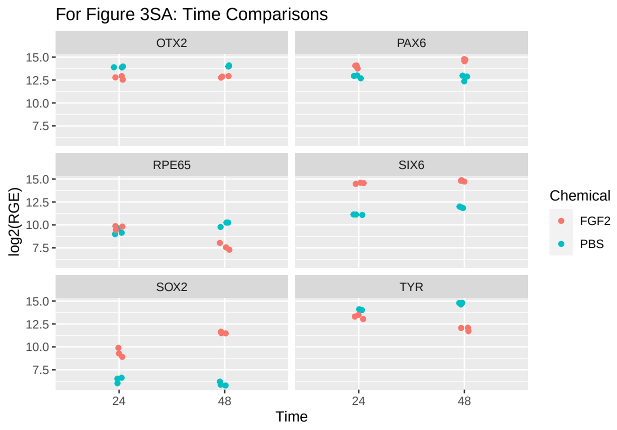

#### Data Analysis (Experiment 9)

We wish to compare PBS vs. FGF2 at each time point for each gene.

```
dS3A_2 <- dS3A %>%
  rename(Time=`Time.(h)`) %>%
  mutate(Time=as.factor(Time))
glimpse(dS3A)
```

```
## Rows: 12
## Columns: 9
## $ Chemical    <chr> "PBS", "PBS", "PBS", "FGF2", "FGF2", "FGF2", "PBS", "PBS", ~
## $ Sample      <dbl> 1, 2, 3, 1, 2, 3, 1, 2, 3, 1, 2, 3
## $ `Time.(h)`  <dbl> 24, 24, 24, 24, 24, 24, 48, 48, 48, 48, 48, 48
## $ SOX2        <dbl> 99.25088, 65.32159, 92.18506, 952.79871, 619.50004, 481.604~
## $ SIX6        <dbl> 2255.023, 2176.349, 2242.821, 24150.727, 24789.424, 22717.0~
## $ PAX6        <dbl> 8205.806, 6594.370, 7890.203, 17391.511, 13764.443, 17093.4~
## $ RPE65       <dbl> 566.6905, 792.1539, 509.1130, 901.5844, 936.3166, 700.2499,~
## $ TYR         <dbl> 17588.749, 16847.786, 16566.075, 8368.197, 11459.573, 10106~
## $ OTX2        <dbl> 16101.053, 15180.530, 14995.786, 5948.323, 7796.750, 7060.8~
```

```
aS3A_SOX2 <- aov(log2(SOX2)~Chemical*Time, data=dS3A_2)
aS3A_SIX6 <- aov(log2(SIX6)~Chemical*Time, data=dS3A_2)
aS3A_PAX6 <- aov(log2(PAX6)~Chemical*Time, data=dS3A_2)
aS3A_RPE65 <- aov(log2(RPE65)~Chemical*Time, data=dS3A_2)
aS3A_TYR <- aov(log2(TYR)~Chemical*Time, data=dS3A_2)
aS3A_OTX2 <- aov(log2(OTX2)~Chemical*Time, data=dS3A_2)
```

```r
plot(aS3A_SOX2,which=1:2)
```

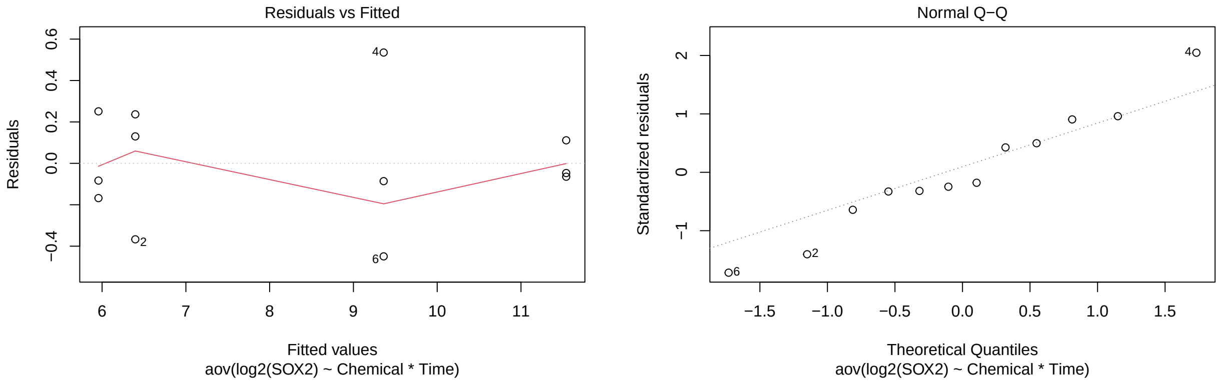

```r
plot(aS3A_SIX6,which=1:2)
```

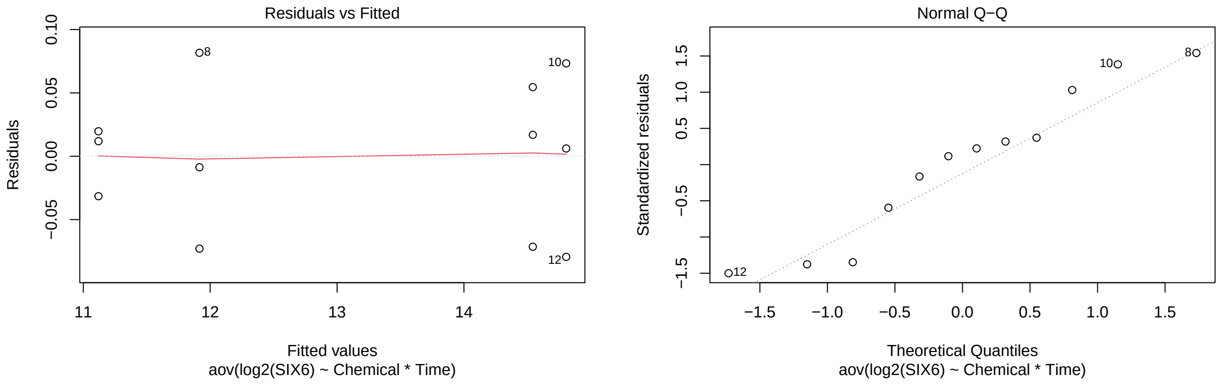

```r
plot(aS3A_PAX6,which=1:2)
```

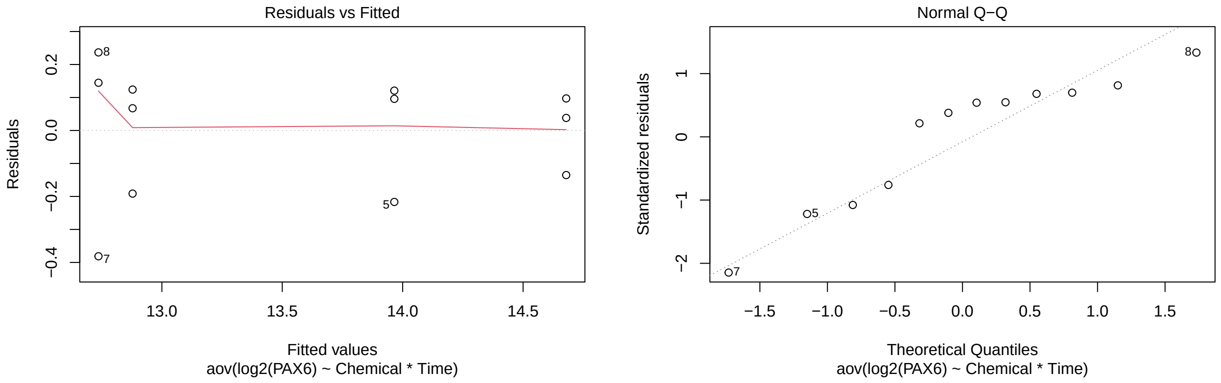

```r
plot(aS3A_RPE65,which=1:2)
```

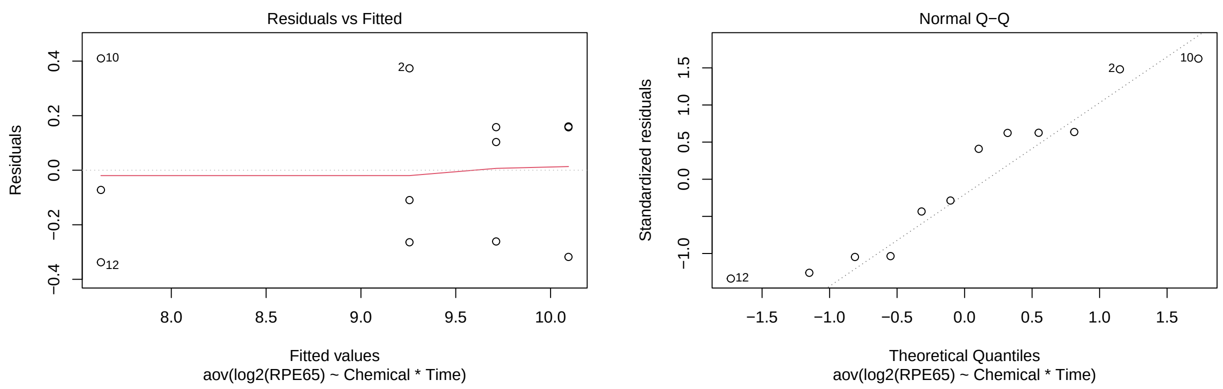

```r
plot(aS3A_TYR,which=1:2)
```

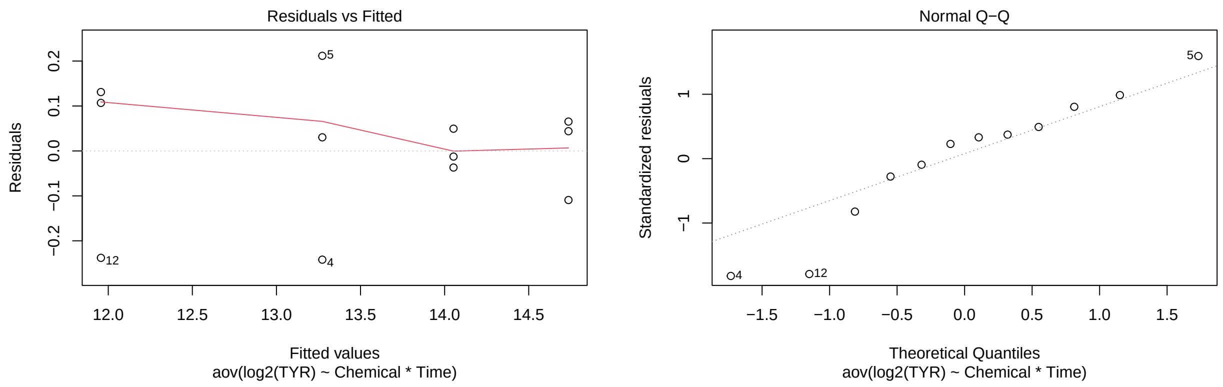

```r
plot(aS3A_OTX2,which=1:2)
```

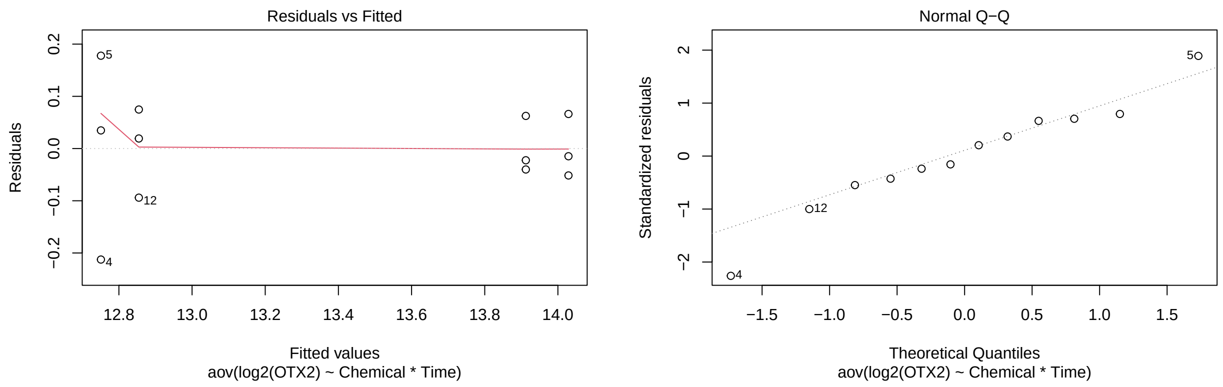

```r
aS3A_tSOX2 <- test(emmeans(aS3A_SOX2, pairwise~Chemical|Time))
aS3A_tSOX2
```

```
## $emmeans
## Time = 24:
##  Chemical emmean    SE df t.ratio p.value
##  FGF2       9.36 0.185  8  50.644  <.0001
##  PBS        6.40 0.185  8  34.605  <.0001
##
## Time = 48:
##  Chemical emmean    SE df t.ratio p.value
##  FGF2      11.54 0.185  8  62.434  <.0001
##  PBS        5.96 0.185  8  32.226  <.0001
##
## Results are given on the log2 (not the response) scale.
##
## $contrasts
## Time = 24:
##  contrast   estimate    SE df t.ratio p.value
##  FGF2 - PBS     2.96 0.261  8  11.341  <.0001
##
## Time = 48:
##  contrast   estimate    SE df t.ratio p.value
##  FGF2 - PBS     5.58 0.261  8  21.360  <.0001
##
## Results are given on the log2 (not the response) scale.
```

```
p.adjust(aS3A_tSOX2$contrasts$p.value, method="fdr")
```

```
## [1] 3.292636e-06 4.855966e-08
```

```
aS3A_tSIX6 <- test(emmeans(aS3A_SIX6, pairwise~Chemical|Time))
aS3A_tSIX6
```

```
## $emmeans
## Time = 24:
##  Chemical emmean      SE df t.ratio p.value
##  FGF2      14.54 0.03745  8 388.282  <.0001
##  PBS       11.12 0.03745  8 296.873  <.0001
##
## Time = 48:
##  Chemical emmean      SE df t.ratio p.value
##  FGF2      14.81 0.03745  8 395.311  <.0001
##  PBS       11.92 0.03745  8 318.125  <.0001
##
## Results are given on the log2 (not the response) scale.
##
## $contrasts
## Time = 24:
##  contrast   estimate    SE df t.ratio p.value
##  FGF2 - PBS     3.42 0.053  8  64.636  <.0001
##
## Time = 48:
##  contrast   estimate    SE df t.ratio p.value
##  FGF2 - PBS     2.89 0.053  8  54.578  <.0001
##
## Results are given on the log2 (not the response) scale.
```

```r
p.adjust(aS3A_tSIX6$contrasts$p.value, method="fdr")
```

```
## [1] 7.302734e-12 1.408874e-11
```

```r
aS3A_tPAX6 <- test(emmeans(aS3A_PAX6, pairwise~Chemical|Time))
aS3A_tPAX6
```

```
## $emmeans
## Time = 24:
##  Chemical emmean    SE df t.ratio p.value
##  FGF2       14.0 0.126  8 111.178  <.0001
##  PBS        12.9 0.126  8 102.526  <.0001
##
## Time = 48:
##  Chemical emmean    SE df t.ratio p.value
##  FGF2       14.7 0.126  8 116.861  <.0001
##  PBS        12.7 0.126  8 101.397  <.0001
##
## Results are given on the log2 (not the response) scale.
##
## $contrasts
## Time = 24:
##  contrast   estimate    SE df t.ratio p.value
##  FGF2 - PBS     1.09 0.178  8   6.118  0.0003
##
## Time = 48:
##  contrast   estimate    SE df t.ratio p.value
##  FGF2 - PBS     1.94 0.178  8  10.934  <.0001
##
## Results are given on the log2 (not the response) scale.
```

```r
p.adjust(aS3A_tPAX6$contrasts$p.value, method="fdr")
```

```
## [1] 2.836904e-04 8.681022e-06
```

```r
aS3A_tRPE65 <- test(emmeans(aS3A_RPE65, pairwise~Chemical|Time))
aS3A_tRPE65
```

```
## $emmeans
## Time = 24:
##  Chemical emmean    SE df t.ratio p.value
##  FGF2       9.71 0.178  8  54.441  <.0001
##  PBS        9.26 0.178  8  51.880  <.0001
##
## Time = 48:
##  Chemical emmean    SE df t.ratio p.value
##  FGF2       7.63 0.178  8  42.759  <.0001
##  PBS       10.09 0.178  8  56.579  <.0001
##
## Results are given on the log2 (not the response) scale.
##
```

```
## $contrasts
## Time = 24:
##  contrast   estimate    SE df t.ratio p.value
##  FGF2 - PBS    0.457 0.252  8   1.811  0.1077
##
## Time = 48:
##  contrast   estimate    SE df t.ratio p.value
##  FGF2 - PBS   -2.466 0.252  8  -9.772  <.0001
##
## Results are given on the log2 (not the response) scale.
```

```
p.adjust(aS3A_tRPE65$contrasts$p.value, method="fdr")
```

```
## [1] 1.076850e-01 2.015788e-05
```

```
aS3A_tTYR <- test(emmeans(aS3A_TYR, pairwise~Chemical|Time))
aS3A_tTYR
```

```
## $emmeans
## Time = 24:
##  Chemical emmean     SE df t.ratio p.value
##  FGF2       13.3 0.0939  8 141.375  <.0001
##  PBS        14.1 0.0939  8 149.686  <.0001
##
## Time = 48:
##  Chemical emmean     SE df t.ratio p.value
##  FGF2       12.0 0.0939  8 127.354  <.0001
##  PBS        14.7 0.0939  8 156.967  <.0001
##
## Results are given on the log2 (not the response) scale.
##
## $contrasts
## Time = 24:
##  contrast   estimate    SE df t.ratio p.value
##  FGF2 - PBS    -0.78 0.133  8  -5.877  0.0004
##
## Time = 48:
##  contrast   estimate    SE df t.ratio p.value
##  FGF2 - PBS    -2.78 0.133  8 -20.940  <.0001
##
## Results are given on the log2 (not the response) scale.
```

```
p.adjust(aS3A_tTYR$contrasts$p.value, method="fdr")
```

```
## [1] 3.714601e-04 5.677615e-08
```

```
aS3A_tOTX2 <- test(emmeans(aS3A_OTX2, pairwise~Chemical|Time))
aS3A_tOTX2
```

```
## $emmeans
## Time = 24:
```

```
##  Chemical emmean     SE df t.ratio p.value
##  FGF2       12.8 0.0665  8 191.775  <.0001
##  PBS        13.9 0.0665  8 209.245  <.0001
##
## Time = 48:
##  Chemical emmean     SE df t.ratio p.value
##  FGF2       12.9 0.0665  8 193.334  <.0001
##  PBS        14.0 0.0665  8 210.998  <.0001
##
## Results are given on the log2 (not the response) scale.
##
## $contrasts
## Time = 24:
##  contrast   estimate    SE df t.ratio p.value
##  FGF2 - PBS    -1.16 0.094  8 -12.353  <.0001
##
## Time = 48:
##  contrast   estimate    SE df t.ratio p.value
##  FGF2 - PBS    -1.17 0.094  8 -12.490  <.0001
##
## Results are given on the log2 (not the response) scale.
```

```
p.adjust(aS3A_tOTX2$contrasts$p.value, method="fdr")
```

```
## [1] 1.718691e-06 1.718691e-06
```

### Experiment 10 (Figure S5B)

A relative gene expression experiment with a Chemical factor (PBS, FGF2, 2DG, and FGF2+2DG) and a Time factor (24 and 48).

Comparisons: PBS vs FGF2; PBS vs 2DG; FGF2 vs FGF2+2DG, and 2DG vs FGF2+2DG at each time point for each gene.

#### Data and Visualization

```
dS5B <- read.xlsx(xlsxFile = "MetabolismPaperRawData_Byran_working.xlsx",
                  colNames = TRUE, sheet="Fig 5S_B")
glimpse(dS5B)
```

```
## Rows: 32
## Columns: 9
## $ Chemical   <chr> "PBS", "PBS", "PBS", "PBS", "FGF2", "FGF2", "FGF2", "FGF2",~
## $ Sample     <dbl> 1, 2, 3, 4, 1, 2, 3, 4, 1, 2, 3, 4, 1, 2, 3, 4, 1, 2, 3, 4,~
## $ `Time.(h)` <dbl> 24, 24, 24, 24, 24, 24, 24, 24, 24, 24, 24, 24, 24, 24, 24,~
## $ SOX2       <dbl> 1.6552989, 0.6335008, 1.0014706, 0.9522220, 2.0540540, 2.19~
## $ SIX6       <dbl> 1.0902879, 0.6457024, 1.0135222, 1.4015001, 11.4740953, 14.~
## $ PAX6       <dbl> 1.2662974, 1.0071927, 0.6287660, 1.2469892, 3.3568731, 2.48~
## $ RPE65      <dbl> 1.1581151, 0.8577921, 0.8225351, 1.2238038, 0.4842760, 0.35~
## $ TYR        <dbl> 1.1443146, 1.0239223, 0.9812298, 0.8697948, 0.5343266, 0.40~
## $ OTX2       <dbl> 1.5452385, 1.0500794, 0.6279589, 0.9814114, 0.5455059, 0.47~
```

```r
# to compare the 4 treatments at each time point
dS5B_long <- dS5B %>%
  rename(Time=`Time.(h)`) %>%
  pivot_longer(cols=4:9,names_to="Gene",values_to="RGE") %>%
  mutate(Time=as.factor(Time))
glimpse(dS5B_long)
```

```
## Rows: 192
## Columns: 5
## $ Chemical <chr> "PBS", "PBS", "PBS", "PBS", "PBS", "PBS", "PBS", "PBS", "PBS"~
## $ Sample   <dbl> 1, 1, 1, 1, 1, 1, 2, 2, 2, 2, 2, 2, 3, 3, 3, 3, 3, 3, 4, 4, 4~
## $ Time     <fct> 24, 24, 24, 24, 24, 24, 24, 24, 24, 24, 24, 24, 24, 24, 24, 2~
## $ Gene     <chr> "SOX2", "SIX6", "PAX6", "RPE65", "TYR", "OTX2", "SOX2", "SIX6~
## $ RGE      <dbl> 1.6552989, 1.0902879, 1.2662974, 1.1581151, 1.1443146, 1.5452~
```

```r
ggplot(dS5B_long, aes(x=Time,y=log2(RGE),color=Chemical)) +
  geom_jitter(width=.05,height=0) +
  facet_wrap(~Gene, nrow=3) +
  #theme(axis.text.x = element_text(angle = 90, vjust = 0.5, hjust=1)) +
  labs(title="For Figure S5B: Time Comparisons")
```

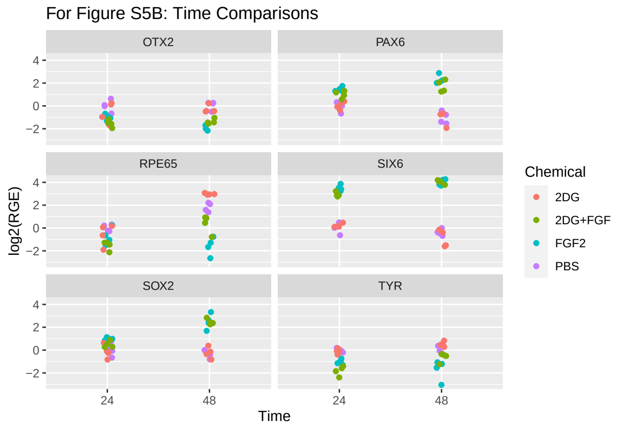

#### Data Analysis (Experiment 10)

```r
dS5B_2 <- dS5B %>%
  rename(Time=`Time.(h)`) %>%
  mutate(Time=as.factor(Time))
glimpse(dS5B_2)
```

```
## Rows: 32
## Columns: 9
## $ Chemical <chr> "PBS", "PBS", "PBS", "PBS", "FGF2", "FGF2", "FGF2", "FGF2", "~
## $ Sample   <dbl> 1, 2, 3, 4, 1, 2, 3, 4, 1, 2, 3, 4, 1, 2, 3, 4, 1, 2, 3, 4, 1~
## $ Time     <fct> 24, 24, 24, 24, 24, 24, 24, 24, 24, 24, 24, 24, 24, 24, 24, 2~
## $ SOX2     <dbl> 1.6552989, 0.6335008, 1.0014706, 0.9522220, 2.0540540, 2.1914~
## $ SIX6     <dbl> 1.0902879, 0.6457024, 1.0135222, 1.4015001, 11.4740953, 14.57~
```

```
## $ PAX6     <dbl> 1.2662974, 1.0071927, 0.6287660, 1.2469892, 3.3568731, 2.4835~
## $ RPE65    <dbl> 1.1581151, 0.8577921, 0.8225351, 1.2238038, 0.4842760, 0.3518~
## $ TYR      <dbl> 1.1443146, 1.0239223, 0.9812298, 0.8697948, 0.5343266, 0.4056~
## $ OTX2     <dbl> 1.5452385, 1.0500794, 0.6279589, 0.9814114, 0.5455059, 0.4789~
```

```
aS5B_SOX2 <- aov(log2(SOX2)~Chemical*Time, data=dS5B_2)
aS5B_SIX6 <- aov(log2(SIX6)~Chemical*Time, data=dS5B_2)
aS5B_PAX6 <- aov(log2(PAX6)~Chemical*Time, data=dS5B_2)
aS5B_RPE65 <- aov(log2(RPE65)~Chemical*Time, data=dS5B_2)
aS5B_TYR <- aov(log2(TYR)~Chemical*Time, data=dS5B_2)
aS5B_OTX2 <- aov(log2(OTX2)~Chemical*Time, data=dS5B_2)

plot(aS5B_SOX2,which=1:2)
```

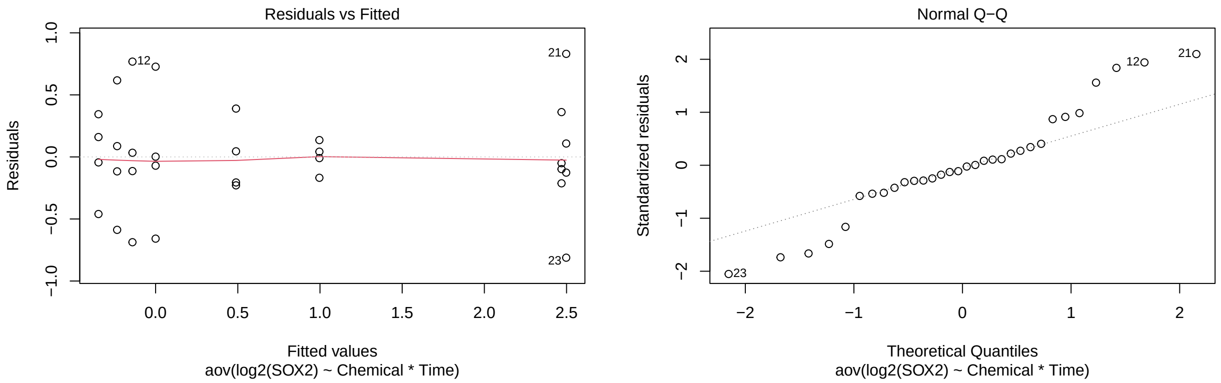

```
plot(aS5B_SIX6,which=1:2)
```

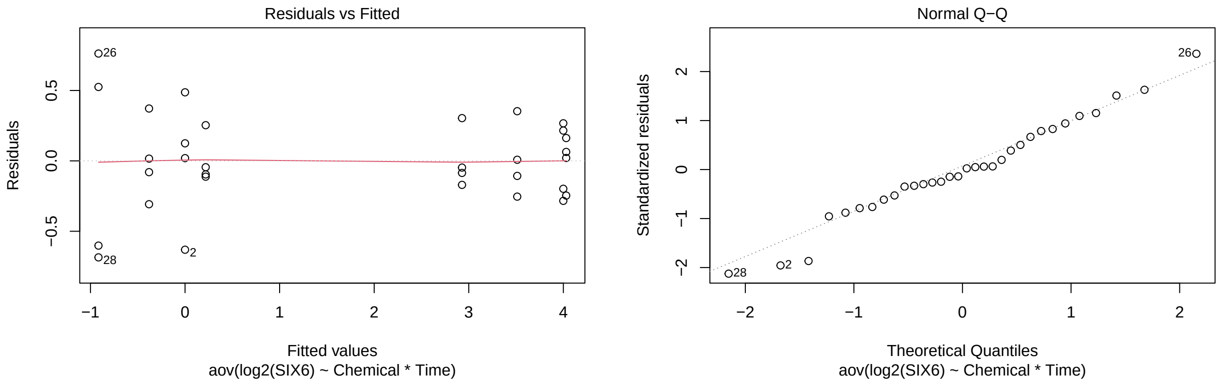

```
plot(aS5B_PAX6,which=1:2)
```

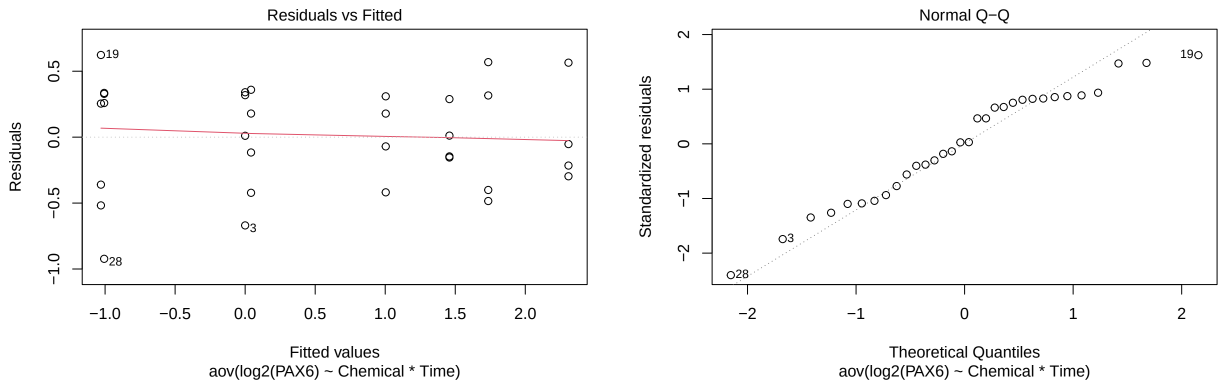

```r
plot(aS5B_RPE65,which=1:2)
```

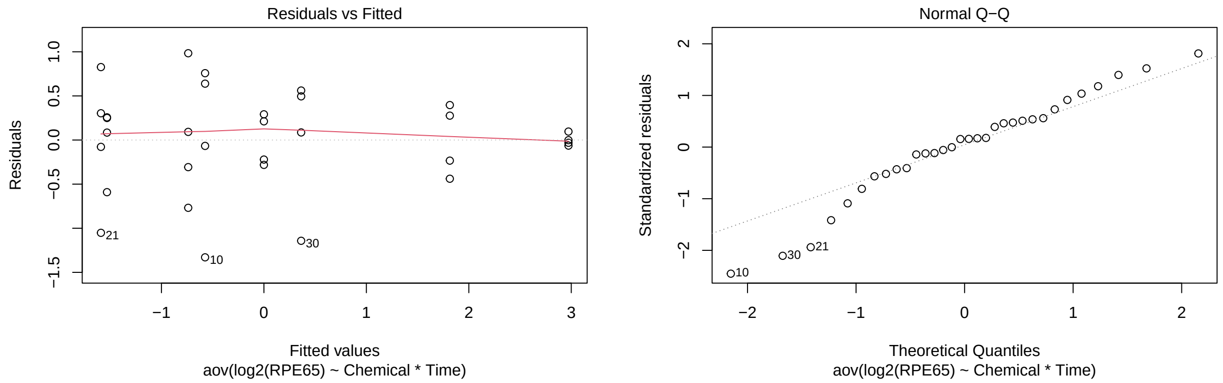

```r
plot(aS5B_TYR,which=1:2)
```

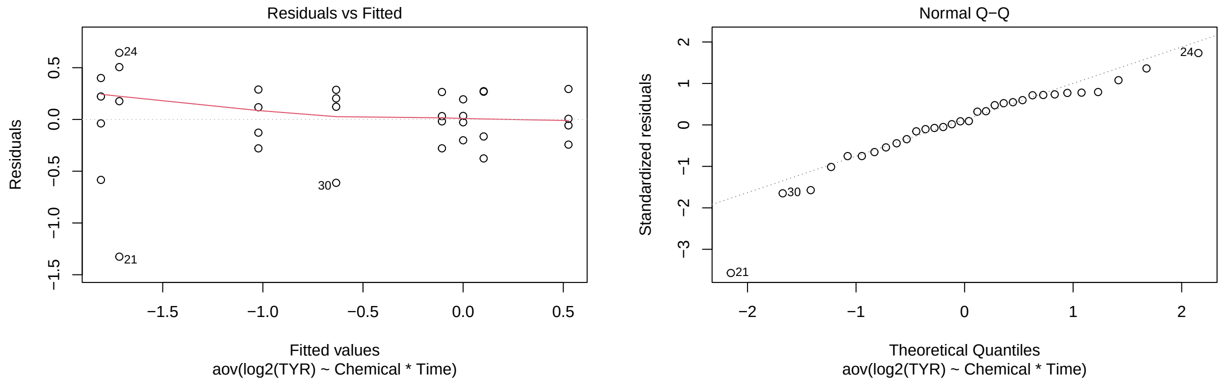

```r
plot(aS5B_OTX2,which=1:2)
```

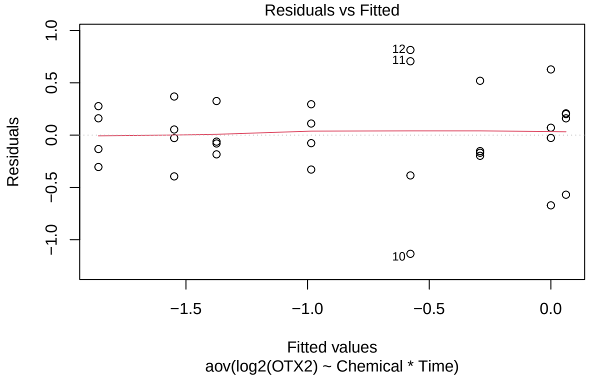

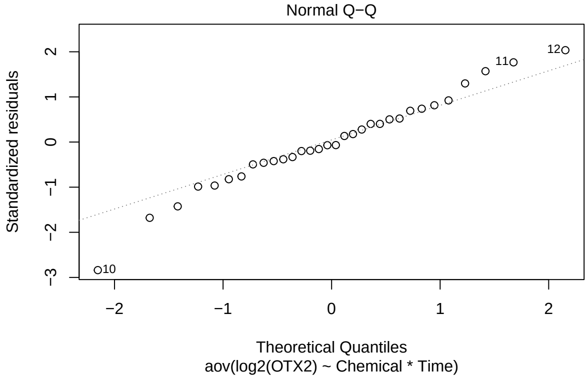

```r
aS5B_tSOX2 <- test(emmeans(aS5B_SOX2, pairwise~Chemical|Time), adjust="none")
aS5B_tSOX2
```

```
## $emmeans
## Time = 24:
##  Chemical emmean    SE df t.ratio p.value
##  2DG      -0.141 0.228 24  -0.617  0.5430
##  2DG+FGF   0.489 0.228 24   2.141  0.0426
##  FGF2      0.996 0.228 24   4.361  0.0002
##  PBS       0.000 0.228 24   0.000  1.0000
##
## Time = 48:
##  Chemical emmean    SE df t.ratio p.value
##  2DG      -0.234 0.228 24  -1.024  0.3159
##  2DG+FGF   2.469 0.228 24  10.810  <.0001
##  FGF2      2.498 0.228 24  10.934  <.0001
##  PBS      -0.348 0.228 24  -1.522  0.1411
##
## Results are given on the log2 (not the response) scale.
##
## $contrasts
## Time = 24:
##  contrast          estimate    SE df t.ratio p.value
##  2DG - (2DG+FGF)    -0.6301 0.323 24  -1.950  0.0629
##  2DG - FGF2         -1.1373 0.323 24  -3.520  0.0018
##  2DG - PBS          -0.1410 0.323 24  -0.436  0.6665
##  (2DG+FGF) - FGF2  -0.5072 0.323 24  -1.570  0.1295
##  (2DG+FGF) - PBS     0.4892 0.323 24   1.514  0.1431
##  FGF2 - PBS          0.9963 0.323 24   3.084  0.0051
##
## Time = 48:
##  contrast          estimate    SE df t.ratio p.value
##  2DG - (2DG+FGF)    -2.7034 0.323 24  -8.368  <.0001
##  2DG - FGF2         -2.7319 0.323 24  -8.456  <.0001
##  2DG - PBS           0.1137 0.323 24   0.352  0.7280
##  (2DG+FGF) - FGF2  -0.0285 0.323 24  -0.088  0.9305
##  (2DG+FGF) - PBS     2.8171 0.323 24   8.720  <.0001
##  FGF2 - PBS          2.8456 0.323 24   8.808  <.0001
##
```

```
## Results are given on the log2 (not the response) scale.
```

```r
p.adjust(aS5B_tSOX2$contrasts$p.value[c(1,3,4,6,7,9,10,12)], method="fdr")
```

```
## [1] 1.257955e-01 8.320515e-01 2.072755e-01 1.354528e-02 5.644461e-08
## [6] 8.320515e-01 9.304616e-01 4.413643e-08
```

```r
aS5B_tSIX6 <- test(emmeans(aS5B_SIX6, pairwise~Chemical|Time), adjust="none")
aS5B_tSIX6
```

```
## $emmeans
## Time = 24:
##  Chemical emmean    SE df t.ratio p.value
##  2DG       0.218 0.186 24   1.173  0.2524
##  2DG+FGF   2.929 0.186 24  15.729  <.0001
##  FGF2      3.512 0.186 24  18.864  <.0001
##  PBS       0.000 0.186 24   0.000  1.0000
##
## Time = 48:
##  Chemical emmean    SE df t.ratio p.value
##  2DG      -0.915 0.186 24  -4.916  0.0001
##  2DG+FGF   4.030 0.186 24  21.645  <.0001
##  FGF2      3.999 0.186 24  21.480  <.0001
##  PBS      -0.380 0.186 24  -2.044  0.0521
##
## Results are given on the log2 (not the response) scale.
##
## $contrasts
## Time = 24:
##  contrast          estimate    SE df t.ratio p.value
##  2DG - (2DG+FGF)    -2.7103 0.263 24 -10.293  <.0001
##  2DG - FGF2         -3.2941 0.263 24 -12.510  <.0001
##  2DG - PBS           0.2184 0.263 24   0.829  0.4151
##  (2DG+FGF) - FGF2   -0.5838 0.263 24  -2.217  0.0363
##  (2DG+FGF) - PBS     2.9286 0.263 24  11.122  <.0001
##  FGF2 - PBS          3.5124 0.263 24  13.339  <.0001
##
## Time = 48:
##  contrast          estimate    SE df t.ratio p.value
##  2DG - (2DG+FGF)    -4.9456 0.263 24 -18.782  <.0001
##  2DG - FGF2         -4.9149 0.263 24 -18.665  <.0001
##  2DG - PBS          -0.5349 0.263 24  -2.031  0.0534
##  (2DG+FGF) - FGF2    0.0307 0.263 24   0.117  0.9082
##  (2DG+FGF) - PBS     4.4106 0.263 24  16.750  <.0001
##  FGF2 - PBS          4.3800 0.263 24  16.634  <.0001
##
## Results are given on the log2 (not the response) scale.
```

```r
p.adjust(aS5B_tSIX6$contrasts$p.value[c(1,3,4,6,7,9,10,12)], method="fdr")
```

```
## [1] 5.578304e-10 4.744248e-01 5.815076e-02 3.618948e-12 5.934725e-15
## [6] 7.123283e-02 9.082017e-01 4.479521e-14
```

```r
aS5B_tPAX6 <- test(emmeans(aS5B_PAX6, pairwise~Chemical|Time), adjust="none")
aS5B_tPAX6
```

```
## $emmeans
## Time = 24:
##  Chemical  emmean    SE df t.ratio p.value
##  2DG       0.0421 0.222 24   0.190  0.8511
##  2DG+FGF   1.0038 0.222 24   4.527  0.0001
##  FGF2      1.4584 0.222 24   6.578  <.0001
##  PBS       0.0000 0.222 24   0.000  1.0000
##
## Time = 48:
##  Chemical  emmean    SE df t.ratio p.value
##  2DG      -1.0066 0.222 24  -4.540  0.0001
##  2DG+FGF   1.7355 0.222 24   7.827  <.0001
##  FGF2      2.3083 0.222 24  10.411  <.0001
##  PBS      -1.0297 0.222 24  -4.644  0.0001
##
## Results are given on the log2 (not the response) scale.
##
## $contrasts
## Time = 24:
##  contrast          estimate    SE df t.ratio p.value
##  2DG - (2DG+FGF)    -0.9617 0.314 24  -3.067  0.0053
##  2DG - FGF2         -1.4164 0.314 24  -4.517  0.0001
##  2DG - PBS           0.0421 0.314 24   0.134  0.8944
##  (2DG+FGF) - FGF2  -0.4547 0.314 24  -1.450  0.1600
##  (2DG+FGF) - PBS    1.0038 0.314 24   3.201  0.0038
##  FGF2 - PBS          1.4584 0.314 24   4.651  0.0001
##
## Time = 48:
##  contrast          estimate    SE df t.ratio p.value
##  2DG - (2DG+FGF)    -2.7421 0.314 24  -8.745  <.0001
##  2DG - FGF2         -3.3149 0.314 24 -10.572  <.0001
##  2DG - PBS           0.0231 0.314 24   0.074  0.9418
##  (2DG+FGF) - FGF2  -0.5728 0.314 24  -1.827  0.0802
##  (2DG+FGF) - PBS    2.7652 0.314 24   8.819  <.0001
##  FGF2 - PBS          3.3380 0.314 24  10.645  <.0001
##
## Results are given on the log2 (not the response) scale.
```

```r
p.adjust(aS5B_tPAX6$contrasts$p.value[c(1,3,4,6,7,9,10,12)], method="fdr")
```

```
## [1] 1.057989e-02 9.418154e-01 2.133602e-01 2.689129e-04 2.521021e-08
## [6] 9.418154e-01 1.283197e-01 1.143616e-09
```

```r
aS5B_tRPE65 <- test(emmeans(aS5B_RPE65, pairwise~Chemical|Time), adjust="none")
aS5B_tRPE65
```

```
## $emmeans
## Time = 24:
##  Chemical emmean    SE df t.ratio p.value
```

```
##   2DG      -0.574 0.313 24  -1.833  0.0792
##   2DG+FGF  -1.532 0.313 24  -4.890  0.0001
##   FGF2     -0.739 0.313 24  -2.359  0.0268
##   PBS       0.000 0.313 24   0.000  1.0000
##
## Time = 48:
##   Chemical emmean    SE df t.ratio p.value
##   2DG       2.973 0.313 24   9.492  <.0001
##   2DG+FGF   0.364 0.313 24   1.161  0.2571
##   FGF2     -1.591 0.313 24  -5.080  <.0001
##   PBS       1.815 0.313 24   5.795  <.0001
##
## Results are given on the log2 (not the response) scale.
##
## $contrasts
## Time = 24:
##   contrast          estimate    SE df t.ratio p.value
##   2DG - (2DG+FGF)      0.957 0.443 24   2.162  0.0408
##   2DG - FGF2           0.165 0.443 24   0.372  0.7134
##   2DG - PBS           -0.574 0.443 24  -1.296  0.2072
##   (2DG+FGF) - FGF2    -0.793 0.443 24  -1.790  0.0861
##   (2DG+FGF) - PBS     -1.532 0.443 24  -3.458  0.0020
##   FGF2 - PBS          -0.739 0.443 24  -1.668  0.1083
##
## Time = 48:
##   contrast          estimate    SE df t.ratio p.value
##   2DG - (2DG+FGF)      2.609 0.443 24   5.891  <.0001
##   2DG - FGF2           4.563 0.443 24  10.304  <.0001
##   2DG - PBS            1.158 0.443 24   2.615  0.0152
##   (2DG+FGF) - FGF2     1.954 0.443 24   4.413  0.0002
##   (2DG+FGF) - PBS     -1.451 0.443 24  -3.277  0.0032
##   FGF2 - PBS          -3.406 0.443 24  -7.690  <.0001
##
## Results are given on the log2 (not the response) scale.
```

```
p.adjust(aS5B_tRPE65$contrasts$p.value[c(1,3,4,6,7,9,10,12)], method="fdr")
```

```
## [1] 6.531428e-02 2.072171e-01 1.147426e-01 1.237979e-01 1.781365e-05
## [6] 3.038937e-02 4.930685e-04 5.050255e-07
```

```
aS5B_tTYR <- test(emmeans(aS5B_TYR, pairwise~Chemical|Time), adjust="none")
aS5B_tTYR
```

```
## $emmeans
## Time = 24:
##   Chemical emmean    SE df t.ratio p.value
##   2DG      -0.106 0.214 24  -0.494  0.6258
##   2DG+FGF  -1.808 0.214 24  -8.440  <.0001
##   FGF2     -1.022 0.214 24  -4.771  0.0001
##   PBS       0.000 0.214 24   0.000  1.0000
##
## Time = 48:
##   Chemical emmean    SE df t.ratio p.value
```

```
##  2DG       0.526 0.214 24   2.454  0.0218
##  2DG+FGF  -0.634 0.214 24  -2.959  0.0068
##  FGF2     -1.717 0.214 24  -8.011  <.0001
##  PBS       0.102 0.214 24   0.478  0.6369
##
## Results are given on the log2 (not the response) scale.
##
## $contrasts
## Time = 24:
##  contrast          estimate    SE df t.ratio p.value
##  2DG - (2DG+FGF)      1.703 0.303 24   5.618  <.0001
##  2DG - FGF2           0.917 0.303 24   3.024  0.0059
##  2DG - PBS           -0.106 0.303 24  -0.349  0.7299
##  (2DG+FGF) - FGF2    -0.786 0.303 24  -2.594  0.0159
##  (2DG+FGF) - PBS     -1.808 0.303 24  -5.968  <.0001
##  FGF2 - PBS          -1.022 0.303 24  -3.374  0.0025
##
## Time = 48:
##  contrast          estimate    SE df t.ratio p.value
##  2DG - (2DG+FGF)      1.160 0.303 24   3.827  0.0008
##  2DG - FGF2           2.243 0.303 24   7.400  <.0001
##  2DG - PBS            0.423 0.303 24   1.397  0.1751
##  (2DG+FGF) - FGF2     1.083 0.303 24   3.573  0.0015
##  (2DG+FGF) - PBS     -0.736 0.303 24  -2.430  0.0229
##  FGF2 - PBS          -1.819 0.303 24  -6.003  <.0001
##
## Results are given on the log2 (not the response) scale.
```

```r
p.adjust(aS5B_tTYR$contrasts$p.value[c(1,3,4,6,7,9,10,12)], method="fdr")
```

```
## [1] 3.507876e-05 7.299165e-01 2.122324e-02 4.024572e-03 2.171020e-03
## [6] 2.001695e-01 3.078087e-03 2.707399e-05
```

```r
aS5B_tOTX2 <- test(emmeans(aS5B_OTX2, pairwise~Chemical|Time), adjust="none")
aS5B_tOTX2
```

```
## $emmeans
## Time = 24:
##  Chemical  emmean    SE df t.ratio p.value
##  2DG      -0.5773 0.231 24  -2.501  0.0196
##  2DG+FGF  -1.5483 0.231 24  -6.708  <.0001
##  FGF2     -0.9852 0.231 24  -4.268  0.0003
##  PBS       0.0000 0.231 24   0.000  1.0000
##
## Time = 48:
##  Chemical  emmean    SE df t.ratio p.value
##  2DG      -0.2910 0.231 24  -1.261  0.2195
##  2DG+FGF  -1.3742 0.231 24  -5.954  <.0001
##  FGF2     -1.8598 0.231 24  -8.057  <.0001
##  PBS       0.0621 0.231 24   0.269  0.7903
##
## Results are given on the log2 (not the response) scale.
##
```

```
## $contrasts
## Time = 24:
##  contrast          estimate    SE df t.ratio p.value
##  2DG - (2DG+FGF)      0.971 0.326 24   2.975  0.0066
##  2DG - FGF2           0.408 0.326 24   1.249  0.2235
##  2DG - PBS           -0.577 0.326 24  -1.769  0.0897
##  (2DG+FGF) - FGF2    -0.563 0.326 24  -1.725  0.0974
##  (2DG+FGF) - PBS     -1.548 0.326 24  -4.743  0.0001
##  FGF2 - PBS          -0.985 0.326 24  -3.018  0.0059
##
## Time = 48:
##  contrast          estimate    SE df t.ratio p.value
##  2DG - (2DG+FGF)      1.083 0.326 24   3.319  0.0029
##  2DG - FGF2           1.569 0.326 24   4.806  0.0001
##  2DG - PBS           -0.353 0.326 24  -1.082  0.2902
##  (2DG+FGF) - FGF2     0.486 0.326 24   1.487  0.1499
##  (2DG+FGF) - PBS     -1.436 0.326 24  -4.400  0.0002
##  FGF2 - PBS          -1.922 0.326 24  -5.888  <.0001
##
## Results are given on the log2 (not the response) scale.
```

```
p.adjust(aS5B_tOTX2$contrasts$p.value[c(1,3,4,6,7,9,10,12)], method="fdr")
```

```
## [1] 1.318462e-02 1.298310e-01 1.298310e-01 1.318462e-02 1.151489e-02
## [6] 2.901841e-01 1.713212e-01 3.595896e-05
```
